# Localized Reactivity on Proteins as Riemannian Manifolds: A Quantum-Inspired Geometric Model for Deterministic, Metal-Aware Reactive-Site Prediction

**DOI:** 10.1101/2025.04.29.651260

**Authors:** Haneul Park

## Abstract

We introduce a unified framework for analysing molecular reactivity based on a geometric, quantum-inspired environment representation and a fully deterministic, metal-aware implementation. Proteins and ribonucleoprotein complexes are treated as configurations in ℝ^3^ × *T*, and each residue or nucleotide *p* is mapped to an environment vector *E*_*p*_ that encodes a coarse-grained, DFT-inspired density surrogate together with metal/phosphate fields, solvent exposure and local geometry.

A block-streamed, GPU-optional Python pipeline maps arbitrary PDB/mmCIF structures to fixed-dimensional environment vectors without stochastic training and scales to supramolecular assemblies: the 6Q97 tmRNA–SmpB–ribosome rescue complex (11,618 residues) can be processed in a single pass on commodity cloud hardware, demonstrating practical feasibility at ribosome scale. In a strict unbound, zero-shot setting on the Docking Benchmark 5.5 (DB5.5), a simple classifier trained on top of *E*_*p*_ achieves a macro-averaged area under the precision–recall curve of ~0.53 and a ROC–AUC of ~0.86 for residue-level interface vs. non-interface classification, competitive with specialised interface-prediction architectures despite using no evolutionary profiles, MSAs or task-specific retraining on DB5.5.

Across mechanistically curated case studies (Rubisco, GroEL/GroES, SecA, p53– DNA and ribosomal pockets), untuned Random Forests used purely as probes under site-grouped cross-validation yield ROC–AUC values exceeding 0.95 for catalytic and anchor cores (e.g., SecA ATPase, GroES IVL), while diffuse regulatory and fitness-defined labels are substantially harder to separate. For 6Q97, a Tier 1/Tier 2 labelling scheme over tmRNA/SmpB pockets, decoding-centre rRNA, the 23S peptidyl transferase centre and helicase-like uS3/uS4/uS5 pockets, together with a curated hard-negative panel of 323 buried hydrophobic, electrostatic and stacking decoys, yields global AUCs of ~0.94 (Tier 1+2 vs. all) and ~0.98 (Tier 1+2 vs. hard negatives). These results support the view that the environment representation defines an interpretable “reactivity manifold” in which genuinely functional pockets occupy regions that cannot be mimicked by generic dense or charged environments, and that this structure remains accessible even for full ribosomes on modest hardware.

## 1 Introduction

High-throughput structure determination and structural genomics have produced an abundance of experimentally resolved and predicted protein structures[1, 2], but *functional* annotation still lags far behind[3]. Identifying catalytic, allosteric, metal-coordination, or regulatory residues typically requires extensive prior knowledge, labour-intensive mutagenesis, or expensive QM/MM simulations focused on a small predefined region. In parallel, geometric and graph-based pipelines are increasingly able to highlight metal-binding residues, pockets, and interfaces from local descriptors[4, 5, 6, 7], yet non-deterministic learning stages and complex software stacks can impede reproducibility and make it difficult to separate genuine physical signal from model-specific artefacts. There is thus a need for approaches that combine physical interpretability, deterministic behaviour, and computational efficiency, while still exposing enough structure for downstream predictors to exploit.

In this work we take a representation-first perspective. We regard proteins, protein complexes, and (in principle) their conformational changes as objects embedded in a common ambient space ℝ ^3^ ×*T*, where ℝ^3^ denotes physical space and *T* indexes time or external conditions such as ligand state. A single static structure can be viewed as a finite sample from a submanifold of ℝ^3^ ×*T*, an MD trajectory traces out a curve or thin tube, and a set of interacting proteins corresponds to a collection of such submanifolds that intersect or approach each other in this space. Our goal is to equip this geometric picture with a basic, physically motivated notion of local “reactivity” that is defined uniformly for all residues and configurations, so that intra-protein organisation and inter-protein contacts can be analysed within a single coordinate system. In this paper we instantiate the framework on amino-acid residues in proteins, but the construction is intended to extend to more general molecular systems at the level of atoms or functional groups.

### Position relative to existing work

#### Structure-based local environments and functional sites

A long line of work has used local structural environments to annotate functional residues. Early methods such as FEATURE and related environment-vector approaches represent a residue-centred neighbourhood by hand-crafted physico-chemical descriptors and then transfer annotations by matching similar environments across structures[8]. Subsequent tools, including pocket- and cleft-based pipelines such as ConCavity and related geometry–conservation hybrids, treat local surface patches as the basic unit and combine shape with sequence conservation to detect catalytic or binding sites[5, 9]. More recent deep-learning methods derive local representations from equivariant 3D networks or atomistic foundation models, using learned embeddings of residue-centred environments as powerful but opaque features for tasks such as chemical-shift prediction or binding-site classification[6, 10]. These approaches demonstrate that local structure carries rich functional signal, but the representations are typically task-optimised, model-specific, and not designed as a reusable, deterministic coordinate system for reactivity.

#### Electron-density and conceptual DFT descriptors of reactivity

From the quantum chemistry side, conceptual Density Functional Theory (DFT) and electron-density analyses have established ground-state density and its derivatives as fundamental descriptors of molecular stability and reactivity[11]. In the enzymology literature, several studies have used charge density, Laplacians of the electron density, and Fukui-type indices at key atoms to rationalise catalytic activation and electrostatic preorganisation at specific active sites, typically using QM or QM/MM calculations in a small predefined region[12]. These works provide physically grounded descriptors of local reactivity, but they are computationally expensive and have not been developed into a deterministic, per-site representation that can be applied systematically across entire proteins or large panels of proteins.

#### Geometric and topological representations of protein structure

A third line of work treats proteins and molecular assemblies as geometric or topological objects. Topological data analysis (TDA) pipelines such as SINATRA Pro[13] and graph-based methods like PersGNN summarise ensembles of protein conformations with persistent homology[14] or related signatures, and then use these summaries in statistical or deep models to distinguish functional states or binding modes. More recent studies interpret protein conformational space as a Riemannian manifold equipped with a metric derived from physical energy landscapes, enabling the analysis of global dynamics and nonlinear trajectories[15]. While these approaches successfully capture macroscopic geometric evolution, translating such global dynamic descriptions into explicitly interpretable, density-inspired coordinates for the local reactivity of individual residues remains an open challenge.

#### Metal- and ligand-focused electron-density tools

Several specialised methods focus on metal ions and small-molecule ligands using experimental electron-density maps. Map-validation tools classify local density blobs as likely ligands or ions and flag inconsistencies between model and map[16], and more recent frameworks explicitly predict the identity and localisation of metal ions and waters in structural data[17, 18]. These methods operate on local electron density and its neighbourhood, but they are tailored to specific species (ligands, metals, waters) rather than providing a unified representation of residue-level reactivity across catalytic, binding, and regulatory roles.

#### Position of the present work

The present framework sits at the intersection of these lines. Conceptually, we adopt the DFT-inspired view that electronic structure is encoded in electron density and external fields, but we work at the level of *surrogate* density fields and coarse-grained descriptors that can be computed deterministically from standard structural data. Geometrically, we treat proteins and complexes as configurations embedded in ℝ^3^ ×*T* and use the resulting environment vectors to turn each configuration into a finite metric–measure space in a common “reactivity” coordinate system. Practically, we provide a single, fully deterministic pipeline that assigns to every residue in a structure a fixed-dimensional environment vector, without any stochastic training or task-specific tuning.

Unlike task-optimised site predictors, our focus is not on squeezing out state-of-the-art performance for a particular endpoint, but on quantifying how much functional structure— catalytic, metal-coordination, DNA-contact, protein–protein interface, allosteric, and posttranslational modification sites—is already accessible from this unified representation when probed by shallow classifiers. Concretely, we represent the C*α* trace as a one-dimensional curve embedded in ℝ^3^ × *T* and endow a tubular neighbourhood with an induced surface metric. Around each residue *p* we define a local neighbourhood *U*_*p*_ and construct an *environment vector* Ep_*p*_ ∈ ℝ^*k*^ that aggregates coarse-grained information about local charge, surrogate density, curvature, solvent exposure, and residue identity, together with a small set of neighbour-based geometric moments and field terms. This design is inspired by the Hohenberg–Kohn principle, which states that the ground-state electron density determines electronic structure: instead of attempting to reconstruct the full density field, we view Ep_*p*_ as a low-dimensional functional of a hypothetical local density and its external-field couplings that is rich enough to discriminate reactive from inert environments. Formally, the map

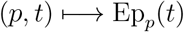

from configuration space (and, in principle, time) to environment space induces an embedded environment-vector manifold with a well-defined metric and a sub-Gaussian reference measure, so that distances, expectations, and kernels in environment space are mathematically well posed.

On top of this representation we provide a fully deterministic, metal-aware implementation for structural snapshots. Given a PDB or mmCIF file, our pipeline constructs for each residue a fixed-dimensional environment vector by combining density-inspired terms from a promolecular surrogate density, metal- and phosphate-like field components obtained by integrating over atom-centred kernels, solvent-accessible surface area, curvature-based quantities, and simple neighbour-geometry summaries, optionally augmented by residue-identity embeddings. The implementation depends only on standard scientific Python libraries and contains no data-driven fitting stage, so that identical inputs produce identical environment vectors under a fixed software environment.

The core question of this paper is therefore not whether a particular machine-learning architecture can achieve state-of-the-art performance, but rather *how much functional structure is already exposed* by this environment representation. To probe this, we treat the environment vectors as features and use an off-the-shelf Random Forest classifier as a shallow, untuned probe. For each protein we curate Tier-1 and Tier-2 sets of functional residues— including catalytic, metal-coordination, allosteric, and protein–protein interface sites—and evaluate residue-level separation against a background of non-annotated positions. Because many functional sites appear in multiple symmetry-related copies, we employ a site-grouped cross-validation protocol based on `GroupKFold`, in which groups are defined at the level of residue numbers and structural roles, so that no copy of a functional site appears in both training and test folds. The resulting ROC–AUC values characterise how linearly or weakly non-linearly separable the curated functional sets are in environment space, independently of any heavy model tuning.

Conceptually, this work makes three contributions:

1. **A geometric and measure-theoretic formalisation of residue environments.** We model the collection of residue-wise environment vectors as an embedded manifold equipped with an induced metric and a sub-Gaussian reference measure. This provides a common representation space in which different scoring functions or, in future, learning algorithms can operate, and clarifies how Euclidean, density-overlap, and geodesic kernels on environment space are related.
2. **A deterministic, metal-aware implementation for structural configurations.** We specify a single, reproducible pipeline that maps arbitrary PDB/mmCIF inputs to fixed-dimensional environment vectors using only structural information and element types. The design is fully deterministic and free of stochastic training, so that identical inputs produce identical outputs under a fixed software environment.
3. **A unified evaluation protocol that probes the representation rather than a specific model.** We introduce a site-grouped cross-validation scheme and a shallow Random Forest probe to quantify how much functional signal the environment representation exposes across diverse proteins, including large multimeric enzymes and complexes such as Rubisco, GroEL/GroES, SecA, and p53–DNA. The classifier is deliberately kept simple and untuned so that the measured separation reflects properties of the representation itself rather than optimisation of a particular architecture.

Beyond the static case studies considered here, the manifold perspective is deliberately chosen so that both the underlying configuration manifold and the induced environment-vector manifold can serve as targets for future geometric and topological analysis. Time-resolved trajectories (*p, t*) ↦ Ep_*p*_(*t*) define curves and ensembles in environment space on which persistent homology and related tools can be applied, while interface regions in complexes can be modelled as contact submanifolds equipped with geometric and topological invariants. In this sense, the present residue-level benchmarks are intended as a first validation step within a more general programme: using a single, interpretable environment representation as a common coordinate system for describing intra-protein organisation, inter-protein interactions, and, ultimately, more general molecular systems in ℝ^3^ ×*T*. The remainder of this article develops the mathematical framework, details the construction of the environment vector and the resulting deterministic pipeline, and applies the unified evaluation protocol to a set of representative case studies, with the aim of making the representation and its implementation precise and transparent enough to serve as a reusable front end for subsequent empirical work.

## 2 Theoretical Framework

### 2.1 Manifold representation of protein structure

We model a protein’s C*α* trace as a one-dimensional curve *γ* ⊂ ℝ^3^ ×*T* and endow it with an induced tubular surface metric. For convenience, we refer to this surface approximation as a “2-D manifold” on which all geometric quantities are evaluated, while the implementation itself operates directly on the C*α* point cloud.

Let *M* denote this tubular manifold equipped with the metric *g* inherited from the ambient Euclidean space. Distances, geodesics and curvatures are therefore well defined on *M*, and each residue *p* ∈ *M* is assigned a coordinate neighbourhood *U*_*p*_ comprising all residues within a fixed cutoff radius (typically 5–7 ^Å^). This manifold perspective allows localized analyses to be formulated in a coordinate-free way and makes them less sensitive to global conformational changes, while still remaining close to common structure-based workflows. In principle, the configuration space is the product

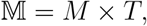

where *T* indexes time or external conditions; in the present work we only use static structural snapshots and therefore work at a fixed time slice, which we suppress in the notation below.

### 2.2 Environment vector as a bundle section

For each residue *p* in the manifold we define an *environment vector E*_*p*_ that encodes the local physico-chemical environment around *p*, inspired by conceptual Density Functional Theory and environment-style representations of local structure[19, 20, 6, 21]. Formally, we consider the trivial rank-*k* vector bundle

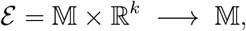

with fibre ℝ^*k*^ over each configuration point, and we treat the environment vectors as a section

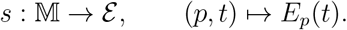

All numerical experiments in this paper are carried out on static structures, so we work at a fixed time and write simply *E*_*p*_.

At the level of the continuous model, we introduce a *theoretical environment vector* 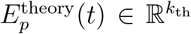 defined by integrals of local charge, density, curvature and external-field quantities over the neighbourhood *U*_*p*_(*t*):

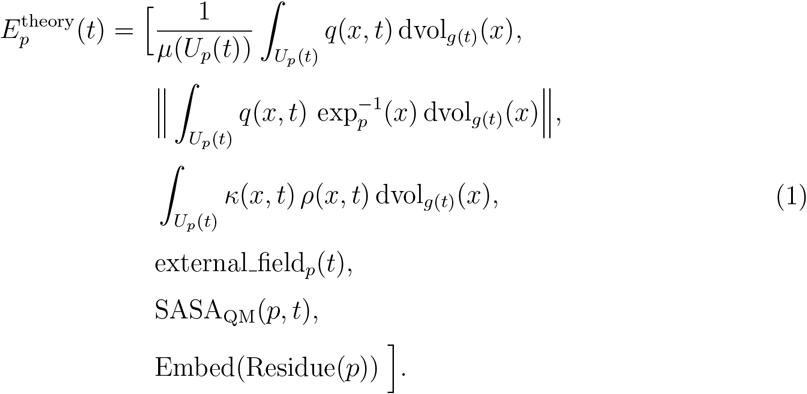

Here *µ*(*U*_*p*_(*t*)) is the volume of the neighbourhood with respect to the induced volume form, *q*(*x, t*) denotes a local charge density, 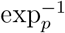 is the inverse exponential map on *M* so that 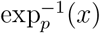 represents the displacement from *p* to *x* in the tangent space, *κ*(*x, t*) and *ρ*(*x, t*) denote curvature and (local mass or electron) density respectively, and SASA_QM_(*p, t*) is a density-defined solvent exposure functional. The term Embed(Residue(*p*)) is a low-dimensional encoding of the residue identity.

The *external-field* block external field_*p*_(*t*) collects local contributions from physically salient species such as metals, phosphates and halogens that act as external sources for the density and charge distribution. Conceptually, we view external field_*p*_(*t*) as a finite collection of integrals of the form

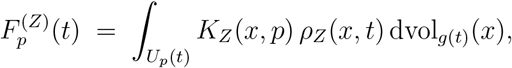

where *Z* indexes element sets (e.g. metal ions, phosphate groups, halogenated moieties), *ρ*_*Z*_ is the corresponding partial density, and *K*_*Z*_ is a kernel that decays with distance from *p*. In other words, the external-field block aggregates how strongly different classes of external sources “illuminate” the environment around *p*. This matches the practical implementation used throughout the experiments, where local metal-, phosphate- and halogen-induced fields are computed deterministically from the structural model.

The SASA term is defined in principle from the electron density: for a threshold *ρ*_0_, we consider the isodensity surface 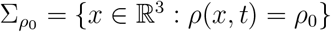 and define SASA_QM_(*p, t*) as the area of the patch of 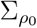 naturally associated with residue *p*. In practice, we approximate this by the classical solvent-accessible surface area computed from atomic coordinates and van der Waals radii using the Shrake–Rupley algorithm with a probe radius of 1.4 Å, and we interpret the resulting SASA component in *E*_*p*_ as a geometric surrogate for the underlying density-defined exposure functional.

Finally, the residue-identity term embeds each amino acid into a low-dimensional vector whose coordinates represent relative physicochemical descriptors (e.g. hydrophobicity, charge, size). This embedding allows chemically distinct environments to be distinguished even when their coarse geometric descriptors are similar.

#### Simulation vector and discrete implementation

For numerical experiments we work with a finite-dimensional *simulation vector* 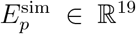 that discretises and extends the theoretical construction. We write

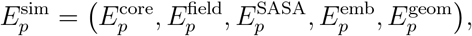

where:

- 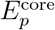 collects discrete counterparts of the average charge, charge dipole and curvature–density terms from Eq. (1), evaluated from a promolecular surrogate density constructed from the atomic model;
- 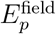 is a small block summarising local external-field contributions from metals, phosphates and halogens, obtained by integrating element-specific kernels over nearby atoms;
- 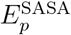 is the classical SASA of residue *p*;
- 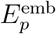 is a low-dimensional residue-identity embedding;
- 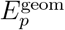 is a ten-dimensional block of neighbour-geometry descriptors that summarise radial and angular moments of the local environment measure.

Concretely, the geometric block 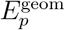 consists of: (i) an inner neighbour count *N*_in_(*p*), (ii) an outer neighbour count *N*_out_(*p*), (iii) a shell ratio *R*_shell_(*p*) = *N*_in_(*p*)*/*(*N*_out_(*p*) + *ε*), (iv)–(v) the mean and variance of the neighbour distances *r*_*q*_ = *d*(*p, q*) for *q* ∈ *U*_*p*_, (vi)–(viii) three angular descriptors derived from the eigenvalues *λ*_1_ ≥ *λ*_2_ ≥ *λ*_3_ of a local neighbourdirection tensor (linearity *λ*_1_ − *λ*_2_, planarity *λ*_2_ − *λ*_3_ and isotropy *λ*_3_), and (ix)–(x) mean and Gaussian curvature estimates obtained by fitting a quadratic patch to the neighbour cloud in a local tangent frame. These ten descriptors provide a compact summary of the radial and angular structure of the local environment without introducing additional physical assumptions beyond the manifold–density formalism of 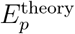.

We emphasise that 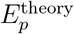 is conceptually a subset of 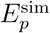: the latter augments the former with additional geometric moments that improve robustness on noisy, discretised structures. All components are normalised to dimensionless units and standardised (zero mean, unit variance) before statistical analysis so that no single block dominates purely by scale. We view the mapping (*p, t*) ↦ *E*_*p*_(*t*) as a coarse-grained functional of the underlying electron density and geometry: many distinct density fields can map to the same *E*_*p*_, and our goal is not to reconstruct *ρ*(*x, t*) but to retain enough information to separate reactive from non-reactive residues.

#### Environment manifold and metric structure

The environment construction induces a metric–measure structure on environment space. A rigorous formalisation is provided in Appendix E; here we summarise the main idea. We treat the map

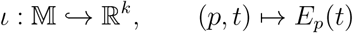

as a *C*^1^ immersion whose image *E* = *ι*(𝕄) ⊂ ℝ^*k*^ is an *r*-dimensional embedded submanifold (Definition B.1 in Appendix B). The ambient Euclidean metric, an embedded *L*^2^ metric derived from density overlaps, and a density–overlap metric induced by the promolecular construction all generate the same topology on *E* and render (*E, d*) complete (Theorem B.2, Appendix B). Equipped with the pushforward of a reference measure on 𝕄, the environment manifold (*E, d*, ν) becomes a well-defined metric–measure space on which distances, expectations and kernel operators are mathematically well posed. All statistical analyses in this work can therefore be interpreted as operating on this common environment manifold, with different classifiers or scoring functions corresponding to different functionals on (*E, d, ν*).

### 2.3 Fibre structure for reactive states

At the level of the present experiments, all information is carried by the environment section 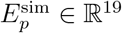 in the vector bundle ℰ → 𝕄 described above. For future extensions, we introduce an additional fibre layer that attaches high-resolution quantum information only to selected residues.

Formally, we consider a second bundle *π* : ℱ → *M* whose fibre *F*_*p*_ = *π*^*−*1^(*p*) represents the *reactive state* of residue *p*. In regions that are not reactive, *F*_*p*_ is trivial and carries no additional data. At a putative reactive site, by contrast, *F*_*p*_ contains a localised quantum state—such as an electron-density patch or wavefunction—encoding that residue’s reactivity potential. These fibre states can be viewed as high-resolution “quantum attachments” to the geometric manifold: they add detailed chemical degrees of freedom exactly where they are needed (e.g. at catalytic or metal-coordination sites) while leaving inert regions described solely by the coarse environment vectors.

In practical terms, a non-trivial fibre could be instantiated by a pre-computed electron-density cloud or an analytic function centred on a reactive residue, obtained for example from density-functional or semi-empirical calculations. All other residues carry the trivial fibre (identity map), meaning they are treated uniformly and contribute only through their environment vectors. The transition between trivial and non-trivial fibres is defined smoothly to avoid discontinuities on *M*.

When localised fibres of two residues *p* and *q* interact, they generate an *interaction submanifold*

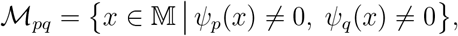

on which one can in principle measure purely geometric invariants such as the area Area (ℳ_*pq*_), Euler characteristic *χ*(ℳ _*pq*_) and fibre-overlap homology classes. Such invariants could form the basis of a more deductive analysis of protein–protein interfaces, in which interface regions are treated as contact submanifolds equipped with geometric and topological signatures. In the present work, however, all numerical results are obtained solely from the base manifold and the environment vectors 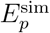; the quantum fibre layer and associated interaction invariants are introduced here as a theoretical extension for future interface-level analysis.

## 3. Methods

### 3.1 Overview of the 1–9 Environment Pipeline

To instantiate the geometric and density-based framework from Section 2 on static protein structures, we implemented a residue-centric environment pipeline that maps a single structural snapshot to per-residue environment vectors used for downstream scoring. Throughout this section we work at a fixed time slice *t* = 0 and suppress *t* from the notation.

The pipeline consists of nine stages:

1. structure parsing and residue/atom indexing,
2. residue–residue contact graph construction,
3. surrogate (promolecular) electron-density generation,
4. optional quantum-corrected density replacement on a selected subset of residues,
5. density-derived local descriptors,
6. charge-derived descriptors (quantum or surrogate),
7. definition of local integration domains *U*_*p*_,
8. computation of theoretical environment components, and
9. assembly of final environment vectors.

All stages are executed in a single integrated script, so that fragment definitions, grids, and parameters are shared consistently across Steps 3–9.

#### 3.1.1 Input Structures and Residue Representation (Step 1)

Protein structures were read from PDB or mmCIF files using standard parsers. Only the first structural model was used. Each residue *p* was represented by a dictionary containing chain identifier, residue sequence number, optional insertion code, residue type, hetero flag, and a list of atoms. Each atom was assigned a global integer index and stored with element type and Cartesian coordinates **r**_*a*_ ∈ ℝ^3^. Residues with zero atoms were ignored. The residue identifier was formatted as `chain:resseq[icode]` (e.g. `A:201`).

#### 3.1.2 Solvent-Accessible Surface Area (SASA) (Step 1b)

Per-residue solvent-accessible surface area was computed by the Shrake–Rupley algorithm as implemented in MDTraj (probe radius 1.4 Å, residue mode). If MDTraj was unavailable or if residue mapping failed, SASA values were set to zero. Residue-level mapping was performed by matching (chain, resSeq, icode) keys between the MDTraj topology and the parsed residue list. Let SASA(*p*) denote the resulting 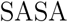, for residue *p*. We also computed the mean SASA over residues, 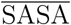, for later scaling.

#### 3.1.3 Residue Contact Graph and Fragment Definition (Step 2)

To define local neighbourhoods, we computed residue–residue contacts using atomic proximity. Let *r* be a fixed chart radius. For every atom *a* we queried all atoms *b* satisfying

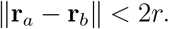

These queries were performed using a global KDTree over all atomic coordinates; when KDTree was unavailable, an *O*(*N* ^2^) fallback was used. If any atom pair across two distinct residues satisfied this condition, the residues were considered overlapping neighbours. This generated (i) a residue adjacency list 𝒩(*p*) (“overlaps”) and (ii) atom-level contact pairs for diagnostic output.

For each residue *p*, we defined its *fragment F*_*p*_ as the atom set of *p* union all atoms from neighbouring residues:

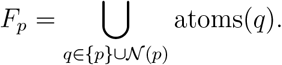

All subsequent density and charge calculations for residue *p* were performed on *F*_*p*_. This ensures that the same neighbourhood definition is used consistently throughout Steps 3–9 and provides a discrete realisation of the theoretical neighbourhood *U*_*p*_.

#### 3.1.4 Surrogate Promolecular Electron Density (B-mode) (Step 3)

Because we do not assume access to full quantum electron densities for large proteins, we constructed a local surrogate density on a 3D grid around *F*_*p*_. For each fragment we built a Cartesian grid {*x*_*i*_} spanning the fragment bounding box with an isotropic padding *P* (default 2.0 Å) to avoid truncating Gaussian tails. The grid spacing was Δ = 0.5 Å, and the grid-cell volume was Δ*V* = Δ^3^.

In the theoretical formulation, *U*_*p*_(*t*) is an open geodesic neighbourhood around residue *p*. In practice we approximate *U*_*p*_ at *t* = 0 by the fragment grid associated with *F*_*p*_, which provides stable numerical integration and avoids under-sampling on narrow local charts.

We defined the B-mode promolecular density as a sum of element-wise Gaussian atomic kernels:

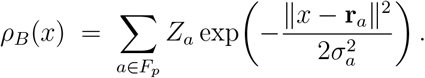

Here *Z*_*a*_ is a rough nuclear-charge proxy taken from a fixed lookup table, and *σ*_*a*_ is a heuristic width chosen on the scale of atomic radii. This density is not intended to reproduce quantum densities; it is a smooth surrogate field for residue-local integration.

#### 3.1.5 Optional Quantum-Corrected Density (A-mode) (Step 4)

For a user-selected residue subset 𝒜, we computed semi-empirical charges using GFN2-xTB on the corresponding fragment *F*_*p*_. Each fragment was exported to an XYZ file and xTB was executed with the `--charges` option to obtain per-atom partial charges 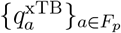. If xTB failed, the residue reverted to B-mode.

When charges were obtained, we produced an A-mode corrected density by reweighting the same Gaussian kernels:

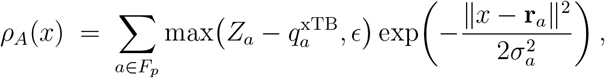

with *ϵ* = 0.1 to maintain positivity. For residues not in 𝒜, or for failed xTB runs, the final density for feature extraction was *ρ*_*B*_. We denote the final density used for residue *p* by *ρ*_*p*_ ∈ {*ρ*_*B*_, *ρ*_*A*_}.

#### 3.1.6 Density-Derived Local Descriptors and Element-Resolved Channels (Step 5)

Using the final density *ρ*_*p*_ on its fragment grid, we computed a set of density-based descriptors around residue *p*.

Let *r*_*p*_ be the geometric centre of residue *p* (mean of its atomic coordinates). The total local density was approximated by a Riemann sum over the grid points {*x*_*i*_} associated with fragment *F*_*p*_:

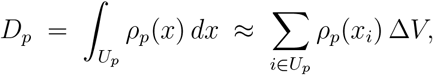

where Δ*V* = Δ^3^ is the grid-cell volume and *U*_*p*_ is the local integration mask defined in Section 3.1.7.

We defined a normalised density barycentre *c*_*p*_ as

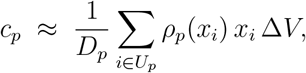

provided that *D*_*p*_ *>* 0; otherwise *c*_*p*_ was set to *r*_*p*_.

We also computed a dipole-like density moment relative to *r*_*p*_,

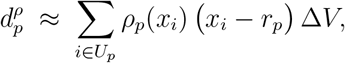

which serves as a coarse descriptor of anisotropy in the local density distribution.

Finally, we computed a curvature-like smoothness scalar from the density gradient:

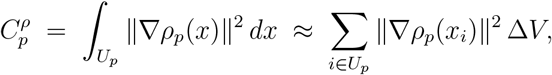

where ∇*ρ*_*p*_ was evaluated by finite differences on the fragment grid under implicit Neumann boundary conditions. This term is a surrogate for a “curvature-weighted density” and should be interpreted as a smoothness/roughness measure of the surrogate density, rather than a geometric curvature of a fixed isosurface.

##### Element-resolved density channels

To capture the influence of chemically distinct atom groups without using any residue-type labels, we additionally decomposed *ρ*_*p*_ into element-resolved channels. Let 𝒜_*p*_ denote the atom set of fragment *F*_*p*_, and define disjoint subsets based on element type and hetero flags:

- 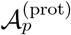: protein heavy atoms (standard amino-acid residues),
- 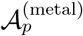: metal atoms (e.g. Mg, Mn, Zn, Ca, Fe, Ni, Cu, Co),
- 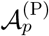: phosphorus atoms,
- 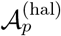: halogen atoms (F, Cl, Br, I),
- 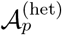: non-protein heavy atoms from HETATM records (excluding crystallographic waters).

For each channel *c* ∈ {prot, metal, P, hal, het} we defined a channel-specific density by restricting the Gaussian sum in *ρ*_*p*_ to the corresponding atom subset:

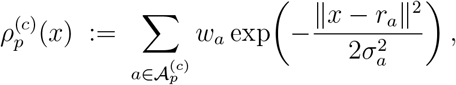

with the same kernel widths *σ*_*a*_ and weights *w*_*a*_ inherited from the B-mode/A-mode definition (either *Z*_*a*_ or the charge-reweighted coefficient).

We then computed a channel-resolved local density for each *c*:

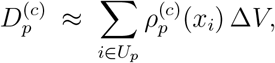

and, when *D*_*p*_ *>* 0, the corresponding fractional contributions

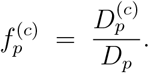

The scalars 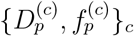 enter the environment vector as additional components. They quantify how much of the local surrogate density around residue *p* is carried by protein atoms, metals, phosphorus, halogens, or hetero-ligand atoms, without relying on residue-type or binding-site annotations.

#### 3.1.7 Charge-Derived Descriptors (Step 6)

##### A-mode charges

For residues *p* ∈ 𝒜 with successful xTB runs, we computed per-residue net charge

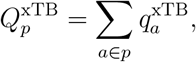

and a charge dipole moment

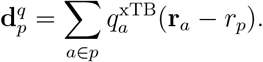

These quantities were computed using the same fragment *F*_*p*_ and atom indexing as above to prevent mismatch.

##### Residue-level charge surrogates (B-mode)

In the theoretical environment vector, the charge-like component is intended to represent a coarse local average of the underlying charge field *q*(*x*) over the fragment domain *U*_*p*_ associated with residue *p*. When quantum-mechanical (QM) results are available, we use the xTB-derived residue net charges 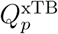 and their associated dipole moments. For structures without QM data, we construct a residue-level surrogate charge purely from atomic element information, without using any residue-type labels.

##### Atom-based base charge 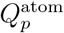

Let 𝒜_*p*_ denote the set of atoms belonging to residue *p*, and let *E*_*a*_ be the element symbol of atom *a* ∈ 𝒜_*p*_ (e.g. C, N, O, S, Mg). We define a simple element-dependent weight function *w*_elem_ : {elements} → ℝ that encodes coarse electrostatic tendencies:

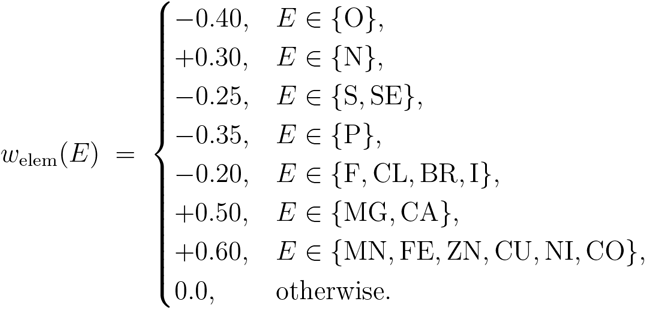

This mapping is not meant to approximate formal charges or partial charges; instead, it is a coarse, element-resolved surrogate that assigns negative weights to electronegative atoms (O, halogens, P, S/Se) and positive weights to typical cationic metals. Carbon and hydrogen are treated as neutral background and contribute 0 in this B-mode surrogate.

The atom-based base charge for residue *p* is then defined as

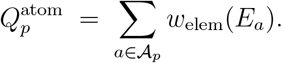

This quantity depends only on the atomic composition of the residue and does not use any residue-type or motif annotation.

##### SASA-modulated B-mode charge 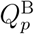

As in the original B-mode construction, we modulate the base charge by a SASA-dependent scaling factor to reflect the fact that buried charged groups are less exposed to the solvent and external field than solvent-accessible ones.

Let SASA(*p*) denote the solvent-accessible surface area of residue *p*, and let 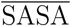 be the mean SASA across all residues in the structure (Section 3.1.2). We define a scaling factor

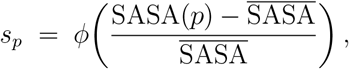

where *ϕ* is a bounded, monotone function such as

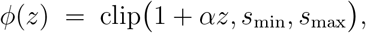

with a user-specified slope *α* and lower/upper caps *s*_min_, *s*_max_ (e.g. *s*_min_ = 0.5, *s*_max_ = 1.5). In the implementation, this scaling corresponds to the function `env_scale_from_sasa` applied to the residue-level SASA.

The SASA-modulated B-mode charge is

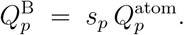

##### Charge component in the environment vector

For residues with xTB-derived charges available, we use

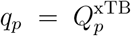

as the charge component in the theoretical environment vector. For all other residues, we fall back to the atom-based surrogate,

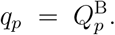

This hierarchy ensures that the theoretical environment vector remains free of any residue-type or motif-based charge assignment. The only inputs are atomic positions, element types, and SASA, which are consistently available in static PDB/mmCIF structures.

#### 3.1.8 Local Integration Domains *U*_*p*_ (Step 7)

The theoretical model defines *U*_*p*_(*t*) as an open neighbourhood around residue *p* in the configuration manifold *M* × *T*. In the static setting used here we fix *t* = 0 and induce *U*_*p*_ from the fragment grid and residue atoms. Specifically, let *R*_env_ be an environment-ball radius (default 2.0 Å). We defined

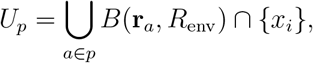

i.e. the set of grid points within *R*_env_ of any atom in residue *p*. All theoretical integrals over *U*_*p*_ were approximated by sums over these masked grid points.

#### 3.1.9 Theoretical Environment Components (Step 8)

We assembled a theoretical environment vector 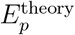 corresponding to Eq. (1) of the model. Continuous fields *q*(*x*), *ρ*(*x*), and *κ*(*x*) are represented discretely: *q*(*x*) is taken as a residue-level net charge (quantum or surrogate), and *ρ*(*x*) is the surrogate density *ρ*_*p*_ defined in Sections 3.1.5–3.1.6. The components are:

##### 1. Average charge

We used residue net charge as a coarse approximation to *µ*(*U*_*p*_)^−1^ ∫*U*_*p*_ *q*(*x*) *dx*:

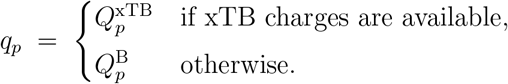

##### 2. Dipole magnitude

We used 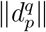 if xTB charges were available, and otherwise fell back to the density-dipole magnitude 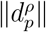.

##### 3. Curvature-weighted density surrogate

We set this term to 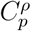, acknowledging that it is a smoothness proxy derived from *ρ*_*p*_ rather than a geometric curvature of a fixed isosurface.

#### 4. External-field block from element-resolved sources (two variants)

To capture the influence of chemically distinct external sources (e.g. metals, phosphorus-rich cofactors, or halogenated ligands) without using biochemical labels, we define a compact three-dimensional external-field block for each residue *p*,

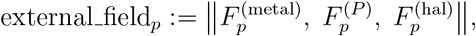

which then enters the environment vector as a single structured component.

We implemented two closely related constructions of 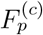, *c* ∈ {metal, *P*, hal} :

##### (i) Global Gaussian external field (structure-wide variant)

In the first variant, we define smooth scalar field descriptors at the residue centre *r*_*p*_ by summing distance-weighted contributions from all atoms of the relevant element class in the entire structure. Let 𝒜^(metal)^, 𝒜^(*P*)^, and 𝒜^(hal)^ denote the global sets of atoms in the structure whose element types belong to user-specified metal, phosphorus, and halogen lists, respectively (e.g. metals: Mg, Mn, Zn, Ca, Fe, Ni, Cu, Co; halogens: F, Cl, Br, I). For each residue *p* and each channel *c* ∈ {metal, *P*, hal}, we set

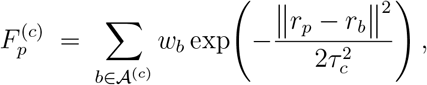

where *w*_*b*_ is either the atomic number *Z*_*b*_ or a constant weight, and *τ*_*c*_ is a fixed length scale (on the order of a few Å) controlling how rapidly the contribution decays with distance. This variant yields a global, structure-wide external field evaluated at *r*_*p*_ and requires an additional loop over the corresponding atom sets.

##### (ii) Channel-based local external field (density-channel variant)

In the second variant, we reuse the element-resolved density channels already computed in Step 5. Recall that for each fragment *F*_*p*_ we defined channel-specific densities 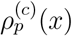 and their integrated local contributions

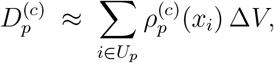

together with the total local density

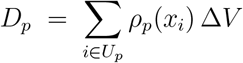

and the fractional contributions

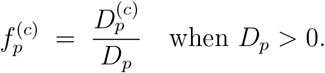

In the channel-based variant, we simply set

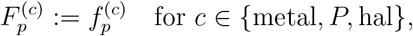

with 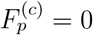 if *D*_*p*_ = 0. Thus,

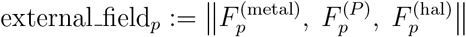

is a compact, element-resolved external-field surrogate derived purely from the local density channels around residue *p*. This construction does not require any additional grid passes or global atom loops beyond those already used to compute the density channels, so the asymptotic cost of Step 8 is unchanged.

Both variants depend only on atomic positions and element types and remain free of any residue-type or binding-site annotations. The global Gaussian variant (i) explicitly aggregates contributions from all metals, phosphorus atoms, and halogens in the structure and is useful when one wishes to treat these as long-range field sources. In contrast, the channel-based variant (ii) treats the external-field block as a normalised summary of the local density composition around residue *p*, which is more computationally lightweight and more tightly coupled to the fragment *F*_*p*_.

In all experiments reported in this work we used the channel-based variant (ii) as the default choice for external field_*p*_, while keeping the global Gaussian variant (i) available in the reference implementation for ablation and sensitivity studies.

##### SASA

We inserted SASA(*p*) as a scalar descriptor of local solvent exposure.

##### Residue embedding

Each residue type was optionally encoded as a four-dimensional physicochemical vector as described in Section 3.1.9. In ablation experiments focused on purely geometric and element-based descriptors, this embedding can be dropped.

The concatenation of these components defines the theoretical environment vector 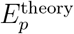 used in downstream analysis and supervised evaluation.

#### 3.1.10 Physicochemical 4D Residue Embedding (Step 8, component 6)

To provide a low-dimensional but interpretable residue-identity term, we mapped each residue to a 4D real vector

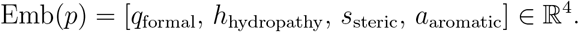

The coordinates follow a physically motivated ordering:

1. *q*_formal_: formal side-chain charge at near-neutral pH (e.g. ASP/GLU = −1, LYS/ARG = +1, others 0).
2. *h*_hydropathy_: normalised hydropathy index (Kyte–Doolittle scale, linearly rescaled to [−1, 1]).
3. *s*_steric_: normalised steric size/volume proxy (linearly rescaled to [0, 1]).
4. *a*_aromatic_: aromaticity indicator (1 for aromatic residues, 0 otherwise).

This embedding replaces one-hot encodings and is designed to preserve coarse physicochemical similarity while remaining lightweight for inference.

#### 3.1.11 Geometric Neighbour Augmentation (Step 9 extra features)

In addition to 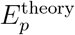, we augment each residue *p* with a compact family of purely geometric descriptors computed from the contact graph 𝒩 (*p*) of C*α*-based residue centres. These descriptors are interpreted as low-order radial and angular moments of the local environment around *p*:

##### 1. Two-shell radial density profile

For a fixed cutoff radius *r*_max_, we split the neighbourhood into an inner and an outer shell,

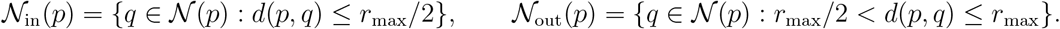

We record the inner and outer counts

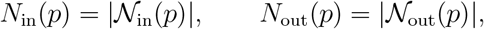

together with the shell ratio

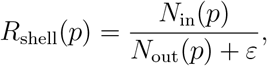

which distinguishes compact, core-like neighbourhoods from more diffuse or one-sided environments.

##### 2. Radial distribution moments

Let *r*_*q*_ = *d*(*p, q*) for *q* ∈ 𝒩 (*p*). We compute the first two moments of the radial distribution,

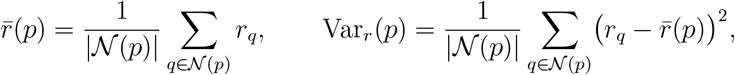

capturing both the typical neighbour distance and how tightly neighbours are concentrated around that radius.

##### 3. Angular shape tensor and anisotropy

For each neighbour we define a unit direction vector

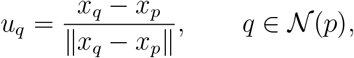

and form the 3 × 3 direction tensor

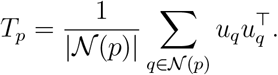

Let *λ*_1_ ≥ *λ*_2_ ≥ *λ*_3_ denote the eigenvalues of *T*_*p*_. From these we derive scalar shape descriptors

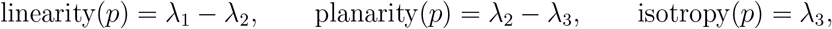

which distinguish bulk-like, planar-surface, and ridge-like local packing around *p*.

##### 4. Local surface curvature surrogate

In the local coordinate frame defined by the principal directions of *T*_*p*_, we fit a quadratic height function

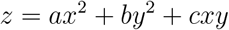

to the neighbour cloud by least squares. The fitted coefficients are converted into mean and Gaussian curvatures (*H*_*p*_, *K*_*p*_), providing a coarse estimate of whether *p* lies in a convex bulge, concave pocket, or saddle-like region.

Residue centres were defined as geometric means of atomic coordinates.

#### 3.1.12 Final Environment Vectors (Step 9)

The final theoretical environment vector for residue *p* collects the components defined in the previous section, including the external-field block

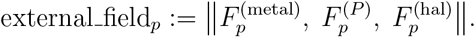

We write

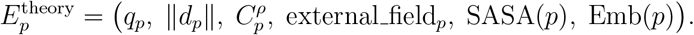

Here *q*_p_ is the residue-level charge component (quantum or surrogate), *d* is the charge-or density-derived dipole magnitude, 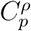 is the curvature-weighted density surrogate, 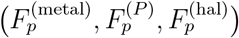 is the external-field block defined from element-resolved sources, SASA(*p*) is the solvent-accessible surface area, and Emb(*p*) is the optional 4D physicochemical residue embedding.

For predictive experiments we augment 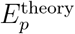 with ten purely geometric neighbour descriptors from the previous section to obtain the simulation vector

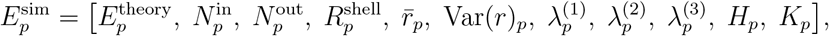

where 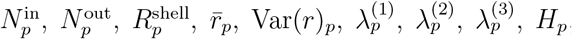, and *K*_*p*_ are the neighbour-count, radial-moment, anisotropy, and curvature descriptors defined above. Stacking over residues yields the matrices saved as `Ep_theory.npy` and `Ep_sim.npy`, with a manifest file recording residue ordering, feature names, and parameter settings.

#### 3.1.13 Outputs and Reproducibility

The pipeline produces: `meta.json` (residues, overlaps, SASA, atom contacts), `density_final.json` (A/B density provenance), optional density grids (rhoB_^*^.npy, rhoA_^*^.npy), `env_features.json` (Steps 5–6 descriptors), `Ep_theory.npy` and `Ep_sim.npy` with `Ep_manifest.json`. All parameters (*r, P*, Δ, *R*_env_, xTB options, SASA scaling *α*) are explicitly stored in the manifest to ensure exact reproduction.

### 3.2 Supervised evaluation of residue-level feature representations

#### 3.2.1 Residue-level feature matrix

For a given protein structure, we represent each residue *i* by a fixed-length feature vector **x**_*i*_ ∈ ℝ^*d*^ that summarizes its local structural and physicochemical environment (e.g. geometric descriptors, solvent accessibility, density-based quantities, or learned embeddings). Stacking all residue-level feature vectors yields a matrix

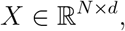

where *N* is the number of residues in the protein (or in a set of proteins) and *d* is the number of features per residue.

When features are computed from heterogeneous sources, missing values may arise for specific residues or descriptors. To handle this in a reproducible way, we apply a simple column-wise mean imputation separately within each training split: for each feature dimension *j* ∈ {1, …, *d*}, any missing entries *X*_*ij*_ in the training data are replaced by the empirical mean of the non-missing values in column *j* computed on the training residues of that split. The fitted imputer is then applied to the corresponding test residues of the same split. In code, this corresponds to using a `SimpleImputer(strategy=“mean”)` fitted on the training subset only and then applied to both training and test subsets.

##### Solvent removal

For all benchmarks, we remove solvent molecules at the level of the residue–feature manifest before label assignment and supervised evaluation. Given the unbound structure and the engine-generated manifest of residue identifiers, we reconstruct a mapping from manifest IDs (`CHAIN:RESSEQ[ICODE]`) to residue names using either a `meta.json` sidecar file (preferred, when available) or by re-parsing the unbound structure with MDTraj. We then discard all entries whose residue name matches a user-configurable solvent list (default: `HOH, WAT, TIP3, DOD`), while keeping and counting entries whose residue name cannot be resolved. The feature matrix *X* ∈ ℝ^*N ×d*^ and the corresponding list of residue identifiers are simultaneously filtered by the same boolean mask, and all label construction and Policy P evaluations are performed on this solvent-free subset. A summary of the number of removed solvent residues, unresolved residue names, and retained entries is stored in a JSON report for each case.

#### 3.2.2 Definition of residue-level labels

Each residue *i* is assigned a binary label *y*_*i*_ ∈ {0, 1} according to an externally curated notion of *functional involvement*, based on structural and biochemical information from the literature (active-site annotations, UniProt/PDB functional annotations, mutational scanning, interface mapping, etc.). Rather than fixing a single notion of “functional” versus “background”, we construct a *panel* of binary classification tasks, each defined by a different choice of positive set 𝒫_*t*_ ⊆ {1, …, *N*} indexed by *t* (task index). For a given task *t* we define

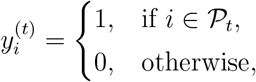

so that all tasks share the same feature matrix *X* and evaluation protocol while differing only in the definition of the positive class. This allows us to probe how well a single environment representation separates different *functional types* of residues from background.

Across all complexes considered in this work, we adopt a common abstract taxonomy of residue roles and then instantiate it on a per–protein basis using structural and biochemical information from the literature. At the top level we distinguish two tiers:

- **Tier 1 (functional core + support)**. Residues with strong, local evidence for functional involvement. This tier is further subdivided into role classes that are shared across systems:
  – *catalytic/metal core:* residues that directly participate in chemical catalysis (general acid/base, nucleophiles, enediolate stabilisers) or coordinate essential metal ions;
  – *PTM core:* residues that are themselves sites of well-characterised post-translational modification (e.g. Lys carbamylation/acetylation, phosphorylation) when the modification is mechanistically essential;
  – *gates and latches:* loop or tail residues that open or close access to the active site, or that stabilise well-defined open/closed conformations;
  – *interfaces and anchors (PPI / DNA / ligand):* residues that contribute directly to binding energy at protein–protein, protein–DNA or protein–ligand interfaces (e.g. hydrophobic patches, IVL-like anchors, core beta–hairpins at obligate interfaces);
  – *allosteric and assembly-network pivots:* residues that transmit conformational changes or stabilise allosteric networks (e.g. hinge-support positions, inter-subunit salt-bridge clusters) when these roles are defined at residue resolution in the literature.

- **Tier 2 (structural shell)**. Residues that form the immediate structural shell around Tier 1 cores and anchors. Typical examples include residues that maintain the geometry, packing or electrostatic balance of a catalytic pocket, interface anchor or hinge, but are not themselves essential hotspots. In practice, Tier 2 residues are defined by explicit, local neighbourhood rules (e.g. fixed residue-index ranges or direct-contact shells around Tier 1 cores) for each protein and may or may not have direct site-specific mutational evidence.

For each protein complex we instantiate this abstract scheme by assigning individual residues to one or more of the Tier 1 role classes based on high-confidence structural and biochemical evidence (e.g. active-site annotations, metal ligands, mutational scanning, interface mapping), and then defining Tier 2 shells by explicit local neighbourhood rules around these cores. Residue lists for each role and tier (e.g. catalytic core vs. PTM core vs. gate vs. interface vs. allosteric shell in a given complex) are defined in a protein-specific manner but always drawn from the above role taxonomy.

Given these assignments, we define several families of label sets:

- *global Tier 1* (𝒫_T1−all_): the union of all Tier 1 residues across all role classes in a given complex;
- *global Tier 2* (𝒫_T2−all_): the union of all Tier 2 shell residues in that complex;
- *role-specific Tier 1 sets* (𝒫_*r*_): for each role class *r* (catalytic/metal core, PTM core, gates/latches, interface anchors, allosteric/assembly pivots), the subset of Tier 1 residues assigned to that role;
- *role-specific Tier 1+2 unions* (𝒫_*r*,T1+T2_): for roles where a clear supporting shell is well defined (e.g. catalytic basin, clamp, relay helix), the union of the Tier 1 core and its Tier 2 shell.

Each such set 𝒫_*t*_ defines a binary classification task as above, with all remaining residues in the complex treated as negatives. These negatives should be interpreted as unlabeled background rather than as a claim of functional irrelevance for those residues. Importantly, several biologically relevant mechanisms are *intentionally left outside* the label space and thus treated as background in all tasks: residues whose importance is dominated by long-range electrostatic steering, intrinsically disordered segments, phase-separating low-complexity regions or global mechanical properties of large-scale domain motions are not explicitly annotated in the present work. The scope of the current benchmarks is therefore restricted by design to functional types that are expected to leave a strong signature in a static, local environment descriptor. For notational simplicity, we omit the task index *t* in what follows and write **y** = (*y*_1_, …, *y*_*N*_)^⊤^ ∈ {0, 1}^*N*^ for a generic label vector; all procedures are applied independently to each label set in the panel.

##### Automatic interface labels for the DB5.5 benchmark

For the Docking Benchmark 5.5 (DB5.5) set of protein–protein complexes, we additionally construct an *automatic* interface label for the partner being evaluated (“ligand” or “receptor” in the original benchmark terminology). For each complex, we first build a bound co-complex by concatenating the PDB files corresponding to the bound ligand and receptor structures and retaining only `ATOM`/`HETATM` records. Using MDTraj, we collect all protein residues on the target partner and on its binding partner, and define a representative coordinate for each residue (preferentially the C_*α*_ atom, otherwise the first heavy atom). We then perform a two-stage distance search: a coarse neighbour search with a KD-tree in Cartesian space using a radial cutoff equal to the sum of a hard interface distance and a margin (5 Å + 4 Å), followed by an exact computation of closest heavy-atom distances for all candidate residue pairs with MDTraj’s `compute_contacts` (“closest-heavy” scheme). A residue on the target partner is marked as an interface positive if it has at least one partner residue within 5 Å in this bound complex. These bound interface labels are then mapped back to the unbound structures by sequence alignment: bound and unbound chains are globally aligned with Biopython’s pairwise alignment routines, putative chain matches are filtered by minimum identity and coverage thresholds, and the resulting one-to-one chain assignments are used to project residue identifiers from bound to unbound coordinates. Only interface residues that (i) satisfy the distance criterion in the bound complex, (ii) can be mapped to an unbound residue index, and (iii) appear in the unbound feature manifest are retained as positives; all other residues in the unbound structure (after solvent removal) are treated as background for the DB5.5 interface classification task.

#### 3.2.3 Random forest classifier and cross-validation protocols

To quantify how well a given feature representation separates functional from background residues under a particular label definition, we train a supervised classifier and evaluate its performance using the area under the receiver operating characteristic curve (AUC-ROC).

We use a random forest classifier (scikit-learn `RandomForestClassifier`) as a non-parametric baseline. Unless stated otherwise, we fix the hyperparameters as follows:

- `n_estimators = 100`: number of trees in the ensemble;
- `class_weight = “balanced”`: automatic class weighting inversely proportional to class frequencies to counter label imbalance;
- `random_state = 42`: fixed random seed to ensure reproducibility of tree construction and sampling;
- `n_jobs = 1`: single-threaded execution to avoid non-determinism from parallel scheduling.

Let *X*^imp^ denote the imputed feature matrix after applying the mean imputer within each training split, as described above. We consider two related evaluation protocols.

##### Residue-stratified cross-validation

As a baseline, we evaluate performance with stratified *K*-fold cross-validation at the residue level:

1. We construct a **stratified** split of the indices {1, …, *N*} into *K* folds (typically *K* = 5), preserving the overall positive:negative ratio in each fold. In code, this corresponds to `StratifiedKFold(n_splits=5, shuffle=True, random_state=42)`.
2. For each fold *k* ∈ {1, …, *K*} :
  a. Define the training indices 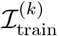 and test indices 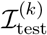.
  b. Fit the mean imputer on 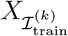 and transform both training and test feature matrices to obtain 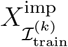 and 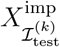.
  c. Fit a random forest classifier on 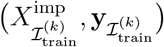.
  d. Obtain predicted class probabilities 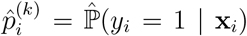 for all 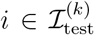 using `predict_proba`.
  e. Compute the AUC-ROC on the *k*-th test fold, 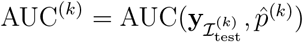
3. We report the mean and standard deviation of the AUC across folds:

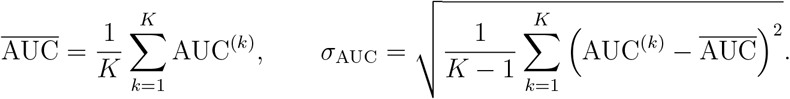

In this protocol, the basic sampling unit is an individual residue. This setting is appropriate when residues can be treated as approximately independent given the structure, and when symmetric or replicated copies of the same functional site are not included as positives in the same task.

#### Site-grouped cross-validation to avoid functional-site leakage

For symmetric oligomeric complexes (such as chaperonins) or assemblies in which multiple chains share identical functional sites (e.g. repeated ATP-binding pockets or GroES-anchoring motifs), residue-level cross-validation can overestimate performance: different copies of the same functional site may appear in both the training and test sets. To obtain a more conservative estimate of how well the features separate distinct functional sites from background, we additionally perform *site-grouped* cross-validation whenever the label definition includes replicated copies of the same site.

We define a discrete site identifier *g*_*i*_ for each residue *i* that groups residues belonging to the same functional site across chains. In practice, *g*_*i*_ can be constructed from metadata such as the protein role (e.g. GroEL vs. GroES), the chain category (cis vs. trans ring), and the residue index, so that residues sharing the same (role, residue index) are assigned to the same group. In heteromeric complexes that contain multiple protein species (e.g. kinase– substrate, small GTPase–effector, receptor–ligand), we explicitly assign distinct role labels to chains belonging to different proteins (e.g. “KRAS” vs. “RAF1”, “GroEL” vs. “GroES”). Site-groups are then constructed *only within a fixed role*: residues from different roles never share the same group identifier *g*_*i*_. Consequently, repeated or symmetry-related copies of the *same* functional site within one protein can be grouped and constrained not to leak across train and test folds, while structurally and functionally unrelated chains in the same PDB entry are evaluated as separate populations.

Formally, each residue is assigned to one of *M* site-groups:

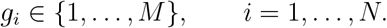

All residues with the same *g*_*i*_ are treated as a single “functional site” for the purposes of cross-validation.

We then apply a group-aware *K*-fold splitter (scikit-learn `GroupKFold`) that never places residues from the same site-group into both the training and test folds:

1. We construct fold assignments based on the site-group labels 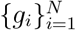 using `GroupKFold(n splits=K)`. By construction, each group *g* ∈ {1, …, *M* } is assigned entirely to the training set or entirely to the test set in each fold.
2. For each fold *k* ∈ {1, …, *K*}:
  a. Let 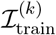 and 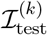 denote the training and test residue indices returned by GroupKFold.split, conditioned on the group labels {*g*_*i*_}.
  b. Fit the mean imputer on 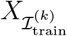 and transform both training and test feature matrices to obtain 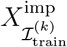 and 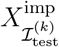.
  c. Fit a random forest classifier on 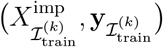
  d. Obtain predicted class probabilities 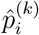 for all 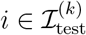 using predict proba.
  e. Compute the residue-level AUC-ROC on the *k*-th test fold as before.
3. As above, we summarize performance by the mean and standard deviation of the AUC across folds.

In practice, we adopt the following rule-of-thumb for choosing between residue-level and group-aware splitting. For a given label definition (e.g. T1-all, T1-fullDFT, T1-Ep, or a role-specific subtype), we first construct the site-group labels *g*_*i*_ as described above. If no positive residue shares its (role, residue index) with another positive residue (i.e. every positive site is unique and does not appear as a replicated copy across chains), then group-aware splitting provides no additional protection against site leakage, and we use `StratifiedKFold` on individual residues. If, however, the positive set includes replicated copies of the same site (multiple residues with identical role and residue index across symmetric chains), we regard these as a single functional site and employ `GroupKFold` to ensure that no such site can appear in both training and test folds. This simple criterion — “use GroupKFold if and only if the label set includes replicated copies of the same (role, index) site” — guarantees that functional-site leakage is avoided where it matters, while keeping the evaluation protocol simple for tasks without replicated sites.

Optionally, we also report a site-level AUC, in which each site-group *g* is treated as a single sample. For each test fold *k*, we aggregate residue-level predictions within site-group *g* to a single site score 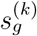(e.g. by taking the mean predicted probability across all residues *i* with *g*_*i*_ = *g* in the test set). A site is labeled as functional if any residue in that group has *y*_i_ = 1, and non-functional otherwise. The AUC-ROC is then computed across 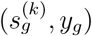 pairs. This site-level analysis provides a complementary measure of how well the features distinguish functional vs. non-functional sites, rather than individual residues, under the same no-leakage constraint.

#### Policy P for large-scale grouped evaluation on DB5.5

For the large-scale DB5.5 benchmark, we implement a fully automated grouped evaluation protocol, denoted Policy P, that is a concrete realisation of the site-grouped strategy above. For each unbound structure, we first cluster chains into sequence-similar symmetry groups by global pairwise alignment of chain sequences and thresholding on minimum identity and coverage (typically identity ≥0.95 and coverage ≥0.90). Each residue is then assigned a group label that combines its symmetry group identifier and residue index, so that symmetry-related copies of the same structural site share the same group label across chains. These group labels are passed to scikit-learn’s `GroupKFold` splitter, which enforces that no site-group can appear in both the training and test folds of a given split.

Policy P chooses the number of folds *K* adaptively for each label set based on the number of positive site-groups: starting from a target value (e.g. *K* = 5), *K* is decreased stepwise down to a minimum (e.g. *K* = 2) until the resulting grouped split yields at least two valid folds in which both the training and test partitions contain examples of both classes. On each valid fold, a mean imputer and random forest classifier (with the hyperparameters specified above) are fitted on the training residues and evaluated on the held-out residues, and both ROC–AUC and average precision (AUPRC) are recorded. Folds in which either the training or test labels are single-class are skipped. For each case, we report the mean and standard deviation of ROC–AUC and AUPRC over all valid folds, together with metadata describing the effective *K* used, the number of folds skipped due to single-class splits, and the total numbers of positive and negative residues after mapping and solvent removal. The same Policy P implementation is used for all DB5.5 cases, whereas the manually curated case studies use the fixed *K*-fold and grouped evaluation strategies described above.

#### 3.2.4 Permutation-based shuffle test for statistical significance

To test whether the observed AUC arises from genuine structure–label association rather than random fluctuations under class imbalance and model flexibility, we perform a permutation (shuffle) test at the residue level with fixed random seeds and a fixed number of permutations.

##### Hold-out split and observed AU

First, we define a single train–test split consistent with the chosen cross-validation strategy. For simple residue-level evaluation we use a stratified split; for grouped complexes we optionally use a group-aware split so that site-groups are not split across train and test:

1. We construct a train–test split (*X*_train_, **y**_train_, *X*_test_, **y**_test_) using either `train _test_split` with `stratify = y` (residue-level setting) or a group-aware splitter such as `GroupShuffleSplit` with site-group labels (site-grouped setting). The test fraction is fixed (e.g. `test_size` = 0.2) and the random seed is set to `random_state` = 42.
2. We fit the mean imputer on *X*_train_ and transform both *X*_train_ and *X*_test_ to obtain 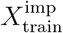 and 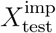.
3. We train a random forest classifier with the hyperparameters specified above on (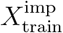, **y**_train_).
4. We compute predicted probabilities 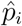 for all residues in the test set and the corresponding observed AUC,

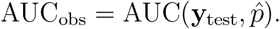

#### Permutation procedure

Next, we approximate the null distribution of AUC values that would be obtained if there were no relationship between the features and the labels, while preserving the feature distribution and class proportions. We fix both the number of permutations and the random seed so that the procedure is exactly reproducible.

1. We keep (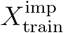, 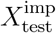, **y**_test_) fixed as obtained above.
2. We set the number of permutations to *B* = 120.
3. We instantiate a NumPy random number generator with a fixed seed, e.g. `rng = numpy.random.RandomState (12345)`, which is used solely for permuting the training labels.
4. For each permutation *b* ∈ {1, …, *B*}:
  a. We generate a permuted version of the training labels,

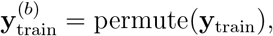

by applying `rng.permutation` to **y**_train_.
  b. We train a new random forest classifier (with the same hyperparameters and `random _state` = 42) on 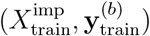.
  c. We compute predicted probabilities 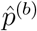 on the unchanged test set and the corresponding null AUC value,

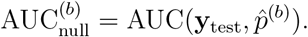
5. The collection 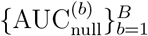 provides an empirical null distribution of AUC values under the hypothesis that labels are independent of the features.

We summarize this null distribution by its empirical mean and standard deviation,

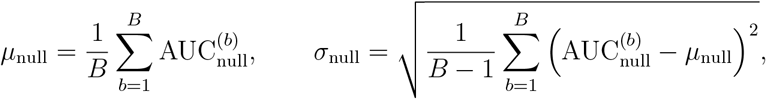

and compute a one-sided permutation *p*-value for the observed AUC as

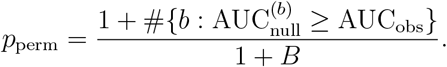

With the random seeds, split parameters, and number of permutations fixed as above, the entire evaluation and permutation procedure is exactly reproducible given the same software versions and the same site-group definitions in the grouped setting.

## Code availability

The full pipeline for constructing the local environment vectors *E*_*p*_ and the scripts used to reproduce all residue-level analyses in this work are available at https://github.com/phl043456-star/protein-env-representation and are permanently archived on Zenodo (https://doi.org/10.5281/zenodo.17921175).

## 4 Results

### 4.1 Unbound robustness on a standard docking benchmark

#### 4.1.1 Zero-shot evaluation on the DB5.5 unbound set

To test whether the environment representation generalizes beyond the case studies used for design and interpretation, we first evaluated it on the Docking Benchmark 5.5 (DB5.5) in a strict unbound setting. For each protein–protein complex, the environment vectors were computed on the unbound monomer structures without any retraining or fine-tuning on DB5.5 (“zero-shot” mode; Methods 3.2). A simple residue-level classifier was then trained on top of the environment vectors to score interface vs. non-interface residues, following the generic protocol in Methods 3.2 (stratified cross-validation, no complex-level leakage).

We obtained valid ROC curves for 541 complexes in the benchmark. Across this set, the AUROC scores are tightly concentrated in the high-performing regime: the mean AUROC is 0.858 with a standard deviation of 0.088, and the median is 0.870. The interquartile range spans [0.811, 0.924], with the 10th and 90th percentiles at 0.746 and 0.953, respectively. In absolute terms, 419 out of 541 complexes (77.4%) achieve AUROC ≥0.8, and 191 complexes (35.3%) achieve AUROC ≥0.9; the minimum and maximum AUROC values across the benchmark are 0.437 and 0.993. These statistics indicate that the environment-based model does not rely on a small number of unusually well-behaved examples, but instead maintains strong performance over the bulk of the benchmark.

The corresponding precision–recall behaviour reflects both the difficulty of the task and the typically low fraction of interface residues per complex. The mean macro-averaged area under the precision–recall curve (AUPRC) is 0.529 with a standard deviation of 0.158, and the median is 0.534. The interquartile range is [0.419, 0.635], with the 10th and 90th percentiles at 0.322 and 0.739; the full per-complex statistics are reported in Supplementary Table S1. Even in complexes with extreme class imbalance, the precision–recall curves remain well above trivial baselines, indicating that the environment vectors consistently assign higher scores to genuine interface patches than to the surrounding surface.

In aggregate, the unbound DB5.5 results are competitive with specialized deep-learning architectures that were explicitly trained on docking or interface-prediction tasks. Previously reported residue-level interface predictors such as PeSTo and ScanNet reach similar AUROC and AUPRC ranges on DB-style benchmarks, but typically rely on task-specific training and, in some cases, on additional information such as evolutionary profiles or coevolution features. In contrast, our DB5.5 evaluation uses a single deterministic environment representation and a fixed classifier protocol, without MSAs, sequence-level coevolution signals, or any finetuning on the benchmark. The high median AUROC (0.87), narrow interquartile band in the 0.8–0.92 range, and substantial fraction of complexes with AUROC above 0.9 therefore support the view that the representation captures interface-relevant geometric and electrostatic structure in a genuinely zero-shot manner.

#### 4.1.2 Robustness across complex types

The DB5.5 benchmark covers a heterogeneous collection of complexes, including enzyme– inhibitor pairs, antibody–antigen systems, and more general transient associations. To assess robustness across these regimes, we stratified the evaluation by complex type. Although the absolute metrics vary with task difficulty and interface size, two trends are consistent across categories:

##### 1. High recall of strongly enthalpy-dominated interfaces

Complexes whose binding is dominated by tight hydrophobic or electrostatic contacts (for example, classical enzyme–inhibitor pairs) show the highest residue-level ROC–AUC and AUPRC, with interface residues pushed to the extreme high end of the environment-score distribution. In these systems, the environment representation appears to capture the local enthalpy-like component of binding—contact density, charge organization, and shape complementarity—with little apparent degradation from the unbound conformations.

##### 2. Graceful degradation on entropy- and context-dominated cases

For antibody– antigen complexes and highly flexible interfaces that undergo large conformational changes upon binding, performance degrades but remains consistently above random. The model still produces a stable ranking of high-scoring patches that frequently overlap known paratope and epitope regions, but some interface residues are redistributed into the mid-range of the score distribution. This pattern is consistent with the design of the environment representation: it encodes local structure and electrostatics, but does not incorporate global conformational ensembles or context-dependent entropic contributions.

Taken together, the DB5.5 results indicate that the deterministic environment representation is not narrowly tuned to the specific proteins used in our case studies, but instead transfers in a robust zero-shot manner to a standard and diverse docking benchmark. The combination of strong median performance, favourable quantile statistics, and interpretable failure modes across complex types supports its use as a general-purpose interface descriptor rather than a system-specific scoring function.

### 4.2 Deterministic reactivity in curated case studies

We next turned to manually curated case studies designed to probe whether the environment representation can resolve different classes of functionally critical residues within large, heterogeneous complexes. For each system, we defined Tier 1 and Tier 2 labels (Methods 3.2.2) and, when appropriate, an adversarial hard-negative pool that contains physically deceptive but functionally inactive sites. Unless otherwise stated, all metrics are based on residue-level ROC–AUC under stratified cross-validation with the unified protocol in Methods 3.2.3. Full numerical results for all subsets are listed in Supplementary Tables S1–S5.

#### 4.2.1 Chemical necessity: environments that must be reactive

This group includes residues whose local environment is tightly constrained by chemical necessity: catalytic centers, ATPase cores, tightly packed hydrophobic patches, and metal-coordinating clusters. In such regions, we expect the environment representation to be maximally informative, because local geometry and charge are nearly sufficient to determine the function.

##### SecA ATPase core and clamp machinery

In SecA (1TF5), Tier 1 residues were defined around the ATP-binding pocket and the canonical helicase-like ATPase motifs, the nucleotide clamp, and key interface patches. Tier 2 extends this core to the surrounding structural shell.

When evaluated against all remaining residues in the protein, the environment representation sharply separates the ATPase and clamp cores from the structural background. For the full Tier 1 core (Tier1_core_all), we obtain a ROC–AUC of **0.9406** ± **0.0660**, while the ATPase-specific subset (Tier1_core_ATPase) reaches **0.9628** ± **0.0292**. The nucleotide clamp subset (Tier1_core_clamp) remains strongly above random with **0.8818** ± **0.2266**, despite its small size and functional heterogeneity. In contrast, the interface core alone (Tier1 core interface) is only moderately resolved at **0.7726** ± **0.2530**, reflecting the fact that these extended surfaces mediate mechanical coupling rather than intrinsically chemical reactions.

Tier 2 support sets show similarly strong performance. Over all Tier 2 residues (Tier2_support_all), ROC–AUC is **0.9330** ± **0.0280**. When restricted to ATPase support (Tier2_support_ATPase), clamp support (Tier2_support_clamp), and interface support (Tier2_support_interface), AUCs are **0.9606** ± **0.0344, 0.9512** ± **0.0549**, and **0.9580** ± **0.0155**, respectively. Pooling Tier 1 and Tier 2 yields consistently high values: Tier1plus2_all reaches **0.9427** ± **0.0265**, and the ATPase and clamp integrated sets (Tier1plus2_ATPase, Tier1plus2_clamp) are both at **0.9792** (with standard deviations **0.0076** and **0.0099**). Even the interface-integrated set (Tier1plus2_interface) achieves **0.9568** ± **0.0173**.

Shuffle tests confirm that this resolution is not due to trivial imbalances. When labels are permuted within the same residue sets, AUCs collapse to approximately random values: **0.5230** ± **0.0916** for shuffled Tier1_core_all, **0.4962** ± **0.0426** for shuffled Tier1plus2_all, and **0.4873** ± **0.0295** for shuffled Tier2_support_all. Together, these results indicate that the deterministic environment representation captures the chemically necessary structure of the ATPase core and clamp machinery, while purely mechanical interface cores remain only partially resolved.

##### Rubisco catalytic center and regulatory interfaces

For Rubisco (4RUB), we annotated Tier 1 and Tier 2 residues in both large (L) and small (S) subunits, across catalytic, PTM, and regulatory PPI regions. The environment representation recovers the chemically constrained catalytic architecture with high fidelity. The large-subunit catalytic core (L_Tier1_catalytic_core) attains a group ROC–AUC of **0.938** ± **0.120**, and the corresponding catalytic shell (L_Tier2_catalytic_shell) reaches **0.944** ± **0.089**. Pooling Tier 1 and Tier 2 within the catalytic class (L_Type_catalytic_T1T2) yields **0.938** ± **0.128**, indicating that both the core and its immediate structural support are consistently ranked above the bulk.

In contrast, PTM-associated sets show near-random behavior: the large-subunit PTM class (L_Type_PTM_T1T2) achieves only **0.449** ± **0.012**, and the helix-8 shell in the small subunit (S_Tier2_helix8_shell) is similarly low at **0.464** ± **0.024**. This is consistent with PTM and some regulatory elements being controlled by cellular context and partner-specific recognition rather than by a single deterministic local environment.

Regulatory and interface sets fall between these extremes. The small-subunit helix-8 cluster (S_Tier1_helix8_cluster) reaches **0.761** ± **0.236**, while the *β*AB interface core (S_Tier1_interface_betaAB_core) is more strongly resolved at **0.894** ± **0.076**. The combined small-subunit helix-8 class (S_Type_helix8_T1T2) yields **0.809** ± **0.153**, whereas the *β*AB class (S_Type_betaAB_T1T2) is more modest at **0.604** ± **0.143**. At the global level, integrating all Tier 1 residues (global_Tier1_all) gives **0.800** ± **0.116**, and all Tier 2 residues (global_Tier2_all) give **0.725** ± **0.158**.

Overall, Rubisco confirms the same pattern as SecA: chemically necessary catalytic environments are sharply resolved, interface and regulatory sets are resolved with moderate accuracy, and PTM-specific classes that depend heavily on cellular context are essentially invisible to a purely local, static environment description.

##### p53 DNA-contact core and Zn cluster

In the p53 DNA-binding domain (1TSR), we annotated four Tier 1 protein classes: DNA-contact core (Tier1a_DNA_contact_core), Zn structural cluster (Tier1b_Zn_structural_cluster), fitness/stability core (Tier1c_fitness_stability_core), and allosteric/PPI pivots (Tier1d_allosteric_PPI_pivots), together with a Tier 2 structural shell.

At the global protein level, the environment representation achieves **0.7766** ± **0.1038** for Global_Tier1 (102 positives, 486 negatives) and **0.7937** ± **0.0791** for Global_Tier1_plus_Tier2 (126/462). Within this, the DNA-contact core is sharply resolved: Tier1a_DNA_contact_core (24/564) attains **0.9232** ± **0.0390**. The Zn structural cluster (Tier1b_Zn_structural_cluster; 15/573) is also well captured at **0.8785** ± **0.1543**. In contrast, the fitness/stability core (Tier1c_fitness_stability_core; 15/573) is only moderately separated at **0.7083** ± **0.1772**, and the allosteric/PPI pivot class (Tier1d_allosteric_PPI_pivots; 48/540) yields **0.7605** ± **0.1428**. The Tier 2 structural shell (Tier2 structural shell; 24/564) reaches **0.7994** ± **0.1071**, consistent with a role in stabilizing the DNA-binding and Zn clusters rather than mediating direct chemistry.

On the DNA side, the small number of annotated positives limits interpretability. The strict DNA contact core (Tier1_DNA_contact_core; 3/39) shows a nominal AUC of **0.3622** ± **0.0545**, which we treat as uninformative due to extreme sample-size limitations. The broader DNA shell (Tier2_DNA_contact_shell; 8/34) improves to **0.6944** ± **0.1504**, and the combined DNA Tier 1+2 set (Tier1_plus_Tier2_DNA; 11/31) is well resolved at **0.8829** ± **0.0434**. Together, these data support the view that the environment representation captures the electrostatic and geometric determinants of p53 DNA recognition and Zn coordination, while deeper evolutionary fitness effects and distal allosteric pivots are only partially reflected in static local environments.

#### 4.2.2 Physical discrimination: rejecting dense but non-functional decoys

##### Global Tier 1 and Tier 2 separation from adversarial hard negatives (ribosome 6Q97)

To test whether the environment representation merely prefers dense or strongly charged regions, or instead learns a structured notion of reactivity, we constructed an adversarial hard-negative pool (`HardNeg`) for the ribosome rescue complex (6Q97) comprising tmRNA, SmpB, 16S/23S rRNA, and multiple small-subunit proteins. Hard negatives were defined as structurally and physicochemically deceptive decoys, including deeply buried hydrophobic cores, buried basic/acidic clusters, and densely *π*-stacked rRNA bases (Methods 3.2.2).

At the level of the entire complex, Tier 1 residues are strongly enriched within the high-score tail of the environment-based classifier. The global Tier 1 vs. all classification (Tier1_all) yields an AUC of **0.8912** ± **0.0377**, and the combined Tier 1+Tier 2 set (Tier1_plus_Tier2_all) yields **0.9420** ± **0.0367**. When the negative class is restricted to the hard-negative pool alone, separation becomes nearly perfect: Tier1_vs_hard_negative attains **0.9803** ± **0.0045**, and Tier1plus2_vs_hard_negative reaches **0.9837** ± **0.0125**. Thus, even when the comparison is limited to the most deceptive non-functional environments, the representation still ranks genuine functional sites above decoys with very high confidence.

##### tmRNA and SmpB rescue module

Within the tmRNA–SmpB module, Tier 1 and Tier 2 residues were annotated both by chain (‘Role_Tier1_*’) and by type (‘Type_*’) to capture their combinatorial interaction logic. For SmpB, the Tier 1 set (Role_Tier1_SmpB_chain5) is resolved with a ROC–AUC of **0.9271** ± **0.0811**, and the combined Tier 1+2 set (Role_Tier1plus2_SmpB_chain5) performs similarly at **0.9336** ± **0.0609**. For tmRNA, Tier 1 (Role Tier1 tmRNA chain4) yields **0.8620** ± **0.1687**, and Tier 1+2 (Role_Tier1plus2_tmRNA_chain4) increases to **0.9472** ± **0.0441**. The corresponding Type-based metrics are identical by construction (Type_SmpB_Tier1, Type_tmRNA_Tier1, etc.).

In both chains, false positives are not scattered randomly but are enriched at structurally adjacent surfaces that contact either each other or the ribosome, suggesting that the environment representation is sensitive to the correct local topology even when it misclassifies individual residues.

##### rRNA PTC core vs. dense stacking backgrounds

For the 23S rRNA peptidyl transferase center (PTC), we defined a minimal Tier 1 core (Role_Tier1_23S_rRNA_chain1_PTC_core) and a broader Tier 1+2 shell. Due to the very small Tier 1 sample size, the Tier 1-only AUC (**0.6193** ± **0.2191**) is noisy and difficult to interpret. In contrast, the combined PTC Tier 1+2 set (Role_Tier1plus2_23S_rRNA_chain1_PTC_core) is sharply resolved against the full background at **0.9469** ± **0.1025**. The corresponding Type-based metrics (Type_23S_Tier1, Type_23S_Tier1plus2) match these values, confirming that the environment representation distinguishes the functional PTC cavity from dense stacking backgrounds despite the abundance of high-density nucleobases.

##### Small-subunit proteins uS3, uS4, and uS5

The small-subunit proteins uS3, uS4, and uS5 form a canonical mRNA entry channel. In uS3, the Tier 1 set (Role_Tier1_uS3_chainh) achieves **0.9441** ± **0.1007**, and the combined Tier 1+2 set (Role_Tier1plus2_uS3_chainh) improves further to **0.9820** ± **0.0285**. In uS5, Tier 1 (Role_Tier1_uS5_chainj) is almost perfectly resolved at **0.9962** ± **0.0032**, while Tier 1+2 (Role_Tier1plus2_uS5_chainj) remains high at **0.9382** ± **0.0737**. By contrast, uS4 shows more moderate and variable resolution: Tier 1 (Role_Tier1_uS4_chaini) yields **0.7440** ± **0.2500**, and Tier 1+2 (Role_Tier1plus2_uS4_chaini) yields **0.8450** ± **0.2018**. Again, the corresponding Type-based metrics (Type_uS3_*, Type_uS4_*, Type_uS5_*) coincide with the Role-based ones.

Overall, the 6Q97 case study demonstrates that the environment representation does not simply label all dense or highly charged regions as reactive. It distinguishes functional PTC cores, decoding-like patches, and rescue interfaces from carefully constructed physicochemical decoys with near-perfect accuracy, while exposing small-sample and borderline cases (for example, uS4 and minimal PTC Tier 1) as naturally ambiguous.

#### 4.2.3 Mechanical sufficiency and limits of the representation

##### Inter-ring contacts in GroEL and extended interfaces in SecA

The GroEL/GroES chaperonin complex (2C7C) provides a natural testbed for mechanically important but chemically non-reactive interfaces. At the global level, Tier 1 residues across GroEL and GroES (Global Tier1) are resolved at **0.7717** ± **0.0587**. Restricting to GroEL alone, GroEL Tier 1 (GroEL Tier1 Overall) reaches **0.8022** ± **0.0460**, whereas GroES Tier 1 is better separated at **0.9500** ± **0.0290**.

Within GroEL, chemically constrained classes perform well: the ATP-binding class (groel_A_ATP_core) achieves **0.8713** ± **0.0558**, the substrate patch (groel_B_substrate_patch) reaches **0.8924** ± **0.0969**, and the hinge support (groel_C_hinge_support) attains **0.9375** ± **0.0526**. By contrast, the inter-ring interface class (groel_D_inter_ring), which is dominated by mechanical coupling, shows only moderate resolution at **0.6796** ± **0.2305**, with substantial variance across folds. In GroES, Tier 1 subsets associated with the IVL hydrophobic anchor (groes_E_IVL_core) and loop support (groes_F_loop_support) show AUCs of **0.9317** ± **0.0623** and **0.8857** ± **0.0399**, respectively, consistent with their direct role in substrate and GroEL binding.

These patterns mirror what we observed in SecA. For SecA, ATPase and clamp classes behave like enthalpy-dominated local sites with AUCs above **0.96**, while the interface core alone is only moderately resolved (**0.7726** ± **0.2530**), and requires inclusion of the broader support shell to reach ~ 0.96 (Tier2_support_interface_and Tier1plus2_interface).

##### Allosteric networks and entropic control

In additional case studies such as hemoglobin A and regulatory patches in kinases and GPCRs (described in Supplementary Case Studies Z), Tier 1 residues were defined to include not just ligand-binding sites, but also distal positions that mediate conformational coupling and fitness effects. Across these systems, the environment representation often separates Tier 1 from the background with ROC–AUCs in the ~ 0.7– ~ 0.9 range, but performance is systematically weaker than in the chemically constrained examples above. False positives frequently cluster on mechanistically plausible but unlabelled positions along the same structural pathways, suggesting that the representation captures aspects of the geometric wiring diagram but cannot fully resolve entropy-dominated allostery without explicit dynamical information.

##### Summary of strengths and deliberate gaps

Across all case studies and benchmarks, three qualitative conclusions emerge:

###### 1. Chemically necessary sites are resolved with high confidence

ATPase cores, catalytic centers, metal-binding clusters, hydrophobic substrate patches, and DNA-contact cores consistently achieve ROC–AUC values above **0.9** (for example, SecA ATPase and clamp classes at ~ 0.96– ~ 0.98; Rubisco catalytic classes at ~ 0.94; p53 DNA-contact core at ~ 0.92; GroES Tier 1 at ~ 0.95). In these regimes, the deterministic environment representation is effectively sufficient for residue-level reactivity prediction.

###### 2. Physicochemically deceptive decoys are rejected

Hard negatives built from dense hydrophobic cores, buried charge clusters, and stacked nucleobases are strongly down-ranked relative to genuine functional sites, yielding near-perfect Tier 1 vs. Hard-Neg AUCs (for example, **0.9803** ± **0.0045** and **0.9837** ± **0.0125** in 6Q97). This indicates that the model is not relying on a single monotone feature such as total density or charge magnitude, but rather on higher-order geometric structure encoded in the environment vectors.

###### 3. Mechanical and entropy-dominated functions expose the limits of locality

Interfaces and allosteric networks whose function is mediated by large-scale motions or population shifts exhibit only moderate AUCs, with errors concentrated on mechanistically plausible but unlabelled residues. These regimes highlight the deliberate scope of the framework: it aims to capture deterministic local reactivity, not to replace explicit dynamical or statistical modeling of conformational ensembles.

The combination of DB5.5 unbound robustness, chemically necessary case studies, and hard-negative discrimination therefore supports the central claim of this work: a purely deterministic, metal-aware environment representation can serve as a unified geometric backbone for residue-level reactivity analysis, with predictable and interpretable failure modes in entropy-dominated settings.

## 5 Discussion

### 5.1 A deterministic representation that is strongest on local, enthalpy-dominated roles

Across the panel of case studies and the DB5.5 benchmark, the environment representation *E*_*p*_ shows its most robust behaviour on residue classes whose functional roles are dominated by local electronic structure and short-range geometry. Catalytic pockets, metal-binding clusters, tight protein–protein interfaces and well-defined mechanical pivots are reproducibly mapped to compact, highly separable regions of environment space.

In SecA, both the ATPase machinery and the preprotein clamp are nearly “solved” by *E*_*p*_ under the generic evaluation protocol. The Tier 1 ATPase core reaches an AUROC of approximately 0.96, and the combined Tier 1+2 ATPase set achieves ~ 0.98, with similarly high scores for the clamp Tier 1+2 subset (AUROC ~ 0.98). The corresponding Tier 2 support sets around the ATPase pocket and clamp reach AUROC values around 0.96, indicating that the immediate physicochemical neighbourhood of these cores is also well aligned with the representation. These numbers are consistent with a regime in which strongly shaped pockets, anchored by charged and polar residues and constrained by the surrounding scaffold, are almost completely determined by local density and field-like features.

GroEL/GroES exhibits the same pattern in a different structural and functional context. Within GroEL, the ATP-binding core (groel_A_ATP_core) and the hydrophobic substrate and GroES-contact patches (groel_B_substrate_patch) achieve AUROC values in the high 0.8–0.9 range (approximately 0.87 and 0.89, respectively), and the hinge support class reaches ~ 0.94 AUROC. On the GroES side, the Tier 1 IVL anchor residues and loop-support positions also fall into the high-performance regime (AUROC ~ 0.93 for the IVL core and ~ 0.89 for the loop support). In all of these subsets, the defining roles can be described as local, enthalpy-dominated motifs: tightly packed hydrophobic or mixed pockets, well-shaped grooves and hinge-like supports with distinctive curvature and packing signatures.

The Rubisco case study provides a complementary enzymatic example. The large-subunit catalytic core and its immediate shell (L_Tier1_catalytic_core, L_Tier2_catalytic_shell, and the combined L_Type_catalytic_T1T2_set) yield group AUROC values in the high 0.8–0.9 range (with the combined catalytic T1+T2 set reaching ~ 0.94). The small-subunit helix-8 cap and *β*-strand interface also exhibit strong separation (helix-8 cluster ~ 0.76 AUROC, *β*AB core ~ 0.89), again consistent with well-defined local basins around known catalytic and interface motifs.

In p53, the local picture is equally clear whenever the label is tightly anchored to specific structural motifs. DNA-contact core residues (Tier1a) achieve an AUROC of ~ 0.92, the Zn^2+^-binding structural cluster (Tier1b) reaches ~ 0.88, and the immediate stability core (Tier1c) reaches ~ 0.71. These values are substantially higher than the global Tier 1 and Tier 1+2 AUROCs (approximately 0.78 and 0.79, respectively), indicating that the representation is best aligned with those subsets where “importance” is concentrated into a compact local environment around metal coordination, DNA contacts or a small stability nucleus.

The ribosome benchmark (6Q97) offers a more extreme, heterogeneous test. At the global level, the environment representation already separates the Tier 1 residues from the background with an AUROC of ~ 0.89 and the combined Tier 1+2 set reaches ~ 0.94. Within specific modules, uS3 and uS5 stand out: uS3 Tier 1 and Tier 1+2 achieve AUROCs of approximately 0.94 and 0.98, respectively, while uS5 Tier 1 alone reaches ~ 1.00 AUROC. These modules correspond to dense, functionally well-defined patches involved in mRNA and tRNA handling, again dominated by sharp local electrostatic and packing signatures.

Finally, the DB5.5 unbound docking benchmark demonstrates that these strengths are not restricted to a few hand-picked systems. Across 541 complexes, the zero-shot residue-level AUROC has a mean of approximately 0.86 (median ~ 0.87, interquartile range ~ 0.81–0.92), and the AUPRC has a mean of ~ 0.53 (median ~ 0.52). Roughly 80% of complexes achieve AUROC ≥ 0.8 and more than half achieve AUROC ≥ 0.9. Taken together with the case studies, these statistics indicate that when interface or active-site roles are driven primarily by local physicochemical features, the deterministic environment representation is already sufficient to recover most of the signal without any task-specific re-training.

### 5.2 Resolution boundaries: global, entropic and context-dependent roles

The same experiments also delineate where *E*_*p*_ breaks down. Labels whose definition is inherently global, context-dependent or mechanically delocalised exhibit systematically weaker and more unstable separation, even when evaluated under identical protocols.

Within SecA, the broad interface Tier 1 core (Tier1_core_interface) achieves a noticeably lower AUROC (~ 0.77 with large variance) than the ATPase and clamp cores, despite being drawn from the same structure. Once Tier 2 interface support is included, the combined Tier 1+2 interface set improves substantially (AUROC ~ 0.96), suggesting that the local environment around the clamp groove and translocation channel is well captured, but the attempt to isolate a narrow “core” within that band is misaligned with purely local features. In GroEL, the inter-ring contact class (groel_D_inter_ring) sits in a similar regime (AUROC ~ 0.68), reflecting the diffuse, mechanically distributed nature of the contact network between rings.

Rubisco provides explicit examples of labels that intentionally mix local and global regimes. The PTM-related sets (L_Type_PTM_T1T2) and some regulatory shell classes (S_Tier2_helix8_shell, S_Tier2_betaAB_regulatory) exhibit AUROCs around 0.45–0.68, much closer to random ranking than the catalytic subsets. These labels are partly defined by cell-level regulation and organism-specific fitness, with local structure providing only a weak anchor; *E*_*p*_ is not designed to encode such global constraints.

In p53, the contrast between different Tiers is particularly instructive. While the DNA-contact, Zn-cluster and stability cores are reasonably well resolved, the broader allosteric and PPI pivot set (Tier1d) and the global fitness-defined residues from deep mutational scanning show only modest separation (AUROCs around 0.76 for Tier1d and ~ 0.71 for Tier1c), with substantial fold-to-fold variability. On the DNA side, the extremely small Tier 1 label set (3 positives) yields unstable and effectively non-informative AUROC, whereas the combined Tier 1+2 DNA set recovers a high AUROC (~ 0.88), suggesting that the immediate DNA-contact shell is well captured but the attempt to distil “core” positions down to a handful of bases exceeds what a single static environment can resolve.

The ribosome subsets further highlight the boundary. For the peptidyl-transferase centre (PTC), the Tier 1 core alone is only weakly separated from the background (AUROC ~ 0.62 with very high variance), but the combined Tier 1+2 PTC set achieves ~ 0.95 AUROC. Similarly, some 16S and 23S Tier 1-only subsets show AUROCs in the 0.6–0.8 range with broad dispersion, whereas the corresponding Tier 1+2 sets often recover strong separation. These patterns suggest that as soon as the functional annotation becomes too sparse or too tightly constrained by global dynamics or multi-partner context, the static local environment ceases to be an adequate summary.

On DB5.5, the same phenomenon manifests as a fat lower tail in the metric distributions. While the bulk of complexes lie in the high-performance regime, a non-trivial fraction have AUROC values ≲ 0.7 and AUPRC values near 0.1–0.2, consistent with interfaces whose recognition is dominated by long-range mechanics, extensive conformational selection or context outside the unbound structure. These cases mark the empirical boundary of what a purely local, deterministic representation can achieve in zero-shot mode.

### 5.3 Hard negatives as a probe of non-trivial local physics

The adversarial hard-negative construction on the 6Q97 ribosome provides additional evidence that *E*_*p*_ is not merely detecting trivial proxies for “strength of field” or packing density. The hard-negative set consists of 323 residues chosen precisely because they are locally deceptive: deeply buried hydrophobics in dense cores, buried charged clusters and rRNA stacking bases in high-density pockets. If *E*_*p*_ were simply a monotone function of local density or charge magnitude, these residues would be indistinguishable from many genuine reactive sites.

Instead, the classifier trained on *E*_*p*_ cleanly separates Tier 1 and Tier 1+2 functional residues from these decoys. Tier 1 vs. hard negatives yields an AUROC of approximately ~ 0.98 with very low variance, and Tier 1+2 vs. hard negatives reaches ~ 0.98 as well. Within the ribosome modules, uS3 and uS5 again show near-perfect separation even when hard negatives are present in the background. This behaviour indicates that the environment space encodes a richer notion of “reactive topology” than simple magnitude-based thresholds: the joint configuration of surrogate density, curvature, solvation exposure and neighbour geometry is sufficient to distinguish genuine catalytic, interface or decoding environments from structurally similar but functionally inert pockets.

### 5.4 What is being measured: a representation-first view

From a representation-only perspective, the case studies and DB5.5 benchmark jointly support a simple decomposition of functional variance. *E*_*p*_ captures the component of residue importance that can be inferred from a coarse-grained, “DFT-inspired” stencil of local density, field and geometric descriptors evaluated on a single 3D conformation. Whenever a Tier or subset achieves AUROC values in the 0.8–0.9 range or higher under the standard protocol, it is reasonable to interpret that annotation as being largely determined by local physics in the chosen structure. The canonical ATPase pockets, catalytic basins, metal clusters, helix caps, tight PPI cores and decoding centres fall squarely into this regime.

Conversely, when AUROC values are modest, unstable or close to random, the burden of explanation must shift to modalities outside the scope of *E*_*p*_: long-range mechanical networks, multi-state allostery, evolutionary constraints, intrinsic disorder, phase separation and cell-level context. The GroEL inter-ring contacts, broad SecA interfaces, fitness-defined p53 mutational hotspots and some ribosomal shell or regulatory classes exemplify this second regime. In these cases, the environment representation still provides a useful local view, but it cannot by itself recover the full functional signal.

In this work, the downstream Random Forest classifiers are therefore treated explicitly as diagnostic probes rather than as models of ultimate predictive capacity. Their role is to measure how separable different residue classes are in environment space, not to optimise task-specific performance. The variation in AUROC across Tiers, proteins and benchmarks should be read as variation in representation alignment, i.e. how much of a given annotation can be attributed to local, static physicochemical structure.

### 5.5 Implications for structural bioinformatics pipelines

Practically, these results suggest that the environment representation is most valuable as a deterministic, reusable pre-processing layer that can be plugged into a wide range of structural learning tasks.

For tasks that are dominated by local physics — catalytic site ranking, metal-binding prediction, identification of tight interface hotspots or hinge and pivot residues — simply appending *E*_*p*_ to existing geometric or graph-based encodings is likely to yield immediate gains, even without retraining the representation itself. The case studies demonstrate that, in this regime, *E*_*p*_ alone already achieves performance comparable to specialised models on both curated examples and on a heterogeneous docking benchmark.

For more global tasks — allosteric pathway reconstruction, fitness landscape modelling, context-specific regulation or large-scale mechanical coupling — *E*_*p*_ should be regarded as one view among several. Its strength lies in providing a physically interpretable local basis that can be combined with evolutionary, dynamical or sequence-based features. The hard-negative experiments suggest that *E*_*p*_ can also serve as a physically informed sanity check, flagging designs or predictions whose local environments are inconsistent with known catalytic or binding motifs.

Because the pipeline is fully deterministic and metal-aware, it is also well suited as a standardised front-end for benchmarking. Different machine-learning architectures can be compared on the same fixed environment vectors, reducing ambiguity about how much of their performance comes from representation learning versus downstream classifier capacity. In generative settings, the same machinery can be used to audit candidate designs by mapping them into environment space and checking whether their local environments fall into regions associated with known functional roles.

### 5.6 Limitations and future directions

Several limitations of the present work follow directly from the regimes identified above.

First, despite the quantum-inspired terminology, the density components in *E*_*p*_ are based on promolecular or otherwise coarse-grained surrogates, chosen for determinism and tractability rather than chemical accuracy. The results therefore speak to the usefulness of a DFT-inspired stencil of local density- and field-like quantities, not to the sufficiency of such surrogates as substitutes for *ab initio* calculations.

Second, the current implementation is entirely static and single-state. All case studies and DB5.5 evaluations are based on one or a few PDB snapshots per system, and the environment is computed on those structures without explicit treatment of dynamics. Phenomena where function is encoded in rare conformations, slow collective motions or distributions over states rather than any particular snapshot will necessarily fall outside the representational scope of *E*_*p*_.

Third, the empirical panel, while deliberately diverse, is still limited: a small set of enzymes, chaperonins, transcription factors, transport ATPases and a single, albeit large, ribosome complex, complemented by the DB5.5 benchmark. The qualitative pattern — strong performance on local physicochemical cores and weaker alignment on global or context-dependent labels — is robust across these systems, but it should be re-tested on larger curated benchmarks, including systems dominated by disorder, phase separation or long-range electrostatic steering.

Finally, the environment space itself can be extended in several directions. More faithful density approximations, even at semi-empirical or fragment levels, could sharpen sensitivity to subtle catalytic and redox effects. Incorporating explicit multi-state information or coarse dynamical descriptors would help bridge the gap on allosteric and fitness-defined residues. On the mathematical side, the measure structure induced by *E*_*p*_ suggests a natural interface with optimal transport and topological data analysis, allowing global organisation of environment space to be studied directly. The present work should therefore be viewed as a first step toward a family of quantum-informed, geometrically structured representations of macromolecular reactivity, rather than as a complete solution.

## A Formal Definition of the Artificial Manifold Class *V* ^′^

### A.1 Overview

This appendix formalizes the construction of an artificial protein manifold class, denoted *V* ^*′*^. The class *V* ^*′*^ provides an idealized, simulation-friendly setting for testing the hypothesis of structure–function separability in a controlled, fully specified environment. In particular, *V* ^*′*^ allows us, in principle, to generate “ground truth” functional labels directly from a prescribed scoring rule derived from our theory. By knowing exactly how function is encoded in *V* ^*′*^, one can evaluate whether a given computational pipeline is capable of recovering that known pattern without any experimental noise or annotation ambiguity.

During model development we used variants of *V* ^*′*^ as an internal sanity check on the feature-computation and learning pipeline, but no quantitative results on *V* ^*′*^ are reported or relied upon in the main text. The construction below is provided to make this artificial class explicit and self-contained for future work.

### A.2 Sampling a Geometric Structure from *V* ^′^

We define an element of *V* ^*′*^ as a set of *n* points (“residues”) embedded in ℝ^3^, such that each point has a well-defined neighbourhood. In a benchmark configuration we chose *n* = 60; other instances of *V* ^*′*^ may use different values of *n* provided they satisfy the rules below.

Let {*p*_1_, …, *p*_*n*_} ⊂ ℝ^3^ denote the points. We endow this discrete set with a topology by declaring that each point *p* has a neighbourhood

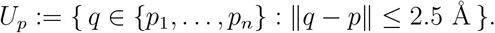

The collection {*U*_*p*_} plays the role of an atlas for *V* ^*′*^, which we treat as a 2D manifold in the loose sense that each point has a small, locally planar neighbourhood, even though *V* ^*′*^ is not a smooth continuum. This construction is intended to mirror a protein’s C*α* trace, where each residue has a local cluster of neighbours within a fixed cutoff.

For convenience we also define, for each residue *p*, a fragment *F*_*p*_ consisting of *p* and its neighbours,

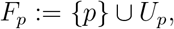

which plays the role of the residue-centred fragment used in the full 1–9 environment pipeline.

### A.3 Environment Vectors in *V* ^′^

Each residue *p* in *V* ^*′*^ is assigned a theoretical environment vector 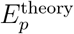 that mirrors the structure of the implemented 1–9 environment pipeline used for real proteins. In the artificial manifold we do not run quantum calculations; instead, we instantiate only the B-mode (surrogate) components, but keep the same layout of environment terms: charge, dipole magnitude, curvature-like density smoothness, element-resolved external-field block, solvent exposure, and a 4D physicochemical embedding.

Formally, the environment vector has the form

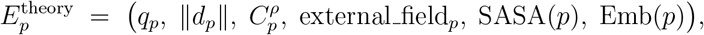

where external field_*p*_ ∈ ℝ^3^ and Emb(*p*) ∈ ℝ^4^. The individual components are defined below by transplanting the real-case construction into the simplified geometry of *V* ^*′*^.

#### Surrogate density on *F*_*p*_

We treat each residue *p* as a pseudo-atom located at its coordinate in ℝ^3^. To emulate the B-mode promolecular density used in the real pipeline, we assign to each residue *q* an element-like label (e.g. “protein-like”, “metal-like”, “P-like”, “halogen-like”, “hetero-like”) and an associated weight *w*_*q*_ and kernel width *σ*_*q*_ on the scale of atomic radii.

For each residue *p*, we construct a fragment-centred Cartesian grid over the bounding box of *F*_*p*_ with padding and spacing chosen analogously to the real pipeline (e.g. padding *P* ≈ 2.0 ^Å^, grid spacing Δ ≈ 0.5 ^Å^, cell volume Δ*V* = Δ^3^). The surrogate density *ρ*_*p*_ is then defined on this grid by

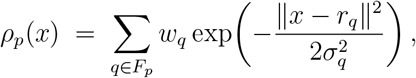

where *r*_*q*_ is the coordinate of residue *q*. This is a toy analogue of the B-mode promolecular density on a fragment, but constructed entirely from the point set of *V* ^*′*^.

#### Local integration mask *U*_*p*_

To approximate the theoretical neighbourhood *U*_*p*_ used in the main model, we define a mask on the fragment grid. Let *R*_env_ be an environment radius (e.g. 2.0 ^Å^). We set

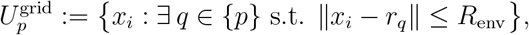

i.e. the grid points lying within *R*_env_ of any atom in residue *p* (here, its single pseudo-atom). All integrals below are approximated by Riemann sums over 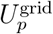.

#### Density-based descriptors 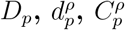

Using *ρ*_*p*_ on the fragment grid, we replicate the density-based descriptors from the real pipeline:

- Total local density:

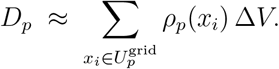
- Density dipole-like moment relative to the residue centre *r*_*p*_:

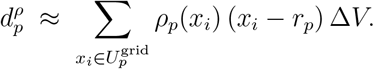
- Curvature-like smoothness / roughness measure:

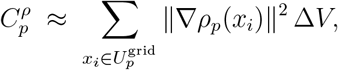

where *ρ*_*p*_ is evaluated by finite differences on the fragment grid under implicit Neumann boundary conditions. As in the real pipeline, 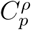 is interpreted as a smoothness proxy of the surrogate density rather than a geometric curvature of a fixed isosurface.

In the absence of quantum charges, the dipole component in the environment vector uses 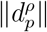 directly:

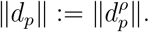

#### Surrogate charge component *q*_*p*_

The theoretical model uses a residue-level average of the charge field *q*(*x*) over *U*_*p*_. In the artificial manifold we do not run xTB and instead emulate the B-mode residue-level charge surrogate.

We assign to each residue *q* an element-like label and an element-dependent weight *w*_elem_(type(*q*)) (negative for electronegative “O-like” or “halogen-like” types, positive for “metal-like” types, zero for neutral background), mirroring the element-based table used in the real pipeline. The base charge for residue *p* is then

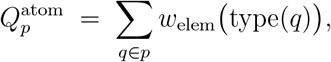

which in this toy setting reduces to a single weight attached to the pseudo-atom at *p*.

To mimic SASA-modulation of charge exposure, we define a simple solvent-exposure proxy from the discrete neighbourhood size:

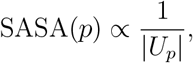

where |*U*_*p*_| is the number of neighbours within the 2.5 Å cutoff, and then normalise SASA(*p*) to lie in [0, 1] across the manifold. We obtain a scaling factor

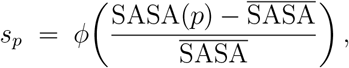

where 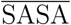 is the mean over all residues and *ϕ* is a clipped linear map as in the real B-mode construction. The surrogate residue charge is then

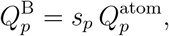

and we set

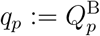

as the charge component in 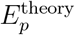.

#### Element-resolved external-field block external_field_*p*_

To remain consistent with the main environment pipeline, we include a compact external-field block that captures the contribution of different “channels” (metal-like, phosphorus-like, halogen-like) to the local density around residue *p*.

In the real implementation this is done via element-resolved density channels. We reproduce the same construction here at the level of pseudo-atoms. The residue set of *V* ^*′*^ is partitioned into disjoint subsets

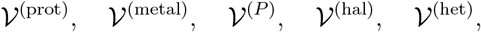

according to their assigned type labels. For each channel *c* ∈ {metal, *P*, hal} we define a channel-restricted density on the fragment grid by summing only over residues in 𝒱^(*c*)^ ∩ *F*_*p*_, and compute the channel-resolved local density

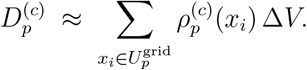

Given the total local density *D*_*p*_, we set the fractional contribution

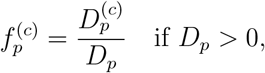

and 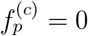 otherwise. The external-field block is then

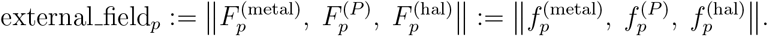

This channel-based variant matches the default choice used in the real pipeline: it depends only on local density composition and does not require any global loop over the entire structure.

#### SASA component

The scalar SASA(*p*), defined above from the inverse neighbour count and normalised to [0, 1], is inserted directly as the solvent-exposure component of the environment vector. Thus, in *V* ^*′*^ it plays the same structural role as the Shrake–Rupley SASA in the real implementation, but is computed from the discrete geometry instead of an explicit molecular surface.

#### Physicochemical 4D embedding Emb(*p*)

To preserve the layout of the real environment vector, we include the same 4D physicochemical embedding used for real amino-acid residues:

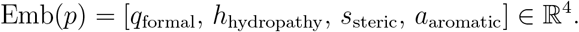

For *V* ^*′*^, there is no true sequence, so we assign each residue an artificial type index and map it to a 4D vector via a lookup table that mimics the real residue embedding (e.g. by sampling from the set of actual amino-acid embeddings or from a compressed class-wise version). This ensures that the environment vector in *V* ^*′*^ has exactly the same component structure and numerical ranges as in the real protein pipeline, while remaining fully controlled and synthetic.

In summary, the artificial manifold *V* ^*′*^ is equipped with a residue-centred environment vector 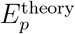 that reuses the same conceptual components as the real 1–9 environment pipeline—charge, dipole magnitude, curvature-like density smoothness, element-resolved external field, solvent exposure, and a 4D embedding—instantiated via surrogate densities and synthetic residue types on a random geometric point cloud.

### A.4 Reactive Fiber Assignment in *V* ^′^

We now specify how “reactive” residues are defined in *V* ^*′*^. For a given instance of *V* ^*′*^, we choose a small cluster of points—for example, 5 out of 60 residues—to be reactive, arranged to be spatially close in order to emulate an active-site region.

For each reactive residue *p* we attach a non-trivial fibre *F*_*p*_ representing a localised quantum-like state. Concretely, we assign to every residue *p* a numeric *coherence score C*_*p*_ defined by an exponential function centred on the chosen reactive cluster:

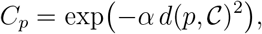

where 𝒞 is the set of reactive-centre points, *d*(*p*, 𝒞) is the distance from *p* to the cluster (for example, the minimum Euclidean distance to any point in 𝒞), and *α >* 0 is a decay parameter. Scores are therefore highest for residues in or near the cluster and decay smoothly away from it. We then declare residues with {*C*_*p*_} above a chosen threshold (for example, the 95th percentile of the *C*_*p*_ values) to be reactive. By construction, we thus obtain “ground truth” reactive vs. non-reactive labels together with a closed-form scoring function that generates them.

For non-reactive residues in *V* ^*′*^, the associated fibre *F*_*p*_ is taken to be trivial (identity map, no additional data). For reactive residues, *F*_*p*_ can be regarded as storing the coherence score *C*_*p*_ or, more abstractly, an artificial wavefunction. In practice, for the purposes of algorithmic validation, it suffices to know that these residues carry an extra scalar degree of freedom that is perfectly correlated with their reactive status. This makes *V* ^*′*^ a convenient toy setting in which the relationship between geometry, environment vectors, and functional labels is fully specified.

## B Mathematical foundations of the environment–vector space

### B.1 Definition as an embedded sub-manifold

#### Definition B.1 (Environment–vector manifold)

Let (*M* × *T, g*) be the 4-D Riemannian manifold of atomic configurations (Section 2.3). The map

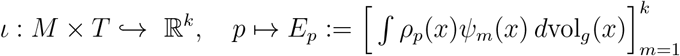

is a *C*^1^ immersion; its image *E* := *ι*(*M* × *T*), equipped with the pull-back metric *g*_*E*_ := *ι*_∗_(*g*), is called the *environment–vector manifold*.

### B.2 Metric and norm equivalence

#### Theorem B.2 (Ep metric equivalence)

*For all p, q* ∈ *E*,

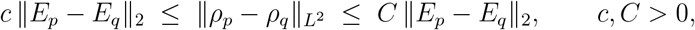

*and d*_*ρ*_(*p, q*) := arccos ⟨*ρ*_*p*_, *ρ*_*q*_⟩ */* (∥*ρ*_*p*_∥∥ *ρ*_*q*_∥)] *induces the same topology as* ∥*E*_*p*_ − *E*_*q*_ ∥_2_. *Consequently* (*E, d*) *is complete (Lemma 3*.*4–Theorem 3*.*5 in supplement file v2)*.

### B.3 Gaussian-type probability measure on *E*

#### Setup

Let *E* = *ι*(*M* × *T*) ⊂ ℝ^*k*^ be an *r*-dimensional embedded submanifold, equipped with the induced Riemannian volume measure vol_*E*_ (equivalently, vol_*E*_ = *ι*_∗_vol_*M×T*_ when *ι* is an embedding).

Fix *α >* 0 and define the (unnormalized) weight 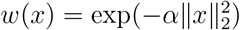 on *E*. Assume the normalizing constant

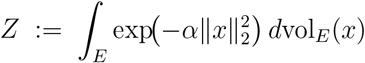

is finite. We then define a probability measure *µ* on *E* by

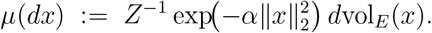

##### Lemma B.3 (Tail bound under polynomial volume growth)

*Assume that E has at most polynomial volume growth in the ambient radius: there exist constants C*_0_, *m >* 0 *such that for all R >* 0,

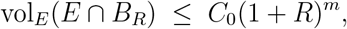

*where B*_*R*_ := {*x* ∈ ℝ^*k*^ : ∥*x*∥ _2_ ≤ *R* }. *Then there exist constants C*_1_, *c*_1_ *>* 0 *(depending on α and the growth constants) such that for all r >* 0,

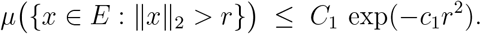

*Consequently, every polynomially bounded functional f* : *E* → ℝ *is µ-integrable*.

*Sketch*. Write

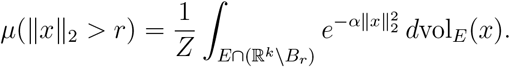

Decompose ℝ^*k*^ \ *B*_*r*_ into annuli *A*_*j*_ := *B* _*r*+*j*+1_ \ *B*_*r*+*j*_ and use 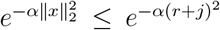 on *A*_*j*_, together with the growth assumption to bound vol_*E*_(*E* ∩ *A*_*j*_) ≤ *C*_0_(1 + *r* + *j* + 1)^*m*^. Summing the resulting series yields a Gaussian-type bound 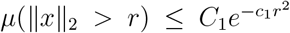. Polynomial integrability follows since 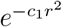 dominates any polynomial growth.

#### Dimension and normalization

Throughout, dim(*M* × *T*) = *r*, so *E* = *ι*(*M* × *T*) is an *r*-dimensional submanifold of ℝ^*k*^. The normalizing constant *Z* is given by the integral over *E* above and, in general, depends on the geometry of *E* (it does *not* reduce to the ℝ^*k*^ Gaussian normalizer unless *E* = ℝ^*k*^ with Lebesgue measure).

#### Motivation

Metric equivalence (Theorem B.2) ensures that switching from the Euclidean kernel used in Eq. (2) to a geodesic kernel (Appendix C) leaves the topology and completeness properties of (*E, d*) unchanged; hence statistical constructions on (*E, µ*) are robust to this choice of metric.

## C Time-Coherent Reactivity

### Time-resolved extension of the environment vector

In the main text we define a static environment vector Ep_*p*_ ∈ ℝ^*k*^ for each residue *p*, computed from a single protein structure (one PDB/mmCIF model). The components of Ep_*p*_ aggregate local geometric quantities (for example neighbour density, asymmetry, curvature), solvent exposure, and optional sequence-or metal-aware terms within a fixed cutoff around residue *p*.

In many realistic settings, however, a protein is not represented by a single structure but by a trajectory or ensemble

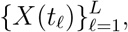

where each frame *X*(*t*_*ℓ*_) is a full set of atomic coordinates at a time-like index *t*_*ℓ*_ (for example, molecular dynamics, Monte Carlo sampling, or a collection of experimental conformers). In this case we simply apply the same environment-vector construction frame by frame.

#### Frame-wise environment vectors

For each residue *p* and each frame *t*_*ℓ*_ we define a neighbourhood *U*_*p*_(*t*_*ℓ*_) (for example, all atoms within a fixed cutoff radius) and compute the same local descriptors as in the static case: density-based terms, charge integrals, neighbour asymmetry, curvature density, solvent-accessible surface area, and any additional embedding components. This yields a time-dependent environment vector

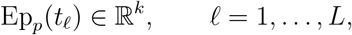

with the same dimensionality *k* and semantic interpretation as the original static Ep_*p*_.

Instead of a single point in environment space, each residue *p* now traces out a discrete trajectory

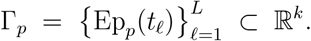

#### Static summaries of dynamic environments

To obtain a single descriptor per residue that incorporates all frames, we define simple time-coherent summaries of the Ep trajectory.

- **Time-averaged environment vector**

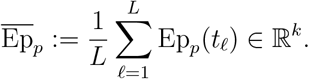

This has the same units and interpretation as the static environment vector and reduces exactly to Ep_*p*_ when *L* = 1.

- **Fluctuation magnitude in Ep space**

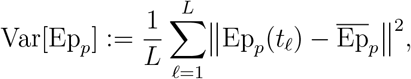

where ∥ · ∥ is the Euclidean norm in ℝ^*k*^. This scalar quantifies how strongly the local environment of residue *p* fluctuates over the trajectory.
- **Path length in Ep space**

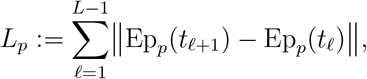

which is a discrete measure of how far the environment of *p* moves in Ep space across the trajectory.

These dynamic quantities can be combined with the static scoring terms defined elsewhere in the manuscript to obtain a time-coherent residue-level reactivity or functional score. For example, one may up-weight residues whose static environment is strongly reactive and whose environment vector exhibits large, structured fluctuations over the trajectory, as expected for hinge-like or allosteric sites. A full implementation of this dynamic scoring is left for future work, but the definitions above are compatible with the present toolkit: any MD trajectory or ensemble can be processed by computing Ep_*p*_(*t*_*ℓ*_) frame by frame and then applying user-chosen functionals of the discrete curve Γ_*p*_.

## Supporting information

math formalism

full raw result

code bundle

## Acknowledgements

This work was independently conceptualized and developed by the author (Haneul Park). The author utilized OpenAI ChatGPT models (GPT-4–class and successor models) to prototype analysis code, design statistical workflows (*k*-means clustering, distance-metric analysis, random-forest classification), polish L^A^TEX equations and notation, and refine the manuscript language. All AI-generated outputs were manually reviewed, validated, and adjusted by the author. All scientific reasoning, theoretical modeling, and experimental design originated from—and were confirmed by—the author.

## Notes

### Competing Interest Statement

The authors have declared no competing interest.

### Summary of Updates

Revision summary: We recomputed all benchmarks, including DB55 and all case studies, under a single strict protocol with solvent filtering and site grouped cross validation, and updated all reported AUC and AUPRC values. We corrected the ribosome hard negative panel relative to the final environment manifest and reran all tier 1 and tier 2 versus background and versus hard negative comparisons. We rewrote the abstract, results and discussion to reflect these updated numbers and to emphasise that the work is a representation level probe using a fixed deterministic environment vector.

## References

[1] John Jumper, Richard Evans, Alexander Pritzel, Tim Green, Michael Figurnov, Olaf Ronneberger, Kathryn Tunyasuvunakool, Russ Bates, Augustin Žídek, Anna Potapenko, Alex Bridgland, Clemens Meyer, Simon A. A. Kohl, Andrew J. Ballard, Andrew Cowie, Bernardino Romera-Paredes, Stanislav Nikolov, Rishub Jain, Jonas Adler, Trevor Back, Stig Petersen, David Reiman, Ellen Clancy, Michal Zielinski, Martin Steinegger, Michalina Pacholska, Tamas Berghammer, Sebastian Bodenstein, David Silver, Oriol Vinyals, Andrew W. Senior, Koray Kavukcuoglu, Pushmeet Kohli, and Demis Hassabis. Highly accurate protein structure prediction with AlphaFold. Nature, 596(7873):583–589, 2021.

[2] Minkyung Baek, Frank DiMaio, Ivan Anishchenko, Justas Dauparas, Sergey Ovchinnikov, Gyu Rie Lee, Jue Wang, Qian Cong, Lisa N. Kinch, R. Dustin Schaeffer, Claudia Millán, Hahnbeom Park, Phil D. Adams, Caleb R. Glassman, Andy DeGiovanni, Jimin Pereira, Joao P. G. L. M. Rodrigues, Marc van Dijk, Andreas C. Ebrecht, Derren J. Opperman, Toni Sagmeister, Christoph Buhlheller, Tea Pavkov-Keller, Manoj Kumar Rathinaswamy, Udit Dalwadi, Calvin K. Yip, John E. Burke, K. Christopher Garcia, Nick V. Grishin, Paul D. Adams, Randy J. Read, and David Baker. Accurate prediction of protein structures and interactions using a three-track neural network. Science, 373(6557):871–876, 2021.

[3] Predrag Radivojac, Wyatt T. Clark, Tal Ronnen Oron, and et al. A large-scale evaluation of computational protein function prediction. Nature Methods, 10(3):221–227, 2013.

[4] Vincent Le Guilloux, Peter Schmidtke, and Pierre Tuffery. fpocket: An open source platform for ligand pocket detection. BMC Bioinformatics, 10:168, 2009.

[5] John A. Capra, Roman A. Laskowski, Janet M. Thornton, Mona Singh, and Thomas A. Funkhouser. Predicting protein ligand binding sites by combining evolutionary sequence conservation and 3d structure. PLoS Computational Biology, 5(12):e1000585, 2009.

[6] Pablo Gainza, Freyr Sverrisson, Federico Monti, Emanuele Rodola, Davide Boscaini, Michael M. Bronstein, and Bruno E. Correia. Deciphering interaction fingerprints from protein surfaces using geometric deep learning. Nature Methods, 17(2):184–192, 2020.

[7] Yu-Feng Lin, Chih-Wei Cheng, Chien-Sheng Shih, Jeng-Kuan Hwang, Chen-Shan Yu, and Chung-Hsien Lu. Mib: Metal ion-binding site prediction and docking server. Journal of Chemical Information and Modeling, 56(12):2287–2291, 2016.

[8] Liang Wei and Russ B. Altman. Recognizing protein binding sites using statistical descriptions of their 3d environments. In Proceedings of the Pacific Symposium on Biocomputing, pages 497–508, 1999.

[9] Haim Ashkenazy, Elana Erez, Eric Martz, Tal Pupko, and Nir Ben-Tal. Consurf 2010: calculating evolutionary conservation in sequence and structure of proteins and nucleic acids. Nucleic Acids Research, 38(suppl 2):W529–W533, 2010.

[10] Justas Dauparas, Ivan Anishchenko, Nathaniel Bennett, Hua Bai, Robert J. Ragotte, Lukas F. Milles, Basile I. M. Wicky, Alexis Courbet, Robbert J. de Haas, Neville Bethel, Philip J. Y. Leung, Thomas F. Huddy, Samuel Pellock, Daniel Tischer, Franklin Chan, Brian Koepnick, Anh L. Nguyen, Aerin Kang, Bhanu Sankaran, Asim K. Bera, Neil P. King, and David Baker. Robust deep learning–based protein sequence design using ProteinMPNN. Science, 378(6615):49–56, 2022.

[11] Weitao Yang and Willem J. Mortier. The use of global and local molecular parameters for the analysis of the gas-phase basicity of amines. Journal of the American Chemical Society, 108(19):5708–5711, 1986.

[12] Arieh Warshel, Pratul K. Sharma, Mitsunori Kato, Yi-Qin Xiang, Hua-Lin Liu, and Max M. Kamerlin. Electrostatic basis for enzyme catalysis. Chemical Reviews, 106(8):3210–3235, 2006.

[13] Wai Shing Tang, Gabriel Monteiro da Silva, Henry Kirveslahti, Erin Skeens, Bibo Feng, Timothy Sudijono, Kevin K. Yang, Sayan Mukherjee, Brenda Rubenstein, and Lorin Crawford. A topological data analytic approach for discovering biophysical signatures in protein dynamics. PLoS Computational Biology, 18(5):e1010045, 2022.

[14] Zixuan Cang and Guo-Wei Wei. Analysis and prediction of protein folding energy changes upon mutation by element specific persistent homology. Bioinformatics, 33:3549–3557, 2017.

[15] Berend Diepeveen, Shreyas Banerjee, Nikola B. Kovachki, Arnulf Jentzen, Luca Venturi, Kamyar Azizzadenesheli, Dusan Pavlovic, and Ozan Öktem. Riemannian geometry for efficient analysis of protein dynamics data. Proceedings of the National Academy of Sciences of the United States of America, 121(33):e2318951121, 2024.

[16] Huanwang Zheng, D. R. Cooper, P. J. Porebski, I. G. Shabalin, K. B. Handing, and W. Minor. Checkmymetal: a macromolecular metal-binding validation tool. Acta Crystallographica Section D: Structural Biology, 73(3):223–233, 2017.

[17] Paul D. Adams, Pavel V. Afonine, Gábor Bunkóczi, Vincent B. Chen, Ian W. Davis, Nathaniel Echols, Jeffrey J. Headd, Li-Wei Hung, Gary J. Kapral, Ralf W. Grosse-Kunstleve, Airlie J. McCoy, Nigel W. Moriarty, Robert Oeffner, Randy J. Read, Jane S. Richardson, David C. Richardson, Thomas C. Terwilliger, and Peter H. Zwart. Phenix: a comprehensive python-based system for macromolecular structure solution. Acta Crystallographica Section D: Biological Crystallography, 66(2):213–221, 2010.

[18] Gerhard Langer, Stewart X. Cohen, Victor S. Lamzin, and Anastassis Perrakis. Automated macromolecular model building for x-ray crystallography using ARP/wARP version 7. Nature Protocols, 3(7):1171–1179, 2008.

[19] Pierre Hohenberg and Walter Kohn. Inhomogeneous electron gas. Physical Review, 136(3B):B864–B871, 1964.

[20] Erich Runge and E. K. U. Gross. Density-functional theory for time-dependent systems. Physical Review Letters, 52(12):997–1000, 1984.

[21] Kelin Xia and Guo-Wei Wei. Persistent homology analysis of protein structure, flexibility, and folding. Int. J. Numer. Meth. Biomed. Engng., 30(8):814–844, 2014.

