## Supplementary material for "Localized Reactivity on Proteins as Riemannian Manifolds: A Quantum-Inspired Geometric Model for Deterministic, Metal-Aware Reactive-Site Prediction": math formalism

### A Geometric and Statistical Framework for Residue-Level Environment Vectors (Ep)

Haneul Park
Department of Biological Sciences and Department of Mathematical Sciences
Seoul National University, Seoul, South Korea


This manuscript formalizes the method originally proposed in a prior preprint [R5], available at https://doi.org/10.1101/2025.04.29.651260.

**Abstract**

Residue-level structural environments are central to protein function, yet the definition of such environments often relies on heuristic features, contact graphs, or black-box embeddings. In this work, we propose a geometric formulation of the local environment vector (Ep) as a well-defined point in a structured topological space. We define a manifold-like framework on the Ep space, introduce a natural distance function between environments, and construct a measure-theoretic foundation for statistical reasoning. This work lays the mathematical groundwork for downstream applications in interaction prediction, embedding-based learning, and electronic density modeling. This work generalizes the preliminary environment vector framework proposed in our previous study [R5], extending it into a rigorously defined metric-topological formalism on Riemannian manifolds.

#### 1 Introduction

The local structural context of a residue in a protein is critical to its biochemical role. Numerous descriptors—such as solvent accessibility, contact maps, and graph-based embeddings—have been used to capture this information. However, these approaches often lack a precise mathematical structure, making them hard to interpret, compare, or extend.

To address this gap, we propose a formal definition of the environment vector (Ep) for each residue as an element of a structured geometric space. By constructing a topological and probabilistic framework around Ep, we aim to provide a mathematically grounded foundation that supports interpretation, statistical analysis, and integration with density-based physical models [R1]

This paper formalizes the structure of the Ep space, introduces a Riemannian metric for comparing environments, and constructs the basis for statistical reasoning over such spaces. throughout this paper, We treat $M \times T$ as a 4-dimensional differentiable manifold, where $T$ represents a time-like coordinate that may, in future formulations, be interpreted within a Minkowski-type or generalized space–time framework. This framework serves as the theoretical backbone for subsequent applications including AI-compatible embeddings and residue-level reactivity predictions.

#### 2 Definition of the Environment Vector (Ep)

Whereas [R5] empirically utilized the environment vector Ep as a descriptor for reactive site prediction, we now lift Ep into a formal object with a well-defined domain, codomain, and metric structure. This section provides the necessary mathematical formulation.

**Definition 2.1** (Ep, Grid-Based)**.** *Let $R_i$ be a residue and $\mathcal{N}_i = \{A_j \mid \|A_j - R_i\| < r\}$ be its spatial neighborhood within cutoff radius $r$. We define the environment vector* $\mathrm{Ep}_i \in \mathbb{R}^k$ *by aggregating local atomic features as:*

$$\mathrm{Ep}_i^{(m)} := \frac{1}{|\mathcal{N}_i|} \sum_{A_j \in \mathcal{N}_i} \phi_m(A_j) \quad \textit{for } m = 1, \ldots, k$$

*Here $\phi_m : \mathbb{M} \times T \to \mathbb{R}$ is a feature function, such as:*

- *Distance from $R_i$*
- *Directionality cosine*
- *Local partial charge*
- *Hydrophobicity*
- *Distance to nearest metal ion*

*The resulting vector* $\mathrm{Ep}_i$ *encodes a statistical average over the residue's atomic environment and forms a structured point in Ep-space $\mathcal{E} \subset \mathbb{R}^k$.*

**Continuity convention.** Instead of a hard cut–off, we weight neighbours with a smooth kernel $w(r) = \exp\big(-r^2/\sigma^2\big)$. Consequently the map $p \mapsto E_p$ is $C^\infty$ on $E$.

**Remark on Generalization.** While Definition 1 constructs the environment vector Ep as an average over discrete atomic features in a fixed neighborhood, Definition 2 provides a continuous generalization using a smoothed electron density function $\rho_i(x)$. This shift from a discrete feature sum to a continuous projection integral allows the Ep representation to be embedded in a functional space, opening the door to differential geometric and density-based analysis. In particular, Definition 2 recovers the structure of Definition 1 in the limiting case where the density $\rho_i(x)$ is composed of narrow atomic kernels centered at the same positions used in Definition 1.

**Definition 2.2** (Feature Function Spaces)**.** *Let $\phi_m : \mathbb{M} \times T \to \mathbb{R}$ be a local atomic feature function used to construct Ep, and let $\psi_m : \mathbb{M} \times T \to \mathbb{R}$ be a projection basis function.*

*We assume the following:*

- $\phi_m \in C^\infty(\mathbb{M} \times T)$ *— infinitely differentiable*
- $\psi_m \in L^2(\mathbb{M} \times T)$ *— square-integrable*
- $\{\psi_m\}_{m=1}^k$ *is an orthonormal set in $L^2$*

**Definition 2.3** (Ep, Density-Based Manifold Interpretation)**.** *Let each residue $R_i$ be associated with a local electron density function $\rho_i : M \times T \to \mathbb{R}_{\geq 0}$, obtained by kernel-smoothing the atomic positions within its neighborhood $\mathcal{N}_i$. We interpret $M \times T$ as a 4-dimensional smooth Riemannian manifold with metric $g$, and denote the induced volume form by* $\mathrm{dvol}_g$.

*Then the environment vector $Ep_i \in \mathbb{R}^k$ is defined via a finite set of projection functionals:*

$$Ep_i^{(m)} := \int_{M \times T} \rho_i(x) \cdot \psi_m(x) \, \mathrm{dvol}_g(x), \quad \textit{for } m = 1, \ldots, k$$

*where $\{\psi_m\}_{m=1}^k \subset C^\infty(M \times T)$ is a fixed set of smooth scalar basis functions (e.g., radial basis functions, Gaussian shells, or low-degree polynomials) defined on the manifold.*

*The space of all such $Ep_i$ forms a submanifold $\mathcal{E} \subset \mathbb{R}^k$, which can be equipped with the standard inner product $\langle Ep_p, Ep_q \rangle := \sum_m Ep_p^{(m)} Ep_q^{(m)}$, and may inherit a Riemannian structure from the ambient Euclidean space.*

> **Note (Generic-Position Assumption).** Throughout this work we assume that atomic configurations are in a *generic* state: thermal/solvent fluctuations and experimental noise (typically $\gtrsim 0.1$ Å) are large enough to destroy any perfectly symmetric arrangement that would lower the Jacobian rank or merge distinct residues into an identical environment vector. Formally, for the map
>
> $$\iota\colon (M \times T) \longrightarrow \mathbb{R}^k, \qquad p \longmapsto E_p,$$
>
> the set of rank-deficient or self-intersecting configurations has Lebesgue measure 0 in configuration space. Hence, with probability one, $\iota$ is an **immersion** of constant rank $r$ and is **globally injective**, so its image $E = \iota(M \times T)$ is an $r$-dimensional embedded **sub-manifold** of $\mathbb{R}^k$. All differential-geometric results in Sections 3–5 are therefore valid on $E$ without further qualification.

#### 2.1 Time-resolved environment vectors on $M \times T$

We first recall the static construction. The protein is modeled as a four-dimensional space–time manifold $M \times T$ equipped with a Riemannian metric $g$ and volume form $\mathrm{dvol}_g$. For each residue $R_i$ we introduce a nonnegative density

$$\rho_i : M \times T \longrightarrow \mathbb{R}_{\geq 0}$$

and a fixed family of basis functions $\{\psi_m\}_{m=1}^k \subset C^\infty(M \times T)$. The static environment vector $\mathrm{Ep}_i \in E \subset \mathbb{R}^k$ is defined component-wise by

$$\mathrm{Ep}_i^{(m)} = \int_{M \times T} \rho_i(x,t)\, \psi_m(x,t)\, \mathrm{dvol}_g(x,t), \qquad m = 1, \dots, k,$$

where $E$ denotes the image of $M \times T$ in $\mathbb{R}^k$ under the Ep map.

In many applications, however, the protein is not observed as a single configuration but as a time-resolved trajectory (for example, a molecular dynamics simulation). In this setting we extend the Ep formalism to handle trajectories without changing the manifold structure or the basis functions.

**Residue trajectories in space–time.** We regard a protein trajectory as a curve in $M \times T$. For each residue $R_i$ we consider its worldline

$$\gamma_i : T \longrightarrow M \times T, \qquad t \longmapsto (x_i(t), t),$$

where $x_i(t) \in M$ denotes the spatial position of residue $i$ at time $t$. In practice, simulations or ensembles are given as a finite sequence of frames

$$\{t_\ell\}_{\ell=1}^L \subset T,$$

so that only the discrete samples $\gamma_i(t_\ell)$ are observed.

The density $\rho_i(x,t)$ is defined on all of $M \times T$. For each fixed time $t \in T$ we may restrict $\rho_i$ to the time-slice $\{t\} \times M$ and view

$$\rho_i(\cdot, t) : M \longrightarrow \mathbb{R}_{\geq 0}, \qquad x \longmapsto \rho_i(x,t)$$

as a spatial density at time $t$. The time-resolved construction below can be regarded as a slice-wise analogue of the static space–time integral above.

**Time-dependent environment vectors.** For each residue $R_i$ and each time $t \in T$ we define a time-resolved environment vector

$$\mathrm{Ep}_i(t) \;=\; \big(\mathrm{Ep}_i^{(1)}(t), \ldots, \mathrm{Ep}_i^{(k)}(t)\big)^\top \in E \subset \mathbb{R}^k$$

by projecting $\rho_i(\cdot, t)$ onto the same family of basis functions $\{\psi_m\}_{m=1}^k$:

$$\mathrm{Ep}_i^{(m)}(t) \;:=\; \int_M \rho_i(x,t)\, \psi_m(x,t)\, \mathrm{dvol}_{g_t}(x), \qquad m = 1, \ldots, k. \tag{1}$$

Here $g_t$ denotes the restriction of the space–time metric $g$ to the spatial slice $\{t\} \times M$, and $\mathrm{dvol}_{g_t}$ is the induced volume form on $M$ at time $t$. In discrete data, this integral is implemented as a quadrature or finite sum over grid points or atoms in a neighbourhood of $x_i(t_\ell)$.

For any fixed time $t$, the map

$$\mathrm{Ep}_t : M \longrightarrow E, \qquad p \longmapsto \mathrm{Ep}(p,t),$$

satisfies the same generic-position and regularity assumptions as in the static construction. In particular, each time-slice image

$$E_t := \mathrm{Ep}_t(M)$$

is an embedded submanifold of $E$. The full trajectory of environments for residue $R_i$ is therefore a curve

$$\Gamma_i : T \longrightarrow E, \qquad t \longmapsto \mathrm{Ep}_i(t),$$

obtained by composing the worldline $\gamma_i$ with the Ep map.

**Discrete trajectories and frame-wise Ep vectors.** For a finite trajectory with frames $\{t_\ell\}_{\ell=1}^L$, we observe the discrete sequence

$$\Gamma_i^{\mathrm{disc}} \;=\; \big\{\mathrm{Ep}_i(t_\ell)\big\}_{\ell=1}^L \;\subset\; E.$$

This is the object used in numerical experiments: the Ep map is applied frame by frame, and no modification of the underlying definition is required.

**Static summaries of dynamic environments.** In many downstream tasks, such as residue-level functional prediction, one ultimately requires a single feature vector per residue, even when a full trajectory $\Gamma_i$ is available. Within this framework, such a reduction corresponds to applying deterministic functionals to the discrete curve $\Gamma_i^{\mathrm{disc}}$.

Given the frame-wise environment vectors $\{\mathrm{Ep}_i(t_\ell)\}_{\ell=1}^L$, we define:

- **Time-averaged environment vector**

$$\overline{\mathrm{Ep}_i} \;:=\; \frac{1}{L} \sum_{\ell=1}^L \mathrm{Ep}_i(t_\ell) \;\in\; \mathbb{R}^k.$$

  Each $\mathrm{Ep}_i(t_\ell)$ lies on the embedded submanifold $E \subset \mathbb{R}^k$, but their Euclidean average $\overline{\mathrm{Ep}_i}$ is, in general, only guaranteed to lie in the ambient space $\mathbb{R}^k$.

- **Fluctuation magnitude in Ep space**

$$\mathrm{Var}[\mathrm{Ep}_i] \;:=\; \frac{1}{L}\sum_{\ell=1}^{L} \big\|\mathrm{Ep}_i(t_\ell) - \overline{\mathrm{Ep}}_i\big\|^2,$$

  where $\|\cdot\|$ is the Euclidean norm in $\mathbb{R}^k$ (equivalently, the norm induced on $E$ by the ambient metric). This scalar measures how strongly the local environment of residue $i$ fluctuates in Ep space over the trajectory.

- **Path length in Ep space**

$$L_i \;:=\; \sum_{\ell=1}^{L-1} \big\|\mathrm{Ep}_i(t_{\ell+1}) - \mathrm{Ep}_i(t_\ell)\big\|,$$

  which is a discrete analogue of the curve length of $\Gamma_i$ in $E$ and quantifies how far the environment of residue $i$ travels in Ep space.

When the ordering of frames reflects a genuine dynamical evolution (for example, molecular dynamics time), one can also define Ep-space auto-correlations at selected lags. For a chosen lag index $h \in \{1, \dots, L-1\}$, with corresponding time increment $\Delta t_h \approx t_{\ell+h} - t_\ell$, set

$$\mathrm{ACF}_i(\Delta t_h) \;:=\; \frac{1}{L-h}\sum_{\ell=1}^{L-h} \Big\langle \mathrm{Ep}_i(t_\ell) - \overline{\mathrm{Ep}}_i,\; \mathrm{Ep}_i(t_{\ell+h}) - \overline{\mathrm{Ep}}_i \Big\rangle,$$

where $\langle \cdot, \cdot \rangle$ is the Euclidean inner product on $\mathbb{R}^k$.

**Relation to the static Ep formalism.** The static, structure-only environment vector is recovered as the special case where the trajectory consists of a single frame ($L = 1$) or where only the time-averaged vector $\overline{\mathrm{Ep}}_i$ is used. The time-resolved construction above therefore extends, rather than replaces, the original Ep formalism: the manifold $E \subset \mathbb{R}^k$, the basis functions $\{\psi_m\}$, and the density-based definition of $\mathrm{Ep}_i$ remain unchanged, while trajectories in $M \times T$ are mapped to curves $\Gamma_i$ in Ep space and summarized by functionals such as $\overline{\mathrm{Ep}}_i$, $\mathrm{Var}[\mathrm{Ep}_i]$, and $L_i$.

### 3 Geometric Structure on the Ep Space

we now work with Definition 2.3 exclusively.

#### 3.1 Smoothness of Ep with Respect to Atomic Coordinates

We now turn to the regularity properties of the Ep vector. Since each component of Ep is constructed from smooth feature functions $\phi_m$ applied to atomic positions, it is natural to ask whether the Ep vector itself varies smoothly as atoms move.

The following lemma formalizes this intuition:

**Lemma 3.1.** *If all feature functions $\phi_m \in C^k(\mathbb{M} \times T)$, and the atomic positions $\mathbf{r}_j$ vary smoothly, then the map $\mathbf{r} \mapsto E_p(\mathbf{r})$ is $C^k$ in the atomic coordinates.*

*Sketch.* Since Ep is defined as a weighted average of $\phi_m(\mathbf{r}_j)$ over atoms $A_j \in \mathcal{N}_p$, and each $\phi_m$ is $C^k$, the sum of smooth functions composed with smooth atomic coordinates is again $C^k$. Therefore, $E_p$ varies smoothly under perturbation of atomic positions. □

The Ep space $\mathcal{E} \subset \mathbb{R}^k$ is endowed with a natural topology inherited from the Euclidean metric. We further define a geometric structure by introducing a distance function between two environment vectors $\mathrm{Ep}_p, \mathrm{Ep}_q \in \mathcal{E}$.

##### 3.2 Distance Function

We define the distance between two environment vectors in $\mathcal{E} \subset R^k$ by the *angular metric*:

$$d\rho(p,q) := \arccos\left( \frac{O(p,q)}{\sqrt{O(p,p)\,O(q,q)}} \right),$$

where the overlap integral is

$$O(p,q) := \int_{M\times T} \rho_p(x)\,\rho_q(x)\,dvol_g(x), \quad \rho_p \in C^\infty(M\times T),\ \rho_q \in C^\infty(M\times T),$$

and $O(p,p) > 0$ for all $p$ by construction (or ensured via $\varepsilon$–regularisation).

**Remark.** Since

$$0 \ \leq\ O(p,q)\ \leq\ \sqrt{O(p,p)\,O(q,q)},$$

the argument of $\arccos(\cdot)$ lies in $[0,1]$, and hence $d\rho(p,q) \in [0, \frac{\pi}{2}]$. The function $\arccos\colon [0,1] \to [0, \frac{\pi}{2}]$ is continuous and strictly decreasing, which together with the properties of $O$ ensures that $d\rho$ satisfies

(i) *non-negativity*: $d\rho(p,q) \geq 0$,

(ii) *identity of indiscernibles*: $d\rho(p,q) = 0 \iff p = q$,

(iii) *symmetry*: $d\rho(p,q) = d\rho(q,p)$,

(iv) *triangle inequality*: $d\rho(p,r) \leq d\rho(p,q) + d\rho(q,r)$.

**Cut-off Distance and Interaction Threshold.** To formalize locality and suppress non-interacting pairs, we define a threshold $\tau > 0$:

$$d'(p,q) := \begin{cases} d\rho(p,q), & d\rho(p,q) \leq \tau, \\ \infty, & \text{otherwise,} \end{cases}$$

which turns $\mathcal{E}$ into a metric space with interaction cut-off.

##### 3.3 Metric and Riemannian Manifold Interpretation

Under the assumption of smooth variation in the Ep vectors across residues and proteins, we may treat $\mathcal{E}$ as a Riemannian manifold. A local chart $\phi : U \to \mathbb{R}^k$ maps an open neighborhood of $\mathcal{E}$ to Euclidean space, and a Riemannian metric $g$ is defined by the inner product on $\mathbb{R}^k$:

$$g_{\mathrm{Ep}_p}(v,w) = \langle v, w \rangle_{\mathbb{R}^k}$$

##### 3.4 Topological Equivalence of Metrics

We have introduced both the Euclidean distance in feature space and the density-overlap based metric. A natural question arises: do these two metrics induce the same topology on $\mathcal{E}$? This matters because downstream statistical or geometric analysis may depend on which metric is used.

The proposition below addresses this question:

**Proposition 3.2.** *Let* $d_{\mathrm{Euc}}(p,q) := \|E_p - E_q\|$ *and*

$$d_\rho(p,q) \;=\; \arccos\!\left(\frac{O(p,q)}{\sqrt{O(p,p)\,O(q,q)}}\right)$$

*with* $O(p,q) := \int \rho_p(x)\rho_q(x)\,dx$, *where* $\rho_p(x)$ *is constructed as a continuous projection from* $E_p$, *i.e.,* $\rho_p(x) = \sum_m E_p^{(m)}\psi_m(x)$ *for fixed basis* $\psi_m \in L^2$.

*Then* $d_{\mathrm{Euc}}$ *and* $d_\rho$ *induce the same topology on* $\mathcal{E}$.

But, before proof of proposition, we need to prove the following lemma and theorem.

**Lemma 3.3** (Injectivity of the density map)**.** *Let*

$$n : \mathbb{R}^k \to L^2(\Omega), \quad n(E_p) = \rho_p(x) = \sum_{m=1}^{n} E_p^{(m)}\,\psi_m(x).$$

*If* $\{\psi_m\}_{m=1}^n$ *is linearly independent, then* $n$ *is injective.*

*Proof.* Suppose $n(E_p) = n(E_q)$. Then

$$0 = \rho_p - \rho_q = \sum_{m=1}^{n} (E_p^{(m)} - E_q^{(m)})\,\psi_m \quad \text{in } L^2(\Omega).$$

Linear independence forces all coefficients to vanish, hence $E_p = E_q$. □

**Lemma 3.4** (Positive-definiteness of $d_{L^2}$)**.** *Define*

$$d_{L^2}(p,q) = \|\rho_p - \rho_q\|_{L^2}.$$

*Then* $d_{L^2}(p,q) = 0 \iff p = q$, *and* $d_{L^2}$ *is a metric on* $E$.

*Proof.* If $d_{L^2}(p,q) = 0$, then $\rho_p = \rho_q$ in $L^2$ and by Lemma 3.3 $p = q$. Symmetry and the triangle inequality follow from the properties of the $L^2$-norm. □

**Theorem 3.5** (Equivalence of Euclidean and $d_{L^2}$ topologies)**.** *There exist constants* $c, C > 0$ *such that for all* $p, q \in E$,

$$c\,\|E_p - E_q\|_{\mathrm{Euc}} \;\le\; \|\rho_p - \rho_q\|_{L^2} \;\le\; C\,\|E_p - E_q\|_{\mathrm{Euc}}.$$

*Hence* $d_{\mathrm{Euc}}$ *and* $d_{L^2}$ *induce the same topology on* $E$.

*Proof.* Since $M : \mathbb{R}^k \to \mathrm{span}\{\psi_m\}$ is a linear isomorphism onto its image in finite dimensions, all norms on that subspace are equivalent. □

**Lemma 3.6** (Overlap–$d_{L^2}$ equivalence)**.** *For all $p, q \in E$ with $\rho_p, \rho_q \neq 0$, define the normalized densities $\tilde{\rho}_p := \rho_p/\|\rho_p\|_{L^2}$ and $\tilde{\rho}_q := \rho_q/\|\rho_q\|_{L^2}$. Let*

$$d_\rho(p,q) := \arccos\left( \frac{\langle \rho_p, \rho_q \rangle_{L^2}}{\|\rho_p\|_{L^2}\,\|\rho_q\|_{L^2}} \right) = \arccos\big( \langle \tilde{\rho}_p, \tilde{\rho}_q \rangle_{L^2} \big),$$

*and let $d_{L^2}(p,q) := \|\rho_p - \rho_q\|_{L^2}$. Then the following bounds hold:*

$$\frac{2}{\pi}\, d_\rho(p,q) \ \leq\ \|\tilde{\rho}_p - \tilde{\rho}_q\|_{L^2} \ \leq\ d_\rho(p,q).$$

*In particular, $d_\rho$ and the chordal distance $\|\tilde{\rho}_p - \tilde{\rho}_q\|_{L^2}$ induce the same topology on the set $\{\rho \in L^2 : \rho \neq 0\}$, hence on $E$ under the identification $p \mapsto \tilde{\rho}_p$.*

*Proof.* Since $\|\tilde{\rho}_p\|_{L^2} = \|\tilde{\rho}_q\|_{L^2} = 1$, we have

$$\|\tilde{\rho}_p - \tilde{\rho}_q\|_{L^2}^2 = 2 - 2\langle \tilde{\rho}_p, \tilde{\rho}_q \rangle_{L^2} = 2 - 2\cos\big(d_\rho(p,q)\big).$$

Thus $\|\tilde{\rho}_p - \tilde{\rho}_q\|_{L^2} = 2\sin\big(d_\rho(p,q)/2\big)$. For $\theta \in [0,\pi]$, the inequalities $\frac{2}{\pi}\theta \leq 2\sin(\theta/2) \leq \theta$ hold, yielding

$$\frac{2}{\pi}\, d_\rho(p,q) \leq \|\tilde{\rho}_p - \tilde{\rho}_q\|_{L^2} \leq d_\rho(p,q).$$

Since $\theta \mapsto 2\sin(\theta/2)$ is continuous and strictly increasing on $[0,\pi]$, the two distances induce the same topology. □

*Proof of Proposition 3.2.* By Theorem 3.5, $d_{\mathrm{Euc}}$ and $d_{L^2}$ are equivalent. By Lemma 3.6, $d_{L^2}$ and $d_\rho$ (overlap) are equivalent. Transitivity of metric equivalence yields $d_{\mathrm{Euc}} \sim d_\rho$, as claimed. □

### 4 Statistical Interpretation on the Ep Space

We now equip the Ep-space $E \subset \mathbb{R}^k$ with a probabilistic structure. Throughout this section, all probability mass is supported on the embedded submanifold

$$E = \iota(M \times T) \subset \mathbb{R}^k,$$

and all densities are understood with respect to the induced volume measure on $E$.

#### 4.1 Probability Measure on $E$ via Push-Forward

**Geometric setup.** Let $(M \times T, g)$ be an $r$-dimensional orientable Riemannian manifold with volume form $\mathrm{vol}_{M\times T}$, and let

$$\iota : M \times T \to \mathbb{R}^k$$

be a $C^1$ immersion. We write $E := \iota(M \times T)$ for the image. When $\iota$ is an embedding (or a $C^1$-diffeomorphism onto its image), the induced volume measure on $E$ is well-defined by push-forward:

$$\mathrm{vol}_E := \iota_*\big(\mathrm{vol}_{M\times T}\big).$$

**A Gaussian-type weight on $M \times T$.** Fix $\alpha > 0$ and define the (unnormalized) weight

$$w(p) := \exp\bigl(-\alpha\|\iota(p)\|^2\bigr), \qquad p \in M \times T.$$

Assume the normalizing constant

$$Z := \int_{M\times T} w(p)\, d\mathrm{vol}_{M\times T}(p)$$

is finite. (Sufficient conditions are given in Remark **??**.) Define a probability measure $\nu$ on $M \times T$ by

$$d\nu(p) := \frac{1}{Z}\, w(p)\, d\mathrm{vol}_{M\times T}(p).$$

**Definition 4.1** (Probability measure on Ep space)**.** *Define the probability measure $\mu$ on $E$ as the push-forward of $\nu$ under $\iota$:*

$$\mu := \iota_*\nu, \qquad \mu(B) := \nu\bigl(\iota^{-1}(B)\bigr), \quad B \subset E \textit{ Borel.}$$

**Proposition 4.2** (Normalization)**.** *The measure $\mu$ is a well-defined probability measure on $E$, i.e. $\mu(E) = 1$.*

*Proof.* By Definition 4.1, $\mu(E) = \nu(\iota^{-1}(E)) = \nu(M \times T) = 1$. □

**Radon–Nikodym derivative on $E$.** If $\iota$ is a $C^1$ embedding (or diffeomorphism onto its image) so that $\mathrm{vol}_E$ is well-defined, then $\mu$ is absolutely continuous with respect to $\mathrm{vol}_E$ and admits the density

$$\frac{d\mu}{d\mathrm{vol}_E}(x) = \frac{1}{Z}\exp\bigl(-\alpha\|x\|^2\bigr), \qquad x \in E.$$

Equivalently, for any bounded measurable $f : E \to \mathbb{R}$,

$$\int_E f(x)\, d\mu(x) = \frac{1}{Z}\int_E f(x)\exp\bigl(-\alpha\|x\|^2\bigr)\, d\mathrm{vol}_E(x).$$

**When is $Z < \infty$?** If $M \times T$ is compact, then $Z < \infty$ holds trivially. More generally, $Z < \infty$ holds whenever the induced volume growth of $E$ is at most polynomial in the ambient radius, i.e. there exist $C, m > 0$ such that $\mathrm{vol}_E(E \cap B_R) \le C(1+R)^m$ for all $R > 0$, since the Gaussian factor $\exp(-\alpha\|x\|^2)$ dominates any polynomial growth.

#### 4.2 Expectation and Variance

Given a measurable functional $f : E \to \mathbb{R}$, we define its expectation under $\mu$ as

$$\mathbb{E}_\mu[f] := \int_E f(x)\, d\mu(x),$$

whenever the integral exists. The variance is defined by

$$\mathrm{Var}_\mu(f) := \mathbb{E}_\mu\Bigl[(f - \mathbb{E}_\mu[f])^2\Bigr],$$

whenever the right-hand side is finite.

**Lemma 4.3** (Integrability of polynomially bounded functionals)**.** *Assume $Z < \infty$ and that $E$ has at most polynomial volume growth as in Remark* **??**. *Let $f : E \to \mathbb{R}$ satisfy*

$$|f(x)| \le C_f\big(1 + \|x\|\big)^m$$

*for some constants $C_f, m > 0$. Then $f \in L^1(E,\mu)$ and $\mathbb{E}_\mu[f]$ is finite.*

*Sketch.* Using the density of $\mu$ with respect to $\mathrm{vol}_E$,

$$\int_E |f(x)|\, d\mu(x) = \frac{1}{Z}\int_E |f(x)| e^{-\alpha\|x\|^2}\, d\mathrm{vol}_E(x).$$

By the assumed polynomial bound on $f$ and polynomial growth of $\mathrm{vol}_E$ on $E \cap B_R$, the integrand is dominated by a polynomial factor times $e^{-\alpha\|x\|^2}$, which is integrable. □

### 5 Toy Example: Alanine Dipeptide

To illustrate the construction and smooth variation of the environment vector $E_p$, we consider a simplified two-residue system resembling alanine dipeptide.

Let residue $R_1$ be centered at the origin with neighboring atoms $\{A_j\}$ at coordinates $\{\mathbf{r}_j \in \mathbb{M} \times T\ \}$. We define feature functions $\phi_1(x) = \|x\|$, $\phi_2(x) = \cos(\theta_x)$, where $\theta_x$ is the angle between $x$ and a fixed direction.

Then the Ep vector is:

$$E_p = \left[\frac{1}{|\mathcal{N}_p|}\sum_{x\in\mathcal{N}_p}\|x\|, \quad \frac{1}{|\mathcal{N}_p|}\sum_{x\in\mathcal{N}_p}\cos(\theta_x)\right]$$

A small perturbation of the atomic coordinates (e.g., thermal noise) results in continuous variation in $E_p$, validating the smoothness assumption.

Furthermore, we can evaluate the Euclidean and overlap-based distances between perturbed and original $E_p$ vectors to compare metric sensitivity.

### 6 Discussion

#### 6.1 Comparison with Traditional Descriptors

The Ep framework contrasts with many existing residue-level structural descriptors. Common features such as solvent-accessible surface area (SASA), contact number, and graph-based embeddings often lack a cohesive geometric interpretation or operate in a black-box fashion.

In contrast, our definition of Ep offers:

- A mathematically structured representation space
- Geometrically interpretable distance and metric properties
- Probabilistic integration for statistical modeling

Moreover, Ep can be extended modularly, incorporating additional physicochemical quantities or data-driven feature transformations without breaking the underlying structure.

##### 6.2 Applications and Extensions

The structured nature of the Ep space enables various downstream applications:

- **Reactivity prediction:** Using Ep vectors to predict active site residues or allosteric regions.
- **Residue clustering:** Applying intrinsic distance metrics for unsupervised learning or embedding.
- **AI feature engineering:** Feeding Ep into graph neural networks (GNNs) or contrastive learning frameworks.
- **Density modeling:** Interpreting electron density overlaps and constructing energy-related functionals [R3].

##### 6.3 Outlook

We envision two distinct directions of extending the Ep formalism: a probabilistic manifold perspective (Section 6.3), and a topological interaction geometry (Section 6.4), each of which opens new pathways for modeling, analysis, and learning.

While this paper avoids invoking the full machinery of probabilistic manifolds, such generalizations are possible. In future work, we plan to construct a probabilistic manifold $\mathcal{M}$, where each point corresponds to a trajectory-level quantum state [R2], extending the current formalism toward more expressive stochastic and quantum modeling.

##### 6.4 Outlook2: Toward Topological Interaction Geometry

**Regular–value assumption.** Fix $\varepsilon > 0$ such that it is a *regular value* of the smooth map

$$\varphi_{pq}(x) \;:=\; |\psi_p(x,t)|^2\,|\psi_q(x,t)|^2, \qquad x \in M \times T,$$

i.e. $\nabla\varphi_{pq}(x) \neq 0$ whenever $\varphi_{pq}(x) = \varepsilon$. Then the level set

$$\partial M^{\varepsilon}_{pq} := \varphi_{pq}^{-1}(\varepsilon)$$

is a closed, embedded hypersurface of codimension 1, and $M^{\varepsilon}_{pq} := \varphi_{pq}^{-1}\big([\varepsilon,\infty)\big)$ is a smooth domain with boundary $\partial M^{\varepsilon}_{pq}$. The Ep framework provides a structured space for residue environments. In future work, we aim to extend this to topological analysis of electron density overlaps.

Given two residues $p, q$, their interaction region is:

$$\mathcal{M}_{pq} := \{x \in \mathbb{M} \times T \mid \rho_p(x) \cdot \rho_q(x) \geq \varepsilon\}$$

This submanifold allows us to compute geometric invariants such as area, Euler characteristic $\chi(\mathcal{M}_{pq})$, or homology classes.

Such topological quantities offer a geometry-informed alternative to scalar scoring, and may eventually inform reactivity prediction or machine learning pipelines. Understanding how Sp and Ep encode or correlate with such invariants is an important direction for future work.

### 7 Conclusion

We have presented a geometric and statistical framework for representing residue-level environments in proteins via the environment vector Ep. The Ep space is structured as a topological and metric space, with an optional Riemannian interpretation under smoothness assumptions.

We defined distance functions, introduced a statistical measure structure, and established functionals such as expectation and variance. These tools allow for mathematical analysis, probabilistic reasoning, and integration with physical quantities such as electron density.

The Ep framework provides a mathematically grounded alternative to heuristic or black-box descriptors and lays a foundation for downstream applications in learning, prediction, and quantum modeling.


#### Acknowledgements

Portions of this manuscript were refined and organized with the assistance of a large language model (GPT-4), particularly in LaTeX structuring and technical phrasing. All mathematical definitions, theoretical constructions, and conceptual directions were independently developed by the author.


#### References


[R1] I. Bengtsson and K. Życzkowski, *Geometry of Quantum States*, Cambridge University Press, 2006.

[R2] M. Nielsen and I. Chuang, *Quantum Computation and Quantum Information*, Cambridge University Press, 2000.

[R3] D. J. Wales, *Energy Landscapes*, Cambridge University Press, 2003.

[R4] L. Wasserman, *All of Statistics*, Springer, 2004.

[R5] H. Park, *Localized Reactivity on Proteins as Riemannian Manifolds: A Geometric and Quantum-Informed Basis for Deterministic, Metal-Aware Reactive-Site Prediction*, bioRxiv (2025), https://doi.org/10.1101/2025.04.29.651260.
