## Supplementary material for "Localized Reactivity on Proteins as Riemannian Manifolds: A Quantum-Inspired Geometric Model for Deterministic, Metal-Aware Reactive-Site Prediction": full raw result

### Supplementary Information for:
### A Geometric and Statistical Framework for Protein Reactivity: Riemannian Manifolds Enable Deterministic, Generalizable Function Discovery

Haneul Park

December 30, 2025

#### Contents

**Supplementary Table S1: Full Benchmark Results** 2

**Supplementary Section S2: Case Studies** 14

#### Supplementary Table S1: Full Benchmark Results

Table 1: **Full Validation Results on DB5.5 Benchmark.** Metrics are reported for strict unbound, zero-shot evaluation under Policy P. (R: Receptor, L: Ligand)

| PDB ID | Chain/Role | ROC-AUC | AUPRC |
|---|---|---|---|
| 1A2K | L | 0.961 | 0.700 |
| 1A2K | R | 0.880 | 0.534 |
| 1ACB | L | 0.799 | 0.618 |
| 1ACB | R | 0.924 | 0.643 |
| 1AHW | L | 0.772 | 0.417 |
| 1AHW | R | 0.859 | 0.343 |
| 1AK4 | L | 0.971 | 0.750 |
| 1AK4 | R | 0.842 | 0.405 |
| 1AKJ | L | 0.895 | 0.618 |
| 1AKJ | R | 0.957 | 0.604 |
| 1ATN | L | 0.911 | 0.611 |
| 1ATN | R | 0.973 | 0.648 |
| 1AVX | L | 0.830 | 0.401 |
| 1AVX | R | 0.892 | 0.617 |
| 1AY7 | L | 0.673 | 0.516 |
| 1AY7 | R | 0.678 | 0.422 |
| 1AZS | L | 0.956 | 0.538 |
| 1AZS | R | 0.845 | 0.502 |
| 1B6C | L | 0.796 | 0.456 |
| 1B6C | R | 0.839 | 0.661 |
| 1BGX | L | 0.966 | 0.726 |
| 1BGX | R | 0.858 | 0.546 |
| 1BJ1 | L | 0.930 | 0.607 |
| 1BJ1 | R | 0.926 | 0.399 |
| 1BKD | L | 0.947 | 0.836 |
| 1BKD | R | 0.950 | 0.560 |
| 1BUH | L | 0.935 | 0.706 |
| 1BUH | R | 0.791 | 0.240 |
| 1BVK | L | 0.810 | 0.643 |
| 1BVK | R | 0.853 | 0.348 |
| 1BVN | L | 0.737 | 0.534 |
| 1BVN | R | 0.880 | 0.418 |
| 1CGI | L | 0.754 | 0.577 |
| 1CGI | R | 0.877 | 0.505 |
| 1CLV | L | 0.800 | 0.860 |
| 1CLV | R | 0.842 | 0.278 |
| 1D6R | L | 0.678 | 0.406 |
| 1D6R | R | 0.912 | 0.687 |
| 1DE4 | L | 0.964 | 0.569 |
| 1DE4 | R | 0.949 | 0.641 |
| 1DFJ | L | 0.899 | 0.432 |
| 1DFJ | R | 0.539 | 0.236 |

Continued on next page

**Table 1 – continued from previous page**

| PDB ID | Chain/Role | ROC-AUC | AUPRC |
|---|---|---|---|
| 1DQJ | L | 0.951 | 0.916 |
| 1DQJ | R | 0.968 | 0.684 |
| 1E4K | L | 0.775 | 0.369 |
| 1E4K | R | 0.887 | 0.435 |
| 1E6E | L | 0.895 | 0.702 |
| 1E6E | R | 0.953 | 0.658 |
| 1E6J | L | 0.787 | 0.640 |
| 1E6J | R | 0.950 | 0.440 |
| 1E96 | L | 0.834 | 0.297 |
| 1E96 | R | 0.978 | 0.822 |
| 1EAW | L | 0.834 | 0.703 |
| 1EAW | R | 0.882 | 0.552 |
| 1EER | L | 0.916 | 0.617 |
| 1EER | R | 0.867 | 0.696 |
| 1EFN | L | 0.740 | 0.642 |
| 1EFN | R | 0.746 | 0.442 |
| 1EWY | L | 0.836 | 0.519 |
| 1EWY | R | 0.898 | 0.762 |
| 1EXB | L | 0.973 | 0.780 |
| 1EXB | R | 0.962 | 0.714 |
| 1EZU | L | 0.818 | 0.341 |
| 1EZU | R | 0.853 | 0.557 |
| 1F34 | L | 0.769 | 0.576 |
| 1F34 | R | 0.913 | 0.536 |
| 1F51 | L | 0.904 | 0.620 |
| 1F51 | R | 0.956 | 0.789 |
| 1F6M | L | 0.790 | 0.540 |
| 1F6M | R | 0.796 | 0.345 |
| 1FAK | L | 0.672 | 0.312 |
| 1FAK | R | 0.878 | 0.436 |
| 1FC2 | L | 0.991 | 0.783 |
| 1FC2 | R | 0.819 | 0.656 |
| 1FCC | L | 0.823 | 0.735 |
| 1FCC | R | 0.929 | 0.562 |
| 1FFW | L | 0.818 | 0.427 |
| 1FFW | R | 0.695 | 0.235 |
| 1FLE | L | 0.736 | 0.614 |
| 1FLE | R | 0.796 | 0.437 |
| 1FQ1 | L | 0.868 | 0.336 |
| 1FQ1 | R | 0.846 | 0.481 |
| 1FQJ | L | 0.830 | 0.547 |
| 1FQJ | R | 0.960 | 0.692 |
| 1FSK | L | 0.936 | 0.755 |
| 1FSK | R | 0.843 | 0.425 |
| 1GCQ | L | 0.576 | 0.372 |
| 1GCQ | R | 0.931 | 0.564 |
| 1GHQ | L | 0.821 | 0.409 |

Continued on next page

**Table 1 – continued from previous page**

| PDB ID | Chain/Role | ROC-AUC | AUPRC |
|---|---|---|---|
| 1GHQ | R | 0.868 | 0.213 |
| 1GL1 | L | 0.505 | 0.523 |
| 1GL1 | R | 0.901 | 0.609 |
| 1GLA | L | 0.858 | 0.438 |
| 1GLA | R | 0.976 | 0.623 |
| 1GP2 | L | 0.741 | 0.425 |
| 1GP2 | R | 0.974 | 0.650 |
| 1GPW | L | 0.815 | 0.439 |
| 1GPW | R | 0.871 | 0.437 |
| 1GRN | L | 0.771 | 0.446 |
| 1GRN | R | 0.868 | 0.688 |
| 1GXD | L | 0.877 | 0.579 |
| 1GXD | R | 0.977 | 0.602 |
| 1H1V | L | 0.910 | 0.481 |
| 1H1V | R | 0.920 | 0.479 |
| 1H9D | L | 0.885 | 0.655 |
| 1H9D | R | 0.731 | 0.494 |
| 1HCF | L | 0.699 | 0.561 |
| 1HCF | R | 0.910 | 0.612 |
| 1HE1 | L | 0.970 | 0.845 |
| 1HE1 | R | 0.825 | 0.628 |
| 1HE8 | L | 0.932 | 0.552 |
| 1HE8 | R | 0.888 | 0.451 |
| 1HIA | L | 0.779 | 0.659 |
| 1HIA | R | 0.907 | 0.553 |
| 1I2M | L | 0.972 | 0.903 |
| 1I2M | R | 0.975 | 0.849 |
| 1I4D | L | 0.882 | 0.574 |
| 1I4D | R | 0.967 | 0.640 |
| 1I9R | L | 0.976 | 0.639 |
| 1I9R | R | 0.779 | 0.229 |
| 1IB1 | L | 0.883 | 0.719 |
| 1IB1 | R | 0.828 | 0.295 |
| 1IBR | L | 0.871 | 0.495 |
| 1IBR | R | 0.748 | 0.475 |
| 1IJK | L | 0.848 | 0.553 |
| 1IJK | R | 0.946 | 0.638 |
| 1IQD | L | 0.942 | 0.759 |
| 1IQD | R | 0.891 | 0.489 |
| 1IRA | L | 0.757 | 0.506 |
| 1IRA | R | 0.874 | 0.478 |
| 1J2J | L | 0.958 | 0.958 |
| 1J2J | R | 0.941 | 0.730 |
| 1JIW | L | 0.927 | 0.779 |
| 1JIW | R | 0.951 | 0.566 |
| 1JK9 | L | 0.869 | 0.806 |
| 1JK9 | R | 0.871 | 0.437 |

Continued on next page

**Table 1 – continued from previous page**

| PDB ID | Chain/Role | ROC-AUC | AUPRC |
|---|---|---|---|
| 1JMO | L | 0.863 | 0.621 |
| 1JMO | R | 0.962 | 0.565 |
| 1JPS | L | 0.860 | 0.505 |
| 1JPS | R | 0.891 | 0.467 |
| 1JTD | L | 0.785 | 0.253 |
| 1JTD | R | 0.942 | 0.606 |
| 1JTG | L | 0.940 | 0.720 |
| 1JTG | R | 0.856 | 0.493 |
| 1JWH | L | 0.946 | 0.490 |
| 1JWH | R | 0.938 | 0.565 |
| 1JZD | L | 0.766 | 0.645 |
| 1JZD | R | 0.925 | 0.518 |
| 1K4C | L | 0.934 | 0.509 |
| 1K4C | R | 0.904 | 0.427 |
| 1K5D | L | 0.884 | 0.617 |
| 1K5D | R | 0.780 | 0.563 |
| 1K74 | L | 0.795 | 0.386 |
| 1K74 | R | 0.890 | 0.615 |
| 1KAC | L | 0.761 | 0.433 |
| 1KAC | R | 0.836 | 0.474 |
| 1KKL | L | 0.871 | 0.711 |
| 1KKL | R | 0.809 | 0.235 |
| 1KLU | L | 0.735 | 0.308 |
| 1KLU | R | 0.918 | 0.583 |
| 1KTZ | L | 0.874 | 0.531 |
| 1KTZ | R | 0.966 | 0.733 |
| 1KXP | L | 0.837 | 0.407 |
| 1KXP | R | 0.936 | 0.567 |
| 1KXQ | L | 0.886 | 0.725 |
| 1KXQ | R | 0.877 | 0.581 |
| 1LFD | L | 0.911 | 0.864 |
| 1LFD | R | 0.861 | 0.514 |
| 1M10 | L | 0.943 | 0.785 |
| 1M10 | R | 0.877 | 0.532 |
| 1M27 | L | 0.779 | 0.362 |
| 1M27 | R | 0.771 | 0.252 |
| 1MAH | L | 0.730 | 0.608 |
| 1MAH | R | 0.850 | 0.338 |
| 1ML0 | L | 0.879 | 0.793 |
| 1ML0 | R | 0.942 | 0.541 |
| 1MLC | L | 0.724 | 0.329 |
| 1MLC | R | 0.969 | 0.578 |
| 1MQ8 | L | 0.860 | 0.408 |
| 1MQ8 | R | 0.923 | 0.524 |
| 1N2C | L | 0.953 | 0.695 |
| 1N2C | R | 0.929 | 0.362 |
| 1NCA | L | 0.882 | 0.364 |

Continued on next page

**Table 1 – continued from previous page**

| PDB ID | Chain/Role | ROC-AUC | AUPRC |
|---|---|---|---|
| 1NCA | R | 0.897 | 0.686 |
| 1NSN | L | 0.936 | 0.609 |
| 1NSN | R | 0.935 | 0.515 |
| 1NW9 | L | 0.836 | 0.678 |
| 1NW9 | R | 0.898 | 0.589 |
| 1OC0 | L | 0.858 | 0.862 |
| 1OC0 | R | 0.811 | 0.280 |
| 1OFU | L | 0.948 | 0.573 |
| 1OFU | R | 0.815 | 0.447 |
| 1OPH | L | 0.927 | 0.665 |
| 1OPH | R | 0.899 | 0.695 |
| 1OYV | L | 0.786 | 0.498 |
| 1OYV | R | 0.710 | 0.408 |
| 1PPE | L | 0.579 | 0.622 |
| 1PPE | R | 0.899 | 0.635 |
| 1PVH | L | 0.736 | 0.335 |
| 1PVH | R | 0.953 | 0.784 |
| 1PXV | L | 0.817 | 0.640 |
| 1PXV | R | 0.558 | 0.266 |
| 1QA9 | L | 0.954 | 0.619 |
| 1QA9 | R | 0.915 | 0.473 |
| 1QFW | L | 0.818 | 0.390 |
| 1QFW | R | 0.852 | 0.530 |
| 1R0R | L | 0.805 | 0.699 |
| 1R0R | R | 0.800 | 0.394 |
| 1R6Q | L | 0.833 | 0.722 |
| 1R6Q | R | 0.822 | 0.487 |
| 1R8S | L | 0.779 | 0.587 |
| 1R8S | R | 0.811 | 0.546 |
| 1RKE | L | 0.783 | 0.532 |
| 1RKE | R | 0.841 | 0.461 |
| 1RLB | L | 0.943 | 0.712 |
| 1RLB | R | 0.933 | 0.588 |
| 1RV6 | L | 0.826 | 0.395 |
| 1RV6 | R | 0.687 | 0.320 |
| 1S1Q | L | 0.831 | 0.721 |
| 1S1Q | R | 0.832 | 0.470 |
| 1S78 | L | 0.924 | 0.496 |
| 1S78 | R | 0.891 | 0.355 |
| 1SBB | L | 0.837 | 0.441 |
| 1SBB | R | 0.931 | 0.741 |
| 1SYX | L | 0.862 | 0.852 |
| 1SYX | R | 0.818 | 0.467 |
| 1T6B | L | 0.870 | 0.594 |
| 1T6B | R | 0.954 | 0.512 |
| 1TMQ | L | 0.785 | 0.475 |
| 1TMQ | R | 0.871 | 0.446 |

Continued on next page

**Table 1 – continued from previous page**

| PDB ID | Chain/Role | ROC-AUC | AUPRC |
|---|---|---|---|
| 1UDI | L | 0.830 | 0.790 |
| 1UDI | R | 0.883 | 0.612 |
| 1US7 | L | 0.993 | 0.957 |
| 1US7 | R | 0.911 | 0.665 |
| 1VFB | L | 0.771 | 0.532 |
| 1VFB | R | 0.893 | 0.554 |
| 1WDW | L | 0.885 | 0.577 |
| 1WDW | R | 0.914 | 0.487 |
| 1WEJ | L | 0.941 | 0.831 |
| 1WEJ | R | 0.880 | 0.388 |
| 1WQ1 | L | 0.946 | 0.842 |
| 1WQ1 | R | 0.903 | 0.548 |
| 1XD3 | L | 0.779 | 0.538 |
| 1XD3 | R | 0.822 | 0.494 |
| 1XQS | L | 0.757 | 0.403 |
| 1XQS | R | 0.948 | 0.672 |
| 1XU1 | L | 0.740 | 0.659 |
| 1XU1 | R | 0.790 | 0.230 |
| 1Y64 | L | 0.878 | 0.581 |
| 1Y64 | R | 0.941 | 0.646 |
| 1YVB | L | 0.913 | 0.704 |
| 1YVB | R | 0.735 | 0.312 |
| 1Z0K | L | 0.860 | 0.713 |
| 1Z0K | R | 0.910 | 0.680 |
| 1Z5Y | L | 0.837 | 0.564 |
| 1Z5Y | R | 0.891 | 0.577 |
| 1ZHH | L | 0.934 | 0.719 |
| 1ZHH | R | 0.819 | 0.256 |
| 1ZHI | L | 0.877 | 0.673 |
| 1ZHI | R | 0.783 | 0.426 |
| 1ZLI | L | 0.625 | 0.425 |
| 1ZLI | R | 0.852 | 0.416 |
| 1ZM4 | L | 0.870 | 0.544 |
| 1ZM4 | R | 0.871 | 0.317 |
| 2A1A | L | 0.714 | 0.375 |
| 2A1A | R | 0.969 | 0.618 |
| 2A5T | L | 0.900 | 0.477 |
| 2A5T | R | 0.946 | 0.646 |
| 2A9K | L | 0.890 | 0.691 |
| 2A9K | R | 0.827 | 0.491 |
| 2ABZ | L | 0.847 | 0.715 |
| 2ABZ | R | 0.773 | 0.381 |
| 2AJF | L | 0.807 | 0.468 |
| 2AJF | R | 0.788 | 0.194 |
| 2AYO | L | 0.732 | 0.575 |
| 2AYO | R | 0.766 | 0.409 |
| 2B42 | L | 0.761 | 0.546 |

Continued on next page

**Table 1 – continued from previous page**

| PDB ID | Chain/Role | ROC-AUC | AUPRC |
|---|---|---|---|
| 2B42 | R | 0.853 | 0.444 |
| 2B4J | L | 0.768 | 0.414 |
| 2B4J | R | 0.919 | 0.478 |
| 2BTF | L | 0.844 | 0.666 |
| 2BTF | R | 0.978 | 0.748 |
| 2C0L | L | 0.848 | 0.629 |
| 2C0L | R | 0.797 | 0.306 |
| 2CFH | L | 0.870 | 0.555 |
| 2CFH | R | 0.685 | 0.286 |
| 2DD8 | L | 0.874 | 0.655 |
| 2DD8 | R | 0.950 | 0.596 |
| 2FD6 | L | 0.855 | 0.492 |
| 2FD6 | R | 0.985 | 0.778 |
| 2FJG | L | 0.730 | 0.371 |
| 2FJG | R | 0.933 | 0.639 |
| 2FJU | L | 0.914 | 0.452 |
| 2FJU | R | 0.964 | 0.651 |
| 2G77 | L | 0.953 | 0.884 |
| 2G77 | R | 0.801 | 0.289 |
| 2GAF | L | 0.776 | 0.313 |
| 2GAF | R | 0.934 | 0.577 |
| 2GTP | L | 0.844 | 0.527 |
| 2GTP | R | 0.974 | 0.836 |
| 2H7V | L | 0.895 | 0.566 |
| 2H7V | R | 0.903 | 0.508 |
| 2HLE | L | 0.892 | 0.732 |
| 2HLE | R | 0.715 | 0.433 |
| 2HMI | L | 0.913 | 0.509 |
| 2HMI | R | 0.910 | 0.612 |
| 2HQS | L | 0.788 | 0.516 |
| 2HQS | R | 0.879 | 0.478 |
| 2HRK | L | 0.780 | 0.534 |
| 2HRK | R | 0.860 | 0.622 |
| 2I25 | L | 0.665 | 0.347 |
| 2I25 | R | 0.753 | 0.312 |
| 2I9B | L | 0.933 | 0.893 |
| 2I9B | R | 0.872 | 0.356 |
| 2IDO | L | 0.763 | 0.526 |
| 2IDO | R | 0.941 | 0.641 |
| 2J0T | L | 0.884 | 0.735 |
| 2J0T | R | 0.704 | 0.404 |
| 2J7P | L | 0.924 | 0.577 |
| 2J7P | R | 0.957 | 0.719 |
| 2JEL | L | 0.755 | 0.410 |
| 2JEL | R | 0.938 | 0.414 |
| 2MTA | L | 0.828 | 0.509 |
| 2MTA | R | 0.844 | 0.404 |

Continued on next page

**Table 1 – continued from previous page**

| PDB ID | Chain/Role | ROC-AUC | AUPRC |
|---|---|---|---|
| 2NZ8 | L | 0.874 | 0.530 |
| 2NZ8 | R | 0.837 | 0.613 |
| 2O3B | L | 0.796 | 0.581 |
| 2O3B | R | 0.800 | 0.371 |
| 2O8V | L | 0.607 | 0.355 |
| 2O8V | R | 0.911 | 0.491 |
| 2OOB | L | 0.812 | 0.618 |
| 2OOB | R | 0.437 | 0.290 |
| 2OOR | L | 0.871 | 0.626 |
| 2OOR | R | 0.952 | 0.441 |
| 2OT3 | L | 0.770 | 0.548 |
| 2OT3 | R | 0.868 | 0.575 |
| 2OUL | L | 0.899 | 0.561 |
| 2OUL | R | 0.778 | 0.353 |
| 2OZA | L | 0.948 | 0.801 |
| 2OZA | R | 0.906 | 0.715 |
| 2PCC | L | 0.948 | 0.926 |
| 2PCC | R | 0.882 | 0.373 |
| 2SIC | L | 0.910 | 0.716 |
| 2SIC | R | 0.795 | 0.394 |
| 2SNI | L | 0.893 | 0.804 |
| 2SNI | R | 0.756 | 0.283 |
| 2UUY | L | 0.885 | 0.755 |
| 2UUY | R | 0.886 | 0.632 |
| 2VDB | L | 0.800 | 0.715 |
| 2VDB | R | 0.837 | 0.196 |
| 2VIS | L | 0.919 | 0.431 |
| 2VIS | R | 0.901 | 0.449 |
| 2VXT | L | 0.912 | 0.633 |
| 2VXT | R | 0.926 | 0.446 |
| 2W9E | L | 0.956 | 0.668 |
| 2W9E | R | 0.884 | 0.330 |
| 2X9A | L | 0.743 | 0.577 |
| 2X9A | R | 0.895 | 0.758 |
| 2YVJ | L | 0.868 | 0.534 |
| 2YVJ | R | 0.941 | 0.435 |
| 2Z0E | L | 0.824 | 0.623 |
| 2Z0E | R | 0.888 | 0.689 |
| 3A4S | L | 0.834 | 0.629 |
| 3A4S | R | 0.741 | 0.308 |
| 3AAA | L | 0.846 | 0.501 |
| 3AAA | R | 0.918 | 0.379 |
| 3AAD | L | 0.824 | 0.455 |
| 3AAD | R | 0.834 | 0.405 |
| 3BIW | L | 0.854 | 0.297 |
| 3BIW | R | 0.990 | 0.842 |
| 3BP8 | L | 0.706 | 0.566 |

Continued on next page

**Table 1 – continued from previous page**

| PDB ID | Chain/Role | ROC-AUC | AUPRC |
|---|---|---|---|
| 3BP8 | R | 0.887 | 0.354 |
| 3BX7 | L | 0.754 | 0.648 |
| 3BX7 | R | 0.912 | 0.770 |
| 3CPH | L | 0.979 | 0.789 |
| 3CPH | R | 0.885 | 0.225 |
| 3D5S | L | 0.800 | 0.630 |
| 3D5S | R | 0.877 | 0.349 |
| 3DAW | L | 0.840 | 0.616 |
| 3DAW | R | 0.914 | 0.560 |
| 3EO1 | L | 0.671 | 0.152 |
| 3EO1 | R | 0.936 | 0.421 |
| 3EOA | L | 0.882 | 0.436 |
| 3EOA | R | 0.979 | 0.776 |
| 3F1P | L | 0.741 | 0.356 |
| 3F1P | R | 0.789 | 0.603 |
| 3FN1 | L | 0.848 | 0.687 |
| 3FN1 | R | 0.858 | 0.587 |
| 3G6D | L | 0.841 | 0.577 |
| 3G6D | R | 0.938 | 0.618 |
| 3H11 | L | 0.906 | 0.678 |
| 3H11 | R | 0.752 | 0.339 |
| 3H2V | L | 0.849 | 0.584 |
| 3H2V | R | 0.840 | 0.319 |
| 3HI6 | L | 0.830 | 0.369 |
| 3HI6 | R | 0.973 | 0.715 |
| 3HMX | L | 0.965 | 0.550 |
| 3HMX | R | 0.942 | 0.455 |
| 3K75 | L | 0.869 | 0.523 |
| 3K75 | R | 0.938 | 0.434 |
| 3L5W | L | 0.940 | 0.617 |
| 3L5W | R | 0.874 | 0.264 |
| 3L89 | L | 0.799 | 0.495 |
| 3L89 | R | 0.932 | 0.516 |
| 3LVK | L | 0.953 | 0.902 |
| 3LVK | R | 0.914 | 0.499 |
| 3MJ9 | L | 0.836 | 0.318 |
| 3MJ9 | R | 0.918 | 0.540 |
| 3MXW | L | 0.886 | 0.662 |
| 3MXW | R | 0.815 | 0.244 |
| 3P57 | L | 0.921 | 0.659 |
| 3P57 | R | 0.905 | 0.486 |
| 3PC8 | L | 0.756 | 0.400 |
| 3PC8 | R | 0.799 | 0.444 |
| 3R9A | L | 0.814 | 0.393 |
| 3R9A | R | 0.989 | 0.617 |
| 3RJQ | L | 0.832 | 0.337 |
| 3RJQ | R | 0.768 | 0.393 |

Continued on next page

**Table 1 – continued from previous page**

| PDB ID | Chain/Role | ROC-AUC | AUPRC |
|---|---|---|---|
| 3RVW | L | 0.861 | 0.442 |
| 3RVW | R | 0.937 | 0.612 |
| 3S9D | L | 0.793 | 0.478 |
| 3S9D | R | 0.912 | 0.541 |
| 3SE8 | L | 0.958 | 0.711 |
| 3SE8 | R | 0.945 | 0.657 |
| 3SGQ | L | 0.713 | 0.570 |
| 3SGQ | R | 0.707 | 0.375 |
| 3SZK | L | 0.666 | 0.302 |
| 3SZK | R | 0.895 | 0.315 |
| 3U7Y | L | 0.940 | 0.682 |
| 3U7Y | R | 0.947 | 0.715 |
| 3V6Z | L | 0.808 | 0.439 |
| 3V6Z | R | 0.850 | 0.508 |
| 3VLB | L | 0.758 | 0.336 |
| 3VLB | R | 0.788 | 0.340 |
| 3WD5 | L | 0.895 | 0.383 |
| 3WD5 | R | 0.957 | 0.597 |
| 4CPA | L | 0.844 | 0.562 |
| 4CPA | R | 0.865 | 0.502 |
| 4DN4 | L | 0.896 | 0.622 |
| 4DN4 | R | 0.911 | 0.439 |
| 4DW2 | L | 0.847 | 0.431 |
| 4DW2 | R | 0.929 | 0.518 |
| 4ETQ | L | 0.873 | 0.540 |
| 4ETQ | R | 0.916 | 0.541 |
| 4FP8 | L | 0.927 | 0.294 |
| 4FP8 | R | 0.574 | 0.019 |
| 4FQI | L | 0.894 | 0.219 |
| 4FQI | R | 0.977 | 0.600 |
| 4FZA | L | 0.878 | 0.296 |
| 4FZA | R | 0.839 | 0.434 |
| 4G6J | L | 0.811 | 0.400 |
| 4G6J | R | 0.960 | 0.653 |
| 4G6M | L | 0.902 | 0.580 |
| 4G6M | R | 0.928 | 0.421 |
| 4GAM | L | 0.837 | 0.807 |
| 4GAM | R | 0.936 | 0.470 |
| 4GXU | L | 0.890 | 0.142 |
| 4GXU | R | 0.954 | 0.685 |
| 4H03 | L | 0.892 | 0.440 |
| 4H03 | R | 0.932 | 0.482 |
| 4HX3 | L | 0.851 | 0.635 |
| 4HX3 | R | 0.872 | 0.460 |
| 4IZ7 | L | 0.911 | 0.640 |
| 4IZ7 | R | 0.796 | 0.254 |
| 4JCV | L | 0.764 | 0.400 |

Continued on next page

**Table 1 – continued from previous page**

| PDB ID | Chain/Role | ROC-AUC | AUPRC |
|---|---|---|---|
| 4JCV | R | 0.891 | 0.267 |
| 4LW4 | L | 0.749 | 0.382 |
| 4LW4 | R | 0.916 | 0.555 |
| 4M3K | L | 0.810 | 0.417 |
| 4M3K | R | 0.794 | 0.505 |
| 4M5Z | L | 0.678 | 0.253 |
| 4M5Z | R | 0.926 | 0.475 |
| 4M76 | L | 0.988 | 0.863 |
| 4M76 | R | 0.790 | 0.206 |
| 4POU | L | 0.869 | 0.513 |
| 4POU | R | 0.886 | 0.526 |
| 4Y7M | L | 0.849 | 0.389 |
| 4Y7M | R | 0.827 | 0.383 |
| 5C7X | L | 0.642 | 0.419 |
| 5C7X | R | 0.883 | 0.296 |
| 5CBA | L | 0.637 | 0.388 |
| 5CBA | R | 0.682 | 0.273 |
| 5E5M | L | 0.888 | 0.572 |
| 5E5M | R | 0.853 | 0.546 |
| 5GRJ | L | 0.939 | 0.625 |
| 5GRJ | R | 0.935 | 0.453 |
| 5HGG | L | 0.906 | 0.570 |
| 5HGG | R | 0.934 | 0.725 |
| 5HYS | L | 0.853 | 0.250 |
| 5HYS | R | 0.729 | 0.375 |
| 5JMO | L | 0.933 | 0.555 |
| 5JMO | R | 0.813 | 0.606 |
| 5KOV | L | 0.897 | 0.483 |
| 5KOV | R | 0.953 | 0.595 |
| 5O14 | L | 0.907 | 0.534 |
| 5O14 | R | 0.910 | 0.414 |
| 5O1R | L | 0.894 | 0.646 |
| 5O1R | R | 0.930 | 0.413 |
| 5SV3 | L | 0.904 | 0.469 |
| 5SV3 | R | 0.784 | 0.451 |
| 5VNW | L | 0.993 | 0.845 |
| 5VNW | R | 0.840 | 0.595 |
| 5WHK | L | 0.822 | 0.403 |
| 5WHK | R | 0.953 | 0.536 |
| 5WK3 | L | 0.902 | 0.701 |
| 5WK3 | R | 0.855 | 0.382 |
| 5WUX | L | 0.899 | 0.507 |
| 5WUX | R | 0.814 | 0.396 |
| 5X0T | L | 0.860 | 0.613 |
| 5X0T | R | 0.842 | 0.260 |
| 5Y9J | L | 0.688 | 0.242 |
| 5Y9J | R | 0.929 | 0.495 |

Continued on next page

**Table 1 – continued from previous page**

| PDB ID | Chain/Role | ROC-AUC | AUPRC |
|---|---|---|---|
| 6A0Z | L | 0.920 | 0.482 |
| 6A0Z | R | 0.933 | 0.611 |
| 6A77 | L | 0.836 | 0.603 |
| 6A77 | R | 0.945 | 0.540 |
| 6AL0 | L | 0.000 | 0.000 |
| 6AL0 | R | 0.971 | 0.682 |
| 6B0S | L | 0.745 | 0.588 |
| 6B0S | R | 0.850 | 0.344 |
| 6BPC | L | 0.889 | 0.313 |
| 6BPC | R | 0.904 | 0.426 |
| 6CWG | L | 0.913 | 0.489 |
| 6CWG | R | 0.839 | 0.444 |
| 6DBG | L | 0.836 | 0.452 |
| 6DBG | R | 0.824 | 0.459 |
| 6EY6 | L | 0.917 | 0.522 |
| 6EY6 | R | 0.772 | 0.406 |
| 6OC3 | L | 0.730 | 0.286 |
| 6OC3 | R | 0.905 | 0.522 |
| 7CEI | L | 0.582 | 0.222 |
| 7CEI | R | 0.721 | 0.448 |
| 9QFW | L | 0.678 | 0.176 |
| 9QFW | R | 0.873 | 0.579 |
| BAAD | L | 0.882 | 0.537 |
| BAAD | R | 0.829 | 0.499 |
| BOYV | L | 0.933 | 0.686 |
| BOYV | R | 0.829 | 0.435 |
| BP57 | L | 0.873 | 0.624 |
| BP57 | R | 0.844 | 0.444 |
| CP57 | L | 0.896 | 0.579 |
| CP57 | R | 0.879 | 0.819 |
| **MEAN** | **All** | **0.858** | **0.529** |

#### Supplementary Section S2: Detailed Performance on Mechanistic Case Studies

In this section, we present the comprehensive evaluation results for all five mechanistic case studies: Rubisco (4RUB), GroEL/ES (2C7C), p53 (1TSR), SecA (1TF5), and the Ribosome rescue complex (6Q97).

Table S2 provides the full performance breakdown for every functional subset defined in the study. Table S3 explicitly lists the ground-truth residue indices used for these evaluations to ensure full reproducibility.

Table 2: **Table S2: Comprehensive Performance Breakdown for All Case Studies.** Evaluation follows Policy P. *Group AUC* denotes the strict grouped-CV metric when grouping is enabled; entries marked with † report non-grouped CV AUC as indicated by the evaluation log (`use_groupkfold=False`).

| **Complex / Subset** | **N (Pos/Neg)** | **Stratified AUC** | **Group AUC** |
|---|---|---|---|
| **1. Rubisco (4RUB)** | | | |
| L_Tier1_catalytic_core | 56 / 2308 | $0.999 \pm 0.002$ | $\mathbf{0.938 \pm 0.120}$ |
| L_Tier1_gate_latch | 12 / 2352 | $1.000 \pm 0.000$ | $0.477 \pm 0.002$ |
| L_Tier1_PPI_Rca_core | 0 / 2364 | N/A | N/A |
| L_Tier2_catalytic_shell | 28 / 2336 | $0.999 \pm 0.001$ | $\mathbf{0.944 \pm 0.089}$ |
| S_Tier1_helix8_cluster | 24 / 2340 | $0.999 \pm 0.000$ | $0.761 \pm 0.236$ |
| S_Tier1_interface_betaAB_core | 12 / 2352 | $0.999 \pm 0.001$ | $0.894 \pm 0.076$ |
| S_Tier2_betaAB_regulatory | 20 / 2344 | $0.945 \pm 0.033$ | $0.676 \pm 0.201$ |
| **Global Tier1 All** | 96 / 2268 | $0.984 \pm 0.031$ | $\mathbf{0.800 \pm 0.116}$ |
| **Global Tier2 All** | 68 / 2296 | $0.982 \pm 0.034$ | $\mathbf{0.725 \pm 0.158}$ |
| L_Type_catalytic_T1T2 | 84 / 2280 | $0.998 \pm 0.002$ | $\mathbf{0.938 \pm 0.128}$ |
| L_Type_PTM_T1T2 | 12 / 2352 | $0.993 \pm 0.015$ | $0.449 \pm 0.012$ |
| **2. GroEL/ES (2C7C)** | | | |
| **Global Tier1 All** | 441 / 7546 | N/A | $\mathbf{0.7717 \pm 0.0587}$ |
| GroEL_Tier1_All (EL-only) | 378 / 6958 | N/A | $0.8022 \pm 0.0460$ |
| GroES_Tier1_All (ES-only) | 63 / 588 | N/A | $0.9500 \pm 0.0290$ |
| groel_A_ATP_core | 140 / 7196 | N/A | $0.8713 \pm 0.0558$ |
| groel_B_substrate_patch | 112 / 7224 | N/A | $0.8924 \pm 0.0969$ |
| groel_C_hinge_core | 14 / 7322 | N/A | N/A |
| groel_C_hinge_support | 28 / 7308 | N/A | $0.9375 \pm 0.0526$ |
| groel_D_inter_ring | 84 / 7252 | N/A | $0.6796 \pm 0.2305$ |
| groes_E_IVL_core | 21 / 630 | N/A | $0.9317 \pm 0.0623$ |
| groes_F_loop_support | 42 / 609 | N/A | $0.8857 \pm 0.0399$ |
| **3. p53 (1TSR)** | | | |
| **p53_Global_Tier1** | 102 / 486 | N/A | $\mathbf{0.7766 \pm 0.1038}$ |
| **p53_Global_Tier1_plus_Tier2** | 126 / 462 | N/A | $\mathbf{0.7937 \pm 0.0791}$ |
| p53_Tier1a_DNA_contact_core | 24 / 564 | N/A | $0.9232 \pm 0.0390$ |
| p53_Tier1b_Zn_structural_cluster | 15 / 573 | N/A | $0.8785 \pm 0.1543$ |
| p53_Tier1c_fitness_stability_core | 15 / 573 | N/A | $0.7083 \pm 0.1772$ |
| p53_Tier1d_allosteric_PPI_pivots | 48 / 540 | N/A | $0.7605 \pm 0.1428$ |
| p53_Tier2_structural_shell | 24 / 564 | N/A | $0.7994 \pm 0.1071$ |
| DNA_Tier1_DNA_contact_core | 3 / 39 | N/A | $0.3622 \pm 0.0545$ |
| DNA_Tier2_DNA_contact_shell | 8 / 34 | N/A | $0.6944 \pm 0.1504$ |

Continued on next page

**Table 2 – continued**

| Complex / Subset | N (Pos/Neg) | Stratified AUC | Group AUC |
|---|---|---|---|
| DNA_Tier1_plus_Tier2_DNA | 11 / 31 | N/A | $0.8829 \pm 0.0434$ |
| **4. SecA (1TF5)** | | | |
| Tier1_core_ATPase | 6 / 769 | N/A | $0.9628 \pm 0.0292$ |
| Tier1_core_clamp | 6 / 769 | N/A | $0.8818 \pm 0.2266$ |
| Tier1_core_interface | 7 / 768 | N/A | $0.7726 \pm 0.2530$ |
| **Tier1_core_all** | 19 / 756 | N/A | $\mathbf{0.9406 \pm 0.0660}$ |
| Tier2_support_ATPase | 35 / 740 | N/A | $0.9606 \pm 0.0344$ |
| Tier2_support_clamp | 20 / 755 | N/A | $0.9512 \pm 0.0549$ |
| Tier2_support_interface | 94 / 681 | N/A | $0.9580 \pm 0.0155$ |
| **Tier2_support_all** | 149 / 626 | N/A | $\mathbf{0.9330 \pm 0.0280}$ |
| Tier1plus2_ATPase | 41 / 734 | N/A | $0.9792 \pm 0.0076$ |
| Tier1plus2_clamp | 26 / 749 | N/A | $0.9792 \pm 0.0099$ |
| Tier1plus2_interface | 101 / 674 | N/A | $0.9568 \pm 0.0173$ |
| **Tier1plus2_all** | 168 / 607 | N/A | $\mathbf{0.9427 \pm 0.0265}$ |
| **5. Ribosome rescue complex (6Q97)** | | | |
| **Tier1_all** | 64 / 11554 | N/A | $\mathbf{0.8912 \pm 0.0377}^\dagger$ |
| **Tier1_plus_Tier2_all** | 150 / 11468 | N/A | $\mathbf{0.9420 \pm 0.0367}^\dagger$ |
| Tier1_vs_hard_negative | 64 / 323 | N/A | $0.9803 \pm 0.0045^\dagger$ |
| Tier1plus2_vs_hard_negative | 150 / 323 | N/A | $0.9837 \pm 0.0125^\dagger$ |
| Type_SmpB_Tier1 | 15 / 11603 | N/A | $0.9271 \pm 0.0811^\dagger$ |
| Type_SmpB_Tier1plus2 | 33 / 11585 | N/A | $0.9336 \pm 0.0609^\dagger$ |
| Type_tmRNA_Tier1 | 14 / 11604 | N/A | $0.8620 \pm 0.1687^\dagger$ |
| Type_tmRNA_Tier1plus2 | 43 / 11575 | N/A | $0.9472 \pm 0.0441^\dagger$ |
| Type_16S_Tier1 | 9 / 11609 | N/A | $0.7903 \pm 0.1862^\dagger$ |
| Type_16S_Tier1plus2 | 11 / 11607 | N/A | $0.6526 \pm 0.1385^\dagger$ |
| Type_23S_Tier1 | 4 / 11614 | N/A | $0.6193 \pm 0.2191^\dagger$ |
| Type_23S_Tier1plus2 | 7 / 11611 | N/A | $0.9469 \pm 0.1025^\dagger$ |
| Type_uS3_Tier1 | 11 / 11607 | N/A | $0.9441 \pm 0.1007^\dagger$ |
| Type_uS3_Tier1plus2 | 32 / 11586 | N/A | $0.9820 \pm 0.0285^\dagger$ |
| Type_uS4_Tier1 | 4 / 11614 | N/A | $0.7440 \pm 0.2500^\dagger$ |
| Type_uS4_Tier1plus2 | 7 / 11611 | N/A | $0.8450 \pm 0.2018^\dagger$ |
| Type_uS5_Tier1 | 7 / 11611 | N/A | $0.9962 \pm 0.0032^\dagger$ |
| Type_uS5_Tier1plus2 | 17 / 11601 | N/A | $0.9382 \pm 0.0737^\dagger$ |

**Notes.** † indicates that the evaluation log reported `use_groupkfold=False`, i.e. the reported AUC is from non-grouped CV rather than GroupKFold. This is a strictness downgrade relative to grouped evaluation and should be interpreted accordingly. For small positive sets (e.g. some 1TSR/DNA subsets), folds with single-class test splits can be skipped; the reported mean±sd is computed over the valid folds only, as recorded in the evaluation log.

Table 3: **Table S3: Ground Truth Label Definitions for All Case Studies.** Residue indices correspond to the canonical sequences in the respective PDB structures. Lists are formatted as ranges when possible.

| **Complex / Subset** | **Residue Indices (Positive Labels)** |
|---|---|
| **1. Rubisco (4RUB)** | |
| L_Tier1_catalytic_core | 20, 60, 65, 123, 175, 177, 201, 203-204, 294-295, 327, 334-335 |
| L_Tier1_gate_latch | 327, 334, 473 |
| L_Tier2_catalytic_shell | 66, 379-381, 402-404 |
| S_Tier1_helix8_cluster | 43, 73, 78-79, 81, 92 |
| S_Tier1_interface_betaAB_core | 16, 18, 32 |
| S_Tier2_betaAB_regulatory | 59, 67-69, 71 |
| **2. GroEL/ES (2C7C)** | |
| Global_Tier1_All | GroEL: 30-34, 51-52, 87, 89, 105, 192, 197, 199, 201, 203-204, 234, 237, 263-264, 374-375, 386, 398, 434, 452, 461; GroES: 23-31 |
| GroEL_Tier1_All (EL-only) | 30-34, 51-52, 87, 89, 105, 192, 197, 199, 201, 203-204, 234, 237, 263-264, 374-375, 386, 398, 434, 452, 461 |
| GroES_Tier1_All (ES-only) | 23-31 |
| groel_A_ATP_core | 30-34, 51-52, 87, 89, 398 |
| groel_B_substrate_patch | 199, 201, 203-204, 234, 237, 263-264 |
| groel_C_hinge_core | 192 |
| groel_C_hinge_support | 374-375 |
| groel_D_inter_ring | 105, 197, 386, 434, 452, 461 |
| groes_E_IVL_core | 25-27 |
| groes_F_loop_support | 23-24, 28-31 |
| **3. p53 (1TSR)** | |
| p53_Tier1a_DNA_contact_core | 120, 241, 248, 273, 276-277, 280, 283 |
| p53_Tier1b_Zn_structural_cluster | 171, 176, 179, 238, 242 |
| p53_Tier1c_fitness_stability_core | 175, 220, 245, 249, 282 |
| p53_Tier1d_allosteric_PPI_pivots | 100, 104, 107, 178, 180-181, 207-213, 243, 247, 268 |
| p53_Tier2_structural_shell | 109, 119, 121, 145-146, 246, 281, 285 |
| p53_Tier1_all | 100, 104, 107, 120, 171, 175-176, 178-181, 207-213, 220, 238, 241-243, 245, 247-249, 268, 273, 276-277, 280, 282-283 |
| p53_Tier1_plus_Tier2 | 100, 104, 107, 109, 119-121, 145-146, 171, 175-176, 178-181, 207-213, 220, 238, 241-243, 245-246, 247-249, 268, 273, 276-277, 280, 281-283, 285 |
| DNA_Tier1_DNA_contact_core | E:12; F:7, F:9 |
| DNA_Tier2_DNA_contact_shell | E:1, E:11, E:13-14; F:6, F:8, F:14, F:16 |
| DNA_Tier1_plus_Tier2_DNA | E:1, E:11-14; F:6-9, F:14, F:16 |
| **4. SecA (1TF5)** | |
| Tier1_core_ATPase | 106-107, 207-208, 215-216 |
| Tier2_support_ATPase | 100-105, 108-119, 200-206, 209-214, 217-220 |
| Tier1_core_clamp | 357, 360-361, 381, 384-385 |
| Tier2_support_clamp | 352-356, 358-359, 362, 375-380, 382-383, 386-389 |
| Tier1_core_interface | 590, 599, 611, 615, 642, 652, 665 |

Continued on next page

**Table 3 – continued**

| **Complex / Subset** | **Residue Indices** |
|---|---|
| Tier2_support_interface | 571-589, 591-598, 600-610, 612-614, 616-619, 623-641, 643-651, 653-664, 666-674 |
| Tier1_core_all | 106-107, 207-208, 215-216, 357, 360-361, 381, 384-385, 590, 599, 611, 615, 642, 652, 665 |
| Tier1plus2_all | 100-119, 200-220, 352-362, 375-389, 571-619, 623-674 |
| **5. Ribosome rescue complex (6Q97)** | |
| Tier1_all | SmpB: 5:HIS22, 5:GLY132, 5:LYS133, 5:LYS134, 5:HIS136, 5:ASP137, 5:LYS138, 5:ARG139, 5:LYS143, 5:ARG145, 5:TRP147, 5:ARG153, 5:ILE154, 5:MET155, 5:LYS156; tmRNA: 4:U119, 4:U120, 4:A121, 4:C127, 4:U128, 4:G129, 4:U131, 4:C183, 4:A184, 4:A185, 4:A186, 4:C361, 4:C362, 4:A363; 16S: 2:G530, 2:A1492, 2:A1493, 2:G693, 2:A790, 2:G926, 2:C1399, 2:C1400, 2:G1401; 23S (PTC): 1:A2451, 1:U2506, 1:U2585, 1:A2602; uS3: h:ARG72, h:PRO73, h:ILE77, h:LYS79, h:LYS80, h:ARG131, h:ARG132, h:LYS135, h:ARG136, h:ASN140, h:LEU144; uS4: i:ARG44, i:ARG47, i:ARG49, i:ARG50; uS5: j:ARG20, j:PHE31, j:PHE33, j:GLU55, j:VAL56, j:ILE60, j:GLN61 |
| Tier2_shells (by role) | SmpB shells: 5:SER20, 5:GLY21, 5:THR23, 5:THR24, 5:LYS25, 5:ARG26, 5:SER140, 5:ASP141, 5:ILE142, 5:GLU144, 5:GLU146, 5:GLN148, 5:VAL149, 5:ASP150, 5:LYS151, 5:ALA152, 5:ASN157, 5:ALA158; tmRNA shells: 4:U115, 4:G116, 4:C117, 4:A118, 4:A122, 4:U123, 4:G124, 4:G125, 4:A126, 4:U130, 4:A132, 4:G175, 4:A176, 4:C177, 4:G178, 4:A179, 4:A180, 4:G181, 4:U182, 4:A187, 4:C188, 4:G189, 4:G190, 4:C355, 4:G356, 4:G357, 4:U358, 4:C359, 4:C360; 16S shell: 2:C518, 2:C1195; 23S shell: 1:C2452, 1:A2601, 1:G2603; uS3 shells: h:LYS68, h:ARG69, h:GLU70, h:GLU71, h:LEU74, h:TYR75, h:GLN76, h:LYS78, h:GLU81, h:GLU82, h:LEU128, h:ARG129, h:GLY130, h:LYS133, h:GLY134, h:ARG137, h:ALA138, h:ALA139, h:LYS141, h:GLY142, h:THR145; uS4 shell: i:GLU35, i:ALA43, i:LYS45; uS5 shells: j:LYS28, j:THR29, j:GLU30, j:GLY32, j:ALA34, j:LYS52, j:ALA53, j:ARG54, j:PRO57, j:ALA58 |

Continued on next page

**Table 3 – continued**

| Complex / Subset | Residue Indices |
|---|---|
| Hard_Negatives (N=323) | 1:A1689, 1:A1701, 1:A1819, 1:A705, 1:G1695, 1:G1702, 1:G1799, 1:U1818, 1:U1820, 2:A246, B:ARG69, B:ASN143, B:HIS142, B:LEU105, B:THR191, B:VAL195, D:ARG102, D:ARG114, D:ARG21, D:ARG44, D:ARG49, D:ARG61, D:ARG67, D:ARG79, D:ARG88, D:ASP116, D:ASP145, D:ASP154, D:ASP22, D:ASP91, D:GLU111, D:GLU122, D:GLU127, D:GLU152, D:GLU16, D:GLU25, D:GLU51, D:HIS92, D:LYS106, D:LYS130, D:LYS132, D:LYS139, D:LYS47, D:LYS57, D:LYS58, D:LYS63, D:LYS74, D:LYS95, D:PHE124, D:PHE158, D:PHE19, D:TYR35, E:ASP10, E:ASP6, E:GLU11, E:GLU19, E:HIS5, E:LYS14, E:LYS15, E:LYS3, E:LYS9, E:PHE20, E:TYR22, E:TYR7, E:TYR8, H:ARG125, H:ARG31, H:ARG42, H:ARG53, H:ARG56, H:ARG61, H:ARG94, H:ASP124, H:ASP29, H:ASP36, H:ASP7, H:ASP74, H:GLU107, H:GLU116, H:GLU14, H:GLU17, H:GLU47, H:GLU65, H:GLU87, H:LYS101, H:LYS105, H:LYS109, H:LYS20, H:LYS43, H:LYS73, H:LYS8, H:LYS97, H:PHE106, H:PHE69, H:PHE76, H:PHE99, I:ARG102, I:ASP115, I:ASP120, I:ASP46, I:ASP63, I:GLU107, I:GLU122, I:LYS44, I:LYS50, I:LYS80, I:LYS9, I:LYS96, I:PHE37, I:PHE66, I:PHE68, I:TYR7, J:ARG120, J:ARG27, J:ARG34, J:ARG37, J:ARG69, J:ARG95, J:ARG96, J:ARG99, J:ASP14, J:ASP19, J:ASP49, J:ASP52, J:ASP60, J:ASP71, J:GLU102, J:GLU31, J:GLU98, J:HIS130, J:HIS132, J:HIS40, J:HIS47, J:HIS76, J:HIS77, J:LYS106, J:LYS12, J:LYS121, J:LYS123, J:LYS23, J:LYS39, J:LYS61, J:LYS68, J:LYS85, J:PHE119, J:PHE4, J:PHE89, J:TYR44, J:TYR74, K:ALA11, K:ALA83, K:ARG108, K:ARG30, K:ARG31, K:ARG49, K:ARG64, K:ARG70, K:ARG71, K:ARG78, K:ASP12, K:ASP80, K:GLU106, K:GLU110, K:GLU4, K:GLU45, K:GLU92, K:HIS29, K:LEU8, K:LYS111, K:LYS114, K:LYS40, K:LYS44, K:LYS53, K:LYS54, K:LYS59, K:LYS66, K:LYS67, K:PHE100, K:PHE112, K:PHE79, K:VAL63, L:ARG123, L:ARG132, L:ARG18, L:ARG21, L:ARG33, L:ARG41, L:ARG59, L:ARG69, L:ARG78, L:ASP91, L:GLU10, L:GLU115, L:GLU136, L:GLU144, L:GLU76, L:GLU86, L:HIS35, L:LYS109, L:LYS141, L:LYS17, L:LYS29, L:LYS63, L:LYS70, L:LYS96, L:PHE107, L:PHE50, L:PHE66, M:ARG50, M:ARG51, M:LYS123, M:LYS71, M:LYS8, M:PHE68, N:ARG46, N:ARG8, N:GLU43, N:HIS31, N:LYS42, N:LYS56, N:TYR94, O:ARG102, O:ARG81, O:ARG9, O:ARG94, O:GLU112, O:GLU80, O:TYR64, O:TYR99, P:ARG39, P:ARG72, P:GLN75, P:GLU27, P:GLU34, P:GLU68, P:GLY23, P:HIS77, P:LYS106, P:LYS111, P:LYS37, P:LYS96, P:TYR99, P:VAL70, Q:ARG30, Q:ARG51, Q:ARG53, Q:ARG64, Q:ARG70, Q:ASP49, Q:GLU111, Q:LYS114, Q:LYS54, Q:LYS78, R:ARG13, R:ARG21, R:ARG78, R:ARG84, R:GLU70, R:LYS48, R:LYS60, R:PHE35, R:TYR2, S:ARG11, S:ARG18, S:ARG8, S:ARG84, S:ARG92, S:ASP34, S:ASP62, S:GLU59, S:LYS16, S:LYS6, S:TYR38, T:ARG12, T:ARG3, T:ARG69, T:GLU4, T:GLU42, T:GLU5, T:GLU56, T:LYS40, T:LYS64, U:ARG7, U:ARG94, U:ASP81, U:ASP9, U:GLU10, U:GLU88, U:LYS17, U:LYS33, U:LYS4, U:PHE85, V:ARG21, V:ARG93, V:HIS88, V:LYS14, V:LYS53, V:PHE56, V:TYR31, W:ARG11, W:ARG14, W:ARG41, W:ARG55, W:ASP15, W:ASP64, W:GLU70, W:GLU85, W:LYS19, W:LYS78, X:ASP60, X:GLU76, X:LYS10, X:TYR78, Y:ARG48, Y:ARG7, Y:ASP49, Y:GLU8, Y:HIS41, Y:LYS60, Y:LYS9, Z:ASP40, Z:LYS19, Z:LYS6 |
