## Supplementary material for "Localized Reactivity on Proteins as Riemannian Manifolds: A Quantum-Inspired Geometric Model for Deterministic, Metal-Aware Reactive-Site Prediction": code bundle: 1TF5test.py

```
#!/usr/bin/env python
"""
SecA (1TF5) residue-level AUC calculation using Ep_sim.npy (Solvent-Excluded Version)

This script reproduces the SecA (PDB: 1TF5) residue-level AUROC values
reported in the paper, using the evaluation protocol described in
Methods 3.2.2 (GroEL-style site-grouped evaluation).

KEY UPGRADE (Solvent-Exclusion Protocol):
  * Automatically filters out solvent residues (HOH, WAT, etc.) based on meta.json.
  * Evaluates performance only on protein residues (Hard Negative Mining).
  * Uses Policy P (Adaptive GroupKFold) to ensure valid splits.

Protocol:
  * Feature matrix X := Ep_sim (with extra geometric features).
  * Labels defined from Tier1_core / Tier2_support residue lists.
  * Site-groups: g_i = (role_i, resno_i) with role_i = "SecA" here.
  * GroupKFold with n_splits <= 5, reduced automatically when needed.
  * RandomForestClassifier, class_weight="balanced", random_state=42.
  * Mean imputation per fold (SimpleImputer).

Assumptions:
- This script is run in the directory containing:
    Ep_manifest.json
    Ep_sim.npy (path stored as manifest["E_sim_path"])
    env_features.json
    meta.json (New Requirement for solvent filtering)
- Ep_manifest["residue_ids"] are keys into env_features.json (e.g. "A:0").
- env_features[rid]["resseq"] is the PDB residue number for SecA (1TF5).
- Labels are defined on chain A; other chains are treated as background.
"""

import json
import numpy as np
from collections import defaultdict

from sklearn.ensemble import RandomForestClassifier
from sklearn.impute import SimpleImputer
from sklearn.metrics import roc_auc_score
from sklearn.model_selection import GroupKFold


# -------------------------------------------------------------------
# 1. Load feature matrix and residue metadata (With Solvent Filtering)
# -------------------------------------------------------------------

MANIFEST_PATH = "Ep_manifest.json"
ENV_FEATURES_PATH = "env_features.json"
META_PATH = "meta.json" # Required for identifying residue types

# Load Data
with open(MANIFEST_PATH, "r") as f:
    manifest = json.load(f)

E_sim = np.load(manifest["E_sim_path"])  # Ep_sim.npy
n_residues_raw, n_features = E_sim.shape
print(f"[INFO] Loaded Raw Ep_sim: {E_sim.shape} (residues × features)")

with open(ENV_FEATURES_PATH, "r") as f:
    env_features = json.load(f)

residue_ids_raw = manifest["residue_ids"]

if len(residue_ids_raw) != n_residues_raw:
    raise ValueError(
        f"Residue count mismatch: len(residue_ids)={len(residue_ids_raw)} "
        f"vs Ep_sim rows={n_residues_raw}"
    )

# =========================================================
# [New] Solvent-Exclusion Filter (Hard Negative Mining)
# =========================================================
print("[INFO] Applying Solvent-Exclusion Filter...")

# 1. Load meta.json to identify residue types
try:
    with open(META_PATH, 'r') as f:
        meta_data = json.load(f)
    
    # Create mapping: ID -> Residue Name
    # Note: Structure depends on meta.json format. Assuming standard list of dicts.
    resname_map = {r['id']: r['resname'] for r in meta_data['residues']}
    
    # 2. Filter indices
    valid_indices = []
    residue_ids = []
    
    # Standard solvent codes to exclude
    solvent_codes = ["HOH", "WAT", "TIP3", "DOD", "SOL"]
    
    removed_count = 0
    
    for i, rid in enumerate(residue_ids_raw):
        # Look up residue name
        rname = resname_map.get(rid, env_features.get(rid, {}).get("resname", "UNKNOWN"))
        
        if rname not in solvent_codes:
            valid_indices.append(i)
            residue_ids.append(rid)
        else:
            removed_count += 1
            
    # 3. Apply Filter to Feature Matrix
    E_sim = E_sim[valid_indices]
    n_residues = len(valid_indices)
    
    print(f"   -> Removed {removed_count} solvent residues.")
    print(f"   -> Retained {n_residues} protein residues for evaluation.")

except FileNotFoundError:
    print("[WARN] meta.json not found! Proceeding WITHOUT solvent filtering.")
    residue_ids = residue_ids_raw
    n_residues = n_residues_raw

print(f"[INFO] Final Ep_sim shape: {E_sim.shape}")
# =========================================================


# Build residue table and mapping (chain, resseq) → indices
# Note: 'indices' now refer to the filtered E_sim rows
rows = []
index_by_chain_resseq = defaultdict(list)

for idx, rid in enumerate(residue_ids):
    info = env_features[rid]
    chain = info["chain"]
    resseq = info["resseq"]  # PDB numbering (1TF5)
    resname = info["resname"]

    row = {
        "idx": idx,
        "id": rid,
        "chain": chain,
        "resseq": resseq,
        "resname": resname,
        "role": "SecA",  # single protein in this case study
    }
    rows.append(row)
    index_by_chain_resseq[(chain, resseq)].append(idx)

rows = sorted(rows, key=lambda r: r["idx"])
print(f"[INFO] Residue table built: {len(rows)} residues")


# -------------------------------------------------------------------
# 2. Define Tier1_core / Tier2_support residue sets (1TF5, chain A)
# -------------------------------------------------------------------

CHAIN_FOR_LABELS = "A"  # evaluation labels are defined for chain A


def expand_ranges(ranges):
    """
    Helper: expand a list of integers and (start, end) tuples into
    a sorted, unique list of residue numbers.
    """
    resnos = []
    for item in ranges:
        if isinstance(item, int):
            resnos.append(item)
        else:
            start, end = item
            resnos.extend(range(start, end + 1))
    return sorted(set(resnos))


# --- ATPase ---

# Tier1_core_ATPase: minimal catalytic core (Walker A/B + catalytic cluster)
TIER1_CORE_ATPASE_RESNOS = {
    106, 107,     # Walker A Lys + neighbour
    207, 208,     # Walker B acidic pair
    215, 216,     # acidic/catalytic cluster just downstream
}

# Tier2_support_ATPase: Walker A / B helical + loop shell around core
ATPASE_SUPPORT_RANGES = [
    (100, 105),   # Walker A upstream loop (core: 106–107)
    (108, 119),   # Walker A helix remainder
    (200, 206),   # Walker B upstream loop
    (208, 214),   # Walker B helix (core 207–208; rest support)
    (217, 220),   # immediate downstream loop
]
_all_atpase_support = expand_ranges(ATPASE_SUPPORT_RANGES)
TIER2_SUPPORT_ATPASE_RESNOS = sorted(
    set(_all_atpase_support) - TIER1_CORE_ATPASE_RESNOS
)


# --- Clamp / preprotein-binding groove ---

# Tier1_core_clamp: hydrophobic grip on two helices
TIER1_CORE_CLAMP_RESNOS = {
    357, 360, 361,   # first helix hydrophobics/aromatics
    381, 384, 385,   # second helix groove-facing hydrophobics
}

# Tier2_support_clamp: same helices + neighbouring loops (minus core)
CLAMP_SUPPORT_RANGES = [
    (352, 356),
    (357, 362),
    (375, 380),
    (377, 386),
    (387, 389),
]
_all_clamp_support = expand_ranges(CLAMP_SUPPORT_RANGES)
TIER2_SUPPORT_CLAMP_RESNOS = sorted(
    set(_all_clamp_support) - TIER1_CORE_CLAMP_RESNOS
)


# --- SecYEG / dimer interface + two-helix finger ---

# Tier1_core_interface: aromatic anchors on interface / finger
TIER1_CORE_INTERFACE_RESNOS = {
    590, 599, 611, 615,   # C-terminal interface helix aromatics
    642,                  # terminal aromatic on C-term helix
    652,                  # finger-loop Trp
    665,                  # downstream Tyr on helix/loop
}

# Tier2_support_interface: same helical/loop bundle around core
INTERFACE_SUPPORT_RANGES = [
    (571, 619),   # long C-terminal helix bundle
    (623, 642),   # next helix (C-term)
    (656, 663),   # following helix
    (643, 655),   # loop around W652
    (664, 674),   # loop around Y665
]
_all_interface_support = expand_ranges(INTERFACE_SUPPORT_RANGES)
TIER2_SUPPORT_INTERFACE_RESNOS = sorted(
    set(_all_interface_support) - TIER1_CORE_INTERFACE_RESNOS
)


# --- Aggregated sets ---

# Tier1_core overall = union of all core sets
TIER1_CORE_ALL_RESNOS = sorted(
    TIER1_CORE_ATPASE_RESNOS
    | TIER1_CORE_CLAMP_RESNOS
    | TIER1_CORE_INTERFACE_RESNOS
)

# Tier2_support overall = union of all support shells
TIER2_SUPPORT_ALL_RESNOS = sorted(
    set(TIER2_SUPPORT_ATPASE_RESNOS)
    | set(TIER2_SUPPORT_CLAMP_RESNOS)
    | set(TIER2_SUPPORT_INTERFACE_RESNOS)
)

# Tier1+Tier2 overall = union of all cores + supports
TIER1_PLUS2_ALL_RESNOS = sorted(
    set(TIER1_CORE_ALL_RESNOS) | set(TIER2_SUPPORT_ALL_RESNOS)
)

print("[INFO] Tier1_core_ATPASE:", sorted(TIER1_CORE_ATPASE_RESNOS))
print("[INFO] Tier2_support_ATPASE:", TIER2_SUPPORT_ATPASE_RESNOS)
print("[INFO] Tier1_core_CLAMP:", sorted(TIER1_CORE_CLAMP_RESNOS))
print("[INFO] Tier2_support_CLAMP:", TIER2_SUPPORT_CLAMP_RESNOS)
print("[INFO] Tier1_core_INTERFACE:", sorted(TIER1_CORE_INTERFACE_RESNOS))
print("[INFO] Tier2_support_INTERFACE:", TIER2_SUPPORT_INTERFACE_RESNOS)
print("[INFO] Tier1_core_all (unique resnos):", TIER1_CORE_ALL_RESNOS)
print("[INFO] Tier1_plus2_all (unique resnos):", TIER1_PLUS2_ALL_RESNOS)


# -------------------------------------------------------------------
# 3. Build site-group labels and generic label vector builder
# -------------------------------------------------------------------

# Site-group labels: g_i = (role_i, resno_i) as strings "SecA:215"
# Note: These are now built from the FILTERED 'rows' list
groups = np.array(
    [f"{r['role']}:{r['resseq']}" for r in rows],
    dtype=object,
)


def build_label_vector_from_resnos(
    resnos,
    chain=CHAIN_FOR_LABELS,
    label_name="",
):
    """
    Given a list/set of PDB residue numbers (resnos) and a chain,
    construct a binary label vector y ∈ {0,1}^N marking positives.

    Any (chain, resno) that does not occur in the manifest is ignored,
    and the missing residue numbers are reported.
    """
    y = np.zeros(len(rows), dtype=int)
    missing = []
    
    # Track found positives to avoid false 'missing' warnings if filtered out intentionally (though core shouldn't be water)
    found_count = 0

    for resno in resnos:
        indices = index_by_chain_resseq.get((chain, resno), [])
        if not indices:
            missing.append(resno)
            continue
        for idx in indices:
            y[idx] = 1
            found_count += 1

    if missing:
        # It's possible some 'support' residues might be missing if the PDB is incomplete,
        # but Core residues should generally be present.
        print(
            f"[WARN] {label_name}: {len(missing)} residue numbers not found on "
            f"chain {chain}: {sorted(set(missing))}"
        )

    n_pos = int(y.sum())
    print(f"[INFO] {label_name}: positives={n_pos}, negatives={len(y) - n_pos}")
    return y


# -------------------------------------------------------------------
# 4. GroupKFold evaluation (Policy P: Adaptive Splits)
# -------------------------------------------------------------------

def evaluate_label_set(
    X,
    y,
    groups,
    label_name,
    random_state=42,
    max_splits=5,
):
    """
    Evaluate a single label set using GroupKFold with site-groups.
    
    POLICY P (Adaptive Splitting):
    - Positive site-groups G+ = { g_i : y_i = 1 }.
    - If |G+| < 2 → no AUC reported (too few positive sites).
    - Otherwise use GroupKFold with n_splits = min(max_splits, |G+|),
      and automatically decrease n_splits until:
         * every test fold contains at least one positive and one negative residue.

    Returns (mean_auc, std_auc) or None if no valid split exists.
    """
    X = np.asarray(X)
    y = np.asarray(y)
    groups = np.asarray(groups)

    pos_groups = set(groups[y == 1])
    n_pos_groups = len(pos_groups)

    if n_pos_groups < 2:
        print(
            f"[SKIP] {label_name}: only {n_pos_groups} positive site group(s); "
            "no AUC reported."
        )
        return None

    # Start with max possible splits given positive groups
    n_splits_start = min(max_splits, n_pos_groups)
    
    print(
        f"[INFO] {label_name}: positive site-groups={n_pos_groups}, "
        f"trying up to {n_splits_start} splits"
    )

    for n_splits in range(n_splits_start, 1, -1):
        gkf = GroupKFold(n_splits=n_splits)
        aucs = []
        valid = True

        for fold_idx, (train_idx, test_idx) in enumerate(
            gkf.split(X, y, groups)
        ):
            y_test = y[test_idx]
            y_train = y[train_idx] # Also check train set

            # Policy P requirement: Both sets must have both classes
            if len(np.unique(y_test)) < 2 or len(np.unique(y_train)) < 2:
                valid = False
                break

            # Standard pipeline: mean imputer + RF (balanced)
            imputer = SimpleImputer(strategy="mean")
            X_train_imp = imputer.fit_transform(X[train_idx])
            X_test_imp = imputer.transform(X[test_idx])

            clf = RandomForestClassifier(
                n_estimators=100,
                class_weight="balanced",
                random_state=random_state,
                n_jobs=1,
            )
            clf.fit(X_train_imp, y_train)
            proba = clf.predict_proba(X_test_imp)[:, 1]

            auc = roc_auc_score(y_test, proba)
            aucs.append(auc)

        if valid and len(aucs) == n_splits:
            mean_auc = float(np.mean(aucs))
            std_auc = float(np.std(aucs, ddof=0))
            print(
                f"[RESULT] {label_name}: n_splits={n_splits}, "
                f"AUC={mean_auc:.4f} ± {std_auc:.4f}"
            )
            return mean_auc, std_auc

        # If invalid, loop continues to try fewer splits (Adaptive K)
        # print(f"   -> n_splits={n_splits} invalid (class imbalance in folds).")

    print(f"[FAIL] {label_name}: no valid GroupKFold split found (even with K=2).")
    return None


# -------------------------------------------------------------------
# 4.b Shuffle test: label–feature leakage check
# -------------------------------------------------------------------

def shuffle_test(X, y, groups, label_name, n_rep=10, random_state=123):
    """
    Run the same evaluation protocol with shuffled labels to obtain
    a baseline AUC distribution.

    - If the implementation is correct and there is no leakage,
      the shuffled-label AUCs should be distributed around ~0.5.
    - If they consistently reach ~0.7–0.8 or higher, there is
      likely some form of label–feature leakage in the setup.
    """
    rng = np.random.RandomState(random_state)
    aucs = []

    print(f"\n[SHUFFLE TEST] {label_name} (n_rep={n_rep})")

    for r in range(n_rep):
        y_shuf = rng.permutation(y)
        res = evaluate_label_set(
            X, y_shuf, groups,
            label_name=f"{label_name}_shuf{r}",
            random_state=random_state + r,
        )
        if res is not None:
            aucs.append(res[0])

    if not aucs:
        print("[SHUFFLE RESULT] no valid split in any repetition.")
        return None

    mean_auc = float(np.mean(aucs))
    std_auc = float(np.std(aucs, ddof=0))
    print(
        f"[SHUFFLE RESULT] {label_name}: "
        f"AUC = {mean_auc:.4f} ± {std_auc:.4f} (over {len(aucs)} reps)"
    )
    return mean_auc, std_auc


# -------------------------------------------------------------------
# 5. Construct label sets and run evaluations
# -------------------------------------------------------------------

# Core-only label vectors
y_tier1_core_atpase = build_label_vector_from_resnos(
    TIER1_CORE_ATPASE_RESNOS,
    chain=CHAIN_FOR_LABELS,
    label_name="Tier1_core_ATPase",
)
y_tier1_core_clamp = build_label_vector_from_resnos(
    TIER1_CORE_CLAMP_RESNOS,
    chain=CHAIN_FOR_LABELS,
    label_name="Tier1_core_clamp",
)
y_tier1_core_interface = build_label_vector_from_resnos(
    TIER1_CORE_INTERFACE_RESNOS,
    chain=CHAIN_FOR_LABELS,
    label_name="Tier1_core_interface",
)
y_tier1_core_all = build_label_vector_from_resnos(
    TIER1_CORE_ALL_RESNOS,
    chain=CHAIN_FOR_LABELS,
    label_name="Tier1_core_all",
)

# Support (Tier2) label vectors
y_tier2_support_atpase = build_label_vector_from_resnos(
    TIER2_SUPPORT_ATPASE_RESNOS,
    chain=CHAIN_FOR_LABELS,
    label_name="Tier2_support_ATPase",
)
y_tier2_support_clamp = build_label_vector_from_resnos(
    TIER2_SUPPORT_CLAMP_RESNOS,
    chain=CHAIN_FOR_LABELS,
    label_name="Tier2_support_clamp",
)
y_tier2_support_interface = build_label_vector_from_resnos(
    TIER2_SUPPORT_INTERFACE_RESNOS,
    chain=CHAIN_FOR_LABELS,
    label_name="Tier2_support_interface",
)
y_tier2_support_all = build_label_vector_from_resnos(
    TIER2_SUPPORT_ALL_RESNOS,
    chain=CHAIN_FOR_LABELS,
    label_name="Tier2_support_all",
)

# Tier1 + Tier2 (per class and all)
y_tier1plus2_atpase = ((y_tier1_core_atpase + y_tier2_support_atpase) > 0).astype(int)
print(
    f"[INFO] Tier1plus2_ATPase: positives={int(y_tier1plus2_atpase.sum())}, "
    f"negatives={len(y_tier1plus2_atpase) - int(y_tier1plus2_atpase.sum())}"
)

y_tier1plus2_clamp = ((y_tier1_core_clamp + y_tier2_support_clamp) > 0).astype(int)
print(
    f"[INFO] Tier1plus2_clamp: positives={int(y_tier1plus2_clamp.sum())}, "
    f"negatives={len(y_tier1plus2_clamp) - int(y_tier1plus2_clamp.sum())}"
)

y_tier1plus2_interface = (
    (y_tier1_core_interface + y_tier2_support_interface) > 0
).astype(int)
print(
    f"[INFO] Tier1plus2_interface: positives={int(y_tier1plus2_interface.sum())}, "
    f"negatives={len(y_tier1plus2_interface) - int(y_tier1plus2_interface.sum())}"
)

y_tier1plus2_all = build_label_vector_from_resnos(
    TIER1_PLUS2_ALL_RESNOS,
    chain=CHAIN_FOR_LABELS,
    label_name="Tier1plus2_all",
)

# Now evaluate each label set
X = E_sim  # feature matrix (Filtered)

results = {}

print("\n=== RUNNING EVALUATIONS ON CLEAN DATA (No Water) ===")

# Core-only
results["Tier1_core_all"] = evaluate_label_set(
    X, y_tier1_core_all, groups, label_name="Tier1_core_all"
)
results["Tier1_core_ATPase"] = evaluate_label_set(
    X, y_tier1_core_atpase, groups, label_name="Tier1_core_ATPase"
)
results["Tier1_core_clamp"] = evaluate_label_set(
    X, y_tier1_core_clamp, groups, label_name="Tier1_core_clamp"
)
results["Tier1_core_interface"] = evaluate_label_set(
    X, y_tier1_core_interface, groups, label_name="Tier1_core_interface"
)

# Tier2-only (optional sanity check)
results["Tier2_support_all"] = evaluate_label_set(
    X, y_tier2_support_all, groups, label_name="Tier2_support_all"
)
results["Tier2_support_ATPase"] = evaluate_label_set(
    X, y_tier2_support_atpase, groups, label_name="Tier2_support_ATPase"
)
results["Tier2_support_clamp"] = evaluate_label_set(
    X, y_tier2_support_clamp, groups, label_name="Tier2_support_clamp"
)
results["Tier2_support_interface"] = evaluate_label_set(
    X, y_tier2_support_interface, groups, label_name="Tier2_support_interface"
)

# Tier1 + Tier2
results["Tier1plus2_all"] = evaluate_label_set(
    X, y_tier1plus2_all, groups, label_name="Tier1plus2_all"
)
results["Tier1plus2_ATPase"] = evaluate_label_set(
    X, y_tier1plus2_atpase, groups, label_name="Tier1plus2_ATPase"
)
results["Tier1plus2_clamp"] = evaluate_label_set(
    X, y_tier1plus2_clamp, groups, label_name="Tier1plus2_clamp"
)
results["Tier1plus2_interface"] = evaluate_label_set(
    X, y_tier1plus2_interface, groups, label_name="Tier1plus2_interface"
)

# -------------------------------------------------------------------
# 6. Shuffle baseline (label–feature leakage sanity check)
# -------------------------------------------------------------------

shuffle_test(X, y_tier1_core_all,     groups, "Tier1_core_all",    n_rep=10)
shuffle_test(X, y_tier1plus2_all,    groups, "Tier1plus2_all",    n_rep=10)
shuffle_test(X, y_tier2_support_all, groups, "Tier2_support_all", n_rep=10)

print("\n=== Summary (No Water / Policy P Applied) ===")
for name, value in results.items():
    if value:
        print(f"{name:24s} -> AUC={value[0]:.4f} ± {value[1]:.4f}")
    else:
        print(f"{name:24s} -> No Result")
```
