## Supplementary material for "Localized Reactivity on Proteins as Riemannian Manifolds: A Quantum-Inspired Geometric Model for Deterministic, Metal-Aware Reactive-Site Prediction": code bundle: 1TSRtest.py

```
#!/usr/bin/env python
"""
p53 (1TSR) + DNA residue-level AUROC evaluation script

Assumptions:
- Ep_sim.npy : (N, d) environment vectors for all residues/nucleotides
- Ep_manifest.json : {"residue_ids": ["A:120", "A:121", ...], ...}
- p53 protein chains = A, B, C
- DNA chains          = E, F

This script:
- loads Ep_sim and Ep_manifest,
- defines Tier-1 / Tier-2 residue sets for p53 (1TSR),
- defines Tier-1 / Tier-2 nucleotide sets for DNA,
- builds site-group labels so that symmetry-related copies
  (e.g. p53 A/B/C with same residue number) are grouped together,
- evaluates AUROC with GroupKFold following the protocol
  used in Methods 3.2.x of the manuscript.

Policy P patch:
- fold validity: BOTH train and test must contain BOTH classes.
- adaptive K: try K = min(K_target, #pos_groups) down to K_min until
  >= min_valid_folds valid folds are obtained; otherwise return NaN.
"""

import json
import numpy as np
from sklearn.model_selection import GroupKFold
from sklearn.ensemble import RandomForestClassifier
from sklearn.impute import SimpleImputer
from sklearn.metrics import roc_auc_score

# ============================
# 0. Paths & chain settings
# ============================

EP_SIM_PATH = "Ep_sim.npy"
MANIFEST_PATH = "Ep_manifest.json"

# For 1TSR; adjust here if needed for a different complex
PROTEIN_CHAINS = ["A", "B", "C"]
DNA_CHAINS = ["E", "F"]

# Policy P knobs (keep fixed across case studies if you want strict policy uniformity)
K_TARGET = 5
K_MIN = 2
MIN_VALID_FOLDS = 2
RF_RANDOM_STATE = 42


# ============================
# 1. Data loading
# ============================

def load_data(ep_sim_path, manifest_path):
    """
    Load environment vectors and residue metadata.

    Returns:
        X        : (N, d) feature matrix
        chain_id : (N,) array of chain IDs (e.g. "A", "E")
        resno    : (N,) array of residue numbers (int)
    """
    X = np.load(ep_sim_path)  # (N, d)

    with open(manifest_path, "r") as f:
        man = json.load(f)
        # =========================================================
# [New] Solvent-Exclusion Filter 
# (Paste this block immediately after loading 'X' and 'manifest')
# =========================================================
import json

# 1. Load meta.json to identify residue types
with open('meta.json', 'r') as f:
    meta_data = json.load(f)

# 2. Create a mapping from residue ID (e.g., "A:100") to residue name (e.g., "HOH")
# Note: Information is located within the 'residues' list in meta.json
resname_map = {r['id']: r['resname'] for r in meta_data['residues']}

# 3. Filter out water/solvent residues (Keep only valid indices)
valid_indices = []
clean_residue_ids = []

# Standard water/solvent codes in PDB
solvent_codes = ["HOH", "WAT", "TIP3", "DOD"]

for i, rid in enumerate(manifest['residue_ids']):
    # Get residue name; default to "UNKNOWN" if not found
    rname = resname_map.get(rid, "UNKNOWN")
    
    # Exclude if it is a known solvent
    if rname not in solvent_codes: 
        valid_indices.append(i)
        clean_residue_ids.append(rid)

# 4. Overwrite data variables with the filtered (clean) subset
X = X[valid_indices]
manifest['residue_ids'] = clean_residue_ids

print(f"-> [Filter Applied] Retained Residues: {len(valid_indices)} (Solvents Removed)")
# =========================================================

    # We assume residue_ids are strings of the form "A:120", "A:121", ...
    residue_ids = man["residue_ids"]
    chain_id = np.array([rid.split(":")[0] for rid in residue_ids], dtype=object)
    resno = np.array([int(rid.split(":")[1]) for rid in residue_ids], dtype=int)

    return X, chain_id, resno


# ============================
# 2. GroupKFold-based AUROC (Policy P)
# ============================

def eval_auc_grouped(
    X,
    y,
    groups,
    n_splits=K_TARGET,
    random_state=RF_RANDOM_STATE,
    k_min=K_MIN,
    min_valid_folds=MIN_VALID_FOLDS,
):
    """
    Evaluate AUROC using GroupKFold with site-level grouping.

    Policy P:
    - A fold is valid only if BOTH train and test contain BOTH classes.
    - Try K = min(n_splits, #positive_groups) down to k_min until
      >= min_valid_folds valid folds are obtained.
    - If still insufficient, return (nan, nan, [], info).

    Args:
        X       : (N_sub, d) feature matrix
        y       : (N_sub,)  0/1 labels
        groups  : (N_sub,)  group IDs, e.g. "P53_120" or "DNA_E_12"
        n_splits: target maximum number of splits to try
        random_state: RF random_state
        k_min: minimum K to try
        min_valid_folds: minimum number of valid folds required to accept a run

    Returns:
        (mean_auc, std_auc, auc_list, info_dict)
        If no valid split is found, returns (nan, nan, [], info_dict).
    """
    y = np.asarray(y).astype(int)
    groups = np.asarray(groups)

    # positive site-groups are the bottleneck for grouped CV
    pos_groups = np.unique(groups[y == 1])
    n_pos_groups = int(len(pos_groups))

    info = {
        "K_target": int(n_splits),
        "K_min": int(k_min),
        "min_valid_folds": int(min_valid_folds),
        "n_pos_groups": int(n_pos_groups),
        "K_used": None,
        "n_total_folds": 0,
        "n_valid_folds": 0,
        "n_skipped_train_single_class": 0,
        "n_skipped_test_single_class": 0,
        "reason": "",
    }

    if n_pos_groups < 2:
        info["reason"] = "Too few positive site-groups (<2) for grouped split."
        return np.nan, np.nan, [], info

    K_start = min(int(n_splits), n_pos_groups)
    if K_start < 2:
        info["reason"] = "K_start < 2 (insufficient positive site-groups)."
        return np.nan, np.nan, [], info

    # Try decreasing K until we get enough valid folds
    for K in range(K_start, int(k_min) - 1, -1):
        gkf = GroupKFold(n_splits=K)
        aucs = []
        total_folds = 0
        skipped_train = 0
        skipped_test = 0

        for train_idx, test_idx in gkf.split(X, y, groups):
            total_folds += 1
            y_train = y[train_idx]
            y_test = y[test_idx]

            # fold validity: both classes must exist in train and test
            if np.unique(y_train).size < 2:
                skipped_train += 1
                continue
            if np.unique(y_test).size < 2:
                skipped_test += 1
                continue

            imputer = SimpleImputer(strategy="mean")
            X_train_imp = imputer.fit_transform(X[train_idx])
            X_test_imp = imputer.transform(X[test_idx])

            clf = RandomForestClassifier(
                n_estimators=100,
                class_weight="balanced",
                random_state=random_state,
                n_jobs=1,
            )
            clf.fit(X_train_imp, y_train)
            y_prob = clf.predict_proba(X_test_imp)[:, 1]

            aucs.append(roc_auc_score(y_test, y_prob))

        info["K_used"] = int(K)
        info["n_total_folds"] = int(total_folds)
        info["n_valid_folds"] = int(len(aucs))
        info["n_skipped_train_single_class"] = int(skipped_train)
        info["n_skipped_test_single_class"] = int(skipped_test)

        if len(aucs) >= int(min_valid_folds):
            info["reason"] = "OK"
            return float(np.mean(aucs)), float(np.std(aucs)), aucs, info

        # else: try smaller K

    info["reason"] = "Insufficient valid folds even after decreasing K."
    return np.nan, np.nan, [], info


def union_lists(*lists):
    """
    Take the union of multiple integer lists and return a sorted list.
    """
    s = set()
    for L in lists:
        s.update(L)
    return sorted(s)


# ============================
# 3. p53 / DNA label definitions
# ============================

# --- p53: Tier sets (1TSR, residue-number level) ---

p53_tiers = {
    "Tier1a_DNA_contact_core": [120, 241, 248, 273, 276, 277, 280, 283],
    "Tier1b_Zn_structural_cluster": [171, 176, 179, 238, 242],
    "Tier1c_fitness_stability_core": [175, 220, 245, 249, 282],
    # Extended PPI / allosteric pivots discussed in the manuscript
    "Tier1d_allosteric_PPI_pivots": [
        100, 104, 107,           # N-term loop / β1 (BCL-xL interface, dimer interface)
        178, 180, 181,           # L2-adjacent positions (cooperativity / ASPP2-related)
        207, 208, 209, 210, 211, 212, 213,  # allosteric loop
        243, 247,                # L3 PPI (ASPP2 contact)
        268,                     # C-side PPI contact (BCL-xL)
    ],
    "Tier2_structural_shell": [109, 119, 121, 145, 146, 246, 281, 285],
}

# --- DNA: nucleotide tiers (chain E/F, 1TSR) ---

DNA_tiers = {
    # Nucleotides jointly contacted by multiple p53 core residues
    "Tier1_DNA_contact_core": [
        ("E", 12),   # DT: contacted by R248 / R273 / R280
        ("F", 7),    # DG: K120 / R283
        ("F", 9),    # DG: K120 / R280
    ],
    # Nucleotides that contact exactly one of the core residues (shell)
    "Tier2_DNA_contact_shell": [
        ("E", 1), ("E", 11), ("E", 13), ("E", 14),
        ("F", 6), ("F", 8), ("F", 14), ("F", 16),
    ],
}


# ============================
# 4. Mask construction
# ============================

def make_mask_p53(resno_sub, pos_resnos):
    """
    Build a boolean mask for p53 residues within a subset.

    Args:
        resno_sub  : (N_sub,) residue numbers in the subset
        pos_resnos : list of residue numbers defining positives

    Returns:
        mask : (N_sub,) bool, True for positives
    """
    pos_set = set(pos_resnos)
    return np.array([r in pos_set for r in resno_sub], dtype=bool)


def make_mask_DNA(chain_sub, resno_sub, pos_pairs):
    """
    Build a boolean mask for DNA nucleotides within a subset.

    Args:
        chain_sub : (N_sub,) chain IDs (e.g. "E", "F")
        resno_sub : (N_sub,) nucleotide numbers (int)
        pos_pairs : list of (chain, resno) tuples that are positive

    Returns:
        mask : (N_sub,) bool, True for positives
    """
    mask = np.zeros_like(resno_sub, dtype=bool)
    pair_set = {(c, int(r)) for (c, r) in pos_pairs}
    for i, (c, r) in enumerate(zip(chain_sub, resno_sub)):
        if (c, int(r)) in pair_set:
            mask[i] = True
    return mask


# ============================
# 5. p53 / DNA evaluation helpers
# ============================

def eval_p53_tier(X, chain_id, resno, pos_resnos, name, n_splits=K_TARGET):
    """
    Evaluate AUROC for a given p53 Tier set.

    - Restricts to protein chains (A/B/C).
    - Groups residues by residue number (resno) across symmetry-related chains,
      so A:120, B:120, C:120 are treated as one site.

    Args:
        X         : (N, d) full feature matrix
        chain_id  : (N,) chain IDs
        resno     : (N,) residue numbers
        pos_resnos: list of p53 residue numbers that are positive
        name      : label set name (for printing)
        n_splits  : maximum GroupKFold splits
    """
    subset_mask = np.isin(chain_id, PROTEIN_CHAINS)
    idx = np.where(subset_mask)[0]

    X_sub = X[idx]
    resno_sub = resno[idx]

    # Site-groups: group by residue number only; A/B/C copies collapse to one site
    groups = np.array([f"P53_{r}" for r in resno_sub], dtype=object)

    y = make_mask_p53(resno_sub, pos_resnos).astype(int)
    n_pos = int(y.sum())
    n_total = int(len(y))

    auc_mean, auc_std, aucs, info = eval_auc_grouped(
        X_sub, y, groups,
        n_splits=n_splits,
        random_state=RF_RANDOM_STATE,
        k_min=K_MIN,
        min_valid_folds=MIN_VALID_FOLDS,
    )

    print(f"[p53 {name}] N={n_total}  #positives={n_pos}  #negatives={n_total - n_pos}")
    print(f"[p53 {name}] AUC = {auc_mean} +/- {auc_std}")
    print(f"[p53 {name}] CV info: {info}")
    print(f"[p53 {name}] fold AUCs: {aucs}\n")


def eval_DNA_tier(X, chain_id, resno, pos_pairs, name, n_splits=K_TARGET):
    """
    Evaluate AUROC for a given DNA Tier set.

    - Restricts to DNA chains (E/F).
    - Groups nucleotides by (chain, resno).

    Args:
        X        : (N, d) full feature matrix
        chain_id : (N,) chain IDs
        resno    : (N,) nucleotide numbers
        pos_pairs: list of (chain, resno) tuples that are positive
        name     : label set name (for printing)
        n_splits : maximum GroupKFold splits
    """
    subset_mask = np.isin(chain_id, DNA_CHAINS)
    idx = np.where(subset_mask)[0]

    X_sub = X[idx]
    resno_sub = resno[idx]
    chain_sub = chain_id[idx]

    # For DNA, treat each nucleotide (chain, resno) as its own site
    groups = np.array([f"DNA_{c}_{r}" for c, r in zip(chain_sub, resno_sub)], dtype=object)

    y = make_mask_DNA(chain_sub, resno_sub, pos_pairs).astype(int)
    n_pos = int(y.sum())
    n_total = int(len(y))

    auc_mean, auc_std, aucs, info = eval_auc_grouped(
        X_sub, y, groups,
        n_splits=n_splits,
        random_state=RF_RANDOM_STATE,
        k_min=K_MIN,
        min_valid_folds=MIN_VALID_FOLDS,
    )

    print(f"[DNA {name}] N={n_total}  #positives={n_pos}  #negatives={n_total - n_pos}")
    print(f"[DNA {name}] AUC = {auc_mean} +/- {auc_std}")
    print(f"[DNA {name}] CV info: {info}")
    print(f"[DNA {name}] fold AUCs: {aucs}\n")


# ============================
# 6. Main: run all evaluations
# ============================

if __name__ == "__main__":
    X, chain_id, resno = load_data(EP_SIM_PATH, MANIFEST_PATH)

    N, d = X.shape
    print(f"[INFO] N (residues/nucleotides): {N}")
    print(f"[INFO] X shape: {X.shape}")
    print(f"[INFO] protein chains: {PROTEIN_CHAINS}, DNA chains: {DNA_CHAINS}\n")

    # ----- p53 Tier sets (residue-number level) -----
    t1a = p53_tiers["Tier1a_DNA_contact_core"]
    t1b = p53_tiers["Tier1b_Zn_structural_cluster"]
    t1c = p53_tiers["Tier1c_fitness_stability_core"]
    t1d = p53_tiers["Tier1d_allosteric_PPI_pivots"]
    t2  = p53_tiers["Tier2_structural_shell"]

    tier1_all = union_lists(t1a, t1b, t1c, t1d)
    tier1_plus_2 = union_lists(tier1_all, t2)

    print("[p53 Tier sets (resno-level)]")
    print("  Tier1a:", t1a)
    print("  Tier1b:", t1b)
    print("  Tier1c:", t1c)
    print("  Tier1d:", t1d)
    print("  Tier2 :", t2)
    print("  Tier1(all):", tier1_all)
    print("  Tier1+Tier2:", tier1_plus_2)
    print()

    # ----- p53 AUROC evaluations -----
    eval_p53_tier(X, chain_id, resno, tier1_all,     "Global_Tier1", n_splits=K_TARGET)
    eval_p53_tier(X, chain_id, resno, tier1_plus_2,  "Global_Tier1_plus_Tier2", n_splits=K_TARGET)

    eval_p53_tier(X, chain_id, resno, t1a, "Tier1a_DNA_contact_core", n_splits=K_TARGET)
    eval_p53_tier(X, chain_id, resno, t1b, "Tier1b_Zn_structural_cluster", n_splits=K_TARGET)
    eval_p53_tier(X, chain_id, resno, t1c, "Tier1c_fitness_stability_core", n_splits=K_TARGET)
    eval_p53_tier(X, chain_id, resno, t1d, "Tier1d_allosteric_PPI_pivots", n_splits=K_TARGET)
    eval_p53_tier(X, chain_id, resno, t2,  "Tier2_structural_shell", n_splits=K_TARGET)

    # ----- DNA Tier sets -----
    dna_t1 = DNA_tiers["Tier1_DNA_contact_core"]
    dna_t2 = DNA_tiers["Tier2_DNA_contact_shell"]
    dna_t1_plus_2 = dna_t1 + dna_t2

    print("[DNA Tier sets]")
    print("  Tier1_DNA_contact_core:", dna_t1)
    print("  Tier2_DNA_contact_shell:", dna_t2)
    print("  Tier1+Tier2:", dna_t1_plus_2)
    print()

    # ----- DNA AUROC evaluations -----
    eval_DNA_tier(X, chain_id, resno, dna_t1,        "Tier1_DNA_contact_core", n_splits=K_TARGET)
    eval_DNA_tier(X, chain_id, resno, dna_t2,        "Tier2_DNA_contact_shell", n_splits=K_TARGET)
    eval_DNA_tier(X, chain_id, resno, dna_t1_plus_2, "Tier1_plus_Tier2_DNA", n_splits=K_TARGET)
```
