## Supplementary material for "Localized Reactivity on Proteins as Riemannian Manifolds: A Quantum-Inspired Geometric Model for Deterministic, Metal-Aware Reactive-Site Prediction": code bundle: 2c7ctest.py

```
#!/usr/bin/env python
import json
import numpy as np
from sklearn.impute import SimpleImputer
from sklearn.ensemble import RandomForestClassifier
from sklearn.model_selection import GroupKFold
from sklearn.metrics import roc_auc_score

# ============================
# 0. File paths
# ============================
EP_SIM_PATH = "Ep_sim.npy"          # Ep_sim.npy file
MANIFEST_PATH = "Ep_manifest.json"  # Ep_manifest.json file

# ============================
# Policy P (fixed)
# ============================
K_TARGET = 5
K_MIN = 2
MIN_VALID_FOLDS = 2
RF_RANDOM_STATE = 42


# ============================
# 1. Data loading
#   - X: (N, d) feature matrix
#   - roles: "EL" or "ES" (GroEL vs GroES)
#   - resnos: residue numbers (int)
#   - groups: GroupKFold groups (role:resno)
#   - residue_ids: strings like "A:34"
# ============================
def load_data(ep_sim_path=EP_SIM_PATH, manifest_path=MANIFEST_PATH):
    # Load environment vectors
    X = np.load(ep_sim_path)

    # Load manifest (we only need residue_ids here)
    with open(manifest_path, "r") as f:
        manifest = json.load(f)

    residue_ids = manifest["residue_ids"]

    N = len(residue_ids)
    if X.shape[0] != N:
        raise ValueError(f"X has {X.shape[0]} rows but manifest lists {N} residues.")

    chains = []
    resnos = []
    for rid in residue_ids:
        chain, res = rid.split(":")
        chains.append(chain)
        resnos.append(int(res))

    chains = np.array(chains, dtype=object)
    resnos = np.array(resnos, dtype=int)

    # Infer GroEL vs GroES role per chain:
    #  - GroEL chains have length ≥ 400
    #  - GroES chains have length ~100
    chain_to_max = {}
    for c, r in zip(chains, resnos):
        chain_to_max[c] = max(chain_to_max.get(c, 0), r)

    roles = []
    for c in chains:
        max_res = chain_to_max[c]
        role = "EL" if max_res >= 400 else "ES"
        roles.append(role)
    roles = np.array(roles, dtype=object)

    # GroupKFold groups: (role, resno) as strings "EL:30" / "ES:27"
    groups = np.array([f"{role}:{res}" for role, res in zip(roles, resnos)], dtype=object)

    return X, roles, resnos, groups, residue_ids


# ============================
# 2. Tier1 class definitions
# ============================
GROEL_TIER1_CLASSES = {
    "groel_A_ATP_core": [30, 31, 32, 33, 34, 51, 52, 87, 89, 398],
    "groel_B_substrate_patch": [199, 201, 203, 204, 234, 237, 263, 264],
    "groel_C_hinge_core": [192],
    "groel_C_hinge_support": [374, 375],
    "groel_D_inter_ring": [105, 197, 386, 434, 452, 461],
}

GROES_TIER1_CLASSES = {
    "groes_E_IVL_core": [25, 26, 27],
    "groes_F_loop_support": [23, 24, 28, 29, 30, 31],
}


# ============================
# 3. Label mask construction
# ============================
def make_label_mask(roles, resnos, class_resnos, role_filter=None):
    """
    Build a boolean mask for a given Tier1 class.

    roles       : (N,)  "EL"/"ES"
    resnos      : (N,)  int residue numbers
    class_resnos: list of residue numbers [30, 31, ...]
    role_filter : "EL" / "ES" / None
    """
    mask = np.isin(resnos, class_resnos)
    if role_filter is not None:
        mask &= (roles == role_filter)
    return mask


# ============================
# 4. Policy P: GroupKFold-based AUROC (adaptive K + fold validity)
# ============================
def _fit_predict_auc_rf(X_train, y_train, X_test, y_test, random_state=RF_RANDOM_STATE):
    imputer = SimpleImputer(strategy="mean")
    X_train_imp = imputer.fit_transform(X_train)
    X_test_imp = imputer.transform(X_test)

    clf = RandomForestClassifier(
        n_estimators=100,
        class_weight="balanced",
        random_state=random_state,
        n_jobs=1,
    )
    clf.fit(X_train_imp, y_train)
    proba = clf.predict_proba(X_test_imp)[:, 1]
    return roc_auc_score(y_test, proba)


def eval_auc_policyP(X, y, groups, desc="",
                     K_target=K_TARGET, K_min=K_MIN, min_valid_folds=MIN_VALID_FOLDS,
                     random_state=RF_RANDOM_STATE):
    """
    Policy P:
      - split validity rule: train/test must both have {0,1}
      - adaptive K: try K from min(K_target, #positive-groups) down to K_min
      - accept first K with >= min_valid_folds valid folds
      - return (mean, std, fold_aucs, info_dict)
    """
    y = np.asarray(y, dtype=int)
    groups = np.asarray(groups, dtype=object)

    n_pos = int((y == 1).sum())
    n_neg = int((y == 0).sum())

    pos_groups = np.unique(groups[y == 1])
    n_pos_groups = int(len(pos_groups))

    info = {
        "desc": str(desc),
        "policy": "P",
        "cv": "GroupKFold",
        "K_target": int(K_target),
        "K_min": int(K_min),
        "min_valid_folds": int(min_valid_folds),
        "n_pos": int(n_pos),
        "n_neg": int(n_neg),
        "n_pos_groups": int(n_pos_groups),
        "K_used": None,
        "n_total_folds": 0,
        "n_valid_folds": 0,
        "n_skipped_train_single_class": 0,
        "n_skipped_test_single_class": 0,
        "reason": "",
    }

    # Need at least 2 positive site-groups for grouped CV to make sense
    if n_pos_groups < 2:
        info["reason"] = "Too few positive groups (<2); grouped CV not feasible."
        return np.nan, np.nan, [], info

    K_start = min(int(K_target), n_pos_groups)
    if K_start < 2:
        info["reason"] = "K_start < 2."
        return np.nan, np.nan, [], info

    for K in range(K_start, int(K_min) - 1, -1):
        gkf = GroupKFold(n_splits=K)
        aucs = []
        total_folds = 0
        skipped_train = 0
        skipped_test = 0

        for train_idx, test_idx in gkf.split(X, y, groups):
            total_folds += 1
            y_tr = y[train_idx]
            y_te = y[test_idx]

            if np.unique(y_tr).size < 2:
                skipped_train += 1
                continue
            if np.unique(y_te).size < 2:
                skipped_test += 1
                continue

            auc = _fit_predict_auc_rf(X[train_idx], y_tr, X[test_idx], y_te, random_state=random_state)
            aucs.append(float(auc))

        info["K_used"] = int(K)
        info["n_total_folds"] = int(total_folds)
        info["n_valid_folds"] = int(len(aucs))
        info["n_skipped_train_single_class"] = int(skipped_train)
        info["n_skipped_test_single_class"] = int(skipped_test)

        if len(aucs) >= int(min_valid_folds):
            info["reason"] = "OK"
            return float(np.mean(aucs)), float(np.std(aucs)), aucs, info

    info["reason"] = "Insufficient valid folds after adaptive K."
    return np.nan, np.nan, [], info


# ============================
# 5. Main: compute all resolutions
# ============================
def main():
    X, roles, resnos, groups, residue_ids = load_data()
    N = X.shape[0]
    print(f"[INFO] N residues: {N}")
    print(f"[INFO] X shape   : {X.shape}")

    # --------------------------------
    # (1) Global Tier1 (all EL + ES)
    # --------------------------------
    y_global = np.zeros(N, dtype=int)
    all_classes = {}
    all_classes.update(GROEL_TIER1_CLASSES)
    all_classes.update(GROES_TIER1_CLASSES)

    for name, res_list in all_classes.items():
        role = "EL" if name.startswith("groel_") else "ES"
        y_global |= make_label_mask(roles, resnos, res_list, role_filter=role).astype(int)

    npos = int(y_global.sum())
    nneg = int(N - npos)
    print(f"\n[Global Tier1] N={N}  #positives={npos}  #negatives={nneg}")

    auc, sd, folds, info = eval_auc_policyP(X, y_global, groups, desc="Global Tier1")
    print(f"[Global Tier1] AUC = {auc} +/- {sd}")
    print(f"[Global Tier1] CV info: {info}")
    print(f"[Global Tier1] fold AUCs: {folds}")

    # --------------------------------
    # (2) GroEL (EL) Tier1 overall
    # --------------------------------
    y_el = np.zeros(N, dtype=int)
    for name, res_list in GROEL_TIER1_CLASSES.items():
        y_el |= make_label_mask(roles, resnos, res_list, role_filter="EL").astype(int)

    mask_el = (roles == "EL")
    N_el = int(mask_el.sum())
    npos_el = int(y_el[mask_el].sum())
    nneg_el = int(N_el - npos_el)

    print(f"\n[GroEL (EL) Tier1] N={N_el}  #positives={npos_el}  #negatives={nneg_el}")
    auc_el, sd_el, folds_el, info_el = eval_auc_policyP(
        X[mask_el], y_el[mask_el], groups[mask_el], desc="GroEL Tier1 (EL-only)"
    )
    print(f"[GroEL (EL) Tier1] AUC = {auc_el} +/- {sd_el}")
    print(f"[GroEL (EL) Tier1] CV info: {info_el}")
    print(f"[GroEL (EL) Tier1] fold AUCs: {folds_el}")

    # --------------------------------
    # (3) GroES (ES) Tier1 overall
    # --------------------------------
    y_es = np.zeros(N, dtype=int)
    for name, res_list in GROES_TIER1_CLASSES.items():
        y_es |= make_label_mask(roles, resnos, res_list, role_filter="ES").astype(int)

    mask_es = (roles == "ES")
    N_es = int(mask_es.sum())
    npos_es = int(y_es[mask_es].sum())
    nneg_es = int(N_es - npos_es)

    print(f"\n[GroES (ES) Tier1] N={N_es}  #positives={npos_es}  #negatives={nneg_es}")
    auc_es, sd_es, folds_es, info_es = eval_auc_policyP(
        X[mask_es], y_es[mask_es], groups[mask_es], desc="GroES Tier1 (ES-only)"
    )
    print(f"[GroES (ES) Tier1] AUC = {auc_es} +/- {sd_es}")
    print(f"[GroES (ES) Tier1] CV info: {info_es}")
    print(f"[GroES (ES) Tier1] fold AUCs: {folds_es}")

    # --------------------------------
    # (4) Per-class AUC (GroEL / EL)
    # --------------------------------
    print("\n[Per-class AUC: GroEL (EL)]")
    for name, res_list in GROEL_TIER1_CLASSES.items():
        y = make_label_mask(roles, resnos, res_list, role_filter="EL").astype(int)
        npos_c = int(y[mask_el].sum())
        nneg_c = int(N_el - npos_c)

        auc_c, sd_c, folds_c, info_c = eval_auc_policyP(
            X[mask_el], y[mask_el], groups[mask_el], desc=name
        )
        print(f"  [{name}] N={N_el}  #positives={npos_c}  #negatives={nneg_c}")
        print(f"  [{name}] AUC = {auc_c} +/- {sd_c}")
        print(f"  [{name}] CV info: {info_c}")
        print(f"  [{name}] fold AUCs: {folds_c}\n")

    # --------------------------------
    # (5) Per-class AUC (GroES / ES)
    # --------------------------------
    print("\n[Per-class AUC: GroES (ES)]")
    for name, res_list in GROES_TIER1_CLASSES.items():
        y = make_label_mask(roles, resnos, res_list, role_filter="ES").astype(int)
        npos_c = int(y[mask_es].sum())
        nneg_c = int(N_es - npos_c)

        auc_c, sd_c, folds_c, info_c = eval_auc_policyP(
            X[mask_es], y[mask_es], groups[mask_es], desc=name
        )
        print(f"  [{name}] N={N_es}  #positives={npos_c}  #negatives={nneg_c}")
        print(f"  [{name}] AUC = {auc_c} +/- {sd_c}")
        print(f"  [{name}] CV info: {info_c}")
        print(f"  [{name}] fold AUCs: {folds_c}\n")


if __name__ == "__main__":
    main()
```
