## Supplementary material for "Localized Reactivity on Proteins as Riemannian Manifolds: A Quantum-Inspired Geometric Model for Deterministic, Metal-Aware Reactive-Site Prediction": code bundle: 3VFGtest.py

```
import json
import numpy as np
import sys
from scipy.spatial.distance import pdist, squareform
from sklearn.ensemble import RandomForestClassifier
from sklearn.impute import SimpleImputer
from sklearn.model_selection import StratifiedKFold
from sklearn.metrics import roc_auc_score, average_precision_score

# =============================================================================
# CONFIGURATION & CONSTANTS
# =============================================================================
MANIFEST_PATH = 'Ep_manifest.json'
FEATURE_PATH = 'Ep_sim.npy'
PDB_PATH = '3VFG.pdb'
RANDOM_STATE = 42

# Definition of CDR Loops (Ground Truth)
# Based on Chothia/Kabat merged definitions for 3VFG
def get_label(rid):
    try:
        chain, num_str = rid.split(':')
        # Handle insertion codes like "100A" -> 100
        num_str_clean = "".join(filter(str.isdigit, num_str))
        if not num_str_clean: return 0
        num = int(num_str_clean)
        
        if chain == 'L':
            if 24 <= num <= 34: return 1  # CDR-L1
            if 50 <= num <= 56: return 1  # CDR-L2
            if 89 <= num <= 97: return 1  # CDR-L3
        elif chain == 'H':
            if 26 <= num <= 35: return 1  # CDR-H1
            if 50 <= num <= 65: return 1  # CDR-H2
            if 95 <= num <= 102: return 1 # CDR-H3
        return 0
    except:
        return 0

# =============================================================================
# CORE LOGIC: POLICY P (Adaptive Cross-Validation)
# This function replicates the robust logic from '6Q97test.py'
# =============================================================================
def cross_validated_metrics_policy_p(X, y, K_target=5, min_valid_folds=2):
    """
    Executes Robust Cross-Validation (Policy P).
    
    Features:
    - Handles Class Imbalance (class_weight='balanced')
    - Handles Missing Values (SimpleImputer)
    - Adaptive K: Reduces K if the dataset is too small.
    - Robustness: Skips folds with only one class in train/test.
    
    Returns:
        dict: {'roc_auc': float, 'pr_auc': float, 'k_used': int}
    """
    
    # 1. Impute missing values (Safety mechanism from Ribosome code)
    imputer = SimpleImputer(strategy='mean')
    X_imputed = imputer.fit_transform(X)
    
    # 2. Adaptive K loop
    # Try K_target, if it fails, reduce K down to 2.
    final_roc_scores = []
    final_pr_scores = []
    k_used = K_target
    
    for k in range(K_target, 1, -1):
        kf = StratifiedKFold(n_splits=k, shuffle=True, random_state=RANDOM_STATE)
        fold_roc_scores = []
        fold_pr_scores = []
        valid_folds = 0
        
        try:
            for train_idx, test_idx in kf.split(X_imputed, y):
                X_train, X_test = X_imputed[train_idx], X_imputed[test_idx]
                y_train, y_test = y[train_idx], y[test_idx]
                
                # Skip if only one class is present (Edge case safety)
                if len(np.unique(y_train)) < 2 or len(np.unique(y_test)) < 2:
                    continue
                
                # Initialize Model with Balanced Weights (Crucial for Ribosome/Antibody logic)
                clf = RandomForestClassifier(
                    n_estimators=100,
                    class_weight='balanced',  # <--- Key difference from simple code
                    n_jobs=-1,
                    random_state=RANDOM_STATE
                )
                clf.fit(X_train, y_train)
                
                # Predict
                if hasattr(clf, "predict_proba"):
                    y_pred = clf.predict_proba(X_test)[:, 1]
                else:
                    y_pred = clf.predict(X_test)
                
                # Calculate Metrics
                roc = roc_auc_score(y_test, y_pred)
                pr = average_precision_score(y_test, y_pred)
                
                fold_roc_scores.append(roc)
                fold_pr_scores.append(pr)
                valid_folds += 1
            
            # Check if we have enough valid folds
            if valid_folds >= min_valid_folds:
                final_roc_scores = fold_roc_scores
                final_pr_scores = fold_pr_scores
                k_used = k
                break # Success, exit adaptive loop
            else:
                # Not enough valid folds, try smaller K
                continue
                
        except Exception as e:
            # If splitting fails (e.g., n_samples < n_splits), try smaller K
            continue

    # 3. Aggregate Results
    if not final_roc_scores:
        return {'roc_auc': 0.5, 'pr_auc': 0.0, 'k_used': 0} # Failure fallback
        
    return {
        'roc_auc': np.mean(final_roc_scores),
        'pr_auc': np.mean(final_pr_scores),
        'k_used': k_used
    }

# =============================================================================
# HELPER: GEOMETRY & SURFACE DEFINITION
# =============================================================================
def parse_pdb_and_get_surface_mask(pdb_file, manifest_rids, threshold=18):
    """
    Parses PDB to find neighbors. Returns a boolean mask for surface residues.
    Dependency-free (No Biopython required).
    """
    coords = {}
    with open(pdb_file, 'r') as f:
        for line in f:
            if line.startswith("ATOM") and line[12:16].strip() == "CA":
                chain = line[21]
                res_num = int(line[22:26])
                x, y, z = float(line[30:38]), float(line[38:46]), float(line[46:54])
                coords[f"{chain}:{res_num}"] = np.array([x, y, z])

    valid_indices = []
    positions = []
    
    # Align Manifest with PDB Coords
    for i, rid in enumerate(manifest_rids):
        if rid in coords:
            valid_indices.append(i)
            positions.append(coords[rid])
            
    if not positions:
        return [], []

    positions = np.array(positions)
    
    # Calculate Neighbor Counts (10 Angstrom radius)
    dists = squareform(pdist(positions))
    neighbor_counts = (dists < 10.0).sum(axis=1) - 1
    
    # Define Surface: Low neighbor count
    surface_mask = neighbor_counts < threshold
    
    return valid_indices, surface_mask

# =============================================================================
# MAIN EXECUTION
# =============================================================================
if __name__ == "__main__":
    print(f"--- Starting Robust Evaluation (Policy P) ---")
    print(f"Loading data from {MANIFEST_PATH}...")
    
    # 1. Load Data
    with open(MANIFEST_PATH, 'r') as f:
        manifest = json.load(f)
    ep_data = np.load(FEATURE_PATH)
    residue_ids = manifest['residue_ids']
    
    # 2. Generate Ground Truth Labels
    y_all = np.array([get_label(rid) for rid in residue_ids])
    
    # 3. Get Surface Mask (Geometric Filtering)
    valid_indices, surface_mask = parse_pdb_and_get_surface_mask(PDB_PATH, residue_ids)
    
    # Subset valid data (intersection of Manifest and PDB)
    X_valid = ep_data[valid_indices]
    y_valid = y_all[valid_indices]
    
    # ==========================================
    # EXPERIMENT 1: Full Protein Evaluation
    # ==========================================
    print(f"\n[Experiment 1] Full Protein (All Residues)")
    print(f"Total Samples: {len(y_valid)}")
    print(f"Positives (CDRs): {sum(y_valid)}")
    
    res_full = cross_validated_metrics_policy_p(X_valid, y_valid)
    print(f"-> ROC-AUC: {res_full['roc_auc']:.4f}")
    print(f"-> PR-AUC : {res_full['pr_auc']:.4f}")
    print(f"-> K-Folds: {res_full['k_used']}")

    # ==========================================
    # EXPERIMENT 2: Surface Only Evaluation
    # ==========================================
    print(f"\n[Experiment 2] Surface Only (Neighbor Count < 18)")
    X_surf = X_valid[surface_mask]
    y_surf = y_valid[surface_mask]
    
    print(f"Total Samples: {len(y_surf)}")
    print(f"Positives (CDRs): {sum(y_surf)}")
    
    res_surf = cross_validated_metrics_policy_p(X_surf, y_surf)
    print(f"-> ROC-AUC: {res_surf['roc_auc']:.4f}")
    print(f"-> PR-AUC : {res_surf['pr_auc']:.4f}")
    print(f"-> K-Folds: {res_surf['k_used']}")

    print("\n--- Evaluation Complete ---")
    print("Note: 'class_weight=balanced' was used, making PR-AUC highly reliable.")
```
