## Supplementary material for "Localized Reactivity on Proteins as Riemannian Manifolds: A Quantum-Inspired Geometric Model for Deterministic, Metal-Aware Reactive-Site Prediction": code bundle: 4.7.py

```
#!/usr/bin/env python
"""
protein_env_full_pipeline_gpu.py

Full pipeline (block-streaming version) with optional GPU acceleration
and explicit tracking of effective GPU density modes per residue.

Steps
-----
1–2 : load structure, build residue list, KDTree-based contacts
      -> meta.json (residues + overlaps + pair_contacts + SASA)
3–4 : hybrid density:
        - B: promolecular surrogate density for all residues
        - A: optional xTB-based correction for a chosen subset of residues
5–6 : per-residue features (env_features.json):
        - density-based: total_density, centroid, dipole_rho, curv_rho
        - charge-based:
            * B-mode: surrogate partial_charge_B (atom-based + SASA scaling)
            * A-mode: partial_charge, dipole_q (if xTB run)
        - SASA
        - field terms: F_metal, F_P, F_hal
7–9 : environment vectors Ep:
        - Ep_theory.npy : [avg_charge, dipole_norm, curv_density,
                          external_field(=||F_metal,F_P,F_hal||),
                          SASA, residue_embedding(4)]
        - Ep_sim.npy    : Ep_theory plus 10 neighbor-geometry features
        - Ep_manifest.json : feature names, residue ids, params, etc.
"""

import os
import json
import argparse
import subprocess
from typing import Dict, List, Tuple, Optional

import numpy as np
import gc
import time
import signal

# --------------------------
# Optional deps
# --------------------------
try:
    from scipy.spatial import cKDTree
    HAS_KDTREE = True
except ImportError:
    HAS_KDTREE = False
    print("[WARN] scipy.spatial.cKDTree not available; falling back to O(N^2) contacts.")

try:
    import mdtraj as mdt
    HAS_MDTRAJ = True
except ImportError:
    HAS_MDTRAJ = False
    print("[ERROR] mdtraj is required for this pipeline.")
    raise SystemExit("mdtraj is required for this pipeline.")

try:
    import torch
    HAS_TORCH = True
except ImportError:
    HAS_TORCH = False
    torch = None
    print("[ERROR] torch is required for this GPU pipeline.")
    raise SystemExit("torch is required for this GPU pipeline.")

try:
    import psutil
    HAS_PSUTIL = True
except ImportError:
    HAS_PSUTIL = False
    psutil = None
    print("[WARN] psutil not available; memory debug helper disabled.")


# =====================================
# Global thresholds for vector GPU mode
# =====================================

MAX_VECTOR_ELEMENTS = 200_000_000
MAX_BATCH_ELEMENTS = 50_000_000


# ==============================
# 0. Utilities
# ==============================

def ensure_dir(path: str):
    os.makedirs(path, exist_ok=True)


def load_json(path: str) -> Dict:
    with open(path, "r") as f:
        return json.load(f)


def log_gpu_usage(tag: str, device: str):
    if not (HAS_TORCH and device == "cuda"):
        return
    if not torch.cuda.is_available():
        return
    try:
        torch.cuda.synchronize()
    except Exception:
        pass
    try:
        alloc = torch.cuda.memory_allocated() / 1024**2
        reserved = torch.cuda.memory_reserved() / 1024**2
        max_alloc = torch.cuda.max_memory_allocated() / 1024**2
    except Exception:
        return
    print(
        f"[GPU] {tag:24s} | "
        f"alloc={alloc:7.1f} MB | "
        f"reserved={reserved:7.1f} MB | "
        f"max={max_alloc:7.1f} MB"
    )


# ==============================
# 1. mdtraj helpers
# ==============================

def get_chain_id(chain) -> str:
    if hasattr(chain, "chain_id") and chain.chain_id is not None:
        return str(chain.chain_id)
    if hasattr(chain, "id") and chain.id is not None:
        return str(chain.id)
    return str(chain.index)


def get_insertion_code(res) -> str:
    raw = None
    if hasattr(res, "insertion_code"):
        raw = res.insertion_code
    elif hasattr(res, "insertionCode"):
        raw = res.insertionCode
    if raw is None:
        raw = ""
    icode = str(raw).strip()
    if icode == "None":
        icode = ""
    return icode


def load_traj(path: str):
    if not HAS_MDTRAJ:
        raise RuntimeError("mdtraj is required for load_traj.")
    ext = os.path.splitext(path)[1].lower()

    if ext in (".cif", ".mmcif"):
        print("[INFO] Detected CIF/mmCIF input → calling mdt.load without standard_names kwarg.")
        try:
            return mdt.load(path)
        except Exception as e:
            msg = str(e)
            print("[ERROR] mdtraj.load(cif) failed.")
            print("        First 200 chars of the error message:")
            print("        ", msg[:200].replace("\n", " "))
            raise

    try:
        return mdt.load(path, standard_names=True)
    except Exception as e:
        msg = str(e)
        print("[WARN] mdtraj.load(..., standard_names=True) failed:")
        print("       ", msg[:200].replace("\n", " "))
        print("[WARN] Retrying with standard_names=False (no canonicalization).")
        return mdt.load(path, standard_names=False)


def collect_residues_from_mdtraj(traj) -> List[Dict]:
    residues: List[Dict] = []
    topo = traj.topology
    coords0 = traj.xyz[0].astype(np.float32)

    for res in topo.residues:
        chain_id = get_chain_id(res.chain)
        resseq = res.resSeq
        icode = get_insertion_code(res)
        suffix = icode if icode else ""
        rid = f"{chain_id}:{resseq}{suffix}"

        atoms: List[Dict] = []
        for atom in res.atoms:
            idx = atom.index
            coord = np.array(coords0[idx, :], dtype=np.float32)
            if atom.element is not None:
                element = atom.element.symbol.upper()
            else:
                element = "?"
            atoms.append(
                {
                    "atom_index": idx,
                    "name": atom.name,
                    "element": element,
                    "coord": coord,
                }
            )

        if not atoms:
            continue

        residues.append(
            {
                "id": rid,
                "chain": chain_id,
                "resseq": resseq,
                "icode": icode,
                "resname": res.name,
                "hetflag": " ",
                "atoms": atoms,
            }
        )
    return residues


def compute_residue_sasa_from_traj(traj, residues: List[Dict]) -> Dict[str, float]:
    resid_to_sasa: Dict[str, float] = {r["id"]: 0.0 for r in residues}
    if not HAS_MDTRAJ:
        return resid_to_sasa

    try:
        sasa_res = mdt.shrake_rupley(traj, probe_radius=1.4, mode="residue")
        sasa_0 = sasa_res[0]
    except Exception as e:
        print("[WARN] mdtraj SASA (residue mode) failed:", e)
        return resid_to_sasa

    topo = traj.topology
    if topo.n_residues != sasa_0.shape[0]:
        print("[WARN] Topology residue count and SASA length mismatch.")
        return resid_to_sasa

    topo_res_sasa: Dict[Tuple[str, int, str], float] = {}
    for i, res in enumerate(topo.residues):
        chain_id = get_chain_id(res.chain)
        resseq = res.resSeq
        icode = get_insertion_code(res)
        key = (chain_id, int(resseq), icode)
        topo_res_sasa[key] = float(sasa_0[i])

    for r in residues:
        key = (str(r["chain"]), int(r["resseq"]), str(r["icode"]))
        if key in topo_res_sasa:
            resid_to_sasa[r["id"]] = topo_res_sasa[key]

    return resid_to_sasa


# ==============================
# 2. KDTree-based residue contacts
# ==============================

def make_pair_key(id_a: str, id_b: str) -> str:
    if id_a <= id_b:
        return f"{id_a}__{id_b}"
    else:
        return f"{id_b}__{id_a}"


def compute_residue_contacts_kdtree(
    residues: List[Dict],
    radius: float,
    store_pairs: bool = True,
) -> Tuple[Dict[str, List[str]], Dict[str, List[Dict]]]:
    atom_coords = []
    atom_res_idx = []
    res_ids = [r["id"] for r in residues]

    for i_res, r in enumerate(residues):
        for a in r["atoms"]:
            atom_coords.append(a["coord"])
            atom_res_idx.append(i_res)

    atom_coords = np.array(atom_coords, dtype=np.float32)
    atom_res_idx = np.array(atom_res_idx, dtype=int)
    n_atoms = atom_coords.shape[0]

    if store_pairs:
        overlaps: Dict[str, List[str]] = {rid: [] for rid in res_ids}
        pair_contacts: Dict[str, List[Dict]] = {}
    else:
        overlaps: Dict[str, set] = {rid: set() for rid in res_ids}
        pair_contacts: Dict[str, List[Dict]] = {}

    cutoff = 2.0 * radius

    if HAS_KDTREE:
        tree = cKDTree(atom_coords)
        for i in range(n_atoms):
            coord_i = atom_coords[i]
            res_i = atom_res_idx[i]
            rid_i = res_ids[res_i]

            idxs = tree.query_ball_point(coord_i, cutoff)
            for j in idxs:
                if j <= i:
                    continue
                res_j = atom_res_idx[j]
                if res_j == res_i:
                    continue
                rid_j = res_ids[res_j]
                d = float(np.linalg.norm(atom_coords[j] - coord_i))
                key = make_pair_key(rid_i, rid_j)

                if store_pairs:
                    if key not in pair_contacts:
                        pair_contacts[key] = []
                    pair_contacts[key].append(
                        {
                            "a_index": int(i),
                            "b_index": int(j),
                            "distance": d,
                        }
                    )

                if store_pairs:
                    overlaps[rid_i].append(rid_j)
                    overlaps[rid_j].append(rid_i)
                else:
                    overlaps[rid_i].add(rid_j)
                    overlaps[rid_j].add(rid_i)
    else:
        for i in range(n_atoms):
            coord_i = atom_coords[i]
            res_i = atom_res_idx[i]
            rid_i = res_ids[res_i]
            for j in range(i + 1, n_atoms):
                coord_j = atom_coords[j]
                res_j = atom_res_idx[j]
                if res_j == res_i:
                    continue
                d = float(np.linalg.norm(coord_j - coord_i))
                if d >= cutoff:
                    continue
                rid_j = res_ids[res_j]
                key = make_pair_key(rid_i, rid_j)

                if store_pairs:
                    if key not in pair_contacts:
                        pair_contacts[key] = []
                    pair_contacts[key].append(
                        {
                            "a_index": int(i),
                            "b_index": int(j),
                            "distance": d,
                        }
                    )

                if store_pairs:
                    overlaps[rid_i].append(rid_j)
                    overlaps[rid_j].append(rid_i)
                else:
                    overlaps[rid_i].add(rid_j)
                    overlaps[rid_j].add(rid_i)

    if store_pairs:
        for rid in overlaps:
            overlaps[rid] = sorted(set(overlaps[rid]))
    else:
        for rid in overlaps:
            overlaps[rid] = sorted(overlaps[rid])

    return overlaps, pair_contacts


def export_meta(
    pdb_name: str,
    radius: float,
    residues: List[Dict],
    overlaps: Dict[str, List[str]],
    pair_contacts: Dict[str, List[Dict]],
    resid_to_sasa: Dict[str, float],
    out_path: str,
):
    json_residues = []
    for r in residues:
        rid = r["id"]
        json_residues.append(
            {
                "id": rid,
                "chain": r["chain"],
                "resseq": r["resseq"],
                "icode": r["icode"],
                "resname": r["resname"],
                "hetflag": r["hetflag"],
                "sasa": float(resid_to_sasa.get(rid, 0.0)),
                "atoms": [
                    {
                        "atom_index": a["atom_index"],
                        "name": a["name"],
                        "element": a["element"],
                        "coord": np.array(a["coord"], dtype=float).tolist(),
                    }
                    for a in r["atoms"]
                ],
            }
        )

    meta = {
        "pdb": pdb_name,
        "radius": radius,
        "residues": json_residues,
        "overlaps": overlaps,
        "pair_contacts": pair_contacts,
    }

    with open(out_path, "w") as f:
        json.dump(meta, f, indent=2)

    print(f"[INFO] meta.json written: {out_path}")


# ==============================
# 3. Hybrid density core
# ==============================

Z_TABLE = {
    "H": 1,
    "C": 6,
    "N": 7,
    "O": 8,
    "S": 16,
    "P": 15,
    "F": 9,
    "CL": 17,
    "BR": 35,
    "I": 53,
    "ZN": 30,
    "MG": 12,
    "CA": 20,
    "FE": 26,
    "MN": 25,
    "CU": 29,
    "NI": 28,
    "CO": 27,
    "MO": 42,
    "W": 74,
    "V": 23,
}

SIGMA_TABLE = {
    "H": 0.7,
    "C": 1.0,
    "N": 0.95,
    "O": 0.9,
    "S": 1.2,
    "P": 1.2,
    "ZN": 1.1,
    "MG": 1.1,
    "CA": 1.3,
    "FE": 1.2,
    "MN": 1.2,
    "CU": 1.2,
    "NI": 1.2,
    "CO": 1.2,
}


def z_of(elem: str) -> int:
    return Z_TABLE.get(elem.upper(), 6)


def sigma_of(elem: str) -> float:
    return SIGMA_TABLE.get(elem.upper(), 1.0)


def build_residue_maps(meta: Dict):
    res_map = {r["id"]: r for r in meta["residues"]}
    overlaps = meta["overlaps"]

    atom_index_to_resid: Dict[int, str] = {}
    resid_to_atom_coords: Dict[str, List[np.ndarray]] = {}
    resid_to_sasa: Dict[str, float] = {}

    for r in meta["residues"]:
        rid = r["id"]
        coords = []
        for a in r["atoms"]:
            idx = int(a["atom_index"])
            atom_index_to_resid[idx] = rid
            coords.append(np.array(a["coord"], dtype=np.float32))
        resid_to_atom_coords[rid] = coords
        resid_to_sasa[rid] = float(r.get("sasa", 0.0))

    return res_map, overlaps, atom_index_to_resid, resid_to_atom_coords, resid_to_sasa


def build_global_atom_arrays(meta: Dict):
    """
    AoS(atom dict) → SoA(parallel arrays).

    Returns
    -------
    atom_coords : (N, 3) float32
    atom_Z      : (N,)   float32  (promolecular Z or surrogate weight)
    atom_sigma  : (N,)   float32  (Gaussian sigma per atom)
    resid_to_atom_indices : dict[rid] -> np.ndarray of atom indices
    """
    coords_list = []
    Z_list = []
    sigma_list = []
    resid_to_atom_indices: Dict[str, List[int]] = {}

    idx = 0
    for r in meta["residues"]:
        rid = r["id"]
        resid_to_atom_indices.setdefault(rid, [])
        for a in r["atoms"]:
            coord = np.asarray(a["coord"], dtype=np.float32)
            elem = str(a.get("element", "C")).strip().upper()
            coords_list.append(coord)
            Z_list.append(float(z_of(elem)))
            sigma_list.append(float(sigma_of(elem)))
            resid_to_atom_indices[rid].append(idx)
            idx += 1

    if len(coords_list) == 0:
        atom_coords = np.zeros((0, 3), dtype=np.float32)
        atom_Z = np.zeros((0,), dtype=np.float32)
        atom_sigma = np.zeros((0,), dtype=np.float32)
    else:
        atom_coords = np.stack(coords_list, axis=0).astype(np.float32)
        atom_Z = np.asarray(Z_list, dtype=np.float32)
        atom_sigma = np.asarray(sigma_list, dtype=np.float32)

    resid_to_atom_indices_np: Dict[str, np.ndarray] = {}
    for rid, idxs in resid_to_atom_indices.items():
        if len(idxs) == 0:
            resid_to_atom_indices_np[rid] = np.zeros((0,), dtype=np.int64)
        else:
            resid_to_atom_indices_np[rid] = np.asarray(idxs, dtype=np.int64)

    return atom_coords, atom_Z, atom_sigma, resid_to_atom_indices_np


def build_fragment_indices(
    center_res_id: str,
    neighbor_res_ids: List[str],
    resid_to_atom_indices: Dict[str, np.ndarray],
) -> np.ndarray:
    idx_arrays = []

    if center_res_id in resid_to_atom_indices:
        idx_arrays.append(resid_to_atom_indices[center_res_id])

    for nid in neighbor_res_ids:
        arr = resid_to_atom_indices.get(nid, None)
        if arr is not None and arr.size > 0:
            idx_arrays.append(arr)

    if not idx_arrays:
        return np.zeros((0,), dtype=np.int64)

    concat = np.concatenate(idx_arrays, axis=0)
    if concat.size == 0:
        return concat
    return np.unique(concat)


def build_fragment_atoms(
    res_map: Dict[str, Dict], center_res_id: str, neighbor_res_ids: List[str]
) -> List[Dict]:
    ids = [center_res_id] + neighbor_res_ids
    atoms: List[Dict] = []
    for rid in ids:
        if rid in res_map:
            atoms.extend(res_map[rid]["atoms"])
    return atoms


def precompute_residue_grids(
    resid_to_atom_coords: Dict[str, List[np.ndarray]],
    R_env: float,
    grid_step: float,
    padding: float,
) -> Tuple[Dict[str, Tuple[np.ndarray, np.ndarray, np.ndarray]], int, int, int]:
    resid_to_grid_axes: Dict[str, Tuple[np.ndarray, np.ndarray, np.ndarray]] = {}
    max_nx = 1
    max_ny = 1
    max_nz = 1

    extra = float(R_env + padding)

    for rid, coords_list in resid_to_atom_coords.items():
        if not coords_list:
            continue
        coords = np.stack(coords_list, axis=0).astype(np.float32)
        mn = coords.min(axis=0) - extra
        mx = coords.max(axis=0) + extra

        xs = np.arange(mn[0], mx[0] + grid_step, grid_step, dtype=np.float32)
        ys = np.arange(mn[1], mx[1] + grid_step, grid_step, dtype=np.float32)
        zs = np.arange(mn[2], mx[2] + grid_step, grid_step, dtype=np.float32)

        nx = xs.size
        ny = ys.size
        nz = zs.size

        resid_to_grid_axes[rid] = (xs, ys, zs)

        if nx > max_nx:
            max_nx = nx
        if ny > max_ny:
            max_ny = ny
        if nz > max_nz:
            max_nz = nz

    return resid_to_grid_axes, max_nx, max_ny, max_nz


# ==============================
# 3c. GPU density workspace
# ==============================

class DensityWorkspaceGPU:
    def __init__(self, max_nx: int, max_ny: int, max_nz: int, device: str):
        if not HAS_TORCH:
            raise RuntimeError("DensityWorkspaceGPU requires torch.")
        self.max_nx = int(max_nx)
        self.max_ny = int(max_ny)
        self.max_nz = int(max_nz)
        self.device = device

        shape = (self.max_nx, self.max_ny, self.max_nz)

        self.X = torch.empty(shape, dtype=torch.float32, device=self.device)
        self.Y = torch.empty_like(self.X)
        self.Z = torch.empty_like(self.X)

        self.rho = torch.empty_like(self.X)
        self.rho_local = torch.empty_like(self.X)

        self.mask = torch.empty(shape, dtype=torch.bool, device=self.device)

        self.dist2 = torch.empty_like(self.X)
        self.dx = torch.empty_like(self.X)
        self.dy = torch.empty_like(self.X)
        self.dz = torch.empty_like(self.X)
        self.tmp = torch.empty_like(self.X)

    def prepare_grid_from_axes(
        self,
        xs: np.ndarray,
        ys: np.ndarray,
        zs: np.ndarray,
    ) -> Tuple[torch.Tensor, torch.Tensor, torch.Tensor, int, int, int]:
        xs = np.asarray(xs, dtype=np.float32)
        ys = np.asarray(ys, dtype=np.float32)
        zs = np.asarray(zs, dtype=np.float32)

        nx = xs.size
        ny = ys.size
        nz = zs.size

        if nx > self.max_nx or ny > self.max_ny or nz > self.max_nz:
            raise ValueError(
                f"Workspace too small for residue grid: ({nx},{ny},{nz}) "
                f"vs ({self.max_nx},{self.max_ny},{self.max_nz})"
            )

        xs_t = torch.from_numpy(xs).to(self.device)
        ys_t = torch.from_numpy(ys).to(self.device)
        zs_t = torch.from_numpy(zs).to(self.device)

        X = self.X[:nx, :ny, :nz]
        Y = self.Y[:nx, :ny, :nz]
        Z = self.Z[:nx, :ny, :nz]

        X[:] = xs_t.view(-1, 1, 1)
        Y[:] = ys_t.view(1, -1, 1)
        Z[:] = zs_t.view(1, 1, -1)

        return X, Y, Z, nx, ny, nz

    def view_rho(self, nx: int, ny: int, nz: int) -> torch.Tensor:
        rho = self.rho[:nx, :ny, :nz]
        rho.zero_()
        return rho

    def view_mask(self, nx: int, ny: int, nz: int) -> torch.Tensor:
        mask = self.mask[:nx, :ny, :nz]
        mask.fill_(False)
        return mask

    def view_rho_local(self, nx: int, ny: int, nz: int) -> torch.Tensor:
        return self.rho_local[:nx, :ny, :nz]

    def view_dist2(self, nx: int, ny: int, nz: int) -> torch.Tensor:
        return self.dist2[:nx, :ny, :nz]

    def view_dx(self, nx: int, ny: int, nz: int) -> torch.Tensor:
        return self.dx[:nx, :ny, :nz]

    def view_dy(self, nx: int, ny: int, nz: int) -> torch.Tensor:
        return self.dy[:nx, :ny, :nz]

    def view_dz(self, nx: int, ny: int, nz: int) -> torch.Tensor:
        return self.dz[:nx, :ny, :nz]

    def view_tmp(self, nx: int, ny: int, nz: int) -> torch.Tensor:
        return self.tmp[:nx, :ny, :nz]


# ==============================
# 3d. GPU promolecular density
# ==============================

def promolecular_density_on_grid_gpu(
    atoms: List[Dict],
    X: torch.Tensor,
    Y: torch.Tensor,
    Z: torch.Tensor,
    rho_out: torch.Tensor,
    dx_ws: torch.Tensor,
    dy_ws: torch.Tensor,
    dz_ws: torch.Tensor,
    dist2_ws: torch.Tensor,
    tmp_ws: torch.Tensor,
):
    if not atoms:
        raise ValueError("atoms list is empty in promolecular_density_on_grid_gpu")

    coords = np.array([a["coord"] for a in atoms], dtype=np.float32)
    Zs = np.array([z_of(a.get("element", "C")) for a in atoms], dtype=np.float32)
    sigmas = np.array([sigma_of(a.get("element", "C")) for a in atoms], dtype=np.float32)

    device = X.device
    coords_t = torch.from_numpy(coords).to(device=device)
    Zs_t = torch.from_numpy(Zs).to(device=device)
    sigmas_t = torch.from_numpy(sigmas).to(device=device)

    rho = rho_out
    rho.zero_()

    dx = dx_ws
    dy = dy_ws
    dz = dz_ws
    dist2 = dist2_ws
    tmp = tmp_ws

    for i in range(coords_t.shape[0]):
        ra = coords_t[i]
        Za = Zs_t[i]
        sa = sigmas_t[i]

        x0, y0, z0 = ra[0], ra[1], ra[2]
        inv_2s2 = -1.0 / (2.0 * (sa * sa))

        dx.copy_(X); dx.add_(-x0)
        dy.copy_(Y); dy.add_(-y0)
        dz.copy_(Z); dz.add_(-z0)

        dist2.copy_(dx); dist2.mul_(dx)
        tmp.copy_(dy); tmp.mul_(dy); dist2.add_(tmp)
        tmp.copy_(dz); tmp.mul_(dz); dist2.add_(tmp)

        tmp.copy_(dist2); tmp.mul_(inv_2s2)
        torch.exp_(tmp)
        tmp.mul_(Za)
        rho.add_(tmp)

    return rho


def promolecular_density_on_grid_gpu_vector(
    atoms: List[Dict],
    X: torch.Tensor,
    Y: torch.Tensor,
    Z: torch.Tensor,
    rho_out: torch.Tensor,
):
    if not atoms:
        raise ValueError("atoms list is empty in promolecular_density_on_grid_gpu_vector")

    coords = np.array([a["coord"] for a in atoms], dtype=np.float32)
    Zs = np.array([z_of(a.get("element", "C")) for a in atoms], dtype=np.float32)
    sigmas = np.array([sigma_of(a.get("element", "C")) for a in atoms], dtype=np.float32)

    device = X.device
    coords_t = torch.from_numpy(coords).to(device=device)
    Zs_t = torch.from_numpy(Zs).to(device=device)
    sigmas_t = torch.from_numpy(sigmas).to(device=device)

    Xb = X.unsqueeze(0)
    Yb = Y.unsqueeze(0)
    Zb = Z.unsqueeze(0)

    x0 = coords_t[:, 0].view(-1, 1, 1, 1)
    y0 = coords_t[:, 1].view(-1, 1, 1, 1)
    z0 = coords_t[:, 2].view(-1, 1, 1, 1)

    dx = Xb - x0
    dy = Yb - y0
    dz = Zb - z0
    dist2 = dx * dx + dy * dy + dz * dz

    sigmas_t = sigmas_t.view(-1, 1, 1, 1)
    inv_2s2 = -1.0 / (2.0 * sigmas_t * sigmas_t)

    tmp = torch.exp(dist2 * inv_2s2)
    Zs_b = Zs_t.view(-1, 1, 1, 1)
    tmp = tmp * Zs_b

    rho = tmp.sum(dim=0)
    rho_out.zero_()
    rho_out.copy_(rho)
    return rho_out


def _atom_batch_density_core(
    coords_t: torch.Tensor,
    weights_t: torch.Tensor,
    sigmas_t: torch.Tensor,
    X: torch.Tensor,
    Y: torch.Tensor,
    Z: torch.Tensor,
    rho_out: torch.Tensor,
    max_batch_elems: int = MAX_BATCH_ELEMENTS,
) -> torch.Tensor:
    if coords_t.numel() == 0:
        raise ValueError("coords_t is empty in _atom_batch_density_core")

    device = X.device
    coords_t = coords_t.to(device=device, dtype=torch.float32)
    weights_t = weights_t.to(device=device, dtype=torch.float32)
    sigmas_t = sigmas_t.to(device=device, dtype=torch.float32)

    nx, ny, nz = X.shape
    rho = rho_out
    rho.zero_()

    natoms = coords_t.shape[0]
    elems_per_grid = int(nx * ny * nz)
    if elems_per_grid <= 0:
        return rho

    max_B = max(1, max_batch_elems // max(elems_per_grid, 1))
    max_B = int(max_B)
    max_B = min(max_B, natoms)
    if max_B < 1:
        max_B = 1

    Xb = X.unsqueeze(0)
    Yb = Y.unsqueeze(0)
    Zb = Z.unsqueeze(0)

    for start in range(0, natoms, max_B):
        end = min(start + max_B, natoms)
        B = end - start

        chunk_coords = coords_t[start:end]
        chunk_w = weights_t[start:end]
        chunk_sigmas = sigmas_t[start:end]

        cx = chunk_coords[:, 0].view(B, 1, 1, 1)
        cy = chunk_coords[:, 1].view(B, 1, 1, 1)
        cz = chunk_coords[:, 2].view(B, 1, 1, 1)

        dist2_b = Xb - cx
        dist2_b.mul_(dist2_b)

        dy_b = Yb - cy
        dy_b.mul_(dy_b)
        dist2_b.add_(dy_b)
        del dy_b

        dz_b = Zb - cz
        dz_b.mul_(dz_b)
        dist2_b.add_(dz_b)
        del dz_b

        s_b = chunk_sigmas.view(B, 1, 1, 1)
        inv_2s2 = -1.0 / (2.0 * (s_b * s_b))

        tmp_b = dist2_b * inv_2s2
        torch.exp_(tmp_b)

        w_b = chunk_w.view(B, 1, 1, 1)
        tmp_b.mul_(w_b)

        rho.add_(tmp_b.sum(dim=0))

        del dist2_b, tmp_b, cx, cy, cz, s_b, inv_2s2, w_b

    return rho


def promolecular_density_on_grid_gpu_mode(
    atoms: List[Dict],
    X: torch.Tensor,
    Y: torch.Tensor,
    Z: torch.Tensor,
    rho_out: torch.Tensor,
    dx_ws: torch.Tensor,
    dy_ws: torch.Tensor,
    dz_ws: torch.Tensor,
    dist2_ws: torch.Tensor,
    tmp_ws: torch.Tensor,
    mode: str,
    rid: str,
) -> Tuple[torch.Tensor, str, bool]:
    nx, ny, nz = X.shape
    n_atoms = len(atoms)
    total_elems = int(n_atoms * nx * ny * nz)

    if mode == "safe":
        rho = promolecular_density_on_grid_gpu(
            atoms, X, Y, Z, rho_out, dx_ws, dy_ws, dz_ws, dist2_ws, tmp_ws
        )
        return rho, "safe", False

    if mode == "batch":
        if not atoms:
            raise ValueError("atoms list is empty in promolecular_density_on_grid_gpu_mode(batch)")
        coords = np.array([a["coord"] for a in atoms], dtype=np.float32)
        Zs = np.array([z_of(a.get("element", "C")) for a in atoms], dtype=np.float32)
        sigmas = np.array([sigma_of(a.get("element", "C")) for a in atoms], dtype=np.float32)

        device = X.device
        coords_t = torch.from_numpy(coords).to(device=device)
        Zs_t = torch.from_numpy(Zs).to(device=device)
        sigmas_t = torch.from_numpy(sigmas).to(device=device)

        rho = _atom_batch_density_core(
            coords_t=coords_t,
            weights_t=Zs_t,
            sigmas_t=sigmas_t,
            X=X,
            Y=Y,
            Z=Z,
            rho_out=rho_out,
            max_batch_elems=MAX_BATCH_ELEMENTS,
        )
        return rho, "batch", False

    if mode == "vector":
        if total_elems > MAX_VECTOR_ELEMENTS:
            print(
                f"[WARN] gpu-density-mode 'vector' skipped for fragment {rid} "
                f"(atoms={n_atoms}, grid={nx}x{ny}x{nz}, elems={total_elems}) "
                f"→ falling back to 'safe' atom-loop to avoid GPU OOM."
            )
            rho = promolecular_density_on_grid_gpu(
                atoms, X, Y, Z, rho_out, dx_ws, dy_ws, dz_ws, dist2_ws, tmp_ws
            )
            return rho, "safe", True

        rho = promolecular_density_on_grid_gpu_vector(
            atoms, X, Y, Z, rho_out
        )
        return rho, "vector", False

    rho = promolecular_density_on_grid_gpu(
        atoms, X, Y, Z, rho_out, dx_ws, dy_ws, dz_ws, dist2_ws, tmp_ws
    )
    return rho, "safe", False


def promolecular_density_on_grid_gpu_mode_arrays(
    atom_indices: np.ndarray,
    atom_coords: np.ndarray,
    atom_Z: np.ndarray,
    atom_sigma: np.ndarray,
    X: torch.Tensor,
    Y: torch.Tensor,
    Z: torch.Tensor,
    rho_out: torch.Tensor,
    dx_ws: torch.Tensor,
    dy_ws: torch.Tensor,
    dz_ws: torch.Tensor,
    dist2_ws: torch.Tensor,
    tmp_ws: torch.Tensor,
    mode: str,
    rid: str,
) -> Tuple[torch.Tensor, str, bool]:
    """
    Legacy approach: Received `atoms` (list[dict]) and reconstructed coords/Z/sigma for every residue.
    Optimized approach: Uses global SoA (Structure of Arrays) + fragment `atom_indices` 
                        for direct tensor operations without Python loops.

    Returns
    -------
    rho      : torch.Tensor, shape (nx, ny, nz)
    eff_mode : "batch" or "vector"
    fallback : True if "vector" was requested but fell back to "batch" due to element limits.
    """
    nx, ny, nz = X.shape

    if atom_indices is None or atom_indices.size == 0:
        rho_out.zero_()
        return rho_out, "batch", False

    # Slice global SoA arrays to get fragment data (pure indexing, no Python loops)
    coords = atom_coords[atom_indices]          # (N, 3)
    Zs = atom_Z[atom_indices]                   # (N,)
    sigmas = atom_sigma[atom_indices]           # (N,)

    device = X.device
    coords_t = torch.from_numpy(coords).to(device=device, dtype=torch.float32)
    Zs_t = torch.from_numpy(Zs).to(device=device, dtype=torch.float32)
    sigmas_t = torch.from_numpy(sigmas).to(device=device, dtype=torch.float32)

    n_atoms = coords_t.shape[0]
    elems_per_grid = int(nx * ny * nz)
    total_elems = int(n_atoms * elems_per_grid)

    # Safety mechanism: Force 'batch' mode if (num_atoms * grid_size) exceeds memory limits
    if mode == "vector" and total_elems > MAX_VECTOR_ELEMENTS:
        print(
            f"[WARN] gpu-density-mode 'vector' skipped for fragment {rid} "
            f"(atoms={n_atoms}, grid={nx}x{ny}x{nz}, elems={total_elems}) "
            f"→ falling back to 'batch' array-mode."
        )
        mode = "batch"
        fallback = True
    else:
        fallback = False

    # Computation: 'batch' mode delegates to _atom_batch_density_core
    if mode in ("safe", "batch"):
        rho = _atom_batch_density_core(
            coords_t=coords_t,
            weights_t=Zs_t,
            sigmas_t=sigmas_t,
            X=X,
            Y=Y,
            Z=Z,
            rho_out=rho_out,
            max_batch_elems=MAX_BATCH_ELEMENTS,
        )
        return rho, "batch", fallback

    # Case mode == "vector": Perform one-shot broadcasting
    Xb = X.unsqueeze(0)  # (1, nx, ny, nz)
    Yb = Y.unsqueeze(0)
    Zb = Z.unsqueeze(0)

    cx = coords_t[:, 0].view(-1, 1, 1, 1)  # (N,1,1,1)
    cy = coords_t[:, 1].view(-1, 1, 1, 1)
    cz = coords_t[:, 2].view(-1, 1, 1, 1)

    dx = Xb - cx
    dy = Yb - cy
    dz = Zb - cz
    dist2 = dx * dx + dy * dy + dz * dz

    sigmas_b = sigmas_t.view(-1, 1, 1, 1)
    inv_2s2 = -1.0 / (2.0 * sigmas_b * sigmas_b)

    tmp = torch.exp(dist2 * inv_2s2)
    Zs_b = Zs_t.view(-1, 1, 1, 1)
    tmp = tmp * Zs_b

    rho = tmp.sum(dim=0)
    rho_out.zero_()
    rho_out.copy_(rho)
    return rho_out, "vector", fallback


# ==============================
# 3e. xTB-based density (A-mode)
# ==============================

def write_xyz(atoms: List[Dict], xyz_path: str, charge: int = 0, mult: int = 1):
    with open(xyz_path, "w") as f:
        f.write(f"{len(atoms)}\n")
        f.write(f"charge={charge} mult={mult}\n")
        for a in atoms:
            elem = a.get("element", "C")
            x, y, z = a["coord"]
            f.write(f"{elem:2s} {x: .6f} {y: .6f} {z: .6f}\n")


def run_xtb(xyz_path: str, workdir: str, xtb_bin: str = "xtb") -> Optional[str]:
    ensure_dir(workdir)
    cmd = [xtb_bin, os.path.basename(xyz_path), "--gfn2", "--charges"]
    try:
        subprocess.run(
            cmd,
            cwd=workdir,
            check=True,
            stdout=subprocess.PIPE,
            stderr=subprocess.PIPE,
            text=True,
        )
    except FileNotFoundError:
        print(f"[xTB ERROR] Binary '{xtb_bin}' not found. Falling back to B-mode.")
        return None
    except subprocess.CalledProcessError as e:
        print("[xTB FAIL]", e.stderr[:300])
        return None

    charges_path = os.path.join(workdir, "charges")
    return charges_path if os.path.exists(charges_path) else None


def parse_xtb_charges(charges_path: str) -> List[float]:
    vals = []
    with open(charges_path, "r") as f:
        for line in f:
            line = line.strip()
            if not line or line.startswith("#"):
                continue
            parts = line.split()
            try:
                vals.append(float(parts[-1]))
            except Exception:
                continue
    return vals


def density_from_charges_on_grid_gpu(
    atoms: List[Dict],
    charges: List[float],
    X: torch.Tensor,
    Y: torch.Tensor,
    Z: torch.Tensor,
    rho_out: torch.Tensor,
    dx_ws: torch.Tensor,
    dy_ws: torch.Tensor,
    dz_ws: torch.Tensor,
    dist2_ws: torch.Tensor,
    tmp_ws: torch.Tensor,
):
    if len(charges) != len(atoms):
        return promolecular_density_on_grid_gpu(
            atoms, X, Y, Z, rho_out, dx_ws, dy_ws, dz_ws, dist2_ws, tmp_ws
        )

    coords = np.array([a["coord"] for a in atoms], dtype=np.float32)
    charges_arr = np.array(charges, dtype=np.float32)

    device = X.device
    coords_t = torch.from_numpy(coords).to(device=device)
    charges_t = torch.from_numpy(charges_arr).to(device=device)

    rho = rho_out
    rho.zero_()

    dx = dx_ws
    dy = dy_ws
    dz = dz_ws
    dist2 = dist2_ws
    tmp = tmp_ws

    for i in range(coords_t.shape[0]):
        r = coords_t[i]
        q = charges_t[i]
        elem = atoms[i].get("element", "C")
        Za = float(z_of(elem))
        sa = float(sigma_of(elem))
        eff = max(Za - float(q.item()), 0.1)
        inv_2s2 = -1.0 / (2.0 * (sa * sa))

        x0, y0, z0 = r[0], r[1], r[2]

        dx.copy_(X); dx.add_(-x0)
        dy.copy_(Y); dy.add_(-y0)
        dz.copy_(Z); dz.add_(-z0)

        dist2.copy_(dx); dist2.mul_(dx)
        tmp.copy_(dy); tmp.mul_(dy); dist2.add_(tmp)
        tmp.copy_(dz); tmp.mul_(dz); dist2.add_(tmp)

        tmp.copy_(dist2); tmp.mul_(inv_2s2)
        torch.exp_(tmp)
        tmp.mul_(eff)
        rho.add_(tmp)

    return rho


def density_from_charges_on_grid_gpu_vector(
    atoms: List[Dict],
    charges: List[float],
    X: torch.Tensor,
    Y: torch.Tensor,
    Z: torch.Tensor,
    rho_out: torch.Tensor,
):
    if len(charges) != len(atoms):
        return promolecular_density_on_grid_gpu_vector(
            atoms, X, Y, Z, rho_out
        )

    coords = np.array([a["coord"] for a in atoms], dtype=np.float32)
    charges_arr = np.array(charges, dtype=np.float32)

    effZ = []
    sigmas = []
    for a, q in zip(atoms, charges_arr):
        elem = a.get("element", "C")
        Za = z_of(elem)
        sa = sigma_of(elem)
        eff = max(Za - float(q), 0.1)
        effZ.append(eff)
        sigmas.append(sa)

    effZ = np.array(effZ, dtype=np.float32)
    sigmas = np.array(sigmas, dtype=np.float32)

    device = X.device
    coords_t = torch.from_numpy(coords).to(device=device)
    effZ_t = torch.from_numpy(effZ).to(device=device)
    sigmas_t = torch.from_numpy(sigmas).to(device=device)

    Xb = X.unsqueeze(0)
    Yb = Y.unsqueeze(0)
    Zb = Z.unsqueeze(0)

    x0 = coords_t[:, 0].view(-1, 1, 1, 1)
    y0 = coords_t[:, 1].view(-1, 1, 1, 1)
    z0 = coords_t[:, 2].view(-1, 1, 1, 1)

    dx = Xb - x0
    dy = Yb - y0
    dz = Zb - z0
    dist2 = dx * dx + dy * dy + dz * dz

    sigmas_t = sigmas_t.view(-1, 1, 1, 1)
    inv_2s2 = -1.0 / (2.0 * sigmas_t * sigmas_t)

    tmp = torch.exp(dist2 * inv_2s2)
    effZ_b = effZ_t.view(-1, 1, 1, 1)
    tmp = tmp * effZ_b

    rho = tmp.sum(dim=0)
    rho_out.zero_()
    rho_out.copy_(rho)
    return rho_out


def density_from_charges_on_grid_gpu_mode(
    atoms: List[Dict],
    charges: List[float],
    X: torch.Tensor,
    Y: torch.Tensor,
    Z: torch.Tensor,
    rho_out: torch.Tensor,
    dx_ws: torch.Tensor,
    dy_ws: torch.Tensor,
    dz_ws: torch.Tensor,
    dist2_ws: torch.Tensor,
    tmp_ws: torch.Tensor,
    mode: str,
    rid: str,
) -> Tuple[torch.Tensor, str, bool]:
    nx, ny, nz = X.shape
    n_atoms = len(atoms)
    total_elems = int(n_atoms * nx * ny * nz)

    if mode == "safe":
        rho = density_from_charges_on_grid_gpu(
            atoms, charges, X, Y, Z, rho_out, dx_ws, dy_ws, dz_ws, dist2_ws, tmp_ws
        )
        return rho, "safe", False

    if mode == "batch":
        if len(charges) != len(atoms):
            rho = density_from_charges_on_grid_gpu(
                atoms, charges, X, Y, Z, rho_out, dx_ws, dy_ws, dz_ws, dist2_ws, tmp_ws
            )
            return rho, "batch", False

        if not atoms:
            raise ValueError("atoms list is empty in density_from_charges_on_grid_gpu_mode(batch)")

        coords = np.array([a["coord"] for a in atoms], dtype=np.float32)
        charges_arr = np.array(charges, dtype=np.float32)

        effZ = []
        sigmas = []
        for a, q in zip(atoms, charges_arr):
            elem = a.get("element", "C")
            Za = z_of(elem)
            sa = sigma_of(elem)
            eff = max(Za - float(q), 0.1)
            effZ.append(eff)
            sigmas.append(sa)

        effZ = np.array(effZ, dtype=np.float32)
        sigmas = np.array(sigmas, dtype=np.float32)

        device = X.device
        coords_t = torch.from_numpy(coords).to(device=device)
        effZ_t = torch.from_numpy(effZ).to(device=device)
        sigmas_t = torch.from_numpy(sigmas).to(device=device)

        rho = _atom_batch_density_core(
            coords_t=coords_t,
            weights_t=effZ_t,
            sigmas_t=sigmas_t,
            X=X,
            Y=Y,
            Z=Z,
            rho_out=rho_out,
            max_batch_elems=MAX_BATCH_ELEMENTS,
        )
        return rho, "batch", False

    if mode == "vector":
        if total_elems > MAX_VECTOR_ELEMENTS:
            print(
                f"[WARN] gpu-density-mode 'vector' (A-mode) skipped for fragment {rid} "
                f"(atoms={n_atoms}, grid={nx}x{ny}x{nz}, elems={total_elems}) "
                f"→ falling back to 'safe' atom-loop to avoid GPU OOM."
            )
            rho = density_from_charges_on_grid_gpu(
                atoms, charges, X, Y, Z, rho_out, dx_ws, dy_ws, dz_ws, dist2_ws, tmp_ws
            )
            return rho, "safe", True

        rho = density_from_charges_on_grid_gpu_vector(
            atoms, charges, X, Y, Z, rho_out
        )
        return rho, "vector", False

    rho = density_from_charges_on_grid_gpu(
        atoms, charges, X, Y, Z, rho_out, dx_ws, dy_ws, dz_ws, dist2_ws, tmp_ws
    )
    return rho, "safe", False


# ==============================
# 4. Density-based features
# ==============================

def density_features_for_residue_gpu(
    rho: torch.Tensor,
    X: torch.Tensor,
    Y: torch.Tensor,
    Z: torch.Tensor,
    ref_point: np.ndarray,
    res_atom_coords: np.ndarray,
    R_env: float,
    mask_ws: torch.Tensor,
    rho_local_ws: torch.Tensor,
    dist2_ws: torch.Tensor,
    tmp_ws: torch.Tensor,
    dx_ws: torch.Tensor,
    dy_ws: torch.Tensor,
    dz_ws: torch.Tensor,
    rid: str,
) -> Dict:
    device = rho.device

    xs = X[:, 0, 0]
    ys = Y[0, :, 0]
    zs = Z[0, 0, :]

    dx_step = float((xs[1] - xs[0]).item()) if xs.numel() > 1 else 1.0
    dy_step = float((ys[1] - ys[0]).item()) if ys.numel() > 1 else 1.0
    dz_step = float((zs[1] - zs[0]).item()) if zs.numel() > 1 else 1.0

    dV = dx_step * dy_step * dz_step
    R2 = float(R_env * R_env)

    mask = mask_ws
    mask.fill_(False)

    coords_res_t = torch.from_numpy(res_atom_coords.astype(np.float32)).to(device=device)

    dist2 = dist2_ws
    tmp = tmp_ws
    dx = dx_ws
    dy = dy_ws
    dz = dz_ws

    for i in range(coords_res_t.shape[0]):
        ra = coords_res_t[i]
        x0, y0, z0 = ra[0], ra[1], ra[2]

        dx.copy_(X); dx.add_(-x0)
        dy.copy_(Y); dy.add_(-y0)
        dz.copy_(Z); dz.add_(-z0)

        dist2.copy_(dx); dist2.mul_(dx)
        tmp.copy_(dy); tmp.mul_(dy); dist2.add_(tmp)
        tmp.copy_(dz); tmp.mul_(dz); dist2.add_(tmp)

        mask |= dist2 <= R2

    rho_local = rho_local_ws
    rho_local.zero_()
    rho_local[mask] = rho[mask]

    total_density = float((rho_local.sum() * dV).item())

    ref_t = torch.from_numpy(ref_point.astype(np.float32)).to(device=device)

    if total_density <= 0.0:
        centroid = ref_point.astype(float)
        dipole = np.zeros(3, dtype=float)
    else:
        cx = float((rho_local * X).sum().mul(dV).div(total_density).item())
        cy = float((rho_local * Y).sum().mul(dV).div(total_density).item())
        cz = float((rho_local * Z).sum().mul(dV).div(total_density).item())
        centroid = np.array([cx, cy, cz], dtype=float)

        RX = X - ref_t[0]
        RY = Y - ref_t[1]
        RZ = Z - ref_t[2]

        px = float((rho_local * RX).sum().mul(dV).item())
        py = float((rho_local * RY).sum().mul(dV).item())
        pz = float((rho_local * RZ).sum().mul(dV).item())
        dipole = np.array([px, py, pz], dtype=float)

    curv_val = 0.0
    curv_failed = False
    try:
        gx, gy, gz = torch.gradient(
            rho_local,
            spacing=(dx_step, dy_step, dz_step),
            edge_order=1,
        )
        curv = (gx * gx + gy * gy + gz * gz).sum().mul(dV)
        curv_val = float(curv.item())
    except Exception as e:
        print(f"[WARN] curv_rho computation failed for residue {rid}: {e}")
        curv_val = 0.0
        curv_failed = True

    return {
        "total_density": total_density,
        "centroid": centroid.tolist(),
        "dipole_rho": dipole.tolist(),
        "curv_rho": curv_val,
        "curv_rho_failed": curv_failed,
    }


# ==============================
# 4b. Charge-based features / surrogate
# ==============================

def charge_features_for_residue_from_fragment(
    rid: str,
    atom_index_to_resid: Dict[int, str],
    resid_to_atom_coords: Dict[str, List[np.ndarray]],
    frag_atoms: List[Dict],
    charges: List[float],
) -> Optional[Dict]:
    if len(frag_atoms) != len(charges):
        return None

    coords_res = resid_to_atom_coords[rid]
    ref_point = np.mean(np.stack(coords_res, axis=0), axis=0)

    q_res = 0.0
    dipole_q = np.zeros(3, dtype=float)

    for a, q in zip(frag_atoms, charges):
        atom_idx = int(a["atom_index"])
        res_of_atom = atom_index_to_resid.get(atom_idx, None)
        if res_of_atom != rid:
            continue
        coord = np.array(a["coord"], dtype=float)
        q_res += q
        dipole_q += q * (coord - ref_point)

    return {
        "partial_charge": float(q_res),
        "dipole_q": dipole_q.tolist(),
    }


def residue_base_charge(resname: str) -> float:
    rn = resname.strip().upper()
    if rn in ("ASP", "GLU"):
        return -1.0
    if rn in ("LYS", "ARG"):
        return +1.0
    if rn == "HIS":
        return +0.1
    return 0.0


def env_scale_from_sasa(sasa: float, mean_sasa: float, alpha: float = 0.4) -> float:
    if mean_sasa <= 1e-8:
        return 1.0
    r = sasa / mean_sasa
    scale = 1.0 + alpha * (r - 1.0)
    return float(np.clip(scale, 0.5, 1.5))


def residue_atom_charge_surrogate(residue) -> float:
    elem_weights = {
        "O":  -0.40,
        "N":  +0.30,
        "S":  -0.25,
        "SE": -0.25,
        "P":  -0.35,
        "F":  -0.20,
        "CL": -0.20,
        "BR": -0.20,
        "I":  -0.20,
        "MG": +0.50,
        "CA": +0.50,
        "MN": +0.60,
        "FE": +0.60,
        "ZN": +0.60,
        "CU": +0.60,
        "NI": +0.60,
        "CO": +0.60,
        # C, H, others → 0.0
    }

    atoms = residue.get("atoms", [])
    q = 0.0
    for atom in atoms:
        elem_raw = atom.get("element", "")
        elem = str(elem_raw).strip().upper()
        q += elem_weights.get(elem, 0.0)

    return float(q)


def compute_b_mode_charge_atom_based(
    res,
    rid,
    resid_to_sasa2,
    mean_sasa,
    sasa_alpha,
    env_scale_from_sasa,
) -> float:
    sasa_val = float(resid_to_sasa2.get(rid, 0.0))
    base_q = residue_atom_charge_surrogate(res)

    if abs(base_q) > 1e-6:
        scale = env_scale_from_sasa(sasa_val, mean_sasa, alpha=sasa_alpha)
    else:
        scale = 1.0

    partial_charge_B = base_q * scale
    return partial_charge_B


# ==============================
# 5. Ep helpers (embedding, neighbor, field)
# ==============================

RES_EMBED = {
    "ALA": [0.0, 0.5, 0.2, 0.0],
    "ARG": [1.0, -1.0, 1.0, 0.0],
    "ASN": [0.0, -0.5, 0.4, 0.0],
    "ASP": [-1.0, -1.0, 0.4, 0.0],
    "ASX": [-0.5, -1.0, 0.4, 0.0],
    "CSO": [0.0, -0.5, 0.4, 0.0],
    "CYS": [0.0, 0.5, 0.4, 0.0],
    "GLN": [0.0, -0.5, 0.6, 0.0],
    "GLU": [-1.0, -1.0, 0.6, 0.0],
    "GLX": [-0.5, -1.0, 0.6, 0.0],
    "GLY": [0.0, 0.0, 0.0, 0.0],
    "HIS": [0.5, -0.5, 0.6, 0.5],
    "HYP": [0.0, -0.5, 0.4, 0.0],
    "ILE": [0.0, 1.0, 0.8, 0.0],
    "LEU": [0.0, 1.0, 0.8, 0.0],
    "LYS": [1.0, -1.0, 0.8, 0.0],
    "MET": [0.0, 0.5, 0.8, 0.0],
    "MSE": [0.0, 0.5, 0.8, 0.0],
    "PHE": [0.0, 1.0, 1.0, 1.0],
    "PRO": [0.0, 0.0, 0.4, 0.0],
    "PYL": [1.0, -1.0, 1.0, 0.0],
    "SEC": [0.0, 0.5, 0.4, 0.0],
    "SER": [0.0, -0.5, 0.2, 0.0],
    "THR": [0.0, -0.5, 0.4, 0.0],
    "TRP": [0.0, 1.0, 1.0, 1.0],
    "TYR": [0.0, 0.5, 1.0, 1.0],
    "VAL": [0.0, 1.0, 0.6, 0.0],
    "XAA": [0.0, 0.0, 0.6, 0.0],
    "XLE": [0.0, 1.0, 0.8, 0.0],
}

DEFAULT_EMBED = np.array([0.0, 0.0, 0.50, 0.50], dtype=float)


def residue_to_embedding(resname: str) -> np.ndarray:
    rn = resname.strip().upper()
    vec = RES_EMBED.get(rn, RES_EMBED["XAA"])
    return np.array(vec, dtype=float)


def build_residue_centers(meta: Dict) -> Dict[str, np.ndarray]:
    centers: Dict[str, np.ndarray] = {}
    for r in meta["residues"]:
        coords = np.array([a["coord"] for a in r["atoms"]], dtype=float)
        centers[r["id"]] = coords.mean(axis=0)
    return centers


# ---------- NEW: 10D geometric neighbor block ----------

def compute_geometric_block(
    center: np.ndarray,
    neighbor_coords: np.ndarray,
    r_max: float,
    eps_shell: float = 1e-6,
    eps_pca: float = 1e-8,
) -> np.ndarray:
    """
    Compute 10 scalar neighbor-geometry features for residue p:

      (7)  N_in(p)          : #neighbors with d <= r_max/2
      (8)  N_out(p)         : #neighbors with r_max/2 < d <= r_max
      (9)  R_shell(p)       : N_in / (N_out + eps_shell)

      (10) r_mean(p)        : mean neighbor distance
      (11) r_var(p)         : variance of neighbor distances

      (12) linearity(p)
      (13) planarity(p)
      (14) isotropy(p)      : eigenvalue-based anisotropy of direction tensor

      (15) H_mean(p)        : mean curvature surrogate
      (16) K_gauss(p)       : Gaussian curvature surrogate

    Curvature is estimated by projecting neighbors into a local PCA frame
    and fitting a quadratic patch z = a x^2 + b y^2 + c x y.
    """
    if neighbor_coords.size == 0:
        return np.zeros(10, dtype=float)

    center = np.asarray(center, dtype=float)
    neighbor_coords = np.asarray(neighbor_coords, dtype=float)

    # distances
    vecs = neighbor_coords - center
    dists = np.linalg.norm(vecs, axis=1)
    if dists.size == 0:
        return np.zeros(10, dtype=float)

    # restrict to neighbors within r_max for shell definition
    # (contacts are already defined by KDTree, so this is mostly a cap)
    if r_max is not None and r_max > 0.0:
        mask_r = dists <= r_max
        if not np.any(mask_r):
            # fallback: use all neighbors anyway
            dists_use = dists
            vecs_use = vecs
        else:
            dists_use = dists[mask_r]
            vecs_use = vecs[mask_r]
    else:
        dists_use = dists
        vecs_use = vecs

    if dists_use.size == 0:
        return np.zeros(10, dtype=float)

    # ---------- 1. Two-shell radial density ----------
    r_cut = r_max if (r_max is not None and r_max > 0.0) else float(dists_use.max())
    if r_cut < eps_shell:
        r_cut = float(dists_use.max())

    r_inner = 0.5 * r_cut

    N_in = float(np.sum(dists_use <= r_inner))
    N_out = float(np.sum((dists_use > r_inner) & (dists_use <= r_cut)))
    R_shell = N_in / (N_out + eps_shell)

    # ---------- 2. Radial distribution moments ----------
    r_mean = float(dists_use.mean())
    r_var = float(dists_use.var()) if dists_use.size > 1 else 0.0

    # ---------- 3. Angular shape tensor / anisotropy ----------
    unit_vecs = []
    for v, d in zip(vecs_use, dists_use):
        if d <= eps_pca:
            continue
        unit_vecs.append(v / d)
    if len(unit_vecs) == 0:
        linearity = 0.0
        planarity = 0.0
        isotropy = 1.0
    else:
        U = np.stack(unit_vecs, axis=0)
        mean_u = U.mean(axis=0)
        centered = U - mean_u
        C = centered.T @ centered / max(U.shape[0], 1)
        evals = np.linalg.eigvalsh(C)
        evals = np.sort(np.maximum(evals, 0.0))
        lam1, lam2, lam3 = evals[-1], evals[-2], evals[-3]

        denom = lam1 + lam2 + lam3 + eps_pca
        linearity = (lam1 - lam2) / denom
        planarity = (lam2 - lam3) / denom
        isotropy = lam3 / denom

    # ---------- 4. Local surface curvature surrogate ----------
    # PCA to get local frame
    if vecs_use.shape[0] >= 3:
        # covariance for PCA
        mean_v = vecs_use.mean(axis=0)
        C_full = (vecs_use - mean_v).T @ (vecs_use - mean_v) / max(vecs_use.shape[0], 1)
        evals_full, evecs_full = np.linalg.eigh(C_full)
        idx = np.argsort(evals_full)[::-1]
        evecs_full = evecs_full[:, idx]

        # local coordinates
        local_coords = (vecs_use @ evecs_full)
        x = local_coords[:, 0]
        y = local_coords[:, 1]
        z = local_coords[:, 2]

        # quadratic fit: z = a x^2 + b y^2 + c x y
        A = np.stack([x * x, y * y, x * y], axis=1)
        try:
            # if underdetermined, lstsq still returns minimal norm solution
            coeffs, *_ = np.linalg.lstsq(A, z, rcond=None)
            a, b, c = coeffs.tolist()
            H = float(a + b)        # mean curvature surrogate
            K = float(a * b - 0.25 * c * c)  # Gaussian curvature surrogate
        except Exception:
            H = 0.0
            K = 0.0
    else:
        H = 0.0
        K = 0.0

    return np.array(
        [
            N_in,        # (7)
            N_out,       # (8)
            R_shell,     # (9)
            r_mean,      # (10)
            r_var,       # (11)
            linearity,   # (12)
            planarity,   # (13)
            isotropy,    # (14)
            H,           # (15)
            K,           # (16)
        ],
        dtype=float,
    )


def geometric_block_from_overlaps(
    rid: str,
    overlaps: Dict[str, List[str]],
    centers: Dict[str, np.ndarray],
    r_max: float,
) -> np.ndarray:
    """
    Build the 10D geometric neighbor feature block for residue `rid`
    from the contact graph overlaps and residue centers.
    """
    if rid not in centers:
        return np.zeros(10, dtype=float)

    neigh_ids = overlaps.get(rid, [])
    if not neigh_ids:
        return np.zeros(10, dtype=float)

    center = centers[rid]
    coords = []
    for nid in neigh_ids:
        if nid in centers:
            coords.append(centers[nid])

    if not coords:
        return np.zeros(10, dtype=float)

    neighbor_coords = np.stack(coords, axis=0)
    return compute_geometric_block(center=center, neighbor_coords=neighbor_coords, r_max=r_max)


FIELD_METALS = {"MG", "CA", "MN", "FE", "ZN", "CU", "NI", "CO", "MO", "W", "V"}
FIELD_P = {"P"}
FIELD_HALOGENS = {"F", "CL", "BR", "I"}


def compute_field_terms_for_residue(
    rid: str,
    res_map: Dict[str, Dict],
    centers: Dict[str, np.ndarray],
    overlaps: Dict[str, List[str]],
    eps: float = 1e-3,
) -> Tuple[float, float, float]:
    """
    Compute local field-like terms around residue p:

      F_metal(p) = sum_{metal atoms a in p ∪ N(p)} 1 / ||r_a - r_p||^2
      F_P(p)     = same for P atoms
      F_hal(p)   = same for halogens (F, Cl, Br, I)

    Only center residue + contact neighbors are used (overlaps).
    """
    if rid not in centers:
        return 0.0, 0.0, 0.0

    rp = centers[rid]
    F_metal = 0.0
    F_P_val = 0.0
    F_hal = 0.0

    neighbor_ids = [rid] + overlaps.get(rid, [])
    for nid in neighbor_ids:
        res = res_map.get(nid)
        if res is None:
            continue
        for a in res["atoms"]:
            elem = str(a.get("element", "")).strip().upper()
            if elem not in FIELD_METALS and elem not in FIELD_P and elem not in FIELD_HALOGENS:
                continue
            coord = np.array(a["coord"], dtype=float)
            d = float(np.linalg.norm(coord - rp))
            d_eff = max(d, eps)
            contrib = 1.0 / (d_eff * d_eff)

            if elem in FIELD_METALS:
                F_metal += contrib
            elif elem in FIELD_P:
                F_P_val += contrib
            elif elem in FIELD_HALOGENS:
                F_hal += contrib

    return float(F_metal), float(F_P_val), float(F_hal)


# ==============================
# 6. Full GPU-accelerated driver
# ==============================

def run_pipeline_gpu(
    pdb_path: str,
    out_dir: str,
    radius: float,
    grid_step: float,
    padding: float,
    xtb_bin: str,
    select_A: List[str],
    save_densities: bool,
    R_env: float,
    sasa_alpha: float,
    no_pair_contacts: bool,
    block_size: int,
    device: str,
    gpu_density_mode: str,
):
    ensure_dir(out_dir)

    traj = load_traj(pdb_path)
    residues = collect_residues_from_mdtraj(traj)
    print(f"[INFO] Residues collected: {len(residues)}")

    resid_to_sasa = compute_residue_sasa_from_traj(traj, residues)
    sasa_vals = list(resid_to_sasa.values())
    mean_sasa = float(np.mean(sasa_vals)) if len(sasa_vals) > 0 else 0.0

    del traj
    gc.collect()

    overlaps, pair_contacts = compute_residue_contacts_kdtree(
        residues,
        radius=radius,
        store_pairs=(not no_pair_contacts),
    )
    if no_pair_contacts:
        pair_contacts = {}

    meta_path = os.path.join(out_dir, "meta.json")
    export_meta(
        pdb_name=os.path.basename(pdb_path),
        radius=radius,
        residues=residues,
        overlaps=overlaps,
        pair_contacts=pair_contacts,
        resid_to_sasa=resid_to_sasa,
        out_path=meta_path,
    )

    del residues
    del resid_to_sasa
    gc.collect()

    meta = load_json(meta_path)
    res_map, overlaps_m, atom_index_to_resid, resid_to_atom_coords, resid_to_sasa2 = \
        build_residue_maps(meta)

    resid_to_grid_axes, max_nx, max_ny, max_nz = precompute_residue_grids(
        resid_to_atom_coords,
        R_env=R_env,
        grid_step=grid_step,
        padding=padding,
    )
    print(f"[INFO] Allocating GPU density workspace: ({max_nx}, {max_ny}, {max_nz}) on {device}")
    density_ws = DensityWorkspaceGPU(max_nx, max_ny, max_nz, device=device)
    log_gpu_usage("after_workspace_alloc", device)

    centers = build_residue_centers(meta)
    
    # ===== NEW: global atom SoA array ready =====
    atom_coords, atom_Z, atom_sigma, resid_to_atom_indices = build_global_atom_arrays(meta)

    A_dir = os.path.join(out_dir, "density_A_overrides")
    ensure_dir(A_dir)

    density_manifest = {
        "A_residues": [],
        "B_only_residues": [],
        "files": {},
        "modes": {},
        "params": {
            "gpu_density_mode_requested": gpu_density_mode,
            "gpu_density_vector_max_elems": int(MAX_VECTOR_ELEMENTS),
            "gpu_density_batch_max_elems": int(MAX_BATCH_ELEMENTS),
        },
        "curv_rho_failed_residues": [],
    }

    env_features: Dict[str, Dict] = {}

    res_ids = list(res_map.keys())
    n_res = len(res_ids)
    feature_dim_theory = 9
    feature_dim_sim = feature_dim_theory + 10  # 9 + 10 = 19

    Ep_theory = np.zeros((n_res, feature_dim_theory), dtype=float)
    Ep_sim = np.zeros((n_res, feature_dim_sim), dtype=float)

    curv_rho_failed_residues: List[str] = []

    for block_start in range(0, n_res, block_size):
        block_end = min(block_start + block_size, n_res)
        print(f"[INFO] GPU block {block_start}..{block_end-1} / {n_res}")
        log_gpu_usage(f"block_{block_start}_{block_end-1}_start", device)

        for idx in range(block_start, block_end):
            rid = res_ids[idx]
            res = res_map[rid]

            neighbor_ids = overlaps_m.get(rid, [])

            frag_atom_indices = build_fragment_indices(
                center_res_id=rid,
                neighbor_res_ids=neighbor_ids,
                resid_to_atom_indices=resid_to_atom_indices,
            )
            frag_atoms = None
            
            if rid in select_A:
                frag_atoms = build_fragment_atoms(res_map, rid, neighbor_ids)

            coords_res_list = resid_to_atom_coords[rid]
            coords_res = np.stack(coords_res_list, axis=0).astype(np.float32)
            ref_point = coords_res.mean(axis=0)

            xs, ys, zs = resid_to_grid_axes[rid]
            X, Y, Z, nx, ny, nz = density_ws.prepare_grid_from_axes(xs, ys, zs)
            rho_view = density_ws.view_rho(nx, ny, nz)

            dx_view = density_ws.view_dx(nx, ny, nz)
            dy_view = density_ws.view_dy(nx, ny, nz)
            dz_view = density_ws.view_dz(nx, ny, nz)
            dist2_view = density_ws.view_dist2(nx, ny, nz)
            tmp_view = density_ws.view_tmp(nx, ny, nz)

            # ----- B-mode density -----
            rho_B, eff_mode_B, fallback_B = promolecular_density_on_grid_gpu_mode_arrays(
                atom_indices=frag_atom_indices,
                atom_coords=atom_coords,
                atom_Z=atom_Z,
                atom_sigma=atom_sigma,
                X=X,
                Y=Y,
                Z=Z,
                rho_out=rho_view,
                dx_ws=dx_view,
                dy_ws=dy_view,
                dz_ws=dz_view,
                dist2_ws=dist2_view,
                tmp_ws=tmp_view,
                mode=gpu_density_mode,
                rid=rid,
            )
            rho_final = rho_B

            rhoB_path = None
            rhoA_path = None
            charges_for_rid = None
            rho_A = None
            eff_mode_A: Optional[str] = None
            fallback_A: bool = False
            xtb_success = False
            xtb_used = False
            charge_length_mismatch = False

            # ----- A-mode (xTB) -----
            if rid in select_A:
                workdir = os.path.join(A_dir, f"work_{rid.replace(':', '_')}")
                xyz_path = os.path.join(workdir, "frag.xyz")
                write_xyz(frag_atoms, xyz_path)

                charges_path = run_xtb(xyz_path, workdir, xtb_bin=xtb_bin)
                if charges_path is not None:
                    charges = parse_xtb_charges(charges_path)
                    if len(charges) == len(frag_atoms):
                        xtb_success = True
                        xtb_used = True
                        charges_for_rid = charges

                        rho_A, eff_mode_A, fallback_A = density_from_charges_on_grid_gpu_mode(
                            atoms=frag_atoms,
                            charges=charges,
                            X=X,
                            Y=Y,
                            Z=Z,
                            rho_out=rho_view,
                            dx_ws=dx_view,
                            dy_ws=dy_view,
                            dz_ws=dz_view,
                            dist2_ws=dist2_view,
                            tmp_ws=tmp_view,
                            mode=gpu_density_mode,
                            rid=rid,
                        )
                        rho_final = rho_A

                        if save_densities:
                            rhoA_path = os.path.join(
                                A_dir, f"rhoA_{rid.replace(':', '_')}.npy"
                            )
                            np.save(rhoA_path, rho_A.detach().cpu().numpy())
                        density_manifest["A_residues"].append(rid)
                    else:
                        charge_length_mismatch = True
                        xtb_success = False
                        xtb_used = False
                        charges_for_rid = None
                        print(
                            f"[WARN] xTB charges length mismatch for {rid}: "
                            f"len(charges)={len(charges)}, len(frag_atoms)={len(frag_atoms)}. "
                            "Using B-mode density only."
                        )
                        density_manifest["B_only_residues"].append(rid)
                else:
                    density_manifest["B_only_residues"].append(rid)
            else:
                density_manifest["B_only_residues"].append(rid)

            if save_densities and rhoA_path is None:
                rhoB_path = os.path.join(out_dir, f"rhoB_{rid.replace(':', '_')}.npy")
                np.save(rhoB_path, rho_B.detach().cpu().numpy())

            density_manifest["files"][rid] = {
                "rhoB": rhoB_path,
                "rhoA": rhoA_path,
            }

            density_manifest["modes"][rid] = {
                "B": {
                    "requested": gpu_density_mode,
                    "effective": eff_mode_B,
                    "fallback_from_vector": bool(fallback_B),
                },
                "A": {
                    "requested": gpu_density_mode if rid in select_A else None,
                    "effective": eff_mode_A,
                    "fallback_from_vector": bool(fallback_A) if eff_mode_A is not None else False,
                    "xtb_used": bool(xtb_used),
                    "xtb_success": bool(xtb_success),
                    "charge_length_mismatch": bool(charge_length_mismatch),
                },
            }

            # ----- Density-based features -----
            mask_view = density_ws.view_mask(nx, ny, nz)
            rho_local_view = density_ws.view_rho_local(nx, ny, nz)

            dens_feats = density_features_for_residue_gpu(
                rho=rho_final,
                X=X,
                Y=Y,
                Z=Z,
                ref_point=ref_point,
                res_atom_coords=coords_res,
                R_env=R_env,
                mask_ws=mask_view,
                rho_local_ws=rho_local_view,
                dist2_ws=dist2_view,
                tmp_ws=tmp_view,
                dx_ws=dx_view,
                dy_ws=dy_view,
                dz_ws=dz_view,
                rid=rid,
            )

            if dens_feats.get("curv_rho_failed", False):
                curv_rho_failed_residues.append(rid)

            # ----- A-mode charge features -----
            charge_feats_A = None
            if charges_for_rid is not None:
                charge_feats_A = charge_features_for_residue_from_fragment(
                    rid=rid,
                    atom_index_to_resid=atom_index_to_resid,
                    resid_to_atom_coords=resid_to_atom_coords,
                    frag_atoms=frag_atoms,
                    charges=charges_for_rid,
                )

            # ----- B-mode atom-based charge (Q_atom) -----
            partial_charge_B = compute_b_mode_charge_atom_based(
                res=res,
                rid=rid,
                resid_to_sasa2=resid_to_sasa2,
                mean_sasa=mean_sasa,
                sasa_alpha=sasa_alpha,
                env_scale_from_sasa=env_scale_from_sasa,
            )

            # ----- Field terms F^{(metal)}, F^{(P)}, F^{(hal)} -----
            F_metal, F_P_val, F_hal = compute_field_terms_for_residue(
                rid=rid,
                res_map=res_map,
                centers=centers,
                overlaps=overlaps_m,
            )

            feats = {
                "id": rid,
                "chain": res["chain"],
                "resseq": res["resseq"],
                "resname": res["resname"],
                "sasa": float(resid_to_sasa2.get(rid, 0.0)),
                "partial_charge_B": partial_charge_B,
                "F_metal": float(F_metal),
                "F_P": float(F_P_val),
                "F_hal": float(F_hal),
            }
            feats.update(dens_feats)
            if charge_feats_A is not None:
                feats.update(charge_feats_A)

            feats["gpu_density_B_requested"] = gpu_density_mode
            feats["gpu_density_B_effective"] = eff_mode_B
            feats["gpu_density_B_fallback_from_vector"] = bool(fallback_B)
            feats["gpu_density_A_requested"] = gpu_density_mode if rid in select_A else None
            feats["gpu_density_A_effective"] = eff_mode_A
            feats["gpu_density_A_fallback_from_vector"] = bool(fallback_A) if eff_mode_A is not None else False
            feats["xtb_A_used"] = bool(charge_feats_A is not None)
            feats["xtb_A_success"] = bool(xtb_success)
            feats["xtb_charge_length_mismatch"] = bool(charge_length_mismatch)

            env_features[rid] = feats

            # ----- Ep rows -----
            if "partial_charge" in feats:
                q_mean = float(feats["partial_charge"])
            else:
                q_mean = float(feats.get("partial_charge_B", 0.0))

            if "dipole_q" in feats:
                dip_vec = np.array(feats["dipole_q"], dtype=float)
            else:
                dip_vec = np.array(feats.get("dipole_rho", [0.0, 0.0, 0.0]), dtype=float)
            dip_norm = float(np.linalg.norm(dip_vec))

            curv_density = float(feats.get("curv_rho", 0.0))

            # combine field components as scalar external_field = ||(F_metal, F_P, F_hal)||
            F_vec = np.array([F_metal, F_P_val, F_hal], dtype=float)
            Efield = float(np.linalg.norm(F_vec))

            sasa = float(feats.get("sasa", 0.0))
            emb = residue_to_embedding(res["resname"]).astype(float)

            E_theory_row = np.concatenate(
                [
                    np.array([q_mean, dip_norm, curv_density, Efield, sasa], dtype=float),
                    emb,
                ]
            )

            # new 10D geometric block from overlaps / centers
            geom_block = geometric_block_from_overlaps(
                rid=rid,
                overlaps=overlaps_m,
                centers=centers,
                r_max=radius,
            )

            E_sim_row = np.concatenate([E_theory_row, geom_block])

            Ep_theory[idx, :] = E_theory_row
            Ep_sim[idx, :] = E_sim_row

            # ----- per-residue cleanup -----
            if 'frag_atom_indices' in locals():
                del frag_atom_indices
            if frag_atoms is not None:
                del frag_atoms
            del coords_res, coords_res_list, ref_point
            if rho_A is not None:
                del rho_A
            if "charges" in locals():
                del charges
                charges = None

        gc.collect()
        if torch is not None and torch.cuda.is_available():
            torch.cuda.empty_cache()
        log_gpu_usage(f"block_{block_start}_{block_end-1}_end", device)

    density_manifest["curv_rho_failed_residues"] = curv_rho_failed_residues

    dens_manifest_path = os.path.join(out_dir, "density_final.json")
    with open(dens_manifest_path, "w") as f:
        json.dump(density_manifest, f, indent=2)
    print(f"[INFO] density_final.json written: {dens_manifest_path}")

    env_path = os.path.join(out_dir, "env_features.json")
    with open(env_path, "w") as f:
        json.dump(env_features, f, indent=2)
    print(f"[INFO] env_features.json written: {env_path}")

    del res_map, overlaps_m, atom_index_to_resid, resid_to_atom_coords, resid_to_sasa2
    gc.collect()

    Ep_theory_path = os.path.join(out_dir, "Ep_theory.npy")
    Ep_sim_path = os.path.join(out_dir, "Ep_sim.npy")
    np.save(Ep_theory_path, Ep_theory)
    np.save(Ep_sim_path, Ep_sim)

    manifest = {
        "pdb": meta["pdb"],
        "n_residues": n_res,
        "residue_ids": res_ids,
        "E_theory_path": os.path.basename(Ep_theory_path),
        "E_sim_path": os.path.basename(Ep_sim_path),
        "feature_dim_theory": int(Ep_theory.shape[1]),
        "feature_dim_sim": int(Ep_sim.shape[1]),
        "features_theory": [
            "avg_charge",
            "dipole_norm",
            "curv_density",
            "external_field",  # ||F_metal, F_P, F_hal||
            "SASA",
            "emb_charge",
            "emb_hydrophobicity",
            "emb_size",
            "emb_aromaticity",
        ],
        "features_extra": [
            # 10D geometric neighbor block:
            "N_in",
            "N_out",
            "R_shell",
            "r_mean",
            "r_var",
            "linearity",
            "planarity",
            "isotropy",
            "H_mean",
            "K_gauss",
        ],
        "params": {
            "radius_contact": float(radius),
            "padding": float(padding),
            "grid_step": float(grid_step),
            "R_env": float(R_env),
            "sasa_alpha": float(sasa_alpha),
            "xtb": {
                "method": "gfn2",
                "charges": True,
                "binary": xtb_bin,
            },
            "block_size": int(block_size),
            "device": device,
            "gpu_density_mode_requested": gpu_density_mode,
            "gpu_density_vector_max_elems": int(MAX_VECTOR_ELEMENTS),
            "gpu_density_batch_max_elems": int(MAX_BATCH_ELEMENTS),
        },
    }
    mani_path = os.path.join(out_dir, "Ep_manifest.json")
    with open(mani_path, "w") as f:
        json.dump(manifest, f, indent=2)

    print(f"[DONE] E_p^theory written: {Ep_theory_path}")
    print(f"[DONE] E_p^sim written:    {Ep_sim_path}")
    print(f"[DONE] Ep manifest:        {mani_path}")


# ==============================
# 6b. Memory debug wrapper
# ==============================

def run_with_mem_monitor(
    pdb_path: str,
    out_dir: str = "pipeline_out",
    extra_args: Optional[List[str]] = None,
    python_bin: str = "python",
    interval: float = 5.0,
    max_seconds: Optional[float] = None,
):
    if not HAS_PSUTIL:
        raise RuntimeError("psutil is not available; cannot use run_with_mem_monitor.")

    if extra_args is None:
        extra_args = []

    script_path = os.path.abspath(__file__)
    cmd = [
        python_bin,
        script_path,
        pdb_path,
        "--out-dir",
        out_dir,
    ] + list(extra_args)

    print("Running:", " ".join(cmd))
    proc = subprocess.Popen(
        cmd,
        stdout=subprocess.PIPE,
        stderr=subprocess.STDOUT,
        text=True,
        bufsize=1,
    )

    p = psutil.Process(proc.pid)
    start = time.time()
    mem_log: List[Tuple[float, float]] = []

    try:
        while proc.poll() is None:
            now = time.time()
            t = now - start
            try:
                rss = p.memory_info().rss
                gb = rss / (1024 ** 3)
                mem_log.append((t, gb))
                print(f"[MEM] {t:7.1f} s : {gb:5.3f} GB")
            except psutil.Error:
                pass

            if proc.stdout is not None:
                try:
                    line = proc.stdout.readline()
                    if line:
                        print("[OUT]", line.rstrip())
                except Exception:
                    pass

            if max_seconds is not None and t > max_seconds:
                print(f"[WARN] max_seconds={max_seconds} exceeded; sending SIGINT to child...")
                try:
                    proc.send_signal(signal.SIGINT)
                except Exception:
                    pass

            time.sleep(interval)

    finally:
        if proc.poll() is None:
            try:
                proc.terminate()
            except Exception:
                pass

    print("Return code:", proc.returncode)
    return mem_log


# ==============================
# 7. CLI
# ==============================

def main():
    ap = argparse.ArgumentParser(
        description="Full 1–9 pipeline with optional GPU acceleration."
    )
    ap.add_argument("pdb", help="Input PDB or mmCIF file")
    ap.add_argument(
        "--out-dir",
        type=str,
        default="pipeline_out",
        help="Output directory for all artifacts",
    )
    ap.add_argument(
        "--radius",
        type=float,
        default=4.7,
        help="Sphere radius (Å) for residue-residue contact detection",
    )
    ap.add_argument(
        "--grid-step",
        type=float,
        default=0.8,
        help="Grid spacing (Å) for density grids",
    )
    ap.add_argument(
        "--padding",
        type=float,
        default=2.0,
        help="Padding (Å) around U_p box when building grids",
    )
    ap.add_argument(
        "--R-env",
        type=float,
        default=2.0,
        help="Environment radius (Å) for defining U_p around each residue",
    )
    ap.add_argument(
        "--sasa-alpha",
        type=float,
        default=0.4,
        help="Scaling factor alpha in SASA-based charge modulation",
    )
    ap.add_argument(
        "--xtb-bin",
        type=str,
        default="xtb",
        help="Path to xTB binary (for A-mode)",
    )
    ap.add_argument(
        "--A-list",
        type=str,
        default="",
        help='JSON list of residue IDs to run A-mode on, e.g. \'["A:201","A:175"]\'',
    )
    ap.add_argument(
        "--save-densities",
        action="store_true",
        help="If set, save rhoB_*.npy and rhoA_*.npy grids to disk",
    )
    ap.add_argument(
        "--no-pair-contacts",
        action="store_true",
        help="If set, do not store atom-level pair_contacts in meta.json.",
    )
    ap.add_argument(
        "--block-size",
        type=int,
        default=200,
        help="Number of residues per density/feature block (for streaming).",
    )
    ap.add_argument(
        "--device",
        type=str,
        default="auto",
        choices=["auto", "cpu", "cuda"],
        help="Device for heavy 3D work (auto=CUDA if available, else CPU).",
    )
    ap.add_argument(
        "--gpu-density-mode",
        type=str,
        default="safe",
        choices=["safe", "batch", "vector"],
        help=(
            "GPU density accumulation mode: "
            "safe=atom-loop, batch=atom-tiled vector kernel, "
            "vector=full broadcast with thresholded fallback."
        ),
    )

    args = ap.parse_args()

    if args.A_list:
        try:
            select_A = json.loads(args.A_list)
            if not isinstance(select_A, list):
                raise ValueError
        except Exception:
            raise SystemExit(
                "A-list must be a JSON list of residue IDs, "
                'e.g. \'["A:201\",\"A:175"]\''
            )
    else:
        select_A = []

    if args.device == "auto":
        if HAS_TORCH and torch.cuda.is_available():
            device = "cuda"
        else:
            device = "cpu"
    elif args.device == "cuda":
        if not (HAS_TORCH and torch.cuda.is_available()):
            raise SystemExit("CUDA requested but torch.cuda.is_available() is False.")
        device = "cuda"
    else:
        device = "cpu"

    if not HAS_TORCH:
        raise SystemExit("torch is required for this GPU pipeline (even on CPU device).")

    print(f"[INFO] Using device: {device}")

    run_pipeline_gpu(
        pdb_path=args.pdb,
        out_dir=args.out_dir,
        radius=args.radius,
        grid_step=args.grid_step,
        padding=args.padding,
        xtb_bin=args.xtb_bin,
        select_A=select_A,
        save_densities=args.save_densities,
        R_env=args.R_env,
        sasa_alpha=args.sasa_alpha,
        no_pair_contacts=args.no_pair_contacts,
        block_size=args.block_size,
        device=device,
        gpu_density_mode=args.gpu_density_mode,
    )


if __name__ == "__main__":
    main()
```
