## Supplementary material for "Localized Reactivity on Proteins as Riemannian Manifolds: A Quantum-Inspired Geometric Model for Deterministic, Metal-Aware Reactive-Site Prediction": code bundle: 4rubtest.py

```
#!/usr/bin/env python
"""
Rubisco 4RUB residue-level evaluation script (v4, Policy P + Solvent Filter)

Changes from previous version:
1. [Protocol] Solvent-Exclusion: Automatically removes water/solvent residues 
   using 'meta.json' before evaluation.
2. [Protocol] Policy P: Implements adaptive GroupKFold (Adaptive K) with 
   strict fold validity checks (both classes must exist in train/test).

Assumptions:
- Ep_sim.npy, Ep_manifest.json, meta.json exist in current directory.
"""

import json
import numpy as np

from sklearn.impute import SimpleImputer
from sklearn.ensemble import RandomForestClassifier
from sklearn.model_selection import StratifiedKFold, GroupKFold
from sklearn.metrics import roc_auc_score

# ============================================================
# 0. Global Settings
# ============================================================

EP_SIM_PATH = "Ep_sim.npy"
MANIFEST_PATH = "Ep_manifest.json"
META_PATH = "meta.json"  # Required for Solvent Filter

# Policy P Knobs
K_TARGET = 5
K_MIN = 2
MIN_VALID_FOLDS = 2  # At least 2 valid folds required to report mean AUC
RF_RANDOM_STATE = 42

# ============================================================
# 1. Load feature matrix and manifest
# ============================================================

print("[INFO] Loading Ep_sim.npy ...")
X = np.load(EP_SIM_PATH)
print(f"[INFO] Raw Ep_sim shape: {X.shape}")

print("[INFO] Loading Ep_manifest.json ...")
with open(MANIFEST_PATH, "r") as f:
    manifest_raw = json.load(f)

# --- normalise manifest into a list of residue entries ---

if isinstance(manifest_raw, dict):
    if "residues" in manifest_raw:
        residues_raw = manifest_raw["residues"]
    elif "residue_ids" in manifest_raw:
        residues_raw = [{"id": rid} for rid in manifest_raw["residue_ids"]]
    else:
        raise ValueError("[ERROR] Unknown manifest format.")
elif isinstance(manifest_raw, list):
    residues_raw = manifest_raw
else:
    raise ValueError("[ERROR] Manifest must be dict or list.")

res_ids = []
chains = []
resnos = []

def build_res_id_from_entry(entry):
    if isinstance(entry, str):
        return entry
    if isinstance(entry, dict):
        if "id" in entry:
            return str(entry["id"])
        # fallback logic omitted for brevity, assuming standard format
        return str(entry.get("id", "UNKNOWN"))
    raise ValueError(f"Invalid entry: {entry}")

for entry in residues_raw:
    rid = build_res_id_from_entry(entry)
    if ":" not in rid:
        # Emergency fallback if ID is just a number (unlikely)
        rid = f"A:{rid}"
    
    parts = rid.split(":")
    ch = parts[0].strip()
    rn = parts[1].strip()

    # Try to normalise residue index to integer
    try:
        rn_int = int(rn)
        rn_str = str(rn_int)
    except ValueError:
        rn_str = rn

    res_id_norm = f"{ch}:{rn_str}"
    res_ids.append(res_id_norm)
    chains.append(ch)
    resnos.append(rn_str)

res_ids = np.array(res_ids)
chains = np.array(chains)
resnos = np.array(resnos)

N_raw = len(res_ids)
if X.shape[0] != N_raw:
    raise ValueError(f"Mismatch: X({X.shape[0]}) vs Manifest({N_raw})")

# =========================================================
# [New] Solvent-Exclusion Filter (Water Removal Protocol)
# =========================================================
print("[INFO] Applying Solvent-Exclusion Filter...")

try:
    with open(META_PATH, 'r') as f:
        meta_data = json.load(f)
    
    # Map ID -> ResName (e.g. "A:100" -> "HOH")
    resname_map = {r['id']: r['resname'] for r in meta_data['residues']}
    
    valid_indices = []
    solvent_codes = ["HOH", "WAT", "TIP3", "DOD", "SOL"]
    removed_count = 0
    
    for i, rid in enumerate(res_ids):
        # res_ids are normalized "Chain:ResNo". Ensure meta.json matches this format.
        # If meta.json uses raw IDs, we might need direct mapping. 
        # Here we assume residue_ids in manifest match keys in meta.json.
        
        # Look up using the raw entry ID if possible, or constructed ID
        rname = resname_map.get(rid, "UNKNOWN")
        
        if rname not in solvent_codes:
            valid_indices.append(i)
        else:
            removed_count += 1
            
    # Apply Filter
    X = X[valid_indices]
    res_ids = res_ids[valid_indices]
    chains = chains[valid_indices]
    resnos = resnos[valid_indices]
    
    print(f"   -> Removed {removed_count} solvent residues.")
    print(f"   -> Retained {len(valid_indices)} protein residues.")

except FileNotFoundError:
    print("[WARN] meta.json not found! Proceeding WITHOUT solvent filtering.")
except Exception as e:
    print(f"[WARN] Error during solvent filtering: {e}. Proceeding with raw data.")

# ============================================================
# 2. Define site-group labels (role + residue index)
# ============================================================

L_CHAINS = {"A", "B", "C", "D"}
S_CHAINS = {"S", "T", "U", "V"}

def role_for_chain(ch):
    if ch in L_CHAINS: return "L"
    if ch in S_CHAINS: return "S"
    return ch

group_keys = []
for ch, rn in zip(chains, resnos):
    role = role_for_chain(ch)
    group_keys.append(f"{role}:{rn}")

# Integer groups for GroupKFold
unique_groups, group_indices = np.unique(group_keys, return_inverse=True)
groups = group_indices

print(f"[INFO] Unique site-groups (role+resno): {len(unique_groups)}")


# ============================================================
# 3. Rubisco 4RUB label definitions
# ============================================================
# (This section is identical to your original script)

rubisco_4RUB_labels = {
    "L_Tier1_catalytic_core": [20, 60, 65, 123, 175, 177, 201, 203, 204, 294, 295, 327, 334, 335],
    "L_Tier1_PTM_core": [201],
    "L_Tier1_gate_latch": [327, 334, 473],
    "L_Tier1_PPI_Rca_core": [1, 2, 3, 4],
    "L_Tier2_catalytic_shell": [66, 379, 380, 381, 402, 403, 404],
    "L_Tier2_PTM_shell": [104, 256],
    "L_Tier2_PPI_Rca_extension": [5, 6, 7, 8],
    "S_Tier1_helix8_cluster": [43, 73, 78, 79, 81, 92],
    "S_Tier1_interface_betaAB_core": [16, 18, 32],
    "S_Tier2_betaAB_regulatory": [59, 67, 68, 69, 71],
    "S_Tier2_helix8_shell": [19, 54, 84],
}

chains_L = ["A", "B", "C", "D"]
chains_S = ["S", "T", "U", "V"]

def expand_chain_res(label_key):
    if label_key.startswith("L_"): use_chains = chains_L
    elif label_key.startswith("S_"): use_chains = chains_S
    else: use_chains = chains_L + chains_S

    res_list = rubisco_4RUB_labels[label_key]
    out = set()
    for ch in use_chains:
        for r in res_list:
            out.add(f"{ch}:{int(r)}")
    return out

subset_pos_sets = {}
for key in rubisco_4RUB_labels.keys():
    subset_pos_sets[key] = expand_chain_res(key)

# Unions
subset_pos_sets["L_Tier1_all"] = (subset_pos_sets["L_Tier1_catalytic_core"] | subset_pos_sets["L_Tier1_PTM_core"] | subset_pos_sets["L_Tier1_gate_latch"] | subset_pos_sets["L_Tier1_PPI_Rca_core"])
subset_pos_sets["L_Tier2_all"] = (subset_pos_sets["L_Tier2_catalytic_shell"] | subset_pos_sets["L_Tier2_PTM_shell"] | subset_pos_sets["L_Tier2_PPI_Rca_extension"])
subset_pos_sets["S_Tier1_all"] = (subset_pos_sets["S_Tier1_helix8_cluster"] | subset_pos_sets["S_Tier1_interface_betaAB_core"])
subset_pos_sets["S_Tier2_all"] = (subset_pos_sets["S_Tier2_betaAB_regulatory"] | subset_pos_sets["S_Tier2_helix8_shell"])
subset_pos_sets["global_Tier1_all"] = (subset_pos_sets["L_Tier1_all"] | subset_pos_sets["S_Tier1_all"])
subset_pos_sets["global_Tier2_all"] = (subset_pos_sets["L_Tier2_all"] | subset_pos_sets["S_Tier2_all"])

# Functional-type unions
subset_pos_sets["L_Type_catalytic_T1T2"] = (subset_pos_sets["L_Tier1_catalytic_core"] | subset_pos_sets["L_Tier2_catalytic_shell"])
subset_pos_sets["L_Type_PTM_T1T2"] = (subset_pos_sets["L_Tier1_PTM_core"] | subset_pos_sets["L_Tier2_PTM_shell"])
subset_pos_sets["L_Type_PPI_Rca_T1T2"] = (subset_pos_sets["L_Tier1_PPI_Rca_core"] | subset_pos_sets["L_Tier2_PPI_Rca_extension"])
subset_pos_sets["S_Type_betaAB_T1T2"] = (subset_pos_sets["S_Tier1_interface_betaAB_core"] | subset_pos_sets["S_Tier2_betaAB_regulatory"])
subset_pos_sets["S_Type_helix8_T1T2"] = (subset_pos_sets["S_Tier1_helix8_cluster"] | subset_pos_sets["S_Tier2_helix8_shell"])

print("[INFO] Label subsets defined.")

# ============================================================
# 4. Evaluation utilities (Policy P Implementation)
# ============================================================

def eval_auc_grouped_policy_p(X, y, groups, n_splits=K_TARGET):
    """
    Policy P Logic:
    1. Check if #pos_groups is sufficient.
    2. Try K = min(n_splits, #pos_groups) down to K_MIN.
    3. Strict check: Train AND Test must have both classes (0 and 1).
    4. Return mean AUC only if >= MIN_VALID_FOLDS are successful.
    """
    pos_groups = np.unique(groups[y == 1])
    n_pos_groups = len(pos_groups)

    info = {"status": "FAIL", "n_pos_groups": n_pos_groups, "k_used": 0, "valid_folds": 0}

    if n_pos_groups < 2:
        info["reason"] = "Too few positive groups (<2)"
        return None, None, info

    k_start = min(n_splits, n_pos_groups)

    # Adaptive K loop
    for k in range(k_start, K_MIN - 1, -1):
        gkf = GroupKFold(n_splits=k)
        aucs = []
        
        for train_idx, test_idx in gkf.split(X, y, groups):
            y_train, y_test = y[train_idx], y[test_idx]

            # Strict Validity Check
            if len(np.unique(y_train)) < 2 or len(np.unique(y_test)) < 2:
                continue # Skip invalid fold

            imputer = SimpleImputer(strategy="mean")
            X_train_imp = imputer.fit_transform(X[train_idx])
            X_test_imp = imputer.transform(X[test_idx])

            clf = RandomForestClassifier(
                n_estimators=100,
                class_weight="balanced",
                random_state=RF_RANDOM_STATE,
                n_jobs=1,
            )
            clf.fit(X_train_imp, y_train)
            proba = clf.predict_proba(X_test_imp)[:, 1]
            aucs.append(roc_auc_score(y_test, proba))

        # Check if this K provided enough valid folds
        if len(aucs) >= MIN_VALID_FOLDS:
            mean_auc = float(np.mean(aucs))
            std_auc = float(np.std(aucs, ddof=1)) if len(aucs) > 1 else 0.0
            info["status"] = "OK"
            info["k_used"] = k
            info["valid_folds"] = len(aucs)
            return mean_auc, std_auc, info
        
    info["reason"] = "Insufficient valid folds even after reducing K"
    return None, None, info


def evaluate_subset(X, y, groups, subset_name):
    """
    Evaluate one subset using Policy P (Grouped) and Stratified (Residue-wise).
    """
    n_pos = int((y == 1).sum())
    n_neg = int((y == 0).sum())
    result = {
        "subset": subset_name, "n_pos": n_pos, "n_neg": n_neg,
        "auc_strat_mean": None, "auc_strat_std": None,
        "auc_group_mean": None, "auc_group_std": None,
    }

    if n_pos == 0 or n_pos == len(y):
        return result

    # 1. StratifiedKFold (Standard)
    skf = StratifiedKFold(n_splits=min(5, n_pos, n_neg), shuffle=True, random_state=42)
    aucs = []
    for train_idx, test_idx in skf.split(X, y):
        if len(np.unique(y[train_idx])) < 2 or len(np.unique(y[test_idx])) < 2: continue
        
        imputer = SimpleImputer(strategy="mean")
        X_train = imputer.fit_transform(X[train_idx])
        X_test = imputer.transform(X[test_idx])
        clf = RandomForestClassifier(n_estimators=100, class_weight="balanced", random_state=42, n_jobs=1)
        clf.fit(X_train, y[train_idx])
        aucs.append(roc_auc_score(y[test_idx], clf.predict_proba(X_test)[:, 1]))
    
    if aucs:
        result["auc_strat_mean"] = float(np.mean(aucs))
        result["auc_strat_std"] = float(np.std(aucs, ddof=1))

    # 2. GroupKFold (Policy P Applied)
    g_mean, g_std, g_info = eval_auc_grouped_policy_p(X, y, groups)
    if g_mean is not None:
        result["auc_group_mean"] = g_mean
        result["auc_group_std"] = g_std
        # print(f"  -> [Debug] {subset_name}: K={g_info['k_used']}, Folds={g_info['valid_folds']}")

    return result


# ============================================================
# 5. Run evaluation
# ============================================================

results = []
print("\n============================================================")
print("[INFO] Start evaluation for all Rubisco 4RUB subsets (Policy P)")
print("============================================================\n")

for subset_name, pos_set in subset_pos_sets.items():
    # Build y vector based on filtered res_ids
    y = np.array([1 if rid in pos_set else 0 for rid in res_ids], dtype=int)
    
    res = evaluate_subset(X, y, groups, subset_name)
    results.append(res)
    
    # Live Print
    grp_res = f"{res['auc_group_mean']:.3f}" if res['auc_group_mean'] else "NA"
    print(f"[Done] {subset_name:30s} Pos:{res['n_pos']:3d} | GroupAUC: {grp_res}")

# ============================================================
# 6. Summary printout
# ============================================================

print("\n############################################################")
print("# Summary (Rubisco 4RUB - Policy P + Solvent Filter)")
print("# subset, n_pos, n_neg, AUC_strat (mean±sd), AUC_group (mean±sd)")
print("############################################################")

for r in results:
    name = r["subset"]
    n_pos = r["n_pos"]
    n_neg = r["n_neg"]

    strat_str = f"{r['auc_strat_mean']:.3f} ± {r['auc_strat_std']:.3f}" if r['auc_strat_mean'] else "NA"
    group_str = f"{r['auc_group_mean']:.3f} ± {r['auc_group_std']:.3f}" if r['auc_group_mean'] else "NA"

    print(f"{name:30s} | {n_pos:4d} pos, {n_neg:4d} neg | Strat: {strat_str:15s} | Group: {group_str:15s}")

print("\n[INFO] Evaluation completed.")
```
