## Supplementary material for "Localized Reactivity on Proteins as Riemannian Manifolds: A Quantum-Inspired Geometric Model for Deterministic, Metal-Aware Reactive-Site Prediction": code bundle: 6Q97test.py

```
#!/usr/bin/env python
"""
6Q97 ribosome case – Tier-1 / Tier-2 / hard-negative evaluation script.

Assumptions:
- Ep_sim.npy         : (N, d) feature matrix (environment vectors)
- Ep_manifest.json   : {"residue_ids": ["CHAIN:RESNUM", ...], ...}
- Tier labels (TIER1, TIER2) and HARD_NEG are defined below.
- Evaluation protocol follows the PDF Methods (random forest, CV, ROC–AUC).

Policy P (patched here):
- Adaptive K: try K_target (default 5), reduce if not feasible
- Skip folds where train/test has single class
- Require at least min_valid_folds (default 2) valid folds; otherwise reduce K and retry
- Report CV diagnostics
"""

import json
import re
from collections import Counter

import numpy as np
from sklearn.ensemble import RandomForestClassifier
from sklearn.impute import SimpleImputer
from sklearn.metrics import roc_auc_score
from sklearn.model_selection import GroupKFold, StratifiedKFold

# ============================
# 1) Paths
# ============================

EP_SIM_PATH = "Ep_sim.npy"
MANIFEST_PATH = "Ep_manifest.json"

# ============================
# 2) Chain mapping (author -> manifest)
# ============================
CHAIN_MAP = {
    "1": "A",
    "2": "B",
    "3": "C",
    "4": "GB",
    "5": "D",
    "6": "E",
    "7": "HB",
    "8": "F",
    "9": "G",
    "B": "H",
    "C": "I",
    "D": "J",
    "E": "K",
    "F": "L",
    "G": "M",
    "H": "N",
    "I": "O",
    "J": "P",
    "K": "Q",
    "L": "R",
    "M": "S",
    "N": "T",
    "O": "U",
    "P": "V",
    "Q": "W",
    "R": "X",
    "S": "Y",
    "T": "Z",
    "U": "AA",
    "V": "BA",
    "W": "CA",
    "X": "DA",
    "Y": "EA",
    "Z": "FA",
    "a": "GA",
    "b": "HA",
    "c": "IA",
    "d": "JA",
    "e": "KA",
    "f": "LA",
    "g": "MA",
    "h": "NA",
    "i": "OA",
    "j": "PA",
    "k": "QA",
    "l": "RA",
    "m": "SA",
    "n": "TA",
    "o": "UA",
    "p": "VA",
    "q": "WA",
    "r": "XA",
    "s": "YA",
    "t": "ZA",
    "u": "AB",
    "v": "BB",
    "w": "CB",
    "x": "DB",
    "y": "EB",
    "z": "FB",
}

# ============================
# 3) Tier-1 / Tier-2 labels (author notation)
# ============================

TIER1 = {
    "SmpB_chain5": [
        "5:HIS22",
        "5:GLY132", "5:LYS133", "5:LYS134",
        "5:HIS136", "5:ASP137", "5:LYS138", "5:ARG139",
        "5:LYS143", "5:ARG145", "5:TRP147",
        "5:ARG153", "5:LYS156",
        "5:ILE154", "5:MET155",
    ],

    "tmRNA_chain4": [
        "4:U119", "4:U120", "4:A121",
        "4:C127", "4:U128", "4:G129",
        "4:U131",
        "4:C183", "4:A184", "4:A185", "4:A186",
        "4:C361", "4:C362", "4:A363",
    ],

    "16S_rRNA_chain2": [
        "2:G530", "2:A1492", "2:A1493",
        "2:G693", "2:A790", "2:G926",
        "2:C1399", "2:C1400", "2:G1401",
    ],

    "23S_rRNA_chain1_PTC_core": [
        "1:A2451", "1:U2506", "1:U2585", "1:A2602",
    ],

    "uS3_chainh": [
        "h:ARG72", "h:PRO73", "h:ILE77",
        "h:LYS79", "h:LYS80",
        "h:ARG131", "h:ARG132", "h:LYS135",
        "h:ARG136", "h:ASN140", "h:LEU144",
    ],

    "uS4_chaini": [
        "i:ARG44", "i:ARG47", "i:ARG49", "i:ARG50",
    ],

    "uS5_chainj": [
        "j:ARG20",
        "j:PHE31", "j:PHE33",
        "j:GLU55", "j:VAL56",
        "j:ILE60", "j:GLN61",
    ],
}

TIER2 = {
    "SmpB_decoding_shell_chain5_20_26": [
        "5:SER20", "5:GLY21",
        "5:THR23", "5:THR24",
        "5:LYS25", "5:ARG26",
    ],

    "SmpB_tail_shell_chain5_140_158": [
        "5:SER140", "5:ASP141", "5:ILE142",
        "5:GLU144", "5:GLU146",
        "5:GLN148", "5:VAL149",
        "5:ASP150", "5:LYS151", "5:ALA152",
        "5:ASN157", "5:ALA158",
    ],

    "tmRNA_H5_shell_chain4_115_132": [
        "4:U115", "4:G116", "4:C117", "4:A118",
        "4:A122", "4:U123", "4:G124", "4:G125",
        "4:A126", "4:U130", "4:A132",
    ],

    "tmRNA_PK2_shell_chain4_175_190": [
        "4:G175", "4:A176", "4:C177", "4:G178",
        "4:A179", "4:A180", "4:G181", "4:U182",
        "4:A187", "4:C188", "4:G189", "4:G190",
    ],

    "tmRNA_acceptor_shell_chain4_355_360": [
        "4:C355", "4:G356", "4:G357",
        "4:U358", "4:C359", "4:C360",
    ],

    "16S_decoding_shell_chain2": [
        "2:C518", "2:C1195",
    ],

    "uS3_KH2_shell_chainh_68_82": [
        "h:LYS68", "h:ARG69", "h:GLU70", "h:GLU71",
        "h:LEU74", "h:TYR75", "h:GLN76",
        "h:LYS78", "h:GLU81", "h:GLU82",
    ],

    "uS3_Cterm_shell_chainh_128_145": [
        "h:LEU128", "h:ARG129", "h:GLY130",
        "h:LYS133", "h:GLY134",
        "h:ARG137", "h:ALA138", "h:ALA139",
        "h:LYS141", "h:GLY142", "h:THR145",
    ],

    "uS4_helicase_related_chaini": [
        "i:GLU35",
        "i:ALA43",
        "i:LYS45",
    ],

    "uS5_shell_chainj_28_34": [
        "j:LYS28", "j:THR29",
        "j:GLU30", "j:GLY32", "j:ALA34",
    ],
    "uS5_shell_chainj_52_58": [
        "j:LYS52", "j:ALA53", "j:ARG54",
        "j:PRO57", "j:ALA58",
    ],

    "23S_PTC_shell_chain1": [
        "1:C2452", "1:A2601", "1:G2603",
    ],
}

# ============================
# 3.5) Role -> Tier2 shells mapping
# ============================

ROLE_TO_TIER2_KEYS = {
    "SmpB_chain5": [
        "SmpB_decoding_shell_chain5_20_26",
        "SmpB_tail_shell_chain5_140_158",
    ],
    "tmRNA_chain4": [
        "tmRNA_H5_shell_chain4_115_132",
        "tmRNA_PK2_shell_chain4_175_190",
        "tmRNA_acceptor_shell_chain4_355_360",
    ],
    "16S_rRNA_chain2": [
        "16S_decoding_shell_chain2",
    ],
    "23S_rRNA_chain1_PTC_core": [
        "23S_PTC_shell_chain1",
    ],
    "uS3_chainh": [
        "uS3_KH2_shell_chainh_68_82",
        "uS3_Cterm_shell_chainh_128_145",
    ],
    "uS4_chaini": [
        "uS4_helicase_related_chaini",
    ],
    "uS5_chainj": [
        "uS5_shell_chainj_28_34",
        "uS5_shell_chainj_52_58",
    ],
}

# ============================
# 4) Hard-negative set (author notation)  [Option A: FULL INLINE]
# ============================

HARD_NEG = [
    "1:A1689", "1:A1701", "1:A1819", "1:A705", "1:G1695",
    "1:G1702", "1:G1799", "1:U1818", "1:U1820", "2:A246",
    "B:ARG69", "B:ASN143", "B:HIS142", "B:LEU105", "B:THR191",
    "B:VAL195", "D:ARG102", "D:ARG114", "D:ARG21", "D:ARG44",
    "D:ARG49", "D:ARG61", "D:ARG67", "D:ARG79", "D:ARG88",
    "D:ASP116", "D:ASP145", "D:ASP154", "D:ASP22", "D:ASP91",
    "D:GLU111", "D:GLU122", "D:GLU127", "D:GLU152", "D:GLU16",
    "D:GLU25", "D:GLU51", "D:HIS92", "D:LYS106", "D:LYS130",
    "D:LYS132", "D:LYS139", "D:LYS47", "D:LYS57", "D:LYS58",
    "D:LYS63", "D:LYS74", "D:LYS95", "D:PHE124", "D:PHE158",
    "D:PHE19", "D:TYR35", "E:ASP10", "E:ASP6", "E:GLU11",
    "E:GLU19", "E:HIS5", "E:LYS14", "E:LYS15", "E:LYS3",
    "E:LYS9", "E:PHE20", "E:TYR22", "E:TYR7", "E:TYR8",
    "H:ARG125", "H:ARG31", "H:ARG42", "H:ARG53", "H:ARG56",
    "H:ARG61", "H:ARG94", "H:ASP124", "H:ASP29", "H:ASP36",
    "H:ASP7", "H:ASP74", "H:GLU107", "H:GLU116", "H:GLU14",
    "H:GLU17", "H:GLU47", "H:GLU65", "H:GLU87", "H:LYS101",
    "H:LYS105", "H:LYS109", "H:LYS20", "H:LYS43", "H:LYS73",
    "H:LYS8", "H:LYS97", "H:PHE106", "H:PHE69", "H:PHE76",
    "H:PHE99", "I:ARG102", "I:ASP115", "I:ASP120", "I:ASP46",
    "I:ASP63", "I:GLU107", "I:GLU122", "I:LYS44", "I:LYS50",
    "I:LYS80", "I:LYS9", "I:LYS96", "I:PHE37", "I:PHE66",
    "I:PHE68", "I:TYR7", "J:ARG120", "J:ARG27", "J:ARG34",
    "J:ARG37", "J:ARG69", "J:ARG95", "J:ARG96", "J:ARG99",
    "J:ASP14", "J:ASP19", "J:ASP49", "J:ASP52", "J:ASP60",
    "J:ASP71", "J:GLU102", "J:GLU31", "J:GLU98", "J:HIS130",
    "J:HIS132", "J:HIS40", "J:HIS47", "J:HIS76", "J:HIS77",
    "J:LYS106", "J:LYS12", "J:LYS121", "J:LYS123", "J:LYS23",
    "J:LYS39", "J:LYS61", "J:LYS68", "J:LYS85", "J:PHE119",
    "J:PHE4", "J:PHE89", "J:TYR44", "J:TYR74", "K:ALA11",
    "K:ALA83", "K:ARG108", "K:ARG30", "K:ARG31", "K:ARG49",
    "K:ARG64", "K:ARG70", "K:ARG71", "K:ARG78", "K:ASP12",
    "K:ASP80", "K:GLU106", "K:GLU110", "K:GLU4", "K:GLU45",
    "K:GLU92", "K:HIS29", "K:LEU8", "K:LYS111", "K:LYS114",
    "K:LYS40", "K:LYS44", "K:LYS53", "K:LYS54", "K:LYS59",
    "K:LYS66", "K:LYS67", "K:PHE100", "K:PHE112", "K:PHE79",
    "K:VAL63", "L:ARG123", "L:ARG132", "L:ARG18", "L:ARG21",
    "L:ARG33", "L:ARG41", "L:ARG59", "L:ARG69", "L:ARG78",
    "L:ASP91", "L:GLU10", "L:GLU115", "L:GLU136", "L:GLU144",
    "L:GLU76", "L:GLU86", "L:HIS35", "L:LYS109", "L:LYS141",
    "L:LYS17", "L:LYS29", "L:LYS63", "L:LYS70", "L:LYS96",
    "L:PHE107", "L:PHE50", "L:PHE66", "M:ARG50", "M:ARG51",
    "M:LYS123", "M:LYS71", "M:LYS8", "M:PHE68", "N:ARG46",
    "N:ARG8", "N:GLU43", "N:HIS31", "N:LYS42", "N:LYS56",
    "N:TYR94", "O:ARG102", "O:ARG81", "O:ARG9", "O:ARG94",
    "O:GLU112", "O:GLU80", "O:TYR64", "O:TYR99", "P:ARG39",
    "P:ARG72", "P:GLN75", "P:GLU27", "P:GLU34", "P:GLU68",
    "P:GLY23", "P:HIS77", "P:LYS106", "P:LYS111", "P:LYS37",
    "P:LYS96", "P:TYR99", "P:VAL70", "Q:ARG30", "Q:ARG51",
    "Q:ARG53", "Q:ARG64", "Q:ARG70", "Q:ASP49", "Q:GLU111",
    "Q:LYS114", "Q:LYS54", "Q:LYS78", "R:ARG13", "R:ARG21",
    "R:ARG78", "R:ARG84", "R:GLU70", "R:LYS48", "R:LYS60",
    "R:PHE35", "R:TYR2", "S:ARG11", "S:ARG18", "S:ARG8",
    "S:ARG84", "S:ARG92", "S:ASP34", "S:ASP62", "S:GLU59",
    "S:LYS16", "S:LYS6", "S:TYR38", "T:ARG12", "T:ARG3",
    "T:ARG69", "T:GLU4", "T:GLU42", "T:GLU5", "T:GLU56",
    "T:LYS40", "T:LYS64", "U:ARG7", "U:ARG94", "U:ASP81",
    "U:ASP9", "U:GLU10", "U:GLU88", "U:LYS17", "U:LYS33",
    "U:LYS4", "U:PHE85", "V:ARG21", "V:ARG93", "V:HIS88",
    "V:LYS14", "V:LYS53", "V:PHE56", "V:TYR31", "W:ARG11",
    "W:ARG14", "W:ARG41", "W:ARG55", "W:ASP15", "W:ASP64",
    "W:GLU70", "W:GLU85", "W:LYS19", "W:LYS78", "X:ASP60",
    "X:GLU76", "X:LYS10", "X:TYR78", "Y:ARG48", "Y:ARG7",
    "Y:ASP49", "Y:GLU8", "Y:HIS41", "Y:LYS60", "Y:LYS9",
    "Z:ASP40", "Z:LYS19", "Z:LYS6",
]

# ============================
# 5) Helper functions
# ============================

def parse_author_label_to_canonical(label: str) -> str | None:
    try:
        chain, rest = label.split(":")
    except ValueError:
        return None

    mapped_chain = CHAIN_MAP.get(chain, chain)

    m = re.search(r"(-?\d+)$", rest)
    if not m:
        return None
    resnum = int(m.group(1))
    return f"{mapped_chain}:{resnum}"


def load_data(ep_sim_path: str, manifest_path: str):
    X = np.load(ep_sim_path)
    with open(manifest_path, "r") as f:
        manifest = json.load(f)
    residue_ids = manifest["residue_ids"]
    return X, residue_ids


def build_index_map(residue_ids):
    return {rid: i for i, rid in enumerate(residue_ids)}


def mark_labels(residue_ids, tier1_dict, tier2_dict, hard_neg_list):
    N = len(residue_ids)
    rid_to_idx = build_index_map(residue_ids)

    y_T1 = np.zeros(N, dtype=int)
    y_T1T2 = np.zeros(N, dtype=int)
    y_HN = np.zeros(N, dtype=int)

    missing_T1, missing_T2, missing_HN = [], [], []

    per_role_T1 = {}
    per_role_T1T2 = {}
    per_type_T1 = {}
    per_type_T1T2 = {}

    per_tier2_mask = {}

    # Tier-1
    for role_name, labels in tier1_dict.items():
        role_mask_T1 = np.zeros(N, dtype=int)
        type_prefix = role_name.split("_")[0]

        if type_prefix not in per_type_T1:
            per_type_T1[type_prefix] = np.zeros(N, dtype=int)
        if type_prefix not in per_type_T1T2:
            per_type_T1T2[type_prefix] = np.zeros(N, dtype=int)

        for lab in labels:
            canonical = parse_author_label_to_canonical(lab)
            if canonical is None or canonical not in rid_to_idx:
                missing_T1.append((role_name, lab, canonical))
                continue
            idx = rid_to_idx[canonical]
            y_T1[idx] = 1
            y_T1T2[idx] = 1
            role_mask_T1[idx] = 1
            per_type_T1[type_prefix][idx] = 1
            per_type_T1T2[type_prefix][idx] = 1

        per_role_T1[role_name] = role_mask_T1

    # Tier-2
    for t2_name, labels in tier2_dict.items():
        t2_mask = np.zeros(N, dtype=int)
        type_prefix = t2_name.split("_")[0]
        if type_prefix not in per_type_T1T2:
            per_type_T1T2[type_prefix] = np.zeros(N, dtype=int)

        for lab in labels:
            canonical = parse_author_label_to_canonical(lab)
            if canonical is None or canonical not in rid_to_idx:
                missing_T2.append((t2_name, lab, canonical))
                continue
            idx = rid_to_idx[canonical]
            if y_T1[idx] == 0:
                y_T1T2[idx] = 1
            t2_mask[idx] = 1
            per_type_T1T2[type_prefix][idx] = 1

        per_tier2_mask[t2_name] = t2_mask

    # Role-level Tier1+2
    for role_name, role_mask_T1 in per_role_T1.items():
        agg_mask = role_mask_T1.copy()
        if role_name in ROLE_TO_TIER2_KEYS:
            for t2_key in ROLE_TO_TIER2_KEYS[role_name]:
                if t2_key not in per_tier2_mask:
                    print(f"[WARN] Role {role_name}: Tier2 key '{t2_key}' not found.")
                    continue
                agg_mask |= per_tier2_mask[t2_key]
        per_role_T1T2[role_name] = agg_mask

    # Hard negatives
    for lab in hard_neg_list:
        canonical = parse_author_label_to_canonical(lab)
        if canonical is None or canonical not in rid_to_idx:
            missing_HN.append((lab, canonical))
            continue
        idx = rid_to_idx[canonical]
        y_HN[idx] = 1

    # Overlap warning (do NOT silently “remove”; just warn)
    pos_mask_any = (y_T1T2 == 1)
    overlap = np.where((y_HN == 1) & pos_mask_any)[0]
    if overlap.size > 0:
        print(f"[WARN] {overlap.size} HN labels overlap with Tier positives (label conflict).")

    # Summary
    print("=== Label summary ===")
    print(f"Total residues         : {N}")
    print(f"Tier-1 positives       : {int(y_T1.sum())}")
    print(f"Tier-1+2 positives     : {int(y_T1T2.sum())}")
    print(f"Hard negatives (HN)    : {int(y_HN.sum())}")

    if missing_T1:
        print(f"[WARN] {len(missing_T1)} Tier-1 labels not found in manifest.")
    if missing_T2:
        print(f"[WARN] {len(missing_T2)} Tier-2 labels not found in manifest.")
    if missing_HN:
        print(f"[WARN] {len(missing_HN)} hard-negative labels not found in manifest.")

    return (
        y_T1,
        y_T1T2,
        y_HN,
        per_role_T1,
        per_role_T1T2,
        per_type_T1,
        per_type_T1T2,
    )

# ============================
# 5.5) Policy P CV engine
# ============================

def _needs_groupkfold(y, groups) -> bool:
    """Use GroupKFold only if positives contain replicated group labels."""
    y = np.asarray(y, dtype=int)
    groups = np.asarray(groups, dtype=object)
    pos_groups = groups[y == 1]
    if pos_groups.size == 0:
        return False
    c = Counter(pos_groups)
    return any(v > 1 for v in c.values())


def eval_auc_policy_p(
    X,
    y,
    groups,
    label_name="",
    K_target=5,
    K_min=2,
    min_valid_folds=2,
    random_state=42,
):
    """
    Policy P:
    - Decide GroupKFold vs StratifiedKFold (based on replicated positive groups).
    - Try K = min(K_target, feasible) down to K_min.
    - Skip folds with single-class train or test.
    - Require >= min_valid_folds valid folds, else reduce K and retry.
    """
    X = np.asarray(X)
    y = np.asarray(y, dtype=int)
    groups = np.asarray(groups, dtype=object)

    n_pos = int((y == 1).sum())
    n_neg = int((y == 0).sum())
    if n_pos == 0 or n_neg == 0:
        return np.nan, np.nan, [], {
            "K_target": K_target,
            "K_min": K_min,
            "min_valid_folds": min_valid_folds,
            "reason": "no_pos_or_no_neg",
            "n_pos": n_pos,
            "n_neg": n_neg,
        }

    use_groupk = _needs_groupkfold(y, groups)

    if use_groupk:
        pos_group_counts = Counter(groups[y == 1])
        n_pos_groups = len(pos_group_counts)
        K_start = min(K_target, n_pos_groups)
    else:
        n_pos_groups = None
        K_start = min(K_target, n_pos, n_neg)

    if K_start < 2:
        return np.nan, np.nan, [], {
            "K_target": K_target,
            "K_min": K_min,
            "min_valid_folds": min_valid_folds,
            "reason": "K_start_lt_2",
            "n_pos": n_pos,
            "n_neg": n_neg,
            "n_pos_groups": n_pos_groups,
        }

    imputer = SimpleImputer(strategy="mean")

    for K in range(K_start, K_min - 1, -1):
        if use_groupk:
            splitter = GroupKFold(n_splits=K)
            split_iter = splitter.split(X, y, groups=groups)
        else:
            splitter = StratifiedKFold(n_splits=K, shuffle=True, random_state=random_state)
            split_iter = splitter.split(X, y)

        aucs = []
        n_total = 0
        n_skip_train = 0
        n_skip_test = 0

        for train_idx, test_idx in split_iter:
            n_total += 1
            y_train = y[train_idx]
            y_test = y[test_idx]

            if len(np.unique(y_train)) < 2:
                n_skip_train += 1
                continue
            if len(np.unique(y_test)) < 2:
                n_skip_test += 1
                continue

            X_train_imp = imputer.fit_transform(X[train_idx])
            X_test_imp = imputer.transform(X[test_idx])

            clf = RandomForestClassifier(
                n_estimators=100,
                class_weight="balanced",
                random_state=random_state,
                n_jobs=1,
            )
            clf.fit(X_train_imp, y_train)
            proba = clf.predict_proba(X_test_imp)[:, 1]
            aucs.append(roc_auc_score(y_test, proba))

        cv_info = {
            "K_target": K_target,
            "K_min": K_min,
            "min_valid_folds": min_valid_folds,
            "use_groupkfold": use_groupk,
            "n_pos": n_pos,
            "n_neg": n_neg,
            "n_pos_groups": n_pos_groups,
            "K_used": K,
            "n_total_folds": n_total,
            "n_valid_folds": len(aucs),
            "n_skipped_train_single_class": n_skip_train,
            "n_skipped_test_single_class": n_skip_test,
        }

        if len(aucs) >= min_valid_folds:
            return float(np.mean(aucs)), float(np.std(aucs)), aucs, {**cv_info, "reason": "OK"}

    return np.nan, np.nan, [], {
        "K_target": K_target,
        "K_min": K_min,
        "min_valid_folds": min_valid_folds,
        "use_groupkfold": use_groupk,
        "n_pos": n_pos,
        "n_neg": n_neg,
        "n_pos_groups": n_pos_groups,
        "reason": "insufficient_valid_folds_after_adaptation",
    }


def cross_validated_auc_policy_p(X, y, groups, label_name, K_target=5):
    """
    Wrapper that prints Policy P CV diagnostics and returns a dict like your previous output style.
    """
    auc_mean, auc_std, aucs, cv_info = eval_auc_policy_p(
        X, y, groups, label_name=label_name, K_target=K_target
    )

    print(f"\n=== {label_name} ===")
    print(f"AUC = {auc_mean} +/- {auc_std}")
    print(f"CV info: {cv_info}")
    print(f"fold AUCs: {aucs}")

    return {
        "auc_residue_mean": auc_mean,
        "auc_residue_std": auc_std,
        "auc_site_mean": auc_mean,   # residue == site in this script
        "auc_site_std": auc_std,
        "fold_aucs": aucs,
        "cv_info": cv_info,
    }


# ============================
# 6) Main evaluation
# ============================

def main():
    X, residue_ids = load_data(EP_SIM_PATH, MANIFEST_PATH)
    N, d = X.shape
    print("=== Data ===")
    print(f"Feature matrix shape: {X.shape}")
    print(f"# residues          : {N}")

    (
        y_T1,
        y_T1T2,
        y_HN,
        per_role_T1,
        per_role_T1T2,
        per_type_T1,
        per_type_T1T2,
    ) = mark_labels(residue_ids, TIER1, TIER2, HARD_NEG)

    # Global groups: if you want true site-grouping beyond residue-id uniqueness,
    # you must define a coarser grouping rule. Current: each residue is its own group.
    groups_global = np.array([f"G:{rid}" for rid in residue_ids], dtype=object)

    results = {}

    # ---- Global metrics ----
    results["Tier1_all"] = cross_validated_auc_policy_p(
        X, y_T1, groups_global, label_name="Tier1_all", K_target=5
    )

    results["Tier1_plus_Tier2_all"] = cross_validated_auc_policy_p(
        X, y_T1T2, groups_global, label_name="Tier1_plus_Tier2_all", K_target=5
    )

    # Tier1 vs HardNeg (only keep samples in {Tier1 union HardNeg})
    mask_T1_HN = ((y_T1 == 1) | (y_HN == 1))
    if mask_T1_HN.sum() > 0:
        y_T1_vs_HN = (y_T1[mask_T1_HN] == 1).astype(int)
        X_T1_vs_HN = X[mask_T1_HN]
        groups_T1_vs_HN = groups_global[mask_T1_HN]
        results["Tier1_vs_hard_negative"] = cross_validated_auc_policy_p(
            X_T1_vs_HN, y_T1_vs_HN, groups_T1_vs_HN,
            label_name="Tier1_vs_hard_negative", K_target=5
        )
    else:
        print("[INFO] Tier1_vs_hard_negative: no samples; skip.")

    # Tier1+2 vs HardNeg
    mask_T1T2_HN = ((y_T1T2 == 1) | (y_HN == 1))
    if mask_T1T2_HN.sum() > 0:
        y_T1T2_vs_HN = (y_T1T2[mask_T1T2_HN] == 1).astype(int)
        X_T1T2_vs_HN = X[mask_T1T2_HN]
        groups_T1T2_vs_HN = groups_global[mask_T1T2_HN]
        results["Tier1plus2_vs_hard_negative"] = cross_validated_auc_policy_p(
            X_T1T2_vs_HN, y_T1T2_vs_HN, groups_T1T2_vs_HN,
            label_name="Tier1plus2_vs_hard_negative", K_target=5
        )
    else:
        print("[INFO] Tier1plus2_vs_hard_negative: no samples; skip.")

    # ---- Role-level metrics ----
    for role_name, role_mask_T1 in per_role_T1.items():
        if role_mask_T1.sum() > 0:
            results[f"Role_Tier1_{role_name}"] = cross_validated_auc_policy_p(
                X, role_mask_T1, groups_global,
                label_name=f"Role_Tier1_{role_name}", K_target=5
            )

        role_mask_T1T2 = per_role_T1T2[role_name]
        if role_mask_T1T2.sum() > 0:
            results[f"Role_Tier1plus2_{role_name}"] = cross_validated_auc_policy_p(
                X, role_mask_T1T2, groups_global,
                label_name=f"Role_Tier1plus2_{role_name}", K_target=5
            )

    # ---- Type-level metrics ----
    for type_prefix, type_mask_T1 in per_type_T1.items():
        if type_mask_T1.sum() > 0:
            results[f"Type_{type_prefix}_Tier1"] = cross_validated_auc_policy_p(
                X, type_mask_T1, groups_global,
                label_name=f"Type_{type_prefix}_Tier1", K_target=5
            )

        type_mask_T1T2 = per_type_T1T2.get(type_prefix, np.zeros_like(type_mask_T1))
        if type_mask_T1T2.sum() > 0:
            results[f"Type_{type_prefix}_Tier1plus2"] = cross_validated_auc_policy_p(
                X, type_mask_T1T2, groups_global,
                label_name=f"Type_{type_prefix}_Tier1plus2", K_target=5
            )

    # Optional: dump results
    # with open("ribosome_auc_results.json", "w") as f:
    #     json.dump(results, f, indent=2)


if __name__ == "__main__":
    main()
```
