## Supplementary material for "Localized Reactivity on Proteins as Riemannian Manifolds: A Quantum-Inspired Geometric Model for Deterministic, Metal-Aware Reactive-Site Prediction": code bundle: benchmarkQQQQ.py

```
#!/usr/bin/env python
# -*- coding: utf-8 -*-

# ==============================================================================
# 0) USER ENGINE CODE (PASTE HERE)
#
# Replace the single "X" line below with the FULL content of your engine script
# (e.g. the original protein_env_full_pipeline_gpu.py).
#
# Requirements:
#   - Your script must accept:
#       --input <PDB_OR_CIF_PATH>
#       --out-dir <OUTPUT_DIR>
#     and produce at least:
#       <OUTPUT_DIR>/Ep_sim.npy
#       <OUTPUT_DIR>/Ep_manifest.json  (with "residue_ids": ["A:100", ...])
#
#   - Docstrings with """ are fine. Avoid using ''' inside your engine,
#     because this block is wrapped in ''' ... '''.
#
#   - Do NOT leave it as just "X". If USER_ENGINE_CODE still contains "X",
#     the script will raise an error.
# ==============================================================================

USER_ENGINE_CODE = r'''
#!/usr/bin/env python
"""
protein_env_full_pipeline_gpu.py

Full pipeline (block-streaming version) with optional GPU acceleration
and explicit tracking of effective GPU density modes per residue.

Steps
-----
1–2 : load structure, build residue list, KDTree-based contacts
      -> meta.json (residues + overlaps + pair_contacts + SASA)
3–4 : hybrid density:
        - B: promolecular surrogate density for all residues
        - A: optional xTB-based correction for a chosen subset of residues
5–6 : per-residue features (env_features.json):
        - density-based: total_density, centroid, dipole_rho, curv_rho
        - charge-based:
            * B-mode: surrogate partial_charge_B (atom-based + SASA scaling)
            * A-mode: partial_charge, dipole_q (if xTB run)
        - SASA
        - field terms: F_metal, F_P, F_hal
7–9 : environment vectors Ep:
        - Ep_theory.npy : [avg_charge, dipole_norm, curv_density,
                          external_field(=||F_metal,F_P,F_hal||),
                          SASA, residue_embedding(4)]
        - Ep_sim.npy    : Ep_theory plus 10 neighbor-geometry features
        - Ep_manifest.json : feature names, residue ids, params, etc.
"""

import os
import json
import argparse
import subprocess
from typing import Dict, List, Tuple, Optional

import numpy as np
import gc
import time
import signal

# --------------------------
# Optional deps
# --------------------------
try:
    from scipy.spatial import cKDTree
    HAS_KDTREE = True
except ImportError:
    HAS_KDTREE = False
    print("[WARN] scipy.spatial.cKDTree not available; falling back to O(N^2) contacts.")

try:
    import mdtraj as mdt
    HAS_MDTRAJ = True
except ImportError:
    HAS_MDTRAJ = False
    print("[ERROR] mdtraj is required for this pipeline.")
    raise SystemExit("mdtraj is required for this pipeline.")

try:
    import torch
    HAS_TORCH = True
except ImportError:
    HAS_TORCH = False
    torch = None
    print("[ERROR] torch is required for this GPU pipeline.")
    raise SystemExit("torch is required for this GPU pipeline.")

try:
    import psutil
    HAS_PSUTIL = True
except ImportError:
    HAS_PSUTIL = False
    psutil = None
    print("[WARN] psutil not available; memory debug helper disabled.")


# =====================================
# Global thresholds for vector GPU mode
# =====================================

MAX_VECTOR_ELEMENTS = 200_000_000
MAX_BATCH_ELEMENTS = 50_000_000


# ==============================
# 0. Utilities
# ==============================

def ensure_dir(path: str):
    os.makedirs(path, exist_ok=True)


def load_json(path: str) -> Dict:
    with open(path, "r") as f:
        return json.load(f)


def log_gpu_usage(tag: str, device: str):
    if not (HAS_TORCH and device == "cuda"):
        return
    if not torch.cuda.is_available():
        return
    try:
        torch.cuda.synchronize()
    except Exception:
        pass
    try:
        alloc = torch.cuda.memory_allocated() / 1024**2
        reserved = torch.cuda.memory_reserved() / 1024**2
        max_alloc = torch.cuda.max_memory_allocated() / 1024**2
    except Exception:
        return
    print(
        f"[GPU] {tag:24s} | "
        f"alloc={alloc:7.1f} MB | "
        f"reserved={reserved:7.1f} MB | "
        f"max={max_alloc:7.1f} MB"
    )


# ==============================
# 1. mdtraj helpers
# ==============================

def get_chain_id(chain) -> str:
    if hasattr(chain, "chain_id") and chain.chain_id is not None:
        return str(chain.chain_id)
    if hasattr(chain, "id") and chain.id is not None:
        return str(chain.id)
    return str(chain.index)


def get_insertion_code(res) -> str:
    raw = None
    if hasattr(res, "insertion_code"):
        raw = res.insertion_code
    elif hasattr(res, "insertionCode"):
        raw = res.insertionCode
    if raw is None:
        raw = ""
    icode = str(raw).strip()
    if icode == "None":
        icode = ""
    return icode


def load_traj(path: str):
    if not HAS_MDTRAJ:
        raise RuntimeError("mdtraj is required for load_traj.")
    ext = os.path.splitext(path)[1].lower()

    if ext in (".cif", ".mmcif"):
        print("[INFO] Detected CIF/mmCIF input → calling mdt.load without standard_names kwarg.")
        try:
            return mdt.load(path)
        except Exception as e:
            msg = str(e)
            print("[ERROR] mdtraj.load(cif) failed.")
            print("        First 200 chars of the error message:")
            print("        ", msg[:200].replace("\n", " "))
            raise

    try:
        return mdt.load(path, standard_names=True)
    except Exception as e:
        msg = str(e)
        print("[WARN] mdtraj.load(..., standard_names=True) failed:")
        print("       ", msg[:200].replace("\n", " "))
        print("[WARN] Retrying with standard_names=False (no canonicalization).")
        return mdt.load(path, standard_names=False)


def collect_residues_from_mdtraj(traj) -> List[Dict]:
    residues: List[Dict] = []
    topo = traj.topology
    coords0 = traj.xyz[0].astype(np.float32)

    for res in topo.residues:
        chain_id = get_chain_id(res.chain)
        resseq = res.resSeq
        icode = get_insertion_code(res)
        suffix = icode if icode else ""
        rid = f"{chain_id}:{resseq}{suffix}"

        atoms: List[Dict] = []
        for atom in res.atoms:
            idx = atom.index
            coord = np.array(coords0[idx, :], dtype=np.float32)
            if atom.element is not None:
                element = atom.element.symbol.upper()
            else:
                element = "?"
            atoms.append(
                {
                    "atom_index": idx,
                    "name": atom.name,
                    "element": element,
                    "coord": coord,
                }
            )

        if not atoms:
            continue

        residues.append(
            {
                "id": rid,
                "chain": chain_id,
                "resseq": resseq,
                "icode": icode,
                "resname": res.name,
                "hetflag": " ",
                "atoms": atoms,
            }
        )
    return residues


def compute_residue_sasa_from_traj(traj, residues: List[Dict]) -> Dict[str, float]:
    resid_to_sasa: Dict[str, float] = {r["id"]: 0.0 for r in residues}
    if not HAS_MDTRAJ:
        return resid_to_sasa

    try:
        sasa_res = mdt.shrake_rupley(traj, probe_radius=1.4, mode="residue")
        sasa_0 = sasa_res[0]
    except Exception as e:
        print("[WARN] mdtraj SASA (residue mode) failed:", e)
        return resid_to_sasa

    topo = traj.topology
    if topo.n_residues != sasa_0.shape[0]:
        print("[WARN] Topology residue count and SASA length mismatch.")
        return resid_to_sasa

    topo_res_sasa: Dict[Tuple[str, int, str], float] = {}
    for i, res in enumerate(topo.residues):
        chain_id = get_chain_id(res.chain)
        resseq = res.resSeq
        icode = get_insertion_code(res)
        key = (chain_id, int(resseq), icode)
        topo_res_sasa[key] = float(sasa_0[i])

    for r in residues:
        key = (str(r["chain"]), int(r["resseq"]), str(r["icode"]))
        if key in topo_res_sasa:
            resid_to_sasa[r["id"]] = topo_res_sasa[key]

    return resid_to_sasa


# ==============================
# 2. KDTree-based residue contacts
# ==============================

def make_pair_key(id_a: str, id_b: str) -> str:
    if id_a <= id_b:
        return f"{id_a}__{id_b}"
    else:
        return f"{id_b}__{id_a}"


def compute_residue_contacts_kdtree(
    residues: List[Dict],
    radius: float,
    store_pairs: bool = True,
) -> Tuple[Dict[str, List[str]], Dict[str, List[Dict]]]:
    atom_coords = []
    atom_res_idx = []
    res_ids = [r["id"] for r in residues]

    for i_res, r in enumerate(residues):
        for a in r["atoms"]:
            atom_coords.append(a["coord"])
            atom_res_idx.append(i_res)

    atom_coords = np.array(atom_coords, dtype=np.float32)
    atom_res_idx = np.array(atom_res_idx, dtype=int)
    n_atoms = atom_coords.shape[0]

    if store_pairs:
        overlaps: Dict[str, List[str]] = {rid: [] for rid in res_ids}
        pair_contacts: Dict[str, List[Dict]] = {}
    else:
        overlaps: Dict[str, set] = {rid: set() for rid in res_ids}
        pair_contacts: Dict[str, List[Dict]] = {}

    cutoff = 2.0 * radius

    if HAS_KDTREE:
        tree = cKDTree(atom_coords)
        for i in range(n_atoms):
            coord_i = atom_coords[i]
            res_i = atom_res_idx[i]
            rid_i = res_ids[res_i]

            idxs = tree.query_ball_point(coord_i, cutoff)
            for j in idxs:
                if j <= i:
                    continue
                res_j = atom_res_idx[j]
                if res_j == res_i:
                    continue
                rid_j = res_ids[res_j]
                d = float(np.linalg.norm(atom_coords[j] - coord_i))
                key = make_pair_key(rid_i, rid_j)

                if store_pairs:
                    if key not in pair_contacts:
                        pair_contacts[key] = []
                    pair_contacts[key].append(
                        {
                            "a_index": int(i),
                            "b_index": int(j),
                            "distance": d,
                        }
                    )

                if store_pairs:
                    overlaps[rid_i].append(rid_j)
                    overlaps[rid_j].append(rid_i)
                else:
                    overlaps[rid_i].add(rid_j)
                    overlaps[rid_j].add(rid_i)
    else:
        for i in range(n_atoms):
            coord_i = atom_coords[i]
            res_i = atom_res_idx[i]
            rid_i = res_ids[res_i]
            for j in range(i + 1, n_atoms):
                coord_j = atom_coords[j]
                res_j = atom_res_idx[j]
                if res_j == res_i:
                    continue
                d = float(np.linalg.norm(coord_j - coord_i))
                if d >= cutoff:
                    continue
                rid_j = res_ids[res_j]
                key = make_pair_key(rid_i, rid_j)

                if store_pairs:
                    if key not in pair_contacts:
                        pair_contacts[key] = []
                    pair_contacts[key].append(
                        {
                            "a_index": int(i),
                            "b_index": int(j),
                            "distance": d,
                        }
                    )

                if store_pairs:
                    overlaps[rid_i].append(rid_j)
                    overlaps[rid_j].append(rid_i)
                else:
                    overlaps[rid_i].add(rid_j)
                    overlaps[rid_j].add(rid_i)

    if store_pairs:
        for rid in overlaps:
            overlaps[rid] = sorted(set(overlaps[rid]))
    else:
        for rid in overlaps:
            overlaps[rid] = sorted(overlaps[rid])

    return overlaps, pair_contacts


def export_meta(
    pdb_name: str,
    radius: float,
    residues: List[Dict],
    overlaps: Dict[str, List[str]],
    pair_contacts: Dict[str, List[Dict]],
    resid_to_sasa: Dict[str, float],
    out_path: str,
):
    json_residues = []
    for r in residues:
        rid = r["id"]
        json_residues.append(
            {
                "id": rid,
                "chain": r["chain"],
                "resseq": r["resseq"],
                "icode": r["icode"],
                "resname": r["resname"],
                "hetflag": r["hetflag"],
                "sasa": float(resid_to_sasa.get(rid, 0.0)),
                "atoms": [
                    {
                        "atom_index": a["atom_index"],
                        "name": a["name"],
                        "element": a["element"],
                        "coord": np.array(a["coord"], dtype=float).tolist(),
                    }
                    for a in r["atoms"]
                ],
            }
        )

    meta = {
        "pdb": pdb_name,
        "radius": radius,
        "residues": json_residues,
        "overlaps": overlaps,
        "pair_contacts": pair_contacts,
    }

    with open(out_path, "w") as f:
        json.dump(meta, f, indent=2)

    print(f"[INFO] meta.json written: {out_path}")


# ==============================
# 3. Hybrid density core
# ==============================

Z_TABLE = {
    "H": 1,
    "C": 6,
    "N": 7,
    "O": 8,
    "S": 16,
    "P": 15,
    "F": 9,
    "CL": 17,
    "BR": 35,
    "I": 53,
    "ZN": 30,
    "MG": 12,
    "CA": 20,
    "FE": 26,
    "MN": 25,
    "CU": 29,
    "NI": 28,
    "CO": 27,
    "MO": 42,
    "W": 74,
    "V": 23,
}

SIGMA_TABLE = {
    "H": 0.7,
    "C": 1.0,
    "N": 0.95,
    "O": 0.9,
    "S": 1.2,
    "P": 1.2,
    "ZN": 1.1,
    "MG": 1.1,
    "CA": 1.3,
    "FE": 1.2,
    "MN": 1.2,
    "CU": 1.2,
    "NI": 1.2,
    "CO": 1.2,
}


def z_of(elem: str) -> int:
    return Z_TABLE.get(elem.upper(), 6)


def sigma_of(elem: str) -> float:
    return SIGMA_TABLE.get(elem.upper(), 1.0)


def build_residue_maps(meta: Dict):
    res_map = {r["id"]: r for r in meta["residues"]}
    overlaps = meta["overlaps"]

    atom_index_to_resid: Dict[int, str] = {}
    resid_to_atom_coords: Dict[str, List[np.ndarray]] = {}
    resid_to_sasa: Dict[str, float] = {}

    for r in meta["residues"]:
        rid = r["id"]
        coords = []
        for a in r["atoms"]:
            idx = int(a["atom_index"])
            atom_index_to_resid[idx] = rid
            coords.append(np.array(a["coord"], dtype=np.float32))
        resid_to_atom_coords[rid] = coords
        resid_to_sasa[rid] = float(r.get("sasa", 0.0))

    return res_map, overlaps, atom_index_to_resid, resid_to_atom_coords, resid_to_sasa


def build_global_atom_arrays(meta: Dict):
    """
    AoS(atom dict) → SoA(parallel arrays).

    Returns
    -------
    atom_coords : (N, 3) float32
    atom_Z      : (N,)   float32  (promolecular Z or surrogate weight)
    atom_sigma  : (N,)   float32  (Gaussian sigma per atom)
    resid_to_atom_indices : dict[rid] -> np.ndarray of atom indices
    """
    coords_list = []
    Z_list = []
    sigma_list = []
    resid_to_atom_indices: Dict[str, List[int]] = {}

    idx = 0
    for r in meta["residues"]:
        rid = r["id"]
        resid_to_atom_indices.setdefault(rid, [])
        for a in r["atoms"]:
            coord = np.asarray(a["coord"], dtype=np.float32)
            elem = str(a.get("element", "C")).strip().upper()
            coords_list.append(coord)
            Z_list.append(float(z_of(elem)))
            sigma_list.append(float(sigma_of(elem)))
            resid_to_atom_indices[rid].append(idx)
            idx += 1

    if len(coords_list) == 0:
        atom_coords = np.zeros((0, 3), dtype=np.float32)
        atom_Z = np.zeros((0,), dtype=np.float32)
        atom_sigma = np.zeros((0,), dtype=np.float32)
    else:
        atom_coords = np.stack(coords_list, axis=0).astype(np.float32)
        atom_Z = np.asarray(Z_list, dtype=np.float32)
        atom_sigma = np.asarray(sigma_list, dtype=np.float32)

    resid_to_atom_indices_np: Dict[str, np.ndarray] = {}
    for rid, idxs in resid_to_atom_indices.items():
        if len(idxs) == 0:
            resid_to_atom_indices_np[rid] = np.zeros((0,), dtype=np.int64)
        else:
            resid_to_atom_indices_np[rid] = np.asarray(idxs, dtype=np.int64)

    return atom_coords, atom_Z, atom_sigma, resid_to_atom_indices_np


def build_fragment_indices(
    center_res_id: str,
    neighbor_res_ids: List[str],
    resid_to_atom_indices: Dict[str, np.ndarray],
) -> np.ndarray:
    idx_arrays = []

    if center_res_id in resid_to_atom_indices:
        idx_arrays.append(resid_to_atom_indices[center_res_id])

    for nid in neighbor_res_ids:
        arr = resid_to_atom_indices.get(nid, None)
        if arr is not None and arr.size > 0:
            idx_arrays.append(arr)

    if not idx_arrays:
        return np.zeros((0,), dtype=np.int64)

    concat = np.concatenate(idx_arrays, axis=0)
    if concat.size == 0:
        return concat
    return np.unique(concat)


def build_fragment_atoms(
    res_map: Dict[str, Dict], center_res_id: str, neighbor_res_ids: List[str]
) -> List[Dict]:
    ids = [center_res_id] + neighbor_res_ids
    atoms: List[Dict] = []
    for rid in ids:
        if rid in res_map:
            atoms.extend(res_map[rid]["atoms"])
    return atoms


def precompute_residue_grids(
    resid_to_atom_coords: Dict[str, List[np.ndarray]],
    R_env: float,
    grid_step: float,
    padding: float,
) -> Tuple[Dict[str, Tuple[np.ndarray, np.ndarray, np.ndarray]], int, int, int]:
    resid_to_grid_axes: Dict[str, Tuple[np.ndarray, np.ndarray, np.ndarray]] = {}
    max_nx = 1
    max_ny = 1
    max_nz = 1

    extra = float(R_env + padding)

    for rid, coords_list in resid_to_atom_coords.items():
        if not coords_list:
            continue
        coords = np.stack(coords_list, axis=0).astype(np.float32)
        mn = coords.min(axis=0) - extra
        mx = coords.max(axis=0) + extra

        xs = np.arange(mn[0], mx[0] + grid_step, grid_step, dtype=np.float32)
        ys = np.arange(mn[1], mx[1] + grid_step, grid_step, dtype=np.float32)
        zs = np.arange(mn[2], mx[2] + grid_step, grid_step, dtype=np.float32)

        nx = xs.size
        ny = ys.size
        nz = zs.size

        resid_to_grid_axes[rid] = (xs, ys, zs)

        if nx > max_nx:
            max_nx = nx
        if ny > max_ny:
            max_ny = ny
        if nz > max_nz:
            max_nz = nz

    return resid_to_grid_axes, max_nx, max_ny, max_nz


# ==============================
# 3c. GPU density workspace
# ==============================

class DensityWorkspaceGPU:
    def __init__(self, max_nx: int, max_ny: int, max_nz: int, device: str):
        if not HAS_TORCH:
            raise RuntimeError("DensityWorkspaceGPU requires torch.")
        self.max_nx = int(max_nx)
        self.max_ny = int(max_ny)
        self.max_nz = int(max_nz)
        self.device = device

        shape = (self.max_nx, self.max_ny, self.max_nz)

        self.X = torch.empty(shape, dtype=torch.float32, device=self.device)
        self.Y = torch.empty_like(self.X)
        self.Z = torch.empty_like(self.X)

        self.rho = torch.empty_like(self.X)
        self.rho_local = torch.empty_like(self.X)

        self.mask = torch.empty(shape, dtype=torch.bool, device=self.device)

        self.dist2 = torch.empty_like(self.X)
        self.dx = torch.empty_like(self.X)
        self.dy = torch.empty_like(self.X)
        self.dz = torch.empty_like(self.X)
        self.tmp = torch.empty_like(self.X)

    def prepare_grid_from_axes(
        self,
        xs: np.ndarray,
        ys: np.ndarray,
        zs: np.ndarray,
    ) -> Tuple[torch.Tensor, torch.Tensor, torch.Tensor, int, int, int]:
        xs = np.asarray(xs, dtype=np.float32)
        ys = np.asarray(ys, dtype=np.float32)
        zs = np.asarray(zs, dtype=np.float32)

        nx = xs.size
        ny = ys.size
        nz = zs.size

        if nx > self.max_nx or ny > self.max_ny or nz > self.max_nz:
            raise ValueError(
                f"Workspace too small for residue grid: ({nx},{ny},{nz}) "
                f"vs ({self.max_nx},{self.max_ny},{self.max_nz})"
            )

        xs_t = torch.from_numpy(xs).to(self.device)
        ys_t = torch.from_numpy(ys).to(self.device)
        zs_t = torch.from_numpy(zs).to(self.device)

        X = self.X[:nx, :ny, :nz]
        Y = self.Y[:nx, :ny, :nz]
        Z = self.Z[:nx, :ny, :nz]

        X[:] = xs_t.view(-1, 1, 1)
        Y[:] = ys_t.view(1, -1, 1)
        Z[:] = zs_t.view(1, 1, -1)

        return X, Y, Z, nx, ny, nz

    def view_rho(self, nx: int, ny: int, nz: int) -> torch.Tensor:
        rho = self.rho[:nx, :ny, :nz]
        rho.zero_()
        return rho

    def view_mask(self, nx: int, ny: int, nz: int) -> torch.Tensor:
        mask = self.mask[:nx, :ny, :nz]
        mask.fill_(False)
        return mask

    def view_rho_local(self, nx: int, ny: int, nz: int) -> torch.Tensor:
        return self.rho_local[:nx, :ny, :nz]

    def view_dist2(self, nx: int, ny: int, nz: int) -> torch.Tensor:
        return self.dist2[:nx, :ny, :nz]

    def view_dx(self, nx: int, ny: int, nz: int) -> torch.Tensor:
        return self.dx[:nx, :ny, :nz]

    def view_dy(self, nx: int, ny: int, nz: int) -> torch.Tensor:
        return self.dy[:nx, :ny, :nz]

    def view_dz(self, nx: int, ny: int, nz: int) -> torch.Tensor:
        return self.dz[:nx, :ny, :nz]

    def view_tmp(self, nx: int, ny: int, nz: int) -> torch.Tensor:
        return self.tmp[:nx, :ny, :nz]


# ==============================
# 3d. GPU promolecular density
# ==============================

def promolecular_density_on_grid_gpu(
    atoms: List[Dict],
    X: torch.Tensor,
    Y: torch.Tensor,
    Z: torch.Tensor,
    rho_out: torch.Tensor,
    dx_ws: torch.Tensor,
    dy_ws: torch.Tensor,
    dz_ws: torch.Tensor,
    dist2_ws: torch.Tensor,
    tmp_ws: torch.Tensor,
):
    if not atoms:
        raise ValueError("atoms list is empty in promolecular_density_on_grid_gpu")

    coords = np.array([a["coord"] for a in atoms], dtype=np.float32)
    Zs = np.array([z_of(a.get("element", "C")) for a in atoms], dtype=np.float32)
    sigmas = np.array([sigma_of(a.get("element", "C")) for a in atoms], dtype=np.float32)

    device = X.device
    coords_t = torch.from_numpy(coords).to(device=device)
    Zs_t = torch.from_numpy(Zs).to(device=device)
    sigmas_t = torch.from_numpy(sigmas).to(device=device)

    rho = rho_out
    rho.zero_()

    dx = dx_ws
    dy = dy_ws
    dz = dz_ws
    dist2 = dist2_ws
    tmp = tmp_ws

    for i in range(coords_t.shape[0]):
        ra = coords_t[i]
        Za = Zs_t[i]
        sa = sigmas_t[i]

        x0, y0, z0 = ra[0], ra[1], ra[2]
        inv_2s2 = -1.0 / (2.0 * (sa * sa))

        dx.copy_(X); dx.add_(-x0)
        dy.copy_(Y); dy.add_(-y0)
        dz.copy_(Z); dz.add_(-z0)

        dist2.copy_(dx); dist2.mul_(dx)
        tmp.copy_(dy); tmp.mul_(dy); dist2.add_(tmp)
        tmp.copy_(dz); tmp.mul_(dz); dist2.add_(tmp)

        tmp.copy_(dist2); tmp.mul_(inv_2s2)
        torch.exp_(tmp)
        tmp.mul_(Za)
        rho.add_(tmp)

    return rho


def promolecular_density_on_grid_gpu_vector(
    atoms: List[Dict],
    X: torch.Tensor,
    Y: torch.Tensor,
    Z: torch.Tensor,
    rho_out: torch.Tensor,
):
    if not atoms:
        raise ValueError("atoms list is empty in promolecular_density_on_grid_gpu_vector")

    coords = np.array([a["coord"] for a in atoms], dtype=np.float32)
    Zs = np.array([z_of(a.get("element", "C")) for a in atoms], dtype=np.float32)
    sigmas = np.array([sigma_of(a.get("element", "C")) for a in atoms], dtype=np.float32)

    device = X.device
    coords_t = torch.from_numpy(coords).to(device=device)
    Zs_t = torch.from_numpy(Zs).to(device=device)
    sigmas_t = torch.from_numpy(sigmas).to(device=device)

    Xb = X.unsqueeze(0)
    Yb = Y.unsqueeze(0)
    Zb = Z.unsqueeze(0)

    x0 = coords_t[:, 0].view(-1, 1, 1, 1)
    y0 = coords_t[:, 1].view(-1, 1, 1, 1)
    z0 = coords_t[:, 2].view(-1, 1, 1, 1)

    dx = Xb - x0
    dy = Yb - y0
    dz = Zb - z0
    dist2 = dx * dx + dy * dy + dz * dz

    sigmas_t = sigmas_t.view(-1, 1, 1, 1)
    inv_2s2 = -1.0 / (2.0 * sigmas_t * sigmas_t)

    tmp = torch.exp(dist2 * inv_2s2)
    Zs_b = Zs_t.view(-1, 1, 1, 1)
    tmp = tmp * Zs_b

    rho = tmp.sum(dim=0)
    rho_out.zero_()
    rho_out.copy_(rho)
    return rho_out


def _atom_batch_density_core(
    coords_t: torch.Tensor,
    weights_t: torch.Tensor,
    sigmas_t: torch.Tensor,
    X: torch.Tensor,
    Y: torch.Tensor,
    Z: torch.Tensor,
    rho_out: torch.Tensor,
    max_batch_elems: int = MAX_BATCH_ELEMENTS,
) -> torch.Tensor:
    if coords_t.numel() == 0:
        raise ValueError("coords_t is empty in _atom_batch_density_core")

    device = X.device
    coords_t = coords_t.to(device=device, dtype=torch.float32)
    weights_t = weights_t.to(device=device, dtype=torch.float32)
    sigmas_t = sigmas_t.to(device=device, dtype=torch.float32)

    nx, ny, nz = X.shape
    rho = rho_out
    rho.zero_()

    natoms = coords_t.shape[0]
    elems_per_grid = int(nx * ny * nz)
    if elems_per_grid <= 0:
        return rho

    max_B = max(1, max_batch_elems // max(elems_per_grid, 1))
    max_B = int(max_B)
    max_B = min(max_B, natoms)
    if max_B < 1:
        max_B = 1

    Xb = X.unsqueeze(0)
    Yb = Y.unsqueeze(0)
    Zb = Z.unsqueeze(0)

    for start in range(0, natoms, max_B):
        end = min(start + max_B, natoms)
        B = end - start

        chunk_coords = coords_t[start:end]
        chunk_w = weights_t[start:end]
        chunk_sigmas = sigmas_t[start:end]

        cx = chunk_coords[:, 0].view(B, 1, 1, 1)
        cy = chunk_coords[:, 1].view(B, 1, 1, 1)
        cz = chunk_coords[:, 2].view(B, 1, 1, 1)

        dist2_b = Xb - cx
        dist2_b.mul_(dist2_b)

        dy_b = Yb - cy
        dy_b.mul_(dy_b)
        dist2_b.add_(dy_b)
        del dy_b

        dz_b = Zb - cz
        dz_b.mul_(dz_b)
        dist2_b.add_(dz_b)
        del dz_b

        s_b = chunk_sigmas.view(B, 1, 1, 1)
        inv_2s2 = -1.0 / (2.0 * (s_b * s_b))

        tmp_b = dist2_b * inv_2s2
        torch.exp_(tmp_b)

        w_b = chunk_w.view(B, 1, 1, 1)
        tmp_b.mul_(w_b)

        rho.add_(tmp_b.sum(dim=0))

        del dist2_b, tmp_b, cx, cy, cz, s_b, inv_2s2, w_b

    return rho


def promolecular_density_on_grid_gpu_mode(
    atoms: List[Dict],
    X: torch.Tensor,
    Y: torch.Tensor,
    Z: torch.Tensor,
    rho_out: torch.Tensor,
    dx_ws: torch.Tensor,
    dy_ws: torch.Tensor,
    dz_ws: torch.Tensor,
    dist2_ws: torch.Tensor,
    tmp_ws: torch.Tensor,
    mode: str,
    rid: str,
) -> Tuple[torch.Tensor, str, bool]:
    nx, ny, nz = X.shape
    n_atoms = len(atoms)
    total_elems = int(n_atoms * nx * ny * nz)

    if mode == "safe":
        rho = promolecular_density_on_grid_gpu(
            atoms, X, Y, Z, rho_out, dx_ws, dy_ws, dz_ws, dist2_ws, tmp_ws
        )
        return rho, "safe", False

    if mode == "batch":
        if not atoms:
            raise ValueError("atoms list is empty in promolecular_density_on_grid_gpu_mode(batch)")
        coords = np.array([a["coord"] for a in atoms], dtype=np.float32)
        Zs = np.array([z_of(a.get("element", "C")) for a in atoms], dtype=np.float32)
        sigmas = np.array([sigma_of(a.get("element", "C")) for a in atoms], dtype=np.float32)

        device = X.device
        coords_t = torch.from_numpy(coords).to(device=device)
        Zs_t = torch.from_numpy(Zs).to(device=device)
        sigmas_t = torch.from_numpy(sigmas).to(device=device)

        rho = _atom_batch_density_core(
            coords_t=coords_t,
            weights_t=Zs_t,
            sigmas_t=sigmas_t,
            X=X,
            Y=Y,
            Z=Z,
            rho_out=rho_out,
            max_batch_elems=MAX_BATCH_ELEMENTS,
        )
        return rho, "batch", False

    if mode == "vector":
        if total_elems > MAX_VECTOR_ELEMENTS:
            print(
                f"[WARN] gpu-density-mode 'vector' skipped for fragment {rid} "
                f"(atoms={n_atoms}, grid={nx}x{ny}x{nz}, elems={total_elems}) "
                f"→ falling back to 'safe' atom-loop to avoid GPU OOM."
            )
            rho = promolecular_density_on_grid_gpu(
                atoms, X, Y, Z, rho_out, dx_ws, dy_ws, dz_ws, dist2_ws, tmp_ws
            )
            return rho, "safe", True

        rho = promolecular_density_on_grid_gpu_vector(
            atoms, X, Y, Z, rho_out
        )
        return rho, "vector", False

    rho = promolecular_density_on_grid_gpu(
        atoms, X, Y, Z, rho_out, dx_ws, dy_ws, dz_ws, dist2_ws, tmp_ws
    )
    return rho, "safe", False


def promolecular_density_on_grid_gpu_mode_arrays(
    atom_indices: np.ndarray,
    atom_coords: np.ndarray,
    atom_Z: np.ndarray,
    atom_sigma: np.ndarray,
    X: torch.Tensor,
    Y: torch.Tensor,
    Z: torch.Tensor,
    rho_out: torch.Tensor,
    dx_ws: torch.Tensor,
    dy_ws: torch.Tensor,
    dz_ws: torch.Tensor,
    dist2_ws: torch.Tensor,
    tmp_ws: torch.Tensor,
    mode: str,
    rid: str,
) -> Tuple[torch.Tensor, str, bool]:
    """
    Legacy approach: Received `atoms` (list[dict]) and reconstructed coords/Z/sigma for every residue.
    Optimized approach: Uses global SoA (Structure of Arrays) + fragment `atom_indices` 
                        for direct tensor operations without Python loops.

    Returns
    -------
    rho      : torch.Tensor, shape (nx, ny, nz)
    eff_mode : "batch" or "vector"
    fallback : True if "vector" was requested but fell back to "batch" due to element limits.
    """
    nx, ny, nz = X.shape

    if atom_indices is None or atom_indices.size == 0:
        rho_out.zero_()
        return rho_out, "batch", False

    # Slice global SoA arrays to get fragment data (pure indexing, no Python loops)
    coords = atom_coords[atom_indices]          # (N, 3)
    Zs = atom_Z[atom_indices]                   # (N,)
    sigmas = atom_sigma[atom_indices]           # (N,)

    device = X.device
    coords_t = torch.from_numpy(coords).to(device=device, dtype=torch.float32)
    Zs_t = torch.from_numpy(Zs).to(device=device, dtype=torch.float32)
    sigmas_t = torch.from_numpy(sigmas).to(device=device, dtype=torch.float32)

    n_atoms = coords_t.shape[0]
    elems_per_grid = int(nx * ny * nz)
    total_elems = int(n_atoms * elems_per_grid)

    # Safety mechanism: Force 'batch' mode if (num_atoms * grid_size) exceeds memory limits
    if mode == "vector" and total_elems > MAX_VECTOR_ELEMENTS:
        print(
            f"[WARN] gpu-density-mode 'vector' skipped for fragment {rid} "
            f"(atoms={n_atoms}, grid={nx}x{ny}x{nz}, elems={total_elems}) "
            f"→ falling back to 'batch' array-mode."
        )
        mode = "batch"
        fallback = True
    else:
        fallback = False

    # Computation: 'batch' mode delegates to _atom_batch_density_core
    if mode in ("safe", "batch"):
        rho = _atom_batch_density_core(
            coords_t=coords_t,
            weights_t=Zs_t,
            sigmas_t=sigmas_t,
            X=X,
            Y=Y,
            Z=Z,
            rho_out=rho_out,
            max_batch_elems=MAX_BATCH_ELEMENTS,
        )
        return rho, "batch", fallback

    # Case mode == "vector": Perform one-shot broadcasting
    Xb = X.unsqueeze(0)  # (1, nx, ny, nz)
    Yb = Y.unsqueeze(0)
    Zb = Z.unsqueeze(0)

    cx = coords_t[:, 0].view(-1, 1, 1, 1)  # (N,1,1,1)
    cy = coords_t[:, 1].view(-1, 1, 1, 1)
    cz = coords_t[:, 2].view(-1, 1, 1, 1)

    dx = Xb - cx
    dy = Yb - cy
    dz = Zb - cz
    dist2 = dx * dx + dy * dy + dz * dz

    sigmas_b = sigmas_t.view(-1, 1, 1, 1)
    inv_2s2 = -1.0 / (2.0 * sigmas_b * sigmas_b)

    tmp = torch.exp(dist2 * inv_2s2)
    Zs_b = Zs_t.view(-1, 1, 1, 1)
    tmp = tmp * Zs_b

    rho = tmp.sum(dim=0)
    rho_out.zero_()
    rho_out.copy_(rho)
    return rho_out, "vector", fallback


# ==============================
# 3e. xTB-based density (A-mode)
# ==============================

def write_xyz(atoms: List[Dict], xyz_path: str, charge: int = 0, mult: int = 1):
    with open(xyz_path, "w") as f:
        f.write(f"{len(atoms)}\n")
        f.write(f"charge={charge} mult={mult}\n")
        for a in atoms:
            elem = a.get("element", "C")
            x, y, z = a["coord"]
            f.write(f"{elem:2s} {x: .6f} {y: .6f} {z: .6f}\n")


def run_xtb(xyz_path: str, workdir: str, xtb_bin: str = "xtb") -> Optional[str]:
    ensure_dir(workdir)
    cmd = [xtb_bin, os.path.basename(xyz_path), "--gfn2", "--charges"]
    try:
        subprocess.run(
            cmd,
            cwd=workdir,
            check=True,
            stdout=subprocess.PIPE,
            stderr=subprocess.PIPE,
            text=True,
        )
    except FileNotFoundError:
        print(f"[xTB ERROR] Binary '{xtb_bin}' not found. Falling back to B-mode.")
        return None
    except subprocess.CalledProcessError as e:
        print("[xTB FAIL]", e.stderr[:300])
        return None

    charges_path = os.path.join(workdir, "charges")
    return charges_path if os.path.exists(charges_path) else None


def parse_xtb_charges(charges_path: str) -> List[float]:
    vals = []
    with open(charges_path, "r") as f:
        for line in f:
            line = line.strip()
            if not line or line.startswith("#"):
                continue
            parts = line.split()
            try:
                vals.append(float(parts[-1]))
            except Exception:
                continue
    return vals


def density_from_charges_on_grid_gpu(
    atoms: List[Dict],
    charges: List[float],
    X: torch.Tensor,
    Y: torch.Tensor,
    Z: torch.Tensor,
    rho_out: torch.Tensor,
    dx_ws: torch.Tensor,
    dy_ws: torch.Tensor,
    dz_ws: torch.Tensor,
    dist2_ws: torch.Tensor,
    tmp_ws: torch.Tensor,
):
    if len(charges) != len(atoms):
        return promolecular_density_on_grid_gpu(
            atoms, X, Y, Z, rho_out, dx_ws, dy_ws, dz_ws, dist2_ws, tmp_ws
        )

    coords = np.array([a["coord"] for a in atoms], dtype=np.float32)
    charges_arr = np.array(charges, dtype=np.float32)

    device = X.device
    coords_t = torch.from_numpy(coords).to(device=device)
    charges_t = torch.from_numpy(charges_arr).to(device=device)

    rho = rho_out
    rho.zero_()

    dx = dx_ws
    dy = dy_ws
    dz = dz_ws
    dist2 = dist2_ws
    tmp = tmp_ws

    for i in range(coords_t.shape[0]):
        r = coords_t[i]
        q = charges_t[i]
        elem = atoms[i].get("element", "C")
        Za = float(z_of(elem))
        sa = float(sigma_of(elem))
        eff = max(Za - float(q.item()), 0.1)
        inv_2s2 = -1.0 / (2.0 * (sa * sa))

        x0, y0, z0 = r[0], r[1], r[2]

        dx.copy_(X); dx.add_(-x0)
        dy.copy_(Y); dy.add_(-y0)
        dz.copy_(Z); dz.add_(-z0)

        dist2.copy_(dx); dist2.mul_(dx)
        tmp.copy_(dy); tmp.mul_(dy); dist2.add_(tmp)
        tmp.copy_(dz); tmp.mul_(dz); dist2.add_(tmp)

        tmp.copy_(dist2); tmp.mul_(inv_2s2)
        torch.exp_(tmp)
        tmp.mul_(eff)
        rho.add_(tmp)

    return rho


def density_from_charges_on_grid_gpu_vector(
    atoms: List[Dict],
    charges: List[float],
    X: torch.Tensor,
    Y: torch.Tensor,
    Z: torch.Tensor,
    rho_out: torch.Tensor,
):
    if len(charges) != len(atoms):
        return promolecular_density_on_grid_gpu_vector(
            atoms, X, Y, Z, rho_out
        )

    coords = np.array([a["coord"] for a in atoms], dtype=np.float32)
    charges_arr = np.array(charges, dtype=np.float32)

    effZ = []
    sigmas = []
    for a, q in zip(atoms, charges_arr):
        elem = a.get("element", "C")
        Za = z_of(elem)
        sa = sigma_of(elem)
        eff = max(Za - float(q), 0.1)
        effZ.append(eff)
        sigmas.append(sa)

    effZ = np.array(effZ, dtype=np.float32)
    sigmas = np.array(sigmas, dtype=np.float32)

    device = X.device
    coords_t = torch.from_numpy(coords).to(device=device)
    effZ_t = torch.from_numpy(effZ).to(device=device)
    sigmas_t = torch.from_numpy(sigmas).to(device=device)

    Xb = X.unsqueeze(0)
    Yb = Y.unsqueeze(0)
    Zb = Z.unsqueeze(0)

    x0 = coords_t[:, 0].view(-1, 1, 1, 1)
    y0 = coords_t[:, 1].view(-1, 1, 1, 1)
    z0 = coords_t[:, 2].view(-1, 1, 1, 1)

    dx = Xb - x0
    dy = Yb - y0
    dz = Zb - z0
    dist2 = dx * dx + dy * dy + dz * dz

    sigmas_t = sigmas_t.view(-1, 1, 1, 1)
    inv_2s2 = -1.0 / (2.0 * sigmas_t * sigmas_t)

    tmp = torch.exp(dist2 * inv_2s2)
    effZ_b = effZ_t.view(-1, 1, 1, 1)
    tmp = tmp * effZ_b

    rho = tmp.sum(dim=0)
    rho_out.zero_()
    rho_out.copy_(rho)
    return rho_out


def density_from_charges_on_grid_gpu_mode(
    atoms: List[Dict],
    charges: List[float],
    X: torch.Tensor,
    Y: torch.Tensor,
    Z: torch.Tensor,
    rho_out: torch.Tensor,
    dx_ws: torch.Tensor,
    dy_ws: torch.Tensor,
    dz_ws: torch.Tensor,
    dist2_ws: torch.Tensor,
    tmp_ws: torch.Tensor,
    mode: str,
    rid: str,
) -> Tuple[torch.Tensor, str, bool]:
    nx, ny, nz = X.shape
    n_atoms = len(atoms)
    total_elems = int(n_atoms * nx * ny * nz)

    if mode == "safe":
        rho = density_from_charges_on_grid_gpu(
            atoms, charges, X, Y, Z, rho_out, dx_ws, dy_ws, dz_ws, dist2_ws, tmp_ws
        )
        return rho, "safe", False

    if mode == "batch":
        if len(charges) != len(atoms):
            rho = density_from_charges_on_grid_gpu(
                atoms, charges, X, Y, Z, rho_out, dx_ws, dy_ws, dz_ws, dist2_ws, tmp_ws
            )
            return rho, "batch", False

        if not atoms:
            raise ValueError("atoms list is empty in density_from_charges_on_grid_gpu_mode(batch)")

        coords = np.array([a["coord"] for a in atoms], dtype=np.float32)
        charges_arr = np.array(charges, dtype=np.float32)

        effZ = []
        sigmas = []
        for a, q in zip(atoms, charges_arr):
            elem = a.get("element", "C")
            Za = z_of(elem)
            sa = sigma_of(elem)
            eff = max(Za - float(q), 0.1)
            effZ.append(eff)
            sigmas.append(sa)

        effZ = np.array(effZ, dtype=np.float32)
        sigmas = np.array(sigmas, dtype=np.float32)

        device = X.device
        coords_t = torch.from_numpy(coords).to(device=device)
        effZ_t = torch.from_numpy(effZ).to(device=device)
        sigmas_t = torch.from_numpy(sigmas).to(device=device)

        rho = _atom_batch_density_core(
            coords_t=coords_t,
            weights_t=effZ_t,
            sigmas_t=sigmas_t,
            X=X,
            Y=Y,
            Z=Z,
            rho_out=rho_out,
            max_batch_elems=MAX_BATCH_ELEMENTS,
        )
        return rho, "batch", False

    if mode == "vector":
        if total_elems > MAX_VECTOR_ELEMENTS:
            print(
                f"[WARN] gpu-density-mode 'vector' (A-mode) skipped for fragment {rid} "
                f"(atoms={n_atoms}, grid={nx}x{ny}x{nz}, elems={total_elems}) "
                f"→ falling back to 'safe' atom-loop to avoid GPU OOM."
            )
            rho = density_from_charges_on_grid_gpu(
                atoms, charges, X, Y, Z, rho_out, dx_ws, dy_ws, dz_ws, dist2_ws, tmp_ws
            )
            return rho, "safe", True

        rho = density_from_charges_on_grid_gpu_vector(
            atoms, charges, X, Y, Z, rho_out
        )
        return rho, "vector", False

    rho = density_from_charges_on_grid_gpu(
        atoms, charges, X, Y, Z, rho_out, dx_ws, dy_ws, dz_ws, dist2_ws, tmp_ws
    )
    return rho, "safe", False


# ==============================
# 4. Density-based features
# ==============================

def density_features_for_residue_gpu(
    rho: torch.Tensor,
    X: torch.Tensor,
    Y: torch.Tensor,
    Z: torch.Tensor,
    ref_point: np.ndarray,
    res_atom_coords: np.ndarray,
    R_env: float,
    mask_ws: torch.Tensor,
    rho_local_ws: torch.Tensor,
    dist2_ws: torch.Tensor,
    tmp_ws: torch.Tensor,
    dx_ws: torch.Tensor,
    dy_ws: torch.Tensor,
    dz_ws: torch.Tensor,
    rid: str,
) -> Dict:
    device = rho.device

    xs = X[:, 0, 0]
    ys = Y[0, :, 0]
    zs = Z[0, 0, :]

    dx_step = float((xs[1] - xs[0]).item()) if xs.numel() > 1 else 1.0
    dy_step = float((ys[1] - ys[0]).item()) if ys.numel() > 1 else 1.0
    dz_step = float((zs[1] - zs[0]).item()) if zs.numel() > 1 else 1.0

    dV = dx_step * dy_step * dz_step
    R2 = float(R_env * R_env)

    mask = mask_ws
    mask.fill_(False)

    coords_res_t = torch.from_numpy(res_atom_coords.astype(np.float32)).to(device=device)

    dist2 = dist2_ws
    tmp = tmp_ws
    dx = dx_ws
    dy = dy_ws
    dz = dz_ws

    for i in range(coords_res_t.shape[0]):
        ra = coords_res_t[i]
        x0, y0, z0 = ra[0], ra[1], ra[2]

        dx.copy_(X); dx.add_(-x0)
        dy.copy_(Y); dy.add_(-y0)
        dz.copy_(Z); dz.add_(-z0)

        dist2.copy_(dx); dist2.mul_(dx)
        tmp.copy_(dy); tmp.mul_(dy); dist2.add_(tmp)
        tmp.copy_(dz); tmp.mul_(dz); dist2.add_(tmp)

        mask |= dist2 <= R2

    rho_local = rho_local_ws
    rho_local.zero_()
    rho_local[mask] = rho[mask]

    total_density = float((rho_local.sum() * dV).item())

    ref_t = torch.from_numpy(ref_point.astype(np.float32)).to(device=device)

    if total_density <= 0.0:
        centroid = ref_point.astype(float)
        dipole = np.zeros(3, dtype=float)
    else:
        cx = float((rho_local * X).sum().mul(dV).div(total_density).item())
        cy = float((rho_local * Y).sum().mul(dV).div(total_density).item())
        cz = float((rho_local * Z).sum().mul(dV).div(total_density).item())
        centroid = np.array([cx, cy, cz], dtype=float)

        RX = X - ref_t[0]
        RY = Y - ref_t[1]
        RZ = Z - ref_t[2]

        px = float((rho_local * RX).sum().mul(dV).item())
        py = float((rho_local * RY).sum().mul(dV).item())
        pz = float((rho_local * RZ).sum().mul(dV).item())
        dipole = np.array([px, py, pz], dtype=float)

    curv_val = 0.0
    curv_failed = False
    try:
        gx, gy, gz = torch.gradient(
            rho_local,
            spacing=(dx_step, dy_step, dz_step),
            edge_order=1,
        )
        curv = (gx * gx + gy * gy + gz * gz).sum().mul(dV)
        curv_val = float(curv.item())
    except Exception as e:
        print(f"[WARN] curv_rho computation failed for residue {rid}: {e}")
        curv_val = 0.0
        curv_failed = True

    return {
        "total_density": total_density,
        "centroid": centroid.tolist(),
        "dipole_rho": dipole.tolist(),
        "curv_rho": curv_val,
        "curv_rho_failed": curv_failed,
    }


# ==============================
# 4b. Charge-based features / surrogate
# ==============================

def charge_features_for_residue_from_fragment(
    rid: str,
    atom_index_to_resid: Dict[int, str],
    resid_to_atom_coords: Dict[str, List[np.ndarray]],
    frag_atoms: List[Dict],
    charges: List[float],
) -> Optional[Dict]:
    if len(frag_atoms) != len(charges):
        return None

    coords_res = resid_to_atom_coords[rid]
    ref_point = np.mean(np.stack(coords_res, axis=0), axis=0)

    q_res = 0.0
    dipole_q = np.zeros(3, dtype=float)

    for a, q in zip(frag_atoms, charges):
        atom_idx = int(a["atom_index"])
        res_of_atom = atom_index_to_resid.get(atom_idx, None)
        if res_of_atom != rid:
            continue
        coord = np.array(a["coord"], dtype=float)
        q_res += q
        dipole_q += q * (coord - ref_point)

    return {
        "partial_charge": float(q_res),
        "dipole_q": dipole_q.tolist(),
    }


def residue_base_charge(resname: str) -> float:
    rn = resname.strip().upper()
    if rn in ("ASP", "GLU"):
        return -1.0
    if rn in ("LYS", "ARG"):
        return +1.0
    if rn == "HIS":
        return +0.1
    return 0.0


def env_scale_from_sasa(sasa: float, mean_sasa: float, alpha: float = 0.4) -> float:
    if mean_sasa <= 1e-8:
        return 1.0
    r = sasa / mean_sasa
    scale = 1.0 + alpha * (r - 1.0)
    return float(np.clip(scale, 0.5, 1.5))


def residue_atom_charge_surrogate(residue) -> float:
    elem_weights = {
        "O":  -0.40,
        "N":  +0.30,
        "S":  -0.25,
        "SE": -0.25,
        "P":  -0.35,
        "F":  -0.20,
        "CL": -0.20,
        "BR": -0.20,
        "I":  -0.20,
        "MG": +0.50,
        "CA": +0.50,
        "MN": +0.60,
        "FE": +0.60,
        "ZN": +0.60,
        "CU": +0.60,
        "NI": +0.60,
        "CO": +0.60,
        # C, H, others → 0.0
    }

    atoms = residue.get("atoms", [])
    q = 0.0
    for atom in atoms:
        elem_raw = atom.get("element", "")
        elem = str(elem_raw).strip().upper()
        q += elem_weights.get(elem, 0.0)

    return float(q)


def compute_b_mode_charge_atom_based(
    res,
    rid,
    resid_to_sasa2,
    mean_sasa,
    sasa_alpha,
    env_scale_from_sasa,
) -> float:
    sasa_val = float(resid_to_sasa2.get(rid, 0.0))
    base_q = residue_atom_charge_surrogate(res)

    if abs(base_q) > 1e-6:
        scale = env_scale_from_sasa(sasa_val, mean_sasa, alpha=sasa_alpha)
    else:
        scale = 1.0

    partial_charge_B = base_q * scale
    return partial_charge_B


# ==============================
# 5. Ep helpers (embedding, neighbor, field)
# ==============================

RES_EMBED = {
    "ALA": [0.0, 0.5, 0.2, 0.0],
    "ARG": [1.0, -1.0, 1.0, 0.0],
    "ASN": [0.0, -0.5, 0.4, 0.0],
    "ASP": [-1.0, -1.0, 0.4, 0.0],
    "ASX": [-0.5, -1.0, 0.4, 0.0],
    "CSO": [0.0, -0.5, 0.4, 0.0],
    "CYS": [0.0, 0.5, 0.4, 0.0],
    "GLN": [0.0, -0.5, 0.6, 0.0],
    "GLU": [-1.0, -1.0, 0.6, 0.0],
    "GLX": [-0.5, -1.0, 0.6, 0.0],
    "GLY": [0.0, 0.0, 0.0, 0.0],
    "HIS": [0.5, -0.5, 0.6, 0.5],
    "HYP": [0.0, -0.5, 0.4, 0.0],
    "ILE": [0.0, 1.0, 0.8, 0.0],
    "LEU": [0.0, 1.0, 0.8, 0.0],
    "LYS": [1.0, -1.0, 0.8, 0.0],
    "MET": [0.0, 0.5, 0.8, 0.0],
    "MSE": [0.0, 0.5, 0.8, 0.0],
    "PHE": [0.0, 1.0, 1.0, 1.0],
    "PRO": [0.0, 0.0, 0.4, 0.0],
    "PYL": [1.0, -1.0, 1.0, 0.0],
    "SEC": [0.0, 0.5, 0.4, 0.0],
    "SER": [0.0, -0.5, 0.2, 0.0],
    "THR": [0.0, -0.5, 0.4, 0.0],
    "TRP": [0.0, 1.0, 1.0, 1.0],
    "TYR": [0.0, 0.5, 1.0, 1.0],
    "VAL": [0.0, 1.0, 0.6, 0.0],
    "XAA": [0.0, 0.0, 0.6, 0.0],
    "XLE": [0.0, 1.0, 0.8, 0.0],
}

DEFAULT_EMBED = np.array([0.0, 0.0, 0.50, 0.50], dtype=float)


def residue_to_embedding(resname: str) -> np.ndarray:
    rn = resname.strip().upper()
    vec = RES_EMBED.get(rn, RES_EMBED["XAA"])
    return np.array(vec, dtype=float)


def build_residue_centers(meta: Dict) -> Dict[str, np.ndarray]:
    centers: Dict[str, np.ndarray] = {}
    for r in meta["residues"]:
        coords = np.array([a["coord"] for a in r["atoms"]], dtype=float)
        centers[r["id"]] = coords.mean(axis=0)
    return centers


# ---------- NEW: 10D geometric neighbor block ----------

def compute_geometric_block(
    center: np.ndarray,
    neighbor_coords: np.ndarray,
    r_max: float,
    eps_shell: float = 1e-6,
    eps_pca: float = 1e-8,
) -> np.ndarray:
    """
    Compute 10 scalar neighbor-geometry features for residue p:

      (7)  N_in(p)          : #neighbors with d <= r_max/2
      (8)  N_out(p)         : #neighbors with r_max/2 < d <= r_max
      (9)  R_shell(p)       : N_in / (N_out + eps_shell)

      (10) r_mean(p)        : mean neighbor distance
      (11) r_var(p)         : variance of neighbor distances

      (12) linearity(p)
      (13) planarity(p)
      (14) isotropy(p)      : eigenvalue-based anisotropy of direction tensor

      (15) H_mean(p)        : mean curvature surrogate
      (16) K_gauss(p)       : Gaussian curvature surrogate

    Curvature is estimated by projecting neighbors into a local PCA frame
    and fitting a quadratic patch z = a x^2 + b y^2 + c x y.
    """
    if neighbor_coords.size == 0:
        return np.zeros(10, dtype=float)

    center = np.asarray(center, dtype=float)
    neighbor_coords = np.asarray(neighbor_coords, dtype=float)

    # distances
    vecs = neighbor_coords - center
    dists = np.linalg.norm(vecs, axis=1)
    if dists.size == 0:
        return np.zeros(10, dtype=float)

    # restrict to neighbors within r_max for shell definition
    # (contacts are already defined by KDTree, so this is mostly a cap)
    if r_max is not None and r_max > 0.0:
        mask_r = dists <= r_max
        if not np.any(mask_r):
            # fallback: use all neighbors anyway
            dists_use = dists
            vecs_use = vecs
        else:
            dists_use = dists[mask_r]
            vecs_use = vecs[mask_r]
    else:
        dists_use = dists
        vecs_use = vecs

    if dists_use.size == 0:
        return np.zeros(10, dtype=float)

    # ---------- 1. Two-shell radial density ----------
    r_cut = r_max if (r_max is not None and r_max > 0.0) else float(dists_use.max())
    if r_cut < eps_shell:
        r_cut = float(dists_use.max())

    r_inner = 0.5 * r_cut

    N_in = float(np.sum(dists_use <= r_inner))
    N_out = float(np.sum((dists_use > r_inner) & (dists_use <= r_cut)))
    R_shell = N_in / (N_out + eps_shell)

    # ---------- 2. Radial distribution moments ----------
    r_mean = float(dists_use.mean())
    r_var = float(dists_use.var()) if dists_use.size > 1 else 0.0

    # ---------- 3. Angular shape tensor / anisotropy ----------
    unit_vecs = []
    for v, d in zip(vecs_use, dists_use):
        if d <= eps_pca:
            continue
        unit_vecs.append(v / d)
    if len(unit_vecs) == 0:
        linearity = 0.0
        planarity = 0.0
        isotropy = 1.0
    else:
        U = np.stack(unit_vecs, axis=0)
        mean_u = U.mean(axis=0)
        centered = U - mean_u
        C = centered.T @ centered / max(U.shape[0], 1)
        evals = np.linalg.eigvalsh(C)
        evals = np.sort(np.maximum(evals, 0.0))
        lam1, lam2, lam3 = evals[-1], evals[-2], evals[-3]

        denom = lam1 + lam2 + lam3 + eps_pca
        linearity = (lam1 - lam2) / denom
        planarity = (lam2 - lam3) / denom
        isotropy = lam3 / denom

    # ---------- 4. Local surface curvature surrogate ----------
    # PCA to get local frame
    if vecs_use.shape[0] >= 3:
        # covariance for PCA
        mean_v = vecs_use.mean(axis=0)
        C_full = (vecs_use - mean_v).T @ (vecs_use - mean_v) / max(vecs_use.shape[0], 1)
        evals_full, evecs_full = np.linalg.eigh(C_full)
        idx = np.argsort(evals_full)[::-1]
        evecs_full = evecs_full[:, idx]

        # local coordinates
        local_coords = (vecs_use @ evecs_full)
        x = local_coords[:, 0]
        y = local_coords[:, 1]
        z = local_coords[:, 2]

        # quadratic fit: z = a x^2 + b y^2 + c x y
        A = np.stack([x * x, y * y, x * y], axis=1)
        try:
            # if underdetermined, lstsq still returns minimal norm solution
            coeffs, *_ = np.linalg.lstsq(A, z, rcond=None)
            a, b, c = coeffs.tolist()
            H = float(a + b)        # mean curvature surrogate
            K = float(a * b - 0.25 * c * c)  # Gaussian curvature surrogate
        except Exception:
            H = 0.0
            K = 0.0
    else:
        H = 0.0
        K = 0.0

    return np.array(
        [
            N_in,        # (7)
            N_out,       # (8)
            R_shell,     # (9)
            r_mean,      # (10)
            r_var,       # (11)
            linearity,   # (12)
            planarity,   # (13)
            isotropy,    # (14)
            H,           # (15)
            K,           # (16)
        ],
        dtype=float,
    )


def geometric_block_from_overlaps(
    rid: str,
    overlaps: Dict[str, List[str]],
    centers: Dict[str, np.ndarray],
    r_max: float,
) -> np.ndarray:
    """
    Build the 10D geometric neighbor feature block for residue `rid`
    from the contact graph overlaps and residue centers.
    """
    if rid not in centers:
        return np.zeros(10, dtype=float)

    neigh_ids = overlaps.get(rid, [])
    if not neigh_ids:
        return np.zeros(10, dtype=float)

    center = centers[rid]
    coords = []
    for nid in neigh_ids:
        if nid in centers:
            coords.append(centers[nid])

    if not coords:
        return np.zeros(10, dtype=float)

    neighbor_coords = np.stack(coords, axis=0)
    return compute_geometric_block(center=center, neighbor_coords=neighbor_coords, r_max=r_max)


FIELD_METALS = {"MG", "CA", "MN", "FE", "ZN", "CU", "NI", "CO", "MO", "W", "V"}
FIELD_P = {"P"}
FIELD_HALOGENS = {"F", "CL", "BR", "I"}


def compute_field_terms_for_residue(
    rid: str,
    res_map: Dict[str, Dict],
    centers: Dict[str, np.ndarray],
    overlaps: Dict[str, List[str]],
    eps: float = 1e-3,
) -> Tuple[float, float, float]:
    """
    Compute local field-like terms around residue p:

      F_metal(p) = sum_{metal atoms a in p ∪ N(p)} 1 / ||r_a - r_p||^2
      F_P(p)     = same for P atoms
      F_hal(p)   = same for halogens (F, Cl, Br, I)

    Only center residue + contact neighbors are used (overlaps).
    """
    if rid not in centers:
        return 0.0, 0.0, 0.0

    rp = centers[rid]
    F_metal = 0.0
    F_P_val = 0.0
    F_hal = 0.0

    neighbor_ids = [rid] + overlaps.get(rid, [])
    for nid in neighbor_ids:
        res = res_map.get(nid)
        if res is None:
            continue
        for a in res["atoms"]:
            elem = str(a.get("element", "")).strip().upper()
            if elem not in FIELD_METALS and elem not in FIELD_P and elem not in FIELD_HALOGENS:
                continue
            coord = np.array(a["coord"], dtype=float)
            d = float(np.linalg.norm(coord - rp))
            d_eff = max(d, eps)
            contrib = 1.0 / (d_eff * d_eff)

            if elem in FIELD_METALS:
                F_metal += contrib
            elif elem in FIELD_P:
                F_P_val += contrib
            elif elem in FIELD_HALOGENS:
                F_hal += contrib

    return float(F_metal), float(F_P_val), float(F_hal)


# ==============================
# 6. Full GPU-accelerated driver
# ==============================

def run_pipeline_gpu(
    pdb_path: str,
    out_dir: str,
    radius: float,
    grid_step: float,
    padding: float,
    xtb_bin: str,
    select_A: List[str],
    save_densities: bool,
    R_env: float,
    sasa_alpha: float,
    no_pair_contacts: bool,
    block_size: int,
    device: str,
    gpu_density_mode: str,
):
    ensure_dir(out_dir)

    traj = load_traj(pdb_path)
    residues = collect_residues_from_mdtraj(traj)
    print(f"[INFO] Residues collected: {len(residues)}")

    resid_to_sasa = compute_residue_sasa_from_traj(traj, residues)
    sasa_vals = list(resid_to_sasa.values())
    mean_sasa = float(np.mean(sasa_vals)) if len(sasa_vals) > 0 else 0.0

    del traj
    gc.collect()

    overlaps, pair_contacts = compute_residue_contacts_kdtree(
        residues,
        radius=radius,
        store_pairs=(not no_pair_contacts),
    )
    if no_pair_contacts:
        pair_contacts = {}

    meta_path = os.path.join(out_dir, "meta.json")
    export_meta(
        pdb_name=os.path.basename(pdb_path),
        radius=radius,
        residues=residues,
        overlaps=overlaps,
        pair_contacts=pair_contacts,
        resid_to_sasa=resid_to_sasa,
        out_path=meta_path,
    )

    del residues
    del resid_to_sasa
    gc.collect()

    meta = load_json(meta_path)
    res_map, overlaps_m, atom_index_to_resid, resid_to_atom_coords, resid_to_sasa2 = \
        build_residue_maps(meta)

    resid_to_grid_axes, max_nx, max_ny, max_nz = precompute_residue_grids(
        resid_to_atom_coords,
        R_env=R_env,
        grid_step=grid_step,
        padding=padding,
    )
    print(f"[INFO] Allocating GPU density workspace: ({max_nx}, {max_ny}, {max_nz}) on {device}")
    density_ws = DensityWorkspaceGPU(max_nx, max_ny, max_nz, device=device)
    log_gpu_usage("after_workspace_alloc", device)

    centers = build_residue_centers(meta)
    
    # ===== NEW: global atom SoA array ready =====
    atom_coords, atom_Z, atom_sigma, resid_to_atom_indices = build_global_atom_arrays(meta)

    A_dir = os.path.join(out_dir, "density_A_overrides")
    ensure_dir(A_dir)

    density_manifest = {
        "A_residues": [],
        "B_only_residues": [],
        "files": {},
        "modes": {},
        "params": {
            "gpu_density_mode_requested": gpu_density_mode,
            "gpu_density_vector_max_elems": int(MAX_VECTOR_ELEMENTS),
            "gpu_density_batch_max_elems": int(MAX_BATCH_ELEMENTS),
        },
        "curv_rho_failed_residues": [],
    }

    env_features: Dict[str, Dict] = {}

    res_ids = list(res_map.keys())
    n_res = len(res_ids)
    feature_dim_theory = 9
    feature_dim_sim = feature_dim_theory + 10  # 9 + 10 = 19

    Ep_theory = np.zeros((n_res, feature_dim_theory), dtype=float)
    Ep_sim = np.zeros((n_res, feature_dim_sim), dtype=float)

    curv_rho_failed_residues: List[str] = []

    for block_start in range(0, n_res, block_size):
        block_end = min(block_start + block_size, n_res)
        print(f"[INFO] GPU block {block_start}..{block_end-1} / {n_res}")
        log_gpu_usage(f"block_{block_start}_{block_end-1}_start", device)

        for idx in range(block_start, block_end):
            rid = res_ids[idx]
            res = res_map[rid]

            neighbor_ids = overlaps_m.get(rid, [])

            frag_atom_indices = build_fragment_indices(
                center_res_id=rid,
                neighbor_res_ids=neighbor_ids,
                resid_to_atom_indices=resid_to_atom_indices,
            )
            frag_atoms = None
            
            if rid in select_A:
                frag_atoms = build_fragment_atoms(res_map, rid, neighbor_ids)

            coords_res_list = resid_to_atom_coords[rid]
            coords_res = np.stack(coords_res_list, axis=0).astype(np.float32)
            ref_point = coords_res.mean(axis=0)

            xs, ys, zs = resid_to_grid_axes[rid]
            X, Y, Z, nx, ny, nz = density_ws.prepare_grid_from_axes(xs, ys, zs)
            rho_view = density_ws.view_rho(nx, ny, nz)

            dx_view = density_ws.view_dx(nx, ny, nz)
            dy_view = density_ws.view_dy(nx, ny, nz)
            dz_view = density_ws.view_dz(nx, ny, nz)
            dist2_view = density_ws.view_dist2(nx, ny, nz)
            tmp_view = density_ws.view_tmp(nx, ny, nz)

            # ----- B-mode density -----
            rho_B, eff_mode_B, fallback_B = promolecular_density_on_grid_gpu_mode_arrays(
                atom_indices=frag_atom_indices,
                atom_coords=atom_coords,
                atom_Z=atom_Z,
                atom_sigma=atom_sigma,
                X=X,
                Y=Y,
                Z=Z,
                rho_out=rho_view,
                dx_ws=dx_view,
                dy_ws=dy_view,
                dz_ws=dz_view,
                dist2_ws=dist2_view,
                tmp_ws=tmp_view,
                mode=gpu_density_mode,
                rid=rid,
            )
            rho_final = rho_B

            rhoB_path = None
            rhoA_path = None
            charges_for_rid = None
            rho_A = None
            eff_mode_A: Optional[str] = None
            fallback_A: bool = False
            xtb_success = False
            xtb_used = False
            charge_length_mismatch = False

            # ----- A-mode (xTB) -----
            if rid in select_A:
                workdir = os.path.join(A_dir, f"work_{rid.replace(':', '_')}")
                xyz_path = os.path.join(workdir, "frag.xyz")
                write_xyz(frag_atoms, xyz_path)

                charges_path = run_xtb(xyz_path, workdir, xtb_bin=xtb_bin)
                if charges_path is not None:
                    charges = parse_xtb_charges(charges_path)
                    if len(charges) == len(frag_atoms):
                        xtb_success = True
                        xtb_used = True
                        charges_for_rid = charges

                        rho_A, eff_mode_A, fallback_A = density_from_charges_on_grid_gpu_mode(
                            atoms=frag_atoms,
                            charges=charges,
                            X=X,
                            Y=Y,
                            Z=Z,
                            rho_out=rho_view,
                            dx_ws=dx_view,
                            dy_ws=dy_view,
                            dz_ws=dz_view,
                            dist2_ws=dist2_view,
                            tmp_ws=tmp_view,
                            mode=gpu_density_mode,
                            rid=rid,
                        )
                        rho_final = rho_A

                        if save_densities:
                            rhoA_path = os.path.join(
                                A_dir, f"rhoA_{rid.replace(':', '_')}.npy"
                            )
                            np.save(rhoA_path, rho_A.detach().cpu().numpy())
                        density_manifest["A_residues"].append(rid)
                    else:
                        charge_length_mismatch = True
                        xtb_success = False
                        xtb_used = False
                        charges_for_rid = None
                        print(
                            f"[WARN] xTB charges length mismatch for {rid}: "
                            f"len(charges)={len(charges)}, len(frag_atoms)={len(frag_atoms)}. "
                            "Using B-mode density only."
                        )
                        density_manifest["B_only_residues"].append(rid)
                else:
                    density_manifest["B_only_residues"].append(rid)
            else:
                density_manifest["B_only_residues"].append(rid)

            if save_densities and rhoA_path is None:
                rhoB_path = os.path.join(out_dir, f"rhoB_{rid.replace(':', '_')}.npy")
                np.save(rhoB_path, rho_B.detach().cpu().numpy())

            density_manifest["files"][rid] = {
                "rhoB": rhoB_path,
                "rhoA": rhoA_path,
            }

            density_manifest["modes"][rid] = {
                "B": {
                    "requested": gpu_density_mode,
                    "effective": eff_mode_B,
                    "fallback_from_vector": bool(fallback_B),
                },
                "A": {
                    "requested": gpu_density_mode if rid in select_A else None,
                    "effective": eff_mode_A,
                    "fallback_from_vector": bool(fallback_A) if eff_mode_A is not None else False,
                    "xtb_used": bool(xtb_used),
                    "xtb_success": bool(xtb_success),
                    "charge_length_mismatch": bool(charge_length_mismatch),
                },
            }

            # ----- Density-based features -----
            mask_view = density_ws.view_mask(nx, ny, nz)
            rho_local_view = density_ws.view_rho_local(nx, ny, nz)

            dens_feats = density_features_for_residue_gpu(
                rho=rho_final,
                X=X,
                Y=Y,
                Z=Z,
                ref_point=ref_point,
                res_atom_coords=coords_res,
                R_env=R_env,
                mask_ws=mask_view,
                rho_local_ws=rho_local_view,
                dist2_ws=dist2_view,
                tmp_ws=tmp_view,
                dx_ws=dx_view,
                dy_ws=dy_view,
                dz_ws=dz_view,
                rid=rid,
            )

            if dens_feats.get("curv_rho_failed", False):
                curv_rho_failed_residues.append(rid)

            # ----- A-mode charge features -----
            charge_feats_A = None
            if charges_for_rid is not None:
                charge_feats_A = charge_features_for_residue_from_fragment(
                    rid=rid,
                    atom_index_to_resid=atom_index_to_resid,
                    resid_to_atom_coords=resid_to_atom_coords,
                    frag_atoms=frag_atoms,
                    charges=charges_for_rid,
                )

            # ----- B-mode atom-based charge (Q_atom) -----
            partial_charge_B = compute_b_mode_charge_atom_based(
                res=res,
                rid=rid,
                resid_to_sasa2=resid_to_sasa2,
                mean_sasa=mean_sasa,
                sasa_alpha=sasa_alpha,
                env_scale_from_sasa=env_scale_from_sasa,
            )

            # ----- Field terms F^{(metal)}, F^{(P)}, F^{(hal)} -----
            F_metal, F_P_val, F_hal = compute_field_terms_for_residue(
                rid=rid,
                res_map=res_map,
                centers=centers,
                overlaps=overlaps_m,
            )

            feats = {
                "id": rid,
                "chain": res["chain"],
                "resseq": res["resseq"],
                "resname": res["resname"],
                "sasa": float(resid_to_sasa2.get(rid, 0.0)),
                "partial_charge_B": partial_charge_B,
                "F_metal": float(F_metal),
                "F_P": float(F_P_val),
                "F_hal": float(F_hal),
            }
            feats.update(dens_feats)
            if charge_feats_A is not None:
                feats.update(charge_feats_A)

            feats["gpu_density_B_requested"] = gpu_density_mode
            feats["gpu_density_B_effective"] = eff_mode_B
            feats["gpu_density_B_fallback_from_vector"] = bool(fallback_B)
            feats["gpu_density_A_requested"] = gpu_density_mode if rid in select_A else None
            feats["gpu_density_A_effective"] = eff_mode_A
            feats["gpu_density_A_fallback_from_vector"] = bool(fallback_A) if eff_mode_A is not None else False
            feats["xtb_A_used"] = bool(charge_feats_A is not None)
            feats["xtb_A_success"] = bool(xtb_success)
            feats["xtb_charge_length_mismatch"] = bool(charge_length_mismatch)

            env_features[rid] = feats

            # ----- Ep rows -----
            if "partial_charge" in feats:
                q_mean = float(feats["partial_charge"])
            else:
                q_mean = float(feats.get("partial_charge_B", 0.0))

            if "dipole_q" in feats:
                dip_vec = np.array(feats["dipole_q"], dtype=float)
            else:
                dip_vec = np.array(feats.get("dipole_rho", [0.0, 0.0, 0.0]), dtype=float)
            dip_norm = float(np.linalg.norm(dip_vec))

            curv_density = float(feats.get("curv_rho", 0.0))

            # combine field components as scalar external_field = ||(F_metal, F_P, F_hal)||
            F_vec = np.array([F_metal, F_P_val, F_hal], dtype=float)
            Efield = float(np.linalg.norm(F_vec))

            sasa = float(feats.get("sasa", 0.0))
            emb = residue_to_embedding(res["resname"]).astype(float)

            E_theory_row = np.concatenate(
                [
                    np.array([q_mean, dip_norm, curv_density, Efield, sasa], dtype=float),
                    emb,
                ]
            )

            # new 10D geometric block from overlaps / centers
            geom_block = geometric_block_from_overlaps(
                rid=rid,
                overlaps=overlaps_m,
                centers=centers,
                r_max=radius,
            )

            E_sim_row = np.concatenate([E_theory_row, geom_block])

            Ep_theory[idx, :] = E_theory_row
            Ep_sim[idx, :] = E_sim_row

            # ----- per-residue cleanup -----
            if 'frag_atom_indices' in locals():
                del frag_atom_indices
            if frag_atoms is not None:
                del frag_atoms
            del coords_res, coords_res_list, ref_point
            if rho_A is not None:
                del rho_A
            if "charges" in locals():
                del charges
                charges = None

        gc.collect()
        if torch is not None and torch.cuda.is_available():
            torch.cuda.empty_cache()
        log_gpu_usage(f"block_{block_start}_{block_end-1}_end", device)

    density_manifest["curv_rho_failed_residues"] = curv_rho_failed_residues

    dens_manifest_path = os.path.join(out_dir, "density_final.json")
    with open(dens_manifest_path, "w") as f:
        json.dump(density_manifest, f, indent=2)
    print(f"[INFO] density_final.json written: {dens_manifest_path}")

    env_path = os.path.join(out_dir, "env_features.json")
    with open(env_path, "w") as f:
        json.dump(env_features, f, indent=2)
    print(f"[INFO] env_features.json written: {env_path}")

    del res_map, overlaps_m, atom_index_to_resid, resid_to_atom_coords, resid_to_sasa2
    gc.collect()

    Ep_theory_path = os.path.join(out_dir, "Ep_theory.npy")
    Ep_sim_path = os.path.join(out_dir, "Ep_sim.npy")
    np.save(Ep_theory_path, Ep_theory)
    np.save(Ep_sim_path, Ep_sim)

    manifest = {
        "pdb": meta["pdb"],
        "n_residues": n_res,
        "residue_ids": res_ids,
        "E_theory_path": os.path.basename(Ep_theory_path),
        "E_sim_path": os.path.basename(Ep_sim_path),
        "feature_dim_theory": int(Ep_theory.shape[1]),
        "feature_dim_sim": int(Ep_sim.shape[1]),
        "features_theory": [
            "avg_charge",
            "dipole_norm",
            "curv_density",
            "external_field",  # ||F_metal, F_P, F_hal||
            "SASA",
            "emb_charge",
            "emb_hydrophobicity",
            "emb_size",
            "emb_aromaticity",
        ],
        "features_extra": [
            # 10D geometric neighbor block:
            "N_in",
            "N_out",
            "R_shell",
            "r_mean",
            "r_var",
            "linearity",
            "planarity",
            "isotropy",
            "H_mean",
            "K_gauss",
        ],
        "params": {
            "radius_contact": float(radius),
            "padding": float(padding),
            "grid_step": float(grid_step),
            "R_env": float(R_env),
            "sasa_alpha": float(sasa_alpha),
            "xtb": {
                "method": "gfn2",
                "charges": True,
                "binary": xtb_bin,
            },
            "block_size": int(block_size),
            "device": device,
            "gpu_density_mode_requested": gpu_density_mode,
            "gpu_density_vector_max_elems": int(MAX_VECTOR_ELEMENTS),
            "gpu_density_batch_max_elems": int(MAX_BATCH_ELEMENTS),
        },
    }
    mani_path = os.path.join(out_dir, "Ep_manifest.json")
    with open(mani_path, "w") as f:
        json.dump(manifest, f, indent=2)

    print(f"[DONE] E_p^theory written: {Ep_theory_path}")
    print(f"[DONE] E_p^sim written:    {Ep_sim_path}")
    print(f"[DONE] Ep manifest:        {mani_path}")


# ==============================
# 6b. Memory debug wrapper
# ==============================

def run_with_mem_monitor(
    pdb_path: str,
    out_dir: str = "pipeline_out",
    extra_args: Optional[List[str]] = None,
    python_bin: str = "python",
    interval: float = 5.0,
    max_seconds: Optional[float] = None,
):
    if not HAS_PSUTIL:
        raise RuntimeError("psutil is not available; cannot use run_with_mem_monitor.")

    if extra_args is None:
        extra_args = []

    script_path = os.path.abspath(__file__)
    cmd = [
        python_bin,
        script_path,
        pdb_path,
        "--out-dir",
        out_dir,
    ] + list(extra_args)

    print("Running:", " ".join(cmd))
    proc = subprocess.Popen(
        cmd,
        stdout=subprocess.PIPE,
        stderr=subprocess.STDOUT,
        text=True,
        bufsize=1,
    )

    p = psutil.Process(proc.pid)
    start = time.time()
    mem_log: List[Tuple[float, float]] = []

    try:
        while proc.poll() is None:
            now = time.time()
            t = now - start
            try:
                rss = p.memory_info().rss
                gb = rss / (1024 ** 3)
                mem_log.append((t, gb))
                print(f"[MEM] {t:7.1f} s : {gb:5.3f} GB")
            except psutil.Error:
                pass

            if proc.stdout is not None:
                try:
                    line = proc.stdout.readline()
                    if line:
                        print("[OUT]", line.rstrip())
                except Exception:
                    pass

            if max_seconds is not None and t > max_seconds:
                print(f"[WARN] max_seconds={max_seconds} exceeded; sending SIGINT to child...")
                try:
                    proc.send_signal(signal.SIGINT)
                except Exception:
                    pass

            time.sleep(interval)

    finally:
        if proc.poll() is None:
            try:
                proc.terminate()
            except Exception:
                pass

    print("Return code:", proc.returncode)
    return mem_log


# ==============================
# 7. CLI
# ==============================

def main():
    ap = argparse.ArgumentParser(
        description="Full 1–9 pipeline with optional GPU acceleration."
    )
    ap.add_argument("pdb", help="Input PDB or mmCIF file")
    ap.add_argument(
        "--out-dir",
        type=str,
        default="pipeline_out",
        help="Output directory for all artifacts",
    )
    ap.add_argument(
        "--radius",
        type=float,
        default=4.7,
        help="Sphere radius (Å) for residue-residue contact detection",
    )
    ap.add_argument(
        "--grid-step",
        type=float,
        default=0.8,
        help="Grid spacing (Å) for density grids",
    )
    ap.add_argument(
        "--padding",
        type=float,
        default=2.0,
        help="Padding (Å) around U_p box when building grids",
    )
    ap.add_argument(
        "--R-env",
        type=float,
        default=2.0,
        help="Environment radius (Å) for defining U_p around each residue",
    )
    ap.add_argument(
        "--sasa-alpha",
        type=float,
        default=0.4,
        help="Scaling factor alpha in SASA-based charge modulation",
    )
    ap.add_argument(
        "--xtb-bin",
        type=str,
        default="xtb",
        help="Path to xTB binary (for A-mode)",
    )
    ap.add_argument(
        "--A-list",
        type=str,
        default="",
        help='JSON list of residue IDs to run A-mode on, e.g. \'["A:201","A:175"]\'',
    )
    ap.add_argument(
        "--save-densities",
        action="store_true",
        help="If set, save rhoB_*.npy and rhoA_*.npy grids to disk",
    )
    ap.add_argument(
        "--no-pair-contacts",
        action="store_true",
        help="If set, do not store atom-level pair_contacts in meta.json.",
    )
    ap.add_argument(
        "--block-size",
        type=int,
        default=1024,
        help="Number of residues per density/feature block (for streaming).",
    )
    ap.add_argument(
        "--device",
        type=str,
        default="auto",
        choices=["auto", "cpu", "cuda"],
        help="Device for heavy 3D work (auto=CUDA if available, else CPU).",
    )
    ap.add_argument(
        "--gpu-density-mode",
        type=str,
        default="safe",
        choices=["safe", "batch", "vector"],
        help=(
            "GPU density accumulation mode: "
            "safe=atom-loop, batch=atom-tiled vector kernel, "
            "vector=full broadcast with thresholded fallback."
        ),
    )

    args = ap.parse_args()

    if args.A_list:
        try:
            select_A = json.loads(args.A_list)
            if not isinstance(select_A, list):
                raise ValueError
        except Exception:
            raise SystemExit(
                "A-list must be a JSON list of residue IDs, "
                'e.g. \'["A:201\",\"A:175"]\''
            )
    else:
        select_A = []

    if args.device == "auto":
        if HAS_TORCH and torch.cuda.is_available():
            device = "cuda"
        else:
            device = "cpu"
    elif args.device == "cuda":
        if not (HAS_TORCH and torch.cuda.is_available()):
            raise SystemExit("CUDA requested but torch.cuda.is_available() is False.")
        device = "cuda"
    else:
        device = "cpu"

    if not HAS_TORCH:
        raise SystemExit("torch is required for this GPU pipeline (even on CPU device).")

    print(f"[INFO] Using device: {device}")

    run_pipeline_gpu(
        pdb_path=args.pdb,
        out_dir=args.out_dir,
        radius=args.radius,
        grid_step=args.grid_step,
        padding=args.padding,
        xtb_bin=args.xtb_bin,
        select_A=select_A,
        save_densities=args.save_densities,
        R_env=args.R_env,
        sasa_alpha=args.sasa_alpha,
        no_pair_contacts=args.no_pair_contacts,
        block_size=args.block_size,
        device=device,
        gpu_density_mode=args.gpu_density_mode,
    )


if __name__ == "__main__":
    main()

'''

# ======================================================================
# 0-1) INTERNAL ENGINE MATERIALIZATION
# ======================================================================

import os
import sys
import re
import json
import shutil
import random
import zipfile
import subprocess
import itertools
import urllib.request
import platform
from datetime import datetime
from dataclasses import dataclass, field
from typing import Dict, List, Tuple, Optional, Any, Set

import numpy as np

_ENGINE_PATH = None  # will be set when first needed


def _ensure_dir(path: str):
    os.makedirs(path, exist_ok=True)


def _materialize_engine(workdir: str) -> str:
    """
    Write USER_ENGINE_CODE to a real .py file once, and reuse it.
    """
    global _ENGINE_PATH
    if _ENGINE_PATH is not None and os.path.exists(_ENGINE_PATH):
        return _ENGINE_PATH

    code = USER_ENGINE_CODE
    if code.strip() == "X" or code.strip() == "":
        raise RuntimeError(
            "USER_ENGINE_CODE still contains placeholder 'X'. "
            "Paste your full engine script into USER_ENGINE_CODE."
        )

    # Write engine into workdir (so everything stays under benchmark_workdir)
    _ensure_dir(workdir)
    engine_path = os.path.join(workdir, "_embedded_engine.py")
    with open(engine_path, "w", encoding="utf-8") as f:
        f.write(code)
    _ENGINE_PATH = engine_path
    print(f"[ENGINE] Embedded engine written to {engine_path}")
    return _ENGINE_PATH


# ==============================================================================
# 1) DEPENDENCY BOOTSTRAP (pip-install if missing)
# ==============================================================================
# This script is designed to run "out of the box" on Colab-like environments.
# If a dependency is missing, we install it via pip at runtime.
#
# NOTE:
# - Installing torch from pip can be large/slow. Colab usually ships with torch preinstalled.
# - If you are in a locked-down environment without pip/network, installs will fail.

import importlib

def _pip_install(pkgs: List[str]):
    cmd = [sys.executable, "-m", "pip", "install", "-q"] + pkgs
    print(f"[PIP] Installing: {' '.join(pkgs)}")
    subprocess.run(cmd, check=True)

def _ensure_import(module: str, pip_name: Optional[str] = None):
    try:
        importlib.import_module(module)
        return
    except Exception as e:
        pkg = pip_name or module.split(".")[0]
        _pip_install([pkg])
        importlib.invalidate_caches()
        importlib.import_module(module)

# Core runtime deps used by the embedded engine and/or evaluation suite
_ensure_import("numpy", "numpy")              # usually present
_ensure_import("mdtraj", "mdtraj")
_ensure_import("scipy", "scipy")
_ensure_import("sklearn", "scikit-learn")
_ensure_import("psutil", "psutil")
_ensure_import("torch", "torch")
_ensure_import("Bio", "biopython")            # Biopython (for optional global alignment)

# Now import the symbols we actually use
import mdtraj as mdt

from sklearn.model_selection import GroupKFold
from sklearn.ensemble import RandomForestClassifier
from sklearn.impute import SimpleImputer
from sklearn.metrics import roc_auc_score, average_precision_score
from sklearn.neighbors import KDTree as SK_KDTree

try:
    from scipy.spatial import cKDTree as SCIPY_KDTree
    _KDTREE_BACKEND = "scipy.cKDTree"
except Exception:
    SCIPY_KDTree = None
    _KDTREE_BACKEND = "sklearn.KDTree"


# 2) BASIC HELPERS
# ==============================================================================


def _collect_run_environment(extra: Optional[Dict[str, Any]] = None) -> Dict[str, Any]:
    """Collect lightweight execution-environment metadata for reproducibility logs."""
    env: Dict[str, Any] = {}
    env["timestamp_local"] = datetime.now().isoformat(timespec="seconds")
    env["python_version"] = sys.version.replace("\n", " ")
    env["python_executable"] = sys.executable
    env["platform"] = platform.platform()
    env["machine"] = platform.machine()
    env["processor"] = platform.processor()
    env["cpu_count"] = os.cpu_count()

    # Key package versions (best-effort)
    versions: Dict[str, Optional[str]] = {}
    try:
        from importlib.metadata import version as _pkg_version  # type: ignore
        def _v(dist: str) -> Optional[str]:
            try:
                return _pkg_version(dist)
            except Exception:
                return None
    except Exception:
        _pkg_version = None  # type: ignore
        def _v(dist: str) -> Optional[str]:
            return None

    for dist in ["numpy", "scipy", "scikit-learn", "mdtraj", "torch", "biopython", "psutil"]:
        versions[dist] = _v(dist)
    env["packages"] = versions

    # Torch / CUDA details (best-effort)
    torch_info: Dict[str, Any] = {}
    try:
        import torch  # type: ignore
        torch_info["torch_version"] = getattr(torch, "__version__", None)
        torch_info["cuda_available"] = bool(torch.cuda.is_available())
        torch_info["cuda_version"] = getattr(torch.version, "cuda", None)
        try:
            torch_info["cudnn_version"] = int(torch.backends.cudnn.version()) if torch.backends.cudnn.is_available() else None
        except Exception:
            torch_info["cudnn_version"] = None
        if torch.cuda.is_available():
            try:
                torch_info["gpu_count"] = int(torch.cuda.device_count())
                torch_info["gpu_name_0"] = torch.cuda.get_device_name(0)
            except Exception:
                pass
            try:
                props = torch.cuda.get_device_properties(0)
                torch_info["gpu_cc_0"] = f"{props.major}.{props.minor}"
                torch_info["gpu_total_mem_mb_0"] = int(props.total_memory / 1024**2)
            except Exception:
                pass
    except Exception as e:
        torch_info["error"] = f"{type(e).__name__}: {e}"
    env["torch"] = torch_info

    # Determinism-relevant env vars (best-effort)
    det_vars = ["PYTHONHASHSEED", "CUBLAS_WORKSPACE_CONFIG", "CUDA_VISIBLE_DEVICES",
                "OMP_NUM_THREADS", "MKL_NUM_THREADS"]
    env["env_vars"] = {k: os.environ.get(k) for k in det_vars if os.environ.get(k) is not None}

    # Internal backend choices
    try:
        env["kdtree_backend"] = _KDTREE_BACKEND  # type: ignore
    except Exception:
        pass

    if extra:
        env["extra"] = extra
    return env

def _read_json(p: str) -> dict:
    with open(p, "r") as f:
        return json.load(f)


def _write_json(p: str, obj: dict):
    with open(p, "w") as f:
        json.dump(obj, f, indent=2)


def _set_reproducible(seed: int = 42):
    os.environ["PYTHONHASHSEED"] = str(seed)
    random.seed(seed)
    np.random.seed(seed)


def _is_url(s: str) -> bool:
    return isinstance(s, str) and (s.startswith("http://") or s.startswith("https://"))


def _looks_like_pdb_id(s: str) -> bool:
    return bool(re.fullmatch(r"[0-9][A-Za-z0-9]{3}", str(s).strip()))


def _download(url: str, out_path: str) -> str:
    _ensure_dir(os.path.dirname(out_path) or ".")
    if not os.path.exists(out_path):
        print(f"[DL] {url} -> {out_path}")
        urllib.request.urlretrieve(url, out_path)
    return out_path


def _resolve_path_or_download(spec: str, cache_dir: str, prefer_ext: Optional[str] = None) -> str:
    if spec is None or str(spec).strip() == "":
        raise ValueError("Empty path spec.")
    s = str(spec).strip()
    if os.path.exists(s):
        return os.path.abspath(s)
    _ensure_dir(cache_dir)

    if _is_url(s):
        base = os.path.basename(s.split("?")[0])
        if (prefer_ext is not None) and (not base.lower().endswith(prefer_ext.lower())):
            base = base + prefer_ext
        out_path = os.path.join(cache_dir, base)
        return _download(s, out_path)

    if _looks_like_pdb_id(s):
        pdb_id = s.upper()
        if prefer_ext is None:
            prefer_ext = ".cif"
        if prefer_ext.lower() == ".pdb":
            url = f"https://files.rcsb.org/download/{pdb_id}.pdb"
            out_path = os.path.join(cache_dir, f"{pdb_id}.pdb")
            return _download(url, out_path)
        else:
            url = f"https://files.rcsb.org/download/{pdb_id}.cif"
            out_path = os.path.join(cache_dir, f"{pdb_id}.cif")
            return _download(url, out_path)

    raise FileNotFoundError(f"Cannot resolve path: {spec} (not local, not URL, not PDB id).")


# ==============================================================================
# 3) RESTORE FROM colab_all.zip
# ==============================================================================

def restore_from_colab_all(zip_path: str, dst_root: Optional[str] = None):
    """
    Restore colab_all.zip into a real directory.

    - If dst_root is None:
        extract into the directory where zip_path actually lives.
    - If dst_root is given:
        extract into that directory explicitly.

    Also handles the common case where the archive contains a top-level
    'content/' folder; in that case, its children are moved one level up.
    """
    if not os.path.exists(zip_path):
        raise FileNotFoundError(f"{zip_path} is not found.")

    if dst_root is None:
        dst_root = os.path.dirname(os.path.abspath(zip_path)) or "."

    abs_zip = os.path.abspath(zip_path)
    abs_dst = os.path.abspath(dst_root)
    print(f"[INFO] Restoring from {abs_zip} to {abs_dst}")
    with zipfile.ZipFile(zip_path, "r") as z:
        z.extractall(dst_root)
    print("[INFO] Restore complete.")

    # Handle nested 'content' directory (e.g., zip created from /content)
    nested = os.path.join(abs_dst, "content")
    if os.path.isdir(nested):
        print(f"[INFO] Detected nested 'content' folder inside zip \u2192 flattening into {abs_dst}")
        for entry in os.listdir(nested):
            src = os.path.join(nested, entry)
            dst = os.path.join(abs_dst, entry)
            if os.path.exists(dst):
                print(f"[WARN] Flatten skip: {dst} already exists, keep existing.")
                continue
            shutil.move(src, dst)


# ==============================================================================
# 3b) RESTORE FROM benchmark5.5.tgz (DB5.5-only mode)
# ==============================================================================

def restore_from_db55_tgz(
    tgz_path: str,
    dataset_unpack_dir: str,
    tmp_extract_root: Optional[str] = None,
):
    """
    Make the script runnable with ONLY a DB5.5-style .tgz.

    Expected common layout inside tgz:
        benchmark5.5/structures/*.pdb
    but we do NOT hardcode it; we search for a folder named 'structures'
    that contains *.pdb files.

    The function copies *.pdb into `dataset_unpack_dir` so the existing
    `build_cases_from_db55_like_dir()` logic works unchanged.
    """
    if not os.path.exists(tgz_path):
        raise FileNotFoundError(f"{tgz_path} is not found.")

    os.makedirs(dataset_unpack_dir, exist_ok=True)

    if tmp_extract_root is None:
        tmp_extract_root = os.path.join(os.path.dirname(os.path.abspath(dataset_unpack_dir)), "_tmp_db55_extract")
    os.makedirs(tmp_extract_root, exist_ok=True)

    print(f"[INFO] Extracting TGZ: {os.path.abspath(tgz_path)} -> {os.path.abspath(tmp_extract_root)}")
    import tarfile
    with tarfile.open(tgz_path, "r:gz") as tf:
        try:

            tf.extractall(tmp_extract_root, filter="data")

        except TypeError:

            tf.extractall(tmp_extract_root)

    # Find a 'structures' directory containing pdbs
    structures_dir = None
    for root, dirs, files in os.walk(tmp_extract_root):
        if os.path.basename(root) == "structures":
            pdbs = [f for f in files if f.lower().endswith(".pdb") and not os.path.basename(f).startswith("._")]
            if len(pdbs) > 0:
                structures_dir = root
                break

    if structures_dir is None:
        # Fallback: any directory with pdbs
        for root, dirs, files in os.walk(tmp_extract_root):
            pdbs = [f for f in files if f.lower().endswith(".pdb") and not os.path.basename(f).startswith("._")]
            if len(pdbs) > 0:
                structures_dir = root
                break

    if structures_dir is None:
        raise RuntimeError(
            "Could not find any *.pdb files inside the extracted tgz.\n"
            "Expected something like 'benchmark5.5/structures/*.pdb'."
        )

    # Copy pdb files into dataset_unpack_dir
    copied = 0
    for fn in os.listdir(structures_dir):
        if not fn.lower().endswith(".pdb"):
            continue
        if fn.startswith("._"):  # macOS metadata files
            continue
        src = os.path.join(structures_dir, fn)
        dst = os.path.join(dataset_unpack_dir, fn)
        shutil.copy2(src, dst)
        copied += 1

    print(f"[INFO] Copied {copied} pdb files from {structures_dir} -> {dataset_unpack_dir}")

# ==============================================================================
# 4) MDTRAJ RESIDUE HELPERS
# ==============================================================================

_AA3_TO_AA1 = {
    "ALA":"A","ARG":"R","ASN":"N","ASP":"D","CYS":"C","GLN":"Q","GLU":"E","GLY":"G",
    "HIS":"H","ILE":"I","LEU":"L","LYS":"K","MET":"M","PHE":"F","PRO":"P","SER":"S",
    "THR":"T","TRP":"W","TYR":"Y","VAL":"V",
    "MSE":"M","SEC":"U","PYL":"O","ASX":"B","GLX":"Z","XAA":"X","XLE":"J",
}


def _get_chain_id_mdtraj(chain) -> str:
    if hasattr(chain, "chain_id") and chain.chain_id is not None:
        return str(chain.chain_id)
    if hasattr(chain, "id") and chain.id is not None:
        return str(chain.id)
    return str(chain.index)


def _get_insertion_code_mdtraj(res) -> str:
    raw = None
    if hasattr(res, "insertion_code"):
        raw = res.insertion_code
    elif hasattr(res, "insertionCode"):
        raw = res.insertionCode
    if raw is None:
        raw = ""
    icode = str(raw).strip()
    if icode == "None":
        icode = ""
    return icode


def _is_protein_residue_mdtraj(res) -> bool:
    rn = str(res.name).strip().upper()
    return rn in _AA3_TO_AA1


def _rid_from_mdtraj_res(res) -> str:
    chain_id = _get_chain_id_mdtraj(res.chain)
    resseq = res.resSeq
    icode = _get_insertion_code_mdtraj(res)
    suffix = icode if icode else ""
    return f"{chain_id}:{resseq}{suffix}"


def _atom_is_heavy(atom) -> bool:
    if hasattr(atom, "element") and atom.element is not None:
        try:
            return atom.element.symbol != "H"
        except Exception:
            pass
    nm = str(atom.name).strip().upper()
    return not nm.startswith("H")


# ==============================================================================
# 5) BOUND COMPLEX BUILDER (L_b + R_b → complex)
# ==============================================================================

def make_bound_complex(l_b_path: str, r_b_path: str, out_path: str) -> str:
    def read_atom_lines(p):
        lines = []
        with open(p, "r") as f:
            for line in f:
                if line.startswith(("ATOM", "HETATM")):
                    lines.append(line.rstrip("\n"))
        return lines

    _ensure_dir(os.path.dirname(out_path) or ".")
    l_lines = read_atom_lines(l_b_path)
    r_lines = read_atom_lines(r_b_path)

    with open(out_path, "w") as w:
        for line in l_lines:
            w.write(line + "\n")
        w.write("TER\n")
        for line in r_lines:
            w.write(line + "\n")
        w.write("END\n")
    return out_path


# ==============================================================================
# 6) SEQUENCE ALIGNMENT
# ==============================================================================

def _global_align(a: str, b: str, match: int = 2, mismatch: int = -1, gap: int = -2) -> Tuple[str, str, int]:
    try:
        from Bio import pairwise2
    except ImportError:
        # Fallback: trivial alignment (no score)
        return a, b, 0
    alignments = pairwise2.align.localms(a, b, match, mismatch, -10, -0.5)
    if not alignments:
        return a, b, 0
    best_aln = alignments[0]
    return best_aln.seqA, best_aln.seqB, int(best_aln.score)


def _alignment_identity_and_coverage(a_seq, b_seq):
    """
    Compute identity and coverage for a pairwise alignment.

    This function is robust to two kinds of inputs:
      1) Already-aligned sequences (same length, with '-' gaps)
      2) Raw sequences (different length, no guaranteed gaps)

    In case (2), it performs a local alignment first and then
    computes identity / coverage on the aligned result.
    """

    # Case 1: already-aligned strings with same length
    if len(a_seq) == len(b_seq):
        a_aln, b_aln = a_seq, b_seq
    else:
        # Case 2: treat as raw sequences → local alignment
        # remove any existing gaps just in case
        a_raw = a_seq.replace("-", "")
        b_raw = b_seq.replace("-", "")

        if not a_raw or not b_raw:
            return 0.0, 0.0


        try:
            from Bio import pairwise2
        except ImportError:
            # Biopython not installed: no reliable alignment possible
            return 0.0, 0.0

        alns = pairwise2.align.localms(a_raw, b_raw, 2, -1, -10, -0.5)
        if not alns:
            return 0.0, 0.0

        a_aln, b_aln, *_ = alns[0]

    # From here on: assume a_aln, b_aln are aligned strings of same length
    if len(a_aln) != len(b_aln):
        # extremely pathological; bail out instead of crashing
        return 0.0, 0.0

    matches = 0
    aligned_len = 0
    core_len_a = 0
    core_len_b = 0

    for aa, bb in zip(a_aln, b_aln):
        if aa != '-':
            core_len_a += 1
        if bb != '-':
            core_len_b += 1

        # only positions where both are non-gap residues count as "aligned"
        if aa != '-' and bb != '-':
            aligned_len += 1
            if aa == bb:
                matches += 1

    if aligned_len == 0:
        return 0.0, 0.0

    core_len = min(core_len_a, core_len_b)
    if core_len == 0:
        return 0.0, 0.0

    ident = matches / aligned_len
    cov = aligned_len / core_len

    return ident, cov


# ==============================================================================
# 7) CHAIN SEQUENCE EXTRACTION
# ==============================================================================

def _extract_chain_sequences(struct_path: str, protein_only: bool = True) -> Dict[str, Dict[str, Any]]:
    traj = mdt.load(struct_path)
    topo = traj.topology

    chains: Dict[str, Dict[str, Any]] = {}
    for res in topo.residues:
        if protein_only and (not _is_protein_residue_mdtraj(res)):
            continue
        c = _get_chain_id_mdtraj(res.chain)
        chains.setdefault(c, {"seq": [], "rids": []})
        aa3 = str(res.name).strip().upper()
        aa1 = _AA3_TO_AA1.get(aa3, "X")
        chains[c]["seq"].append(aa1)
        chains[c]["rids"].append(_rid_from_mdtraj_res(res))

    for c in list(chains.keys()):
        chains[c]["seq"] = "".join(chains[c]["seq"])
        if len(chains[c]["seq"]) == 0:
            del chains[c]
    return chains


# ==============================================================================
# 8) SYMMETRY DETECTION
# ==============================================================================

@dataclass
class SymmetryPolicy:
    min_identity: float = 0.95
    min_coverage: float = 0.90


def detect_symmetry_groups(chains: Dict[str, Dict[str, Any]], sym_policy: SymmetryPolicy) -> Dict[str, int]:
    chain_ids = list(chains.keys())
    n = len(chain_ids)
    if n == 0:
        return {}

    adj = [[False] * n for _ in range(n)]
    for i in range(n):
        adj[i][i] = True

    for i in range(n):
        for j in range(i + 1, n):
            a = chains[chain_ids[i]]["seq"]
            b = chains[chain_ids[j]]["seq"]
            a_aln, b_aln, _ = _global_align(a, b)
            ident, cov = _alignment_identity_and_coverage(a_aln, b_aln)
            eq = (ident >= sym_policy.min_identity) and (cov >= sym_policy.min_coverage)
            adj[i][j] = eq
            adj[j][i] = eq

    group_id = 0
    visited = [False] * n
    chain_to_group: Dict[str, int] = {}
    for i in range(n):
        if visited[i]:
            continue
        stack = [i]
        visited[i] = True
        members = []
        while stack:
            u = stack.pop()
            members.append(u)
            for v in range(n):
                if (not visited[v]) and adj[u][v]:
                    visited[v] = True
                    stack.append(v)
        for idx in members:
            chain_to_group[chain_ids[idx]] = group_id
        group_id += 1
    return chain_to_group


# ==============================================================================
# 9) CHAIN ASSIGNMENT + RESIDUE MAP (BOUND → UNBOUND)
# ==============================================================================
from dataclasses import dataclass

@dataclass
class ChainAssignPolicy:
    min_chain_identity: float = 0.70
    min_chain_coverage: float = 0.10
    exhaustive_max_n: int = 0


def assign_chains_bound_to_unbound(
    bound_chains: Dict[str, Dict[str, Any]],
    unbound_chains: Dict[str, Dict[str, Any]],
    policy: ChainAssignPolicy,
    chain_hints: Optional[Dict[str, str]] = None,
) -> Tuple[Dict[str, str], Dict[str, Any]]:
    chain_hints = chain_hints or {}
    b_ids = list(bound_chains.keys())
    u_ids = list(unbound_chains.keys())

    score = {}
    detail = {}
    for bc in b_ids:
        for uc in u_ids:
            a = bound_chains[bc]["seq"]
            b = unbound_chains[uc]["seq"]
            a_aln, b_aln, _ = _global_align(a, b)
            ident, cov = _alignment_identity_and_coverage(a_aln, b_aln)
            w = float(ident * cov)
            score[(bc, uc)] = w
            detail[(bc, uc)] = {"identity": float(ident), "coverage": float(cov), "weight": float(w)}

    mapping: Dict[str, str] = {}
    used_u = set()
    forced = []
    for bc, uc in (chain_hints or {}).items():
        if bc in bound_chains and uc in unbound_chains:
            mapping[bc] = uc
            used_u.add(uc)
            forced.append((bc, uc))

    remaining_b = [bc for bc in b_ids if bc not in mapping]
    remaining_u = [uc for uc in u_ids if uc not in used_u]

    def valid_pair(bc, uc) -> bool:
        d = detail[(bc, uc)]
        return (d["identity"] >= policy.min_chain_identity) and (d["coverage"] >= policy.min_chain_coverage)

    best_assign = None
    best_weight = -1.0

    if len(remaining_b) > 0 and len(remaining_u) > 0:
        if (len(remaining_b) <= policy.exhaustive_max_n) and (len(remaining_u) <= policy.exhaustive_max_n):
            for perm in itertools.permutations(remaining_u, r=min(len(remaining_b), len(remaining_u))):
                wsum = 0.0
                ok = True
                local = {}
                for i, bc in enumerate(remaining_b[:len(perm)]):
                    uc = perm[i]
                    if not valid_pair(bc, uc):
                        ok = False
                        break
                    wsum += score[(bc, uc)]
                    local[bc] = uc
                if ok and wsum > best_weight:
                    best_weight = wsum
                    best_assign = local
        else:
            pairs = []
            for bc in remaining_b:
                for uc in remaining_u:
                    pairs.append((score[(bc, uc)], bc, uc))
            pairs.sort(reverse=True, key=lambda x: x[0])
            local = {}
            used = set()
            for w, bc, uc in pairs:
                if bc in local or uc in used:
                    continue
                if not valid_pair(bc, uc):
                    continue
                local[bc] = uc
                used.add(uc)
            best_assign = local
            best_weight = sum(score[(bc, uc)] for bc, uc in local.items())

    if best_assign:
        for bc, uc in best_assign.items():
            mapping[bc] = uc
            used_u.add(uc)

    ambiguity = []
    for bc in b_ids:
        cands = sorted(
            [{"unbound": uc, **detail[(bc, uc)]} for uc in u_ids],
            reverse=True, key=lambda x: x["weight"]
        )
        if len(cands) >= 2:
            topw = cands[0]["weight"]
            secw = cands[1]["weight"]
            if topw > 0 and (secw / topw) >= 0.95:
                ambiguity.append({
                    "bound_chain": bc,
                    "top": cands[0]["unbound"],
                    "top_weight": float(topw),
                    "second": cands[1]["unbound"],
                    "second_weight": float(secw),
                })

    unmatched_bound = [bc for bc in b_ids if bc not in mapping]
    matched_unbound = set(mapping.values())
    unmatched_unbound = [uc for uc in u_ids if uc not in matched_unbound]

    report = {
        "forced_hints": forced,
        "unmatched_bound_chains": unmatched_bound,
        "unmatched_unbound_chains": unmatched_unbound,
        "ambiguity": ambiguity,
        "pair_details_top": {
            bc: sorted(
                [{"unbound": uc, **detail[(bc, uc)]} for uc in u_ids],
                reverse=True, key=lambda x: x["weight"]
            )[:3]
            for bc in b_ids
        },
    }
    return mapping, report


def build_bound_to_unbound_residue_map(
    bound_path: str,
    unbound_path: str,
    chain_hints: Optional[Dict[str, str]] = None,
    protein_only: bool = True,
    assign_policy: Optional[ChainAssignPolicy] = None,
) -> Tuple[Dict[str, str], Dict[str, Any]]:
    assign_policy = assign_policy or ChainAssignPolicy()
    bound_chains = _extract_chain_sequences(bound_path, protein_only=protein_only)
    unbound_chains = _extract_chain_sequences(unbound_path, protein_only=protein_only)

    chain_map, chain_report = assign_chains_bound_to_unbound(
        bound_chains=bound_chains,
        unbound_chains=unbound_chains,
        policy=assign_policy,
        chain_hints=chain_hints,
    )

    b2u: Dict[str, str] = {}
    dropped_chains = []
    for bc, uc in chain_map.items():
        bseq = bound_chains[bc]["seq"]
        useq = unbound_chains[uc]["seq"]
        a_aln, b_aln, _ = _global_align(bseq, useq)
        ident, cov = _alignment_identity_and_coverage(a_aln, b_aln)
        if ident < assign_policy.min_chain_identity or cov < assign_policy.min_chain_coverage:
            dropped_chains.append({"bound_chain": bc, "unbound_chain": uc, "identity": ident, "coverage": cov})
            continue

        brids = bound_chains[bc]["rids"]
        urids = unbound_chains[uc]["rids"]

        bi = 0
        ui = 0
        for x, y in zip(a_aln, b_aln):
            if x != "-" and y != "-":
                if bi < len(brids) and ui < len(urids):
                    b2u[brids[bi]] = urids[ui]
                bi += 1
                ui += 1
            elif x != "-" and y == "-":
                bi += 1
            elif x == "-" and y != "-":
                ui += 1

    report = {
        "bound_to_unbound_chain_map": chain_map,
        "chain_assignment": chain_report,
        "dropped_chains_low_quality": dropped_chains,
        "n_bound_chains_total": int(len(bound_chains)),
        "n_unbound_chains_total": int(len(unbound_chains)),
    }
    return b2u, report


def map_bound_labels_to_unbound_indices(
    bound_positive_rids: List[str],
    bound_to_unbound_rid: Dict[str, str],
    unbound_manifest_rids: List[str],
) -> Tuple[np.ndarray, List[str], Dict[str, Any]]:
    rid_to_idx = {rid: i for i, rid in enumerate(unbound_manifest_rids)}
    y = np.zeros((len(unbound_manifest_rids),), dtype=np.int32)

    mapped = []
    dropped = []
    for b_rid in bound_positive_rids:
        u_rid = bound_to_unbound_rid.get(b_rid)
        if u_rid is None:
            dropped.append({"bound": b_rid, "reason": "no_bound_to_unbound_map"})
            continue
        idx = rid_to_idx.get(u_rid)
        if idx is None:
            dropped.append({"bound": b_rid, "unbound": u_rid, "reason": "unbound_not_in_manifest"})
            continue
        y[idx] = 1
        mapped.append({"bound": b_rid, "unbound": u_rid, "idx": int(idx)})

    unbound_positive_rids = [unbound_manifest_rids[i] for i in np.where(y == 1)[0].tolist()]

    report = {
        "n_bound_labels": int(len(bound_positive_rids)),
        "n_mapped": int(len(mapped)),
        "n_dropped": int(len(dropped)),
        "mapped_examples": mapped[:20],
        "dropped_examples": dropped[:20],
    }
    return y, unbound_positive_rids, report


# ==============================================================================
# 10) AUTO-LABEL PPI INTERFACE FROM BOUND COMPLEX
# ==============================================================================

@dataclass
class InterfaceLabelPolicy:
    cutoff_angstrom: float = 5.0
    coarse_margin_angstrom: float = 4.0
    protein_only: bool = True
    scheme: str = "closest-heavy"


def infer_ppi_interface_labels_from_bound(
    bound_structure_path: str,
    target_bound_chains: Set[str],
    policy: InterfaceLabelPolicy,
) -> Tuple[List[str], Dict[str, Any]]:
    traj = mdt.load(bound_structure_path)
    topo = traj.topology

    target_res = []
    other_res = []
    for res in topo.residues:
        if policy.protein_only and (not _is_protein_residue_mdtraj(res)):
            continue
        cid = _get_chain_id_mdtraj(res.chain)
        if cid in target_bound_chains:
            target_res.append(res)
        else:
            other_res.append(res)

    if len(target_res) == 0 or len(other_res) == 0:
        return [], {
            "error": "empty_target_or_partner_residues_after_filter",
            "n_target_res": int(len(target_res)),
            "n_other_res": int(len(other_res)),
            "target_bound_chains": sorted(list(target_bound_chains)),
        }

    xyz = traj.xyz[0]

    def rep_coord(res) -> np.ndarray:
        ca = None
        first_heavy = None
        first_any = None
        for atom in res.atoms:
            if first_any is None:
                first_any = atom.index
            if _atom_is_heavy(atom) and first_heavy is None:
                first_heavy = atom.index
            if str(atom.name).strip().upper() == "CA":
                ca = atom.index
                break
        idx = ca if ca is not None else (first_heavy if first_heavy is not None else first_any)
        return xyz[idx]

    target_coords = np.stack([rep_coord(r) for r in target_res], axis=0)
    other_coords = np.stack([rep_coord(r) for r in other_res], axis=0)

    radius_nm = (policy.cutoff_angstrom + policy.coarse_margin_angstrom) / 10.0

    if SCIPY_KDTree is not None:
        tree = SCIPY_KDTree(other_coords)
        neighs = tree.query_ball_point(target_coords, r=radius_nm)
        candidate_pairs = []
        for i_t, nbrs in enumerate(neighs):
            t_res = target_res[i_t]
            for j in nbrs:
                o_res = other_res[j]
                candidate_pairs.append((t_res.index, o_res.index))
    else:
        tree = SK_KDTree(other_coords)
        neighs = tree.query_radius(target_coords, r=radius_nm, return_distance=False)
        candidate_pairs = []
        for i_t, nbrs in enumerate(neighs):
            t_res = target_res[i_t]
            for j in neighs:
                o_res = other_res[int(j)]
                candidate_pairs.append((t_res.index, o_res.index))

    if len(candidate_pairs) == 0:
        return [], {
            "cutoff_angstrom": float(policy.cutoff_angstrom),
            "coarse_margin_angstrom": float(policy.coarse_margin_angstrom),
            "n_candidate_pairs": 0,
            "n_interface_target_res": 0,
            "note": "No coarse candidates found.",
            "backend_kdtree": _KDTREE_BACKEND,
        }

    dists_nm = mdt.compute_contacts(traj, contacts=candidate_pairs, scheme=policy.scheme)[0][0]
    dists_A = dists_nm * 10.0

    interface_target_indices = set()
    cutoff = float(policy.cutoff_angstrom)
    for (pair, dA) in zip(candidate_pairs, dists_A.tolist()):
        if dA <= cutoff:
            t_res_idx, _ = pair
            interface_target_indices.add(t_res_idx)

    interface_rids = []
    for res in topo.residues:
        if res.index in interface_target_indices:
            if policy.protein_only and (not _is_protein_residue_mdtraj(res)):
                continue
            interface_rids.append(_rid_from_mdtraj_res(res))

    report = {
        "cutoff_angstrom": float(policy.cutoff_angstrom),
        "coarse_margin_angstrom": float(policy.coarse_margin_angstrom),
        "scheme": str(policy.scheme),
        "protein_only": bool(policy.protein_only),
        "target_bound_chains": sorted(list(target_bound_chains)),
        "n_target_res": int(len(target_res)),
        "n_other_res": int(len(other_res)),
        "n_candidate_pairs": int(len(candidate_pairs)),
        "n_interface_target_res": int(len(set(interface_rids))),
        "backend_kdtree": _KDTREE_BACKEND,
    }
    # FIXED: return tuple (list, dict), not sorted(..., report)
    return sorted(list(set(interface_rids))), report


# ==============================================================================
# 11) SOLVENT REMOVAL FROM ENGINE OUTPUTS
# ==============================================================================

def _build_resname_map_from_mdtraj(struct_path: str) -> Dict[str, str]:
    traj = mdt.load(struct_path)
    topo = traj.topology
    m: Dict[str, str] = {}
    for res in topo.residues:
        m[_rid_from_mdtraj_res(res)] = str(res.name).strip().upper()
    return m


def _build_resname_map_from_meta_json(meta_json_path: str) -> Dict[str, str]:
    meta = _read_json(meta_json_path)
    if isinstance(meta, dict) and "residues" in meta and isinstance(meta["residues"], list):
        out = {}
        for r in meta["residues"]:
            if isinstance(r, dict) and ("id" in r) and ("resname" in r):
                out[str(r["id"])] = str(r["resname"]).strip().upper()
        if out:
            return out
    if isinstance(meta, dict) and "residue_map" in meta and isinstance(meta["residue_map"], dict):
        return {str(k): str(v).strip().upper() for k, v in meta["residue_map"].items()}
    raise ValueError(f"Unrecognized meta.json schema: {meta_json_path}")


def filter_solvent_from_engine_outputs(
    X: np.ndarray,
    residue_ids: List[str],
    solvent_codes: List[str],
    unbound_structure_path: Optional[str],
    meta_json_path: Optional[str],
) -> Tuple[np.ndarray, List[str], np.ndarray, Dict[str, Any]]:
    solvent_set = {c.strip().upper() for c in solvent_codes}
    resname_map: Dict[str, str] = {}
    source = None

    if meta_json_path is not None and os.path.exists(meta_json_path):
        try:
            resname_map = _build_resname_map_from_meta_json(meta_json_path)
            source = "meta.json"
        except Exception:
            resname_map = {}

    if not resname_map and unbound_structure_path is not None:
        try:
            resname_map = _build_resname_map_from_mdtraj(unbound_structure_path)
            source = "mdtraj_structure"
        except Exception:
            resname_map = {}

    keep = np.ones((len(residue_ids),), dtype=bool)
    unknown = 0
    removed = 0
    for i, rid in enumerate(residue_ids):
        rn = resname_map.get(rid)
        if rn is None:
            unknown += 1
            continue
        if rn in solvent_set:
            keep[i] = False
            removed += 1

    X2 = X[keep]
    r2 = [rid for rid, k in zip(residue_ids, keep) if k]
    report = {
        "solvent_filter_applied": True,
        "source": source,
        "n_total": int(len(residue_ids)),
        "n_removed": int(removed),
        "n_unknown_resname": int(unknown),
        "n_retained": int(len(r2)),
        "solvent_codes": list(solvent_set),
    }
    return X2, r2, keep, report


# ==============================================================================
# 12) POLICY P (GROUPKFOLD AUROC/AUPRC)
# ==============================================================================

@dataclass
class PolicyP:
    K_TARGET: int = 5
    K_MIN: int = 2
    MIN_VALID_FOLDS: int = 2
    RF_RANDOM_STATE: int = 42
    RF_N_ESTIMATORS: int = 100
    IMPUTER_STRATEGY: str = "mean"
    RF_N_JOBS: int = 1


def eval_metrics_grouped_policy_p(
    X: np.ndarray,
    y: np.ndarray,
    groups: np.ndarray,
    policy: PolicyP,
) -> Tuple[Dict[str, Any], Dict[str, Any]]:
    y = np.asarray(y).astype(int)
    groups = np.asarray(groups)

    pos_groups = np.unique(groups[y == 1])
    n_pos_groups = int(len(pos_groups))

    info = {
        "K_target": int(policy.K_TARGET),
        "K_min": int(policy.K_MIN),
        "min_valid_folds": int(policy.MIN_VALID_FOLDS),
        "n_pos_groups": int(n_pos_groups),
        "K_used": None,
        "n_total_folds": 0,
        "n_valid_folds": 0,
        "n_skipped_train_single_class": 0,
        "n_skipped_test_single_class": 0,
        "reason": "",
    }

    if n_pos_groups < 2:
        info["reason"] = "Too few positive site-groups (<2) for grouped split."
        return {
            "AUROC_mean": float("nan"), "AUROC_std": float("nan"), "fold_AUROCs": [],
            "AUPRC_mean": float("nan"), "AUPRC_std": float("nan"), "fold_AUPRCs": [],
        }, info

    K_start = min(int(policy.K_TARGET), n_pos_groups)
    if K_start < 2:
        info["reason"] = "K_start < 2 (insufficient positive site-groups)."
        return {
            "AUROC_mean": float("nan"), "AUROC_std": float("nan"), "fold_AUROCs": [],
            "AUPRC_mean": float("nan"), "AUPRC_std": float("nan"), "fold_AUPRCs": [],
        }, info

    for K in range(K_start, int(policy.K_MIN) - 1, -1):
        gkf = GroupKFold(n_splits=K)
        aurocs: List[float] = []
        auprcs: List[float] = []
        total_folds = 0
        skipped_train = 0
        skipped_test = 0

        for train_idx, test_idx in gkf.split(X, y, groups):
            total_folds += 1
            y_train = y[train_idx]
            y_test = y[test_idx]

            if np.unique(y_train).size < 2:
                skipped_train += 1
                continue
            if np.unique(y_test).size < 2:
                skipped_test += 1
                continue

            imputer = SimpleImputer(strategy=policy.IMPUTER_STRATEGY)
            X_train_imp = imputer.fit_transform(X[train_idx])
            X_test_imp = imputer.transform(X[test_idx])

            clf = RandomForestClassifier(
                n_estimators=int(policy.RF_N_ESTIMATORS),
                class_weight="balanced",
                random_state=int(policy.RF_RANDOM_STATE),
                n_jobs=int(policy.RF_N_JOBS),
            )
            clf.fit(X_train_imp, y_train)
            y_prob = clf.predict_proba(X_test_imp)[:, 1]

            aurocs.append(float(roc_auc_score(y_test, y_prob)))
            auprcs.append(float(average_precision_score(y_test, y_prob)))

        info["K_used"] = int(K)
        info["n_total_folds"] = int(total_folds)
        info["n_valid_folds"] = int(len(aurocs))
        info["n_skipped_train_single_class"] = int(skipped_train)
        info["n_skipped_test_single_class"] = int(skipped_test)

        if len(aurocs) >= int(policy.MIN_VALID_FOLDS):
            info["reason"] = "OK"
            return {
                "AUROC_mean": float(np.mean(aurocs)),
                "AUROC_std": float(np.std(aurocs)),
                "fold_AUROCs": [float(x) for x in aurocs],
                "AUPRC_mean": float(np.mean(auprcs)),
                "AUPRC_std": float(np.std(auprcs)),
                "fold_AUPRCs": [float(x) for x in auprcs],
            }, info

    info["reason"] = "Insufficient valid folds even after decreasing K."
    return {
        "AUROC_mean": float("nan"), "AUROC_std": float("nan"), "fold_AUROCs": [],
        "AUPRC_mean": float("nan"), "AUPRC_std": float("nan"), "fold_AUPRCs": [],
    }, info


# ==============================================================================
# 13) CaseSpec + grouping helpers
# ==============================================================================

@dataclass
class CaseSpec:
    name: str
    unbound_structure_path: str
    bound_structure_path: str

    bound_positive_residue_ids: Optional[List[str]] = None

    chain_hints: Dict[str, str] = field(default_factory=dict)
    protein_only: bool = True

    symmetry_policy: SymmetryPolicy = field(default_factory=SymmetryPolicy)
    collapse_only_within_symmetry_groups: bool = True

    chain_assign_policy: ChainAssignPolicy = field(default_factory=ChainAssignPolicy)

    remove_solvent: bool = True
    solvent_codes: List[str] = field(default_factory=lambda: ["HOH", "WAT", "TIP3", "DOD"])

    interface_policy: InterfaceLabelPolicy = field(default_factory=InterfaceLabelPolicy)
    policy_p: PolicyP = field(default_factory=PolicyP)


def _parse_rid(rid: str) -> Tuple[str, str]:
    try:
        c, right = rid.split(":", 1)
        return c, right
    except Exception:
        return "?", rid


# ==============================================================================
# 14) BenchmarkSuite (ENGINE FROM USER_ENGINE_CODE)
# ==============================================================================

class BenchmarkSuite:
    def __init__(self, workdir: str, cache_dirname: str = "_suite_cache"):
        self.workdir = os.path.abspath(workdir)
        self.cache_dir = os.path.join(self.workdir, cache_dirname)
        _ensure_dir(self.workdir)
        _ensure_dir(self.cache_dir)

    def resolve_case_paths(self, case: CaseSpec) -> CaseSpec:
        u = _resolve_path_or_download(case.unbound_structure_path, cache_dir=self.cache_dir, prefer_ext=".cif")
        b = _resolve_path_or_download(case.bound_structure_path, cache_dir=self.cache_dir, prefer_ext=".cif")
        case.unbound_structure_path = u
        case.bound_structure_path = b
        return case

    def run_engine_unbound(self, unbound_path: str, out_dir: str, engine_args: Optional[List[str]] = None):
        """
        Engine stage for each case:
          - If Ep_sim.npy / Ep_manifest.json already exist in out_dir, reuse them.
          - Otherwise, materialize USER_ENGINE_CODE into _embedded_engine.py and call it.
        """
        _ensure_dir(out_dir)
        ep_path = os.path.join(out_dir, "Ep_sim.npy")
        mani_path = os.path.join(out_dir, "Ep_manifest.json")

        if os.path.exists(ep_path) and os.path.exists(mani_path):
            print(f"[SKIP] Engine(unbound): using precomputed Ep_sim / Ep_manifest in {out_dir}")
            return

        engine_path = _materialize_engine(self.workdir)

        # Default engine options: no pair-contacts, vector mode
        effective_args = ["--no-pair-contacts", "--gpu-density-mode", "vector"]
        if engine_args is not None:
            effective_args = list(engine_args)

        # Legacy compatibility: older wrappers may pass "--vector-mode" (not supported by the embedded engine).
        # Map it to "--gpu-density-mode vector".
        if "--vector-mode" in effective_args:
            # remove all occurrences
            effective_args = [a for a in effective_args if a != "--vector-mode"]
            # only add if not already specified
            if "--gpu-density-mode" not in effective_args:
                effective_args += ["--gpu-density-mode", "vector"]

        cmd = [
            sys.executable,
            engine_path,
            unbound_path,
            "--out-dir", out_dir,
        ] + effective_args

        print("[RUN] Engine(unbound):", " ".join(cmd))
        subprocess.run(cmd, check=True)

    def _build_groups_safely(self, unbound_structure_path: str, rids_unbound: List[str], case: CaseSpec) -> Tuple[np.ndarray, Dict[str, Any]]:
        unbound_chains = _extract_chain_sequences(unbound_structure_path, protein_only=case.protein_only)
        sym_map = detect_symmetry_groups(unbound_chains, case.symmetry_policy)

        groups = []
        for rid in rids_unbound:
            c, right = _parse_rid(rid)
            if case.collapse_only_within_symmetry_groups:
                gid = sym_map.get(c, None)
                if gid is None:
                    groups.append(f"SITE_{c}_{right}")
                else:
                    groups.append(f"SITE_G{gid}_{right}")
            else:
                groups.append(f"SITE_{c}_{right}")

        report = {
            "symmetry_groups": sym_map,
            "symmetry_policy": {
                "min_identity": case.symmetry_policy.min_identity,
                "min_coverage": case.symmetry_policy.min_coverage,
            },
            "collapse_only_within_symmetry_groups": bool(case.collapse_only_within_symmetry_groups),
        }
        return np.array(groups, dtype=object), report

    def evaluate_policy_p(
        self,
        unbound_out_dir: str,
        case: CaseSpec,
        b2u: Dict[str, str],
        map_report_extra: Dict[str, Any],
        bound_positive_rids_used: List[str],
        bound_interface_report: Dict[str, Any],
    ) -> Dict[str, Any]:
        ep_path = os.path.join(unbound_out_dir, "Ep_sim.npy")
        mani_path = os.path.join(unbound_out_dir, "Ep_manifest.json")
        if not os.path.exists(ep_path) or not os.path.exists(mani_path):
            return {"case": case.name, "error": "engine_outputs_missing",
                    "missing": [p for p in [ep_path, mani_path] if not os.path.exists(p)]}

        X = np.load(ep_path)
        mani = _read_json(mani_path)
        rids_unbound = mani.get("residue_ids", [])
        if not isinstance(rids_unbound, list) or len(rids_unbound) == 0:
            return {"case": case.name, "error": "empty_manifest_residue_ids"}

        solvent_report = {"solvent_filter_applied": False}
        if case.remove_solvent:
            meta_json_path = os.path.join(unbound_out_dir, "meta.json")
            X, rids_unbound, keep_mask, solvent_report = filter_solvent_from_engine_outputs(
                X=X,
                residue_ids=rids_unbound,
                solvent_codes=case.solvent_codes,
                unbound_structure_path=case.unbound_structure_path,
                meta_json_path=meta_json_path,
            )

        y, unbound_pos_rids, map_report = map_bound_labels_to_unbound_indices(
            bound_positive_rids=bound_positive_rids_used,
            bound_to_unbound_rid=b2u,
            unbound_manifest_rids=rids_unbound,
        )

        n_pos = int(y.sum())
        n_neg = int(len(y) - n_pos)
        if n_pos == 0 or n_neg == 0:
            return {
                "case": case.name,
                "error": "degenerate_labels_after_mapping",
                "n_pos": n_pos, "n_neg": n_neg,
                "bound_positive_residue_ids_used": bound_positive_rids_used,
                "bound_interface_report": bound_interface_report,
                "mapping": map_report,
                "mapping_extra": map_report_extra,
                "solvent": solvent_report,
            }

        groups, sym_report = self._build_groups_safely(case.unbound_structure_path, rids_unbound, case)

        metrics, info = eval_metrics_grouped_policy_p(
            X=X, y=y, groups=groups, policy=case.policy_p
        )

        label_out_path = os.path.join(os.path.dirname(unbound_out_dir), "unbound_positive_residue_ids.json")
        _write_json(label_out_path, {"unbound_positive_residue_ids": unbound_pos_rids})

        return {
            "case": case.name,
            **metrics,
            "policyP_info": info,
            "n_pos": n_pos,
            "n_neg": n_neg,
            "unbound_positive_residue_ids": unbound_pos_rids,
            "unbound_positive_residue_ids_path": label_out_path,
            "bound_positive_residue_ids_used": bound_positive_rids_used,
            "bound_interface_report": bound_interface_report,
            "mapping": map_report,
            "mapping_extra": map_report_extra,
            "symmetry": sym_report,
            "solvent": solvent_report,
            "unbound_out_dir": unbound_out_dir,
            "unbound_structure": case.unbound_structure_path,
            "bound_structure": case.bound_structure_path,
        }

    def run_case_bound_labels_unbound_features(self, case: CaseSpec, engine_args: Optional[List[str]] = None) -> Dict[str, Any]:
        case = self.resolve_case_paths(case)
        case_dir = os.path.join(self.workdir, case.name)
        out_unbound = os.path.join(case_dir, "out_unbound")
        _ensure_dir(case_dir)

        self.run_engine_unbound(case.unbound_structure_path, out_unbound, engine_args=engine_args)

        b2u, map_report_extra = build_bound_to_unbound_residue_map(
            bound_path=case.bound_structure_path,
            unbound_path=case.unbound_structure_path,
            chain_hints=case.chain_hints,
            protein_only=case.protein_only,
            assign_policy=case.chain_assign_policy,
        )

        chain_map = map_report_extra.get("bound_to_unbound_chain_map", {})
        target_bound_chains = set(chain_map.keys())

        bound_interface_report = {"auto_interface_labels": False}
        if case.bound_positive_residue_ids is None or len(case.bound_positive_residue_ids) == 0:
            if len(target_bound_chains) == 0:
                bound_positive_rids_used = []
                bound_interface_report = {
                    "auto_interface_labels": True,
                    "error": "no_target_bound_chains_matched_unbound",
                    "chain_map": chain_map,
                }
            else:
                bound_positive_rids_used, bound_interface_report = infer_ppi_interface_labels_from_bound(
                    bound_structure_path=case.bound_structure_path,
                    target_bound_chains=target_bound_chains,
                    policy=case.interface_policy,
                )
                bound_interface_report["auto_interface_labels"] = True
        else:
            bound_positive_rids_used = list(case.bound_positive_residue_ids)
            bound_interface_report = {
                "auto_interface_labels": False,
                "n_bound_labels_user": int(len(bound_positive_rids_used)),
            }

        result = self.evaluate_policy_p(
            unbound_out_dir=out_unbound,
            case=case,
            b2u=b2u,
            map_report_extra=map_report_extra,
            bound_positive_rids_used=bound_positive_rids_used,
            bound_interface_report=bound_interface_report,
        )

        _write_json(os.path.join(case_dir, "result_bound_labels_unbound_features.json"), result)
        return result

    def run_suite(self, cases: List[CaseSpec], engine_args: Optional[List[str]] = None) -> Dict[str, Any]:
        results = []
        for c in cases:
            res = self.run_case_bound_labels_unbound_features(c, engine_args=engine_args)
            results.append(res)

        aurocs = [r.get("AUROC_mean") for r in results
                  if isinstance(r.get("AUROC_mean"), (int, float)) and not np.isnan(r.get("AUROC_mean"))]
        auprcs = [r.get("AUPRC_mean") for r in results
                  if isinstance(r.get("AUPRC_mean"), (int, float)) and not np.isnan(r.get("AUPRC_mean"))]

        summary = {
            "n_cases": int(len(results)),
            "n_cases_with_valid_AUROC": int(len(aurocs)),
            "n_cases_with_valid_AUPRC": int(len(auprcs)),
            "suite_AUROC_mean_over_cases": float(np.mean(aurocs)) if len(aurocs) > 0 else float("nan"),
            "suite_AUPRC_mean_over_cases": float(np.mean(auprcs)) if len(auprcs) > 0 else float("nan"),
        }
        out = {"run_environment": _collect_run_environment({"workdir": self.workdir}), "summary": summary, "results": results}
        _write_json(os.path.join(self.workdir, "suite_results.json"), out)
        return out


# ==============================================================================
# 15) BUILD DB5.5-LIKE CASES FROM DIRECTORY
# ==============================================================================

def build_cases_from_db55_like_dir(root: str, name_prefix: str = "DB55") -> List[CaseSpec]:
    pat = re.compile(r"^([0-9A-Za-z]{4})_([lr])_([bu])\.pdb$")
    struct_dirs: Dict[str, Dict[str, str]] = {}
    for r, _, files in os.walk(root):
        for fn in files:
            m = pat.match(fn)
            if not m:
                continue
            cid, lr, bu = m.group(1), m.group(2), m.group(3)
            struct_dirs.setdefault(cid, {})[f"{lr}_{bu}"] = os.path.join(r, fn)

    cases: List[CaseSpec] = []
    for cid, d in sorted(struct_dirs.items()):
        need = ["l_b", "l_u", "r_b", "r_u"]
        if not all(k in d for k in need):
            continue

        lb = d["l_b"]
        rb = d["r_b"]
        out_dir = os.path.dirname(lb)
        bound_complex = os.path.join(out_dir, f"{cid}_bound_complex.pdb")
        if not os.path.exists(bound_complex):
            try:
                make_bound_complex(lb, rb, bound_complex)
            except Exception as e:
                print(f"[WARN] Failed to make bound complex for {cid}: {e}")
                continue

        cases.append(CaseSpec(
            name=f"{name_prefix}_{cid}_L",
            unbound_structure_path=d["l_u"],
            bound_structure_path=bound_complex,
            bound_positive_residue_ids=None,
        ))
        cases.append(CaseSpec(
            name=f"{name_prefix}_{cid}_R",
            unbound_structure_path=d["r_u"],
            bound_structure_path=bound_complex,
            bound_positive_residue_ids=None,
        ))
    return cases


# ==============================================================================
# 16) MAIN
# ==============================================================================

def _dataset_unpacked_has_db55_cases(dataset_dir: str) -> bool:
    """Return True if dataset_dir contains DB5.5-style pdb filenames like XXXX_l_u.pdb."""
    if not os.path.isdir(dataset_dir):
        return False
    pat = re.compile(r"^[0-9A-Za-z]{4}_[lr]_[bu]\.pdb$")
    for root, _, files in os.walk(dataset_dir):
        for fn in files:
            if pat.match(fn):
                return True
    return False


if __name__ == "__main__":
    _set_reproducible(42)

    import argparse

    parser = argparse.ArgumentParser(
        description="Universal Protein Benchmark runner (supports colab_all.zip OR DB5.5 .tgz only)."
    )
    parser.add_argument(
        "--tgz",
        type=str,
        default="benchmark5.5.tgz",
        help="Path to DB5.5-style tgz (default: benchmark5.5.tgz).",
    )
    parser.add_argument(
        "--colab_zip",
        type=str,
        default="colab_all.zip",
        help="Path to colab_all.zip (default: colab_all.zip). If present, it is preferred.",
    )
    parser.add_argument(
        "--workdir",
        type=str,
        default="benchmark_workdir",
        help="Working directory (default: benchmark_workdir).",
    )
    parser.add_argument(
        "--block-size",
        type=int,
        default=1024,
        help="Engine block size (number of residues per streaming block). Default: 1024.",
    )
    args = parser.parse_args()

    WORKDIR = args.workdir
    os.makedirs(WORKDIR, exist_ok=True)
    DATASET_UNPACK_DIR = os.path.join(WORKDIR, "_dataset_unpacked")
    os.makedirs(DATASET_UNPACK_DIR, exist_ok=True)


    # If _dataset_unpacked already exists with DB5.5-style files, skip restore/unpack and go straight to evaluation.
    if _dataset_unpacked_has_db55_cases(DATASET_UNPACK_DIR):
        print(f"[INFO] Found existing dataset in {DATASET_UNPACK_DIR} -> skipping restore/unpack and starting evaluation.")
    else:
        # Prefer colab_all.zip if available (backward compatible)
        if os.path.exists(args.colab_zip):
            print(f"[INFO] Found {args.colab_zip} -> restoring legacy workdir.")
            restore_from_colab_all(args.colab_zip)
            if not os.path.isdir(WORKDIR):
                raise FileNotFoundError(
                    "benchmark_workdir directory was not found after restoring colab_all.zip.\n"
                    "Make sure colab_all.zip contains the full previous work folder."
                )
            if not os.path.isdir(DATASET_UNPACK_DIR):
                raise FileNotFoundError(
                    "benchmark_workdir/_dataset_unpacked was not found after restoring colab_all.zip.\n"
                    "Make sure your archive contains the unpacked dataset directory."
                )
        else:
            # TGZ-only mode
            if not os.path.exists(args.tgz):
                raise FileNotFoundError(
                    f"Neither {args.colab_zip} nor {args.tgz} was found.\n"
                    "Put benchmark5.5.tgz next to this script, or pass --tgz PATH."
                )
            print(f"[INFO] colab_all.zip not found -> using TGZ-only mode with {args.tgz}")
            restore_from_db55_tgz(args.tgz, dataset_unpack_dir=DATASET_UNPACK_DIR)

    suite = BenchmarkSuite(workdir=WORKDIR)
    cases = build_cases_from_db55_like_dir(DATASET_UNPACK_DIR, name_prefix="DB55")
    if len(cases) == 0:
        raise RuntimeError(
            "No DB5.5-like cases were found.\n"
            "Expected files like XXXX_l_u.pdb, XXXX_l_b.pdb, XXXX_r_u.pdb, XXXX_r_b.pdb."
        )
    print(f"[INFO] Built {len(cases)} cases from {DATASET_UNPACK_DIR}")

    # Default engine options preserved + block size controlled from CLI
    ENGINE_ARGS = ["--no-pair-contacts", "--gpu-density-mode", "vector", "--block-size", str(args.block_size)]
    out = suite.run_suite(cases, engine_args=ENGINE_ARGS)
    summary_path = os.path.join(WORKDIR, "suite_results.json")
    print("[DONE] suite_results.json written to:", summary_path)
    print(json.dumps(out["summary"], indent=2))
```
